# CURE-ating the substrate scope and functional residues of nonheme iron(II) α-ketoglutarate dependent hydroxylase BesE

**DOI:** 10.1101/2025.10.10.681649

**Authors:** Austin R. Hopiavuori, Beau S. Andre, Emely D. Avalos, Surina L. Beal, George R. Beck, Charles J. Choi, Elizabeth M. Dolzhansky, Michelle X. Du, Isabella S. Gallardo, Cyrus J. Ghorbani, Kyle M. Hinaga, Analynn T. Nguyen, Edward H. T. Pham, Shaun M. K. McKinnie

## Abstract

Nonheme iron (II) α-ketoglutarate-dependent dioxygenases (Fe/αKGs) play important roles in functionalizing biological substrates from individual amino acids up to macromolecules. BesE, a unique homolog from the actinobacterial β-ethynylserine biosynthetic pathway, catalyzes a highly selective hydroxylation on dipeptide substrate γ-L-glutamyl-L-propargylglycine (**1**). Inspired by this transformation, our year-long Course-based Undergraduate Research Experience (CURE) laboratory interrogated BesE catalysis using an interdisciplinary approach. After establishing a modular chemical synthesis of **1**, we rationally designed fourteen non-native analogs with key alterations to specific substrate moieties putatively involved in enzymatic recognition. Following *in vitro* enzymology and the application of chemical derivatization techniques compatible with all substrate analogs and putative products, we determined that the terminal alkyne moiety is not essential while reinforcing the significance of the γ-glutamyl moiety for hydroxylation activity. Thirteen rationally designed BesE mutants established the importance of polar active site residues for substrate recognition and catalysis. This work establishes a baseline of BesE recognition from both a chemical and biochemical perspective and contributes to the growing understanding about Fe/αKG recognition on biomedically relevant targets. Moreover, this contributes to the growing examples of using natural products and their biosynthetic enzymology as a vibrant platform for the interdisciplinary training of early career biomedical researchers.

**Graphical Abstract:** 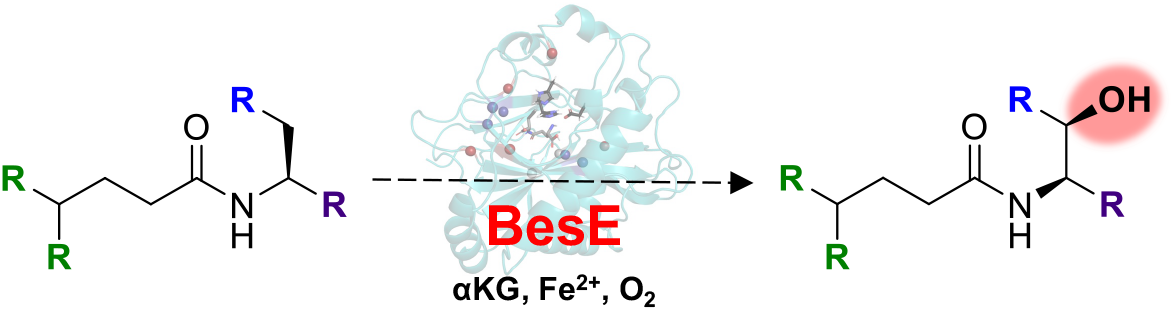

## Introduction

Nonheme iron(II) α-ketoglutarate dependent dioxygenases (Fe/αKG) are a diverse family of enzymes that perform important roles in both primary and secondary metabolism.^1^ Within primary metabolism, these enzymes are involved in the modification of small molecules up to macromolecules and play important physiological regulatory roles, including oxygen-sensing, stabilizing structural proteins like collagen, and modifying nucleic acid bases.^2^ Fe/αKG enzymes exhibit expanded chemical reactivity within secondary (or specialized) metabolism, and contribute broadly to the diversity observed within natural product biosynthesis.^3,4^ Through a conserved catalytic cycle involving co-factor iron (II), and co-substrates α-ketoglutarate and molecular oxygen, Fe/αKGs generate a high valency Fe(IV) oxo species capable of radical hydrogen abstraction, typically from an unactivated, aliphatic C-H bond.^5^ Following substrate radical formation, native or engineered Fe/αKGs can perform fascinating and synthetically useful reactions, including: halogenation, epimerization, desaturation, endoperoxidation, carbon-nitrogen, and carbon-carbon bond formations.^3,6,7^ However the most common reaction pathway involves radical rebound from the Fe(III)-OH intermediate to form hydroxylated products with high regio- and stereoselectivity. We have been interested in understanding how this family of oxidative enzymes modulates substrate hydroxylation and other chemistries within natural product biosynthetic and focused biocatalytic directions.^8^

We were particularly inspired by the biosynthetic pathway of known antimetabolite β-ethynylserine (βes) from *Streptomyces cattleya*, discovered and fully elucidated by the Chang group in 2019 (**Figures 1A, S1**).^9^ Through the action of three biosynthetic enzymes (BesBCD), L-lysine is transformed into the terminal alkyne-containing L-propargylglycine (L-Pra). Subsequent ligation of this non-canonical amino acid (ncAA) to the γ-carboxylate of L-glutamic acid by ATP-dependent ligase BesA formed dipeptide γ-L-glutamyl-L-propargylglycine (γ-L-Glu-L-Pra, **1**). BesE performs the final stereoselective β-hydroxylation to generate γ-L-glutamyl-L-β-ethynylserine (γ-L-Glu-L-βes, **2**). Two Fe/αKGs with diverging chemistries are highlighted within **2** biosynthesis: BesD initiates the biosynthetic pathway by performing a stereoselective γ-chlorination on L-lysine; whereas BesE catalyzes a conventional hydroxylation reaction to create the threonine-mimicking antimetabolite pharmacophore.^10^ Considerable mechanistic, structural, and protein engineering efforts have provided insight into BesD halogenation catalysis and how it diverges from conventional L-lysine hydroxylation.^11,12^ We were instead interested in the unusual features associated with the more canonical hydroxylase BesE. As noted in its original characterization, this Fe/αKG did not appreciably hydroxylate L-Pra *in vitro*, suggesting an important role for the γ-L-glutamyl moiety.^9^ Moreover, the ability of BesE to catalyze radical hydrogen abstraction without rearrangement of the adjacent alkyne moiety was unusual from a chemical reactivity perspective (**Figure S2**). BesE itself was characterized via a coupled assay during its initial discovery^9^ and subsequent cell-free lysate assays,^13^ thus providing an opportunity to specifically interrogate it individually. And given the structural similarity of βes mimicking L-threonine, we were curious if native or engineered BesE could produce other non-native antimetabolites in appreciable yields.

**Figure 1.**
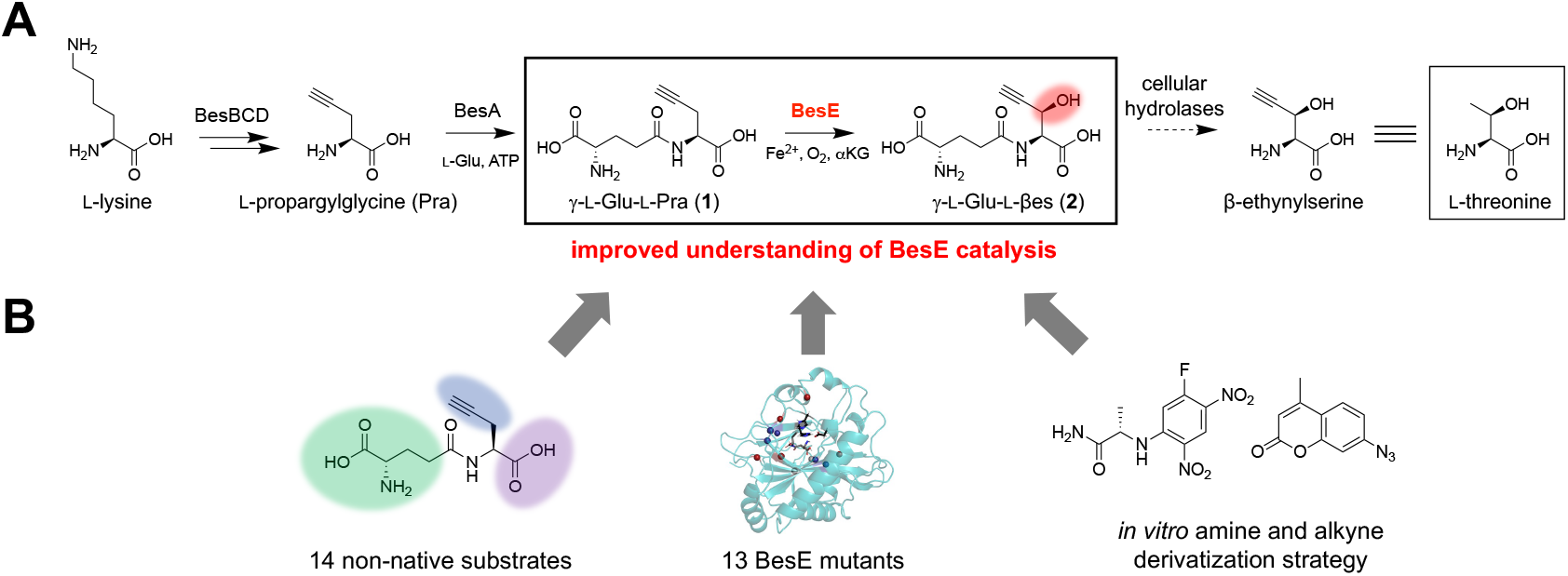
**A)** BesE catalyzed β-hydroxylation of γ -L-glutamyl-L-propargylglycine (γ-L-Glu-L-Pra, **1**) to form γ - L-glutamyl-L-β-ethynylserine (γ -L-Glu-L-βes, **2**) in the context of the β-ethynylserine biosynthetic pathway. The structure of L-threonine is shown for comparison to the related antimetabolite β-ethynylserine. **B)** General project overview for our Course-based Undergraduate Research Experience to interrogate an improved chemical, biochemical, and analytical understanding of BesE catalysis.

Through this study, our year-long Course-based Undergraduate Research Experience (CURE) lab of twelve undergraduate researchers investigated the features necessary for BesE catalysis using both a chemical and biochemical approach (**Figure 1B**). After developing a modular four-step chemical synthesis of **1**, we extended this strategy to 14 non-native analogs to assess the extent of substrate engineering. We also heterologously expressed and purified native BesE and generated 13 mutants to probe the biochemical features of hydroxylation catalysis. Moreover, we provided a robust analytical assay and chemical derivatization approach to detect and quantify the extent of conversion of multiple, polar, non-native substrates via *in vitro* BesE assays. These cumulative efforts provide an improved understanding of this unique and significant Fe/αKG hydroxylase homolog, but also for this broader enzyme family.

## Results

The chemical synthesis of native substrate **1** was initiated by an acid-catalyzed methyl ester formation of L-propargylglycine in 2,2-dimethoxypropane (**Figure 2A**). The resulting L-propargylglycine methyl ester hydrochloride salt (**3**) was subsequently coupled to the γ-carboxylic acid side chain of Boc-L-Glu-OtBu using hexafluorophosphate azabenzotriazole tetramethyl uronium (HATU) in the presence of triethylamine, isolating protected amide product **4** following silica flash column chromatography. Sequential methyl ester hydrolysis using aqueous lithium hydroxide followed by trifluoroacetic acid (TFA) deprotection of Boc- and *tert*-butyl ester groups afforded **1** as a TFA salt in appreciable yield.

**Figure 2.**
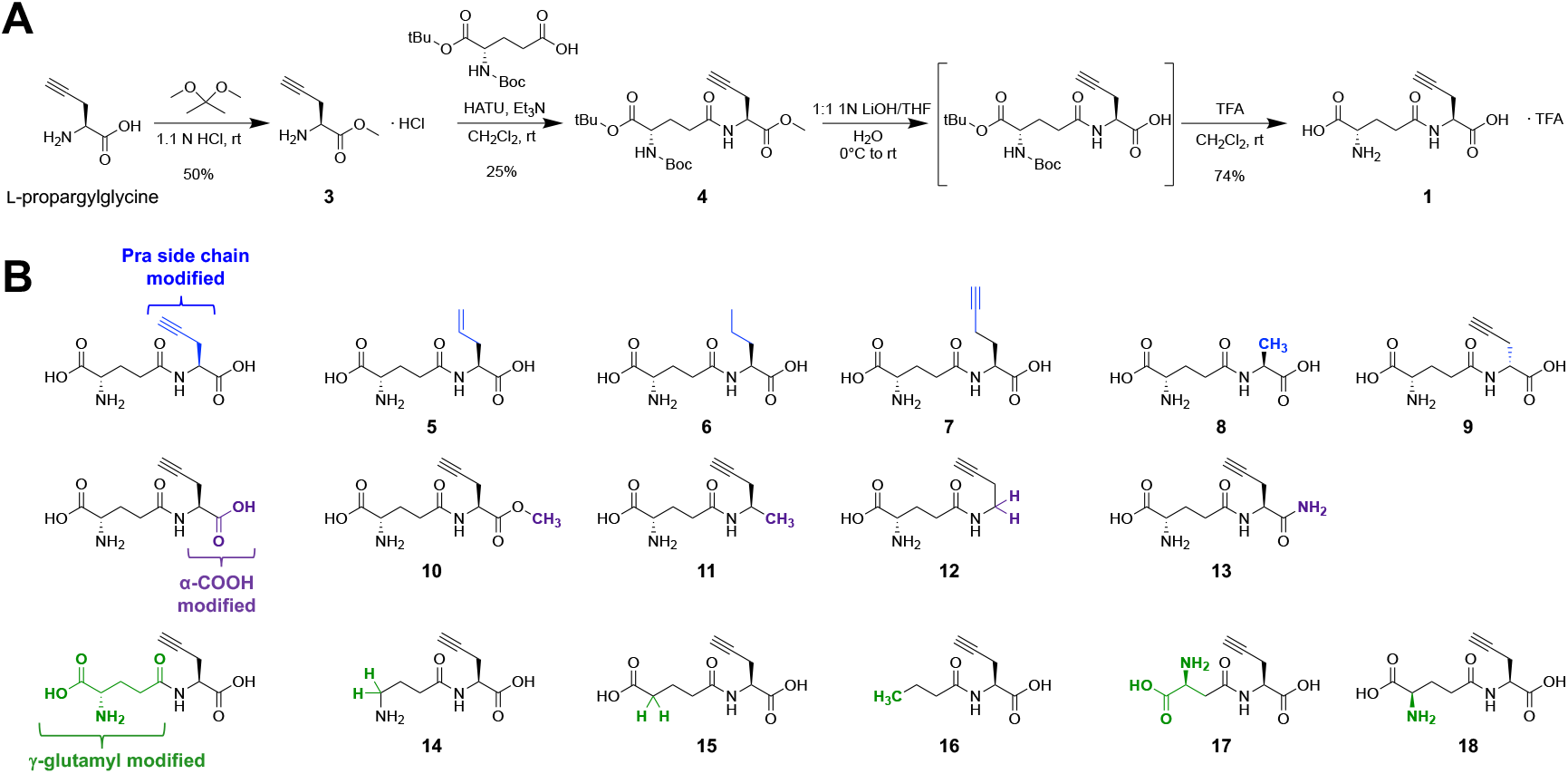
**A)** Modular four step synthesis of native BesE substrate **1. B)** Structures of non-native BesE substrate analogs (**5-18**) grouped by region of modification at either the: Pra side chain (blue); Pra α-carboxylic acid (purple); γ-glutamyl moiety (green).

Following the successful synthesis of **1**, a series of non-native analogs were designed with modifications to either the glutamic acid or propargylglycine moieties to assess the importance of key features for BesE catalysis and selectivity (**Figure 2B**). Alteration of the propargylglycine amino acid side chain generated five novel side chain modified analogs nearest the site of hydroxylation. Adapting the previous synthetic scheme to use either L-allylglycine (L-Alg) or L-norvaline (L-Nva) allowed the generation of substrate analogs with the same carbon chain length but either a terminal alkene (γ-L-Glu-L-Alg, **5**) or a linear saturated alkane (γ-L-Glu-L-Nva, **6**) respectively. Other amino acid starting materials were selected, beginning with L-homopropargylglycine (L-Hpg) to retain the terminal alkyne feature but extend it by one methylene unit (γ-L-Glu-L-Hpg, **7**), and a simplified L-alanine containing analog (γ-L-Glu-L-Ala, **8**) were created to assess hydroxylation activity lacking this unique chemical feature. The importance of the Pra L-stereocenter was investigated by adapting the same synthetic workflow with D-propargylglycine to generate epimerized analog (γ-L-Glu-D-Pra, **9**). Efforts to replace the terminal alkyne with a nitrile were stymied due to chemical incompatibility during the methyl esterification reaction on β-cyano-L-alanine (**Figure S3**), and this analog was not pursued further.

Another series of four analogs were developed to interrogate the significance of the L-Pra α-carboxylic acid for BesE recognition. Intermediate **4** was directly subjected to TFA deprotection, generating methyl esterified analog γ-L-Glu-L-Pra-OMe (**10**). Coupling Boc-L-Glu-OtBu with chiral amine (*R*)-pent-4-yn-2-amine (Pya), purification of the amide intermediate, and subsequent TFA deprotection enabled the synthesis of an analog **11** (γ-L-Glu-Pya, **11**) that replaced this moiety with an α-methyl group while retaining the desired propargyl group stereochemistry. Adapting a similar workflow beginning with but-3-yn-1-amine (Bya) generated analog **12** (γ-L-Glu-Bya) that lacked this C-terminal functionality altogether. Lastly, C-terminal amide analog γ-L-Glu-L-Pra-NH_2_ (**13**) was synthesized by Fmoc-solid phase peptide synthesis on Rink Amide resin by the following sequence: loading Fmoc-L-Pra-OH on the resin by HATU coupling; Fmoc-deprotection using piperidine; HATU coupling of Boc-L-Glu-OtBu; and global TFA deprotection and concomitant resin cleavage.

The last series of analogs were designed to modify chemical features on the γ-L-glutamic acid side of the substrate. Four putative substrates were synthesized by HATU coupling key intermediates to **3** then following necessary deprotection steps, resulting in the removal of: the α-carboxylic acid (GABA-L-Pra, **14**); the α-amine (glutarate-L-Pra, **15**); both α-moieties (But-L-Pra, **16**); or a side chain methylene (β-L-Asp-L-Pra, **17**). Lastly the same synthetic workflow to **1** was altered to use Boc-D-Glu-OtBu, generating epimerized analog (γ-D-Glu-L-Pra, **18**) to investigate the importance of the Glu stereocenter. With established substrate **1** and 14 novel analogs in place, we next focused on the biochemical and analytical aspects of BesE interrogation.

To generate our recombinant BesE enzyme, we synthesized the gene as an *E. coli* optimized construct and cloned it into a pET28a vector with an N-terminal hexahistidine tag. Recombinant BesE was expressed in *E. coli* BL21(DE3) cells following conventional IPTG-induction and purified to near homogeneity using Ni-NTA affinity chromatography, producing typical yields of 40 mg/L. Any co-purified iron(II) was removed by treatment with EDTA prior to size exclusion chromatography to minimize any potential aerobic BesE oxidation and putative inactivation, a strategy that we routinely use for other Fe/αKG homologs.^14^ *In vitro* BesE assays were established following adapted literature conditions from either the one pot BesAE^9^ or cell free protein synthesis^13^ publications and analyzed by ultra-high performance liquid chromatography-mass spectrometry (UPLC-MS). Initial assays with **1** were set up in 50 mM K_2_HPO_4_ pH 8.0 buffer with 4 mol% BesE and all necessary cofactors and co-substrates (FeSO_4_, αKG, L-ascorbate, and O_2_). Subtle optimization efforts revealed that increasing the concentrations of both L-ascorbate and FeSO_4_ improved the overall *in vitro* turnover of BesE based on relative extracted ion chromatogram (EIC) intensities (**Figures S4, S5**). Despite this optimization, complete *in vitro* consumption of **1** was not achieved, which was consistent with previous literature efforts.^9,13^ To improve the reproducibility of peak shapes, intensities, and retention times for polar analytes **1** and **2** following *in vitro* assays, we adopted two chemical derivatization methods (**Figure 3A**). Primary amines were derivatized with 2,4-dinitro-5-fluorophenyl-L-alanine amide (L-FDAA, Marfey’s reagent) following established conditions (**Figure S6**),^15,16^ and terminal alkyne-containing substrates were reacted via copper(I)-catalyzed azide alkyne cycloaddition (CuAAC) with coumarin azide (**Figure S7**).^9^ Importantly, at least one derivatization approach would be compatible with each of the synthetic non-native substrates and their putative hydroxylated products to improve chromatographic retention properties and analytical reproducibility.

**Figure 3.**
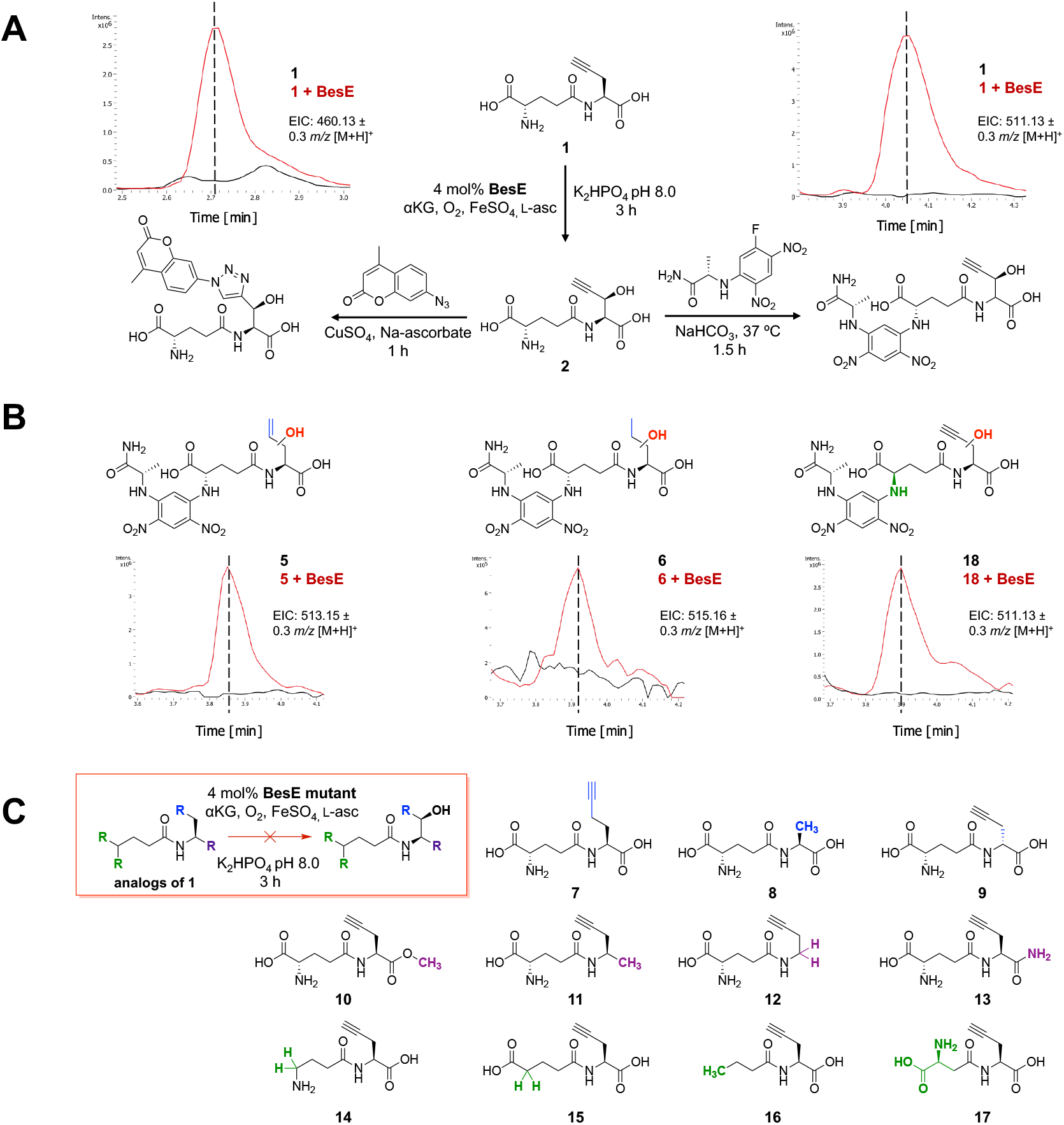
**A)** BesE *in vitro* enzyme assay with native substrate **1** and subsequent derivatization with either 7-azido-4-methylcoumarin (left) or Marfey’s reagent (right). Extracted ion chromatograms of derivatized β-hydroxylated product **2** for assays incubated in the presence (red) and absence (black) of purified BesE (EICs: 460.13; 511.13 ± 0.3 *m/z* [M+H]^+^ for 7-azido-4-methylcoumarin and Marfey derivatized products, respectively). **B)** Extracted ion chromatograms of Marfey derivatized hydroxylated substrate analogs **5, 6**, and **18**, for assays incubated in the presence (red) and absence (black) of purified BesE (EICs: 513.15; 515.16; 511.13 ± 0.3 *m/z* [M+H]^+^ respectively). **C)** Structures of substrate analogs which were not appreciably hydroxylated by BesE *in vitro*.

With *in vitro* assay and derivatization conditions in hand, we next looked to identify the scope of BesE hydroxylation biochemistry on non-native substrates. Each non-native substrate (**5 – 18**) was incubated with 4 mol% wild type BesE for three hours under optimized co-factor and co-substrate concentrations and an aerobic environment (**Figures S8 – S21**). BesE assays in the presence of **1**, and in the absence of enzyme corresponded to positive and negative controls, respectively. Following chemically appropriate derivatization and UPLC-MS analysis, we identified that three analogs (**5, 6**, and **18**) showed a significant [M+16] *m/z* feature that was absent in the no enzyme control (**Figures 3B, S8, S9, S21**). Substrates **5** and **6** had modifications directly near the native site of hydroxylation, with allyl- and *n*-propyl-amino acid side chains respectively. The epimerized γ-D-Glu-L-Pra derivative **18** also showed putative activity supportive of hydroxylation. Analogs that failed to be appreciably hydroxylated *in vitro* included: methylene extended **7**; simplified L-Ala derivative **8**; epimerized D-Pra-containing **9**; any alteration to the Pra carboxylate (**10 – 13**); and all other modifications to the γ-L-Glu moiety (**14 – 17**) (**Figures 3C, S10 – S20**). While modest from a substrate scope perspective, these experiments highlighted the strong preference of BesE for its native substrate **1**, and underscored the importance of the γ-Glu moiety for enzymatic recognition and to direct hydroxylation chemistry.

After assessing the extent of hydroxylation chemistry on non-native substrates, we next examined the biochemical features that are significant for BesE catalysis. In the absence of a crystal structure, we used an AlphaFold 3 model^17^ of BesE, generated a structural model of **1** using Avogadro,^18^ and docked the substrate into the putative active site using AutoDock.^19^ Using the 25 lowest energy docked structures, and filtering for orientations that positioned the L-Pra β-hydrogen in a productive position (12 of 25 structures), we hypothesized key BesE amino acid residues that could be contributing to substrate recognition and catalysis. Sequence and structural alignment identified the conserved catalytic active site triad (H153, D155, H238) involved in binding iron (II) for oxidative chemistry. While it is difficult to precisely predict where the organic substrate binds, especially given the dynamics associated with Fe/αKG catalysis, this enabled us to generate testable hypotheses to probe BesE substrate recognition. A total of thirteen residues were selected for substitution based on their proximity (within 6 Å) to docked **1** (**Figures 4A, S22**) and sequence conservation in other putative actinobacterial *besE* homologs (**Figure S23**). The two facial triad histidines (H153, H238) were left untouched, however D155 was selected given that its substitution contributes to halide coordination in Fe/αKG halogenase homologs.^20^ Other selected residues were broadly categorized into putative hydrophobic pocket shaping (M70, V146, A175, F232, V240, F255), or hydrogen bonding residues (R144, T150, Q173, N177, Y207, R249). Most selected residues were mutated to alanine with the exception of A175W, which was designed to assess if a bulkier side chain impeded **1** hydroxylation activity.

**Figure 4.**
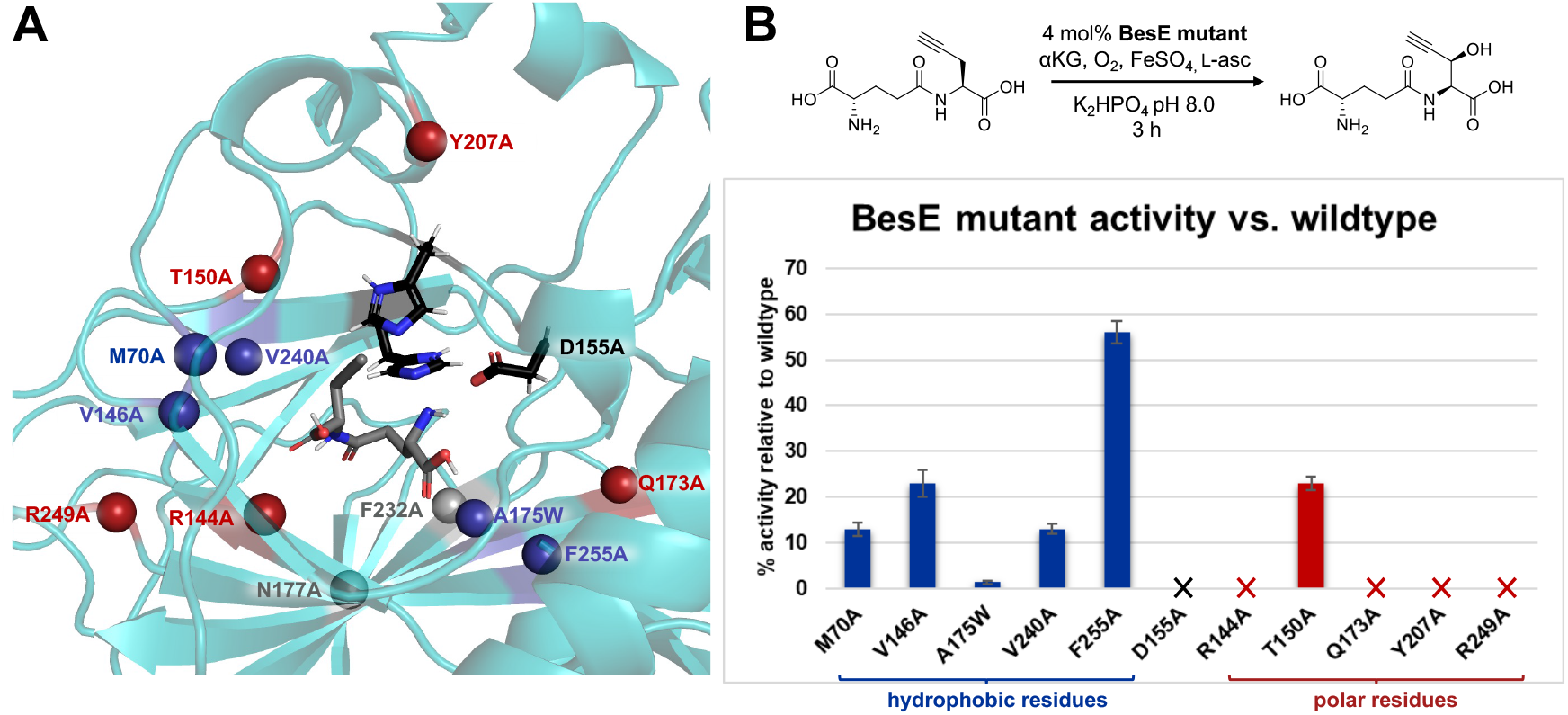
**A)** Depiction of the BesE AlphaFold 3 model with mutant residues assayed shown as spheres. Mutation sites of hydrophobic residues are shown in blue; mutation sites of polar residues are shown in red. Mutations N177A and F232A which were produced in lower titers and were not able to be assayed are shown in grey. The catalytic triad residues (black) and docked substrate **1** (grey) are shown as sticks. **B)** Bar graph displaying *in vitro* mutant activity in comparison to wild type BesE. Mutants are grouped and colored by hydrophobic (blue) and polar side chains (red) in the same manner as above. Variants that did not show any conversion to product are indicated with an ‘X’. Assays were run in quadruplicate (n = 4) for each BesE mutant.

Following successful site-directed mutagenesis (**Table S1**) and sequence confirmation, BesE mutants were transformed into *E. coli* BL21(DE3) cells and successfully heterologously expressed and purified following analogous protocols. All thirteen purified BesE variants were isolated in >90% purity by SDS-PAGE (**Figure S24**). However, mutants N177A and F232A were isolated in lower concentrations (< 5 μM), showed poor solubility, precipitated during efforts to further concentrate them, and were excluded from being functionally assessed *in vitro*. The remaining eleven BesE variants were incubated with **1** in quadruplicate following previously established *in vitro* assay conditions, L-FDAA derivatization, and analysis by UPLC-MS. The relative production of derivatized **2** was standardized to an internal chloramphenicol control to correct for deviations in ionization efficiency and compared to wild type BesE activity (**Figure 4B**). While all mutants showed reduced *in vitro* **1** hydroxylation activity (**Figures S25 – S35**), an interesting trend was uncovered. Alanine substitution of hydrophobic pocket shaping residues retained some **2** production, with F255A showing the highest relative activity (56%) and a continued decrease across V146A, M70A, and V240A BesE variants. These indicate that these residues are important for efficient substrate recognition but not essential for catalysis. The A175W mutant had detectable hydroxylation activity but at ∼1% wild type conversion. This suggested that this normally small residue is helping to shape a critical portion of the substrate bonding pocket and putatively occludes **1** from constructive binding. In contrast, interrogating alanine mutations of residues with polar side chains nearly abolished **1** hydroxylation activity. T150A was the only variant from this second group of mutants that retained any *in vitro* activity (23% wild type), potentially due to its more distant position from the predicted active site. All other polar side chain BesE mutants (R144A, D155A, Q173A, Y207A, R249A) showed a complete loss of **2** production, indicating their importance in **1** recognition via hydrogen bonding or ion pairing interactions. We assessed D155A for its halogenation potential on **1** by extracting for a putative chlorinated product *m/z* value, however no appreciable signal was observed (**Figure S36**). This corroborates multiple experimental and computational reports that additional engineering efforts are needed in on top of mutating the facial triad aspartic acid to enable efficient Fe/αKG halogenase activity.^11,12,20–22^ While none of our single point mutations improved *in vitro* BesE activity, they provided valuable functional insight into the significance of key hydrogen bonding and pocket shaping residues for efficient **1** hydroxylation catalysis. Moreover, they highlight the increasing utility of using structural prediction models to form experimentally testable hypotheses.

## Discussion

Overall, our study identified the extent of *in vitro* BesE hydroxylation catalysis on a non-native substrate library and suggested putative structural features important for **2** production. These focused substrates helped identify that BesE has limited tolerance for alteration at key regions of native substrate **1**. None of our analogs with substitutions to the propargylglycine α-carboxylic acid (**10 – 13**) were appreciably hydroxylated by BesE *in vitro*. While epimeric γ-D-Glu analog **18** showed some hydroxylation, negligible activity was observed on other γ-glutamyl modified analogs **14 – 17**. This highlighted that the presence of both α-amine and α-carboxylate moieties on the glutamyl moiety are essential for BesE recognition. Moreover, the absence of a putative hydroxylated product on the methylene-truncated β-L-Asp-L-Pra **17** indicates that the chain length of the glutamyl moiety is important. These results complement the BesE mutant assays where the alanine substitution of polar residues in the putative substrate binding pocket abolishes efficient **1** hydroxylation. While prediction in the absence of structural data is hypothetical, these cumulative data suggest there are important recognition elements, potentially hydrogen bonding or ionic interactions, that must be satisfied on both the substrate and enzyme end to afford this level of specificity. This further corroborates the initial report that L-Pra itself does not serve as a BesE substrate, underscoring the importance of this conserved γ-glutamylation step by BesA prior to stereoselective hydroxylation. The addition of chemical handles to standalone amino acid substrates prior to Fe/αKG reactions has been previously observed for the SadA amino acid β-hydroxylase,^23^ which showed promiscuity across a variety of *N*-succinylated hydrophobic amino acids and has served as a vibrant platform for further biocatalytic diversification.^24,25^ The presence of a dedicated hydrolase to liberate the hydroxylated product^26^ does not seem to be observed within the *bes* BGC but could speak to additional recognition, self-protection, or assisting with export of the precursor to the βes antimetabolite.

Intriguingly, substrate modifications near the site of hydroxylation occurs were tolerated provided the side chain carbon length was conserved. Both alkene (**5**) and alkane (**6**) derivatives showed reproducible hydroxylation albeit with a relative decrease in conversion compared to native substrate. Although slightly counterintuitive, this indicates that the terminal alkyne functionality itself is not essential for BesE recognition and catalysis. However, the absence of hydroxylated products for methylene extended analog **7**, alanine-derived analog **8**, and epimerized analog **9**, show the limitations of additional side chain modification. While mutation of hydrophobic BesE residues retained some level of *in vitro* **2** production, both valine mutations V146A and V240A displayed substantially reduced activity despite being two of the more conservative mutations screened in this study. Molecular docking suggested that these residues flank a hydrophobic pocket near the catalytic site that could be significant for properly orienting the propargyl side chain. Additional targeted mutagenesis fueled by structural insight would provide further credibility to this hypothesis. While low *in vitro* turnover prevented the isolation of hydroxylated **5** and **6** products, efforts to structurally characterize them and assess their ability to serve as antimetabolites in other biological systems are underway.

## Conclusion

Our interdisciplinary investigation into BesE catalysis provides insight into necessary features on both the substrate and macromolecular levels. Terminal alkyne containing scaffolds are relatively scarce in Nature and, when present, are typically observed at the end of fatty acid chains.^27,28^ The ability of this unique Fe/αKG homolog to selectively modify alkyne-containing amino acids generates additional tools for their application in natural product and biorthogonal labeling contexts.^29^ While this work provides an initial baseline into the biocatalytic capacity of BesE, building upon these results with experimentally validated structures and expanded mutant and substrate libraries would comprehensively identify its potential applications for generating hydroxylated terminal alkyne-containing amino acids and peptides. Additional efforts via directed evolution^30,31^ and high-throughput screening methods^32^ would rapidly diversify the testable BesE variants in relation to this goal. Non-heme iron enzymes have proven to have broad utility in both biocatalysis and chemoenzymatic synthesis;^33–36^ moreover, a remarkable array of inspiring transformations have also been observed from alteration of Fe/αKGs on complex meroterpenoid^37^ and tropolone^38^ substrates. An improved understanding of their fundamental biochemical and chemical scope features empowers these impactful applications on diverse non-native substrates. The application of multiple interdisciplinary technical skills while encouraging individual goal setting and completion on a focused research project creates a vibrant platform for the training of early career biomedical researchers (**Table S2**). Moreover, this study adds to the growing examples of leveraging natural products and their associated metabolomics, genomics, molecular biology, protein biochemistry, and biosynthetic enzymology in CURE or comparable upper division laboratory settings.^39–42^

## Supporting information

BesE-CURE-SI-PDF

## Acknowledgements

This work was supported by the University of California, Santa Cruz Physical and Biological Sciences division (CURE lab start up funding) and the National Institutes of Health (R21-GM-148870, R35-GM-147235) for general McKinnie lab work in the field of non-heme iron enzymology and other biosynthetic metalloenzymes respectively. We gratefully acknowledge S. Rubin, F. Pavlovici, P. Ngoi, and M. Membreño for additional biochemical support and supervision, H.-W. Lee for maintenance of nuclear magnetic resonance spectroscopy facilities, and R. Dunkin for support conceptualizing CURE lab assessments and pedagogy (all University of California, Santa Cruz). The authors also acknowledge the National Science Foundation Division of Undergraduate Education (DUE 2150444) for supporting general CURE lab research at the University of California Santa Cruz.

## References

(1) Hausinger, R. P. Fe(II)/α-Ketoglutarate-Dependent Hydroxylases and Related Enzymes. Crit. Rev. Biochem. Mol. Biol. 2004, 39 (1), 21–68. 10.1080/10409230490440541.

(2) Islam, S.; Leissing, T. M.; Chowdhury, R.; Hopkinson, R. J.; Schofield, C. J. 2-Oxoglutarate-Dependent Oxygenases. Annu. Rev. Biochem. 2018, 87, 620–. 10.1146/annurev-biochem061516-044724.

(3) Ushimaru, R.; Abe, I. Unusual Dioxygen-Dependent Reactions Catalyzed by Nonheme Iron Enzymes in Natural Product Biosynthesis. ACS Catal. 2023, 13 (2), 1045–1076. 10.1021/acscatal.2c05247.

(4) Zwick, C. R.; Renata, H. Overview of Amino Acid Modifications by Iron- and α-Ketoglutarate-Dependent Enzymes. ACS Catal. 2023, 13 (7), 4853–4865. 10.1021/acscatal.3c00424.

(5) Martinez, S.; Hausinger, R. P. Catalytic Mechanisms of Fe(II)- and 2-Oxoglutarate-Dependent Oxygenases. J. Biol. Chem. 2015, 290 (34), 20702–20711. 10.1074/jbc.R115.648691.

(6) Bunno, R.; Awakawa, T.; Mori, T.; Abe, I. Aziridine Formation by a Fe^II^ /α-Ketoglutarate Dependent Oxygenase and 2-Aminoisobutyrate Biosynthesis in Fungi. Angew. Chem. Int. Ed. 2021, 60 (29), 15827–15831. 10.1002/anie.202104644.

(7) Brunson, J. K.; McKinnie, S. M. K.; Chekan, J. R.; McCrow, J. P.; Miles, Z. D.; Bertrand, E. M.; Bielinski, V. A.; Luhavaya, H.; Oborník, M.; Smith, G. J.; Hutchins, D. A.; Allen, A. E.; Moore, B. S. Biosynthesis of the Neurotoxin Domoic Acid in a Bloom-Forming Diatom. Science 2018, 361 (6409), 1356–1358. 10.1126/science.aau0382.

(8) Hopiavuori, A. R.; McKinnie, S. M. K. Algal Kainoid Synthases Exhibit Substrate-Dependent Hydroxylation and Cyclization Activities. ACS Chem. Biol. 2023, 18 (12), 2457–2463. 10.1021/acschembio.3c00596.

(9) Marchand, J. A.; Neugebauer, M. E.; Ing, M. C.; Lin, C.-I.; Pelton, J. G.; Chang, M. C. Y. Discovery of a Pathway for Terminal-Alkyne Amino Acid Biosynthesis. Nature 2019, 567 (7748), 420–424. 10.1038/s41586-019-1020-y.

(10) Sanada, M.; Miyano, T.; Iwadare, S. β-Ethynylserine, an Antimetabolite of L-Threonine, from Streptomyces cattleya. J. Antibiot. (Tokyo) 39, 304–305. 10.7164/antibiotics.39.304.

(11) Neugebauer, M. E.; Kissman, E. N.; Marchand, J. A.; Pelton, J. G.; Sambold, N. A.; Millar, D. C.; Chang, M. C. Y. Reaction Pathway Engineering Converts a Radical Hydroxylase into a Halogenase. Nat. Chem. Biol. 2022, 18 (2), 171–179. 10.1038/s41589-021-00944-x.

(12) Kissman, E. N.; Neugebauer, M. E.; Sumida, K. H.; Swenson, C. V.; Sambold, N. A.; Marchand, J. A.; Millar, D. C.; Chang, M. C. Y. Biocatalytic Control of Site-Selectivity and Chain Length-Selectivity in Radical Amino Acid Halogenases. Proc. Natl. Acad. Sci. 2023, 120 (12), e2214512120. 10.1073/pnas.2214512120.

(13) Chen, Y.; Liu, W.-Q.; Zheng, X.; Liu, Y.; Ling, S.; Li, J. Cell-Free Biosynthesis of Lysine-Derived Unnatural Amino Acids with Chloro, Alkene, and Alkyne Groups. ACS Synth. Biol. 2023, 12 (4), 1349–1357. 10.1021/acssynbio.3c00132.

(14) Hopiavuori, A. R.; Huffman, R. T.; McKinnie, S. M. K. Expression, Purification, and Biochemical Characterization of Micro- and Macroalgal Kainoid Synthases. In Methods in Enzymology; Elsevier, 2024; Vol. 704, pp 233–258. 10.1016/bs.mie.2024.05.017.

(15) Marfey, P. Determination of D-Amino Acids. II. Use of a Bifunctional Reagent, 1,5-Difluoro-2,4-Dinitrobenzene. Carlesberg Res. Commun. 1984, 49, 596–.

(16) Cordoza, J. L.; Chen, P. Y.-T.; Blaustein, L. R.; Lima, S. T.; Fiore, M. F.; Chekan, J. R.; Moore, B. S.; McKinnie, S. M. K. Mechanistic and Structural Insights into a Divergent PLP-Dependent L - Enduracididine Cyclase from a Toxic Cyanobacterium. ACS Catal. 2023, 13 (14), 9817–9828. 10.1021/acscatal.3c01294.

(17) Abramson, J.; Adler, J.; Dunger, J.; Evans, R.; Green, T.; Pritzel, A.; Ronneberger, O.; Willmore, L.; Ballard, A. J.; Bambrick, J.; Bodenstein, S. W.; Evans, D. A.; Hung, C.-C.; O’Neill, M.; Reiman, D.; Tunyasuvunakool, K.; Wu, Z.; Žemgulytė, A.; Arvaniti, E.; Beattie, C.; Bertolli, O.; Bridgland, A.; Cherepanov, A.; Congreve, M.; Cowen-Rivers, A. I.; Cowie, A.; Figurnov, M.; Fuchs, F. B.; Gladman, H.; Jain, R.; Khan, Y. A.; Low, C. M. R.; Perlin, K.; Potapenko, A.; Savy, P.; Singh, S.; Stecula, A.; Thillaisundaram, A.; Tong, C.; Yakneen, S.; Zhong, E. D.; Zielinski, M.; Žídek, A.; Bapst, V.; Kohli, P.; Jaderberg, M.; Hassabis, D.; Jumper, J. M. Accurate Structure Prediction of Biomolecular Interactions with AlphaFold 3. Nature 2024, 630 (8016), 493–500. 10.1038/s41586-024-07487-w.

(18) Hanwell, M. D.; Curtis, D. E.; Lonie, D. C.; Vandermeersch, T.; Zurek, E.; Hutchison, G. R. Avogadro: An Advanced Semantic Chemical Editor, Visualization, and Analysis Platform. J. Cheminformatics 2012, 4 (1), 17. 10.1186/1758-2946-4-17.

(19) Trott, O.; Olson, A. J. AutoDock Vina: Improving the Speed and Accuracy of Docking with a New Scoring Function, Efficient Optimization, and Multithreading. J. Comput. Chem. 2010, 31 (2), 455–461. 10.1002/jcc.21334.

(20) Papadopoulou, A.; Meyer, F.; Buller, R. M. Engineering Fe(II)/α-Ketoglutarate-Dependent Halogenases and Desaturases. Biochemistry 2023, 62 (2), 229–240. 10.1021/acs.biochem.2c00115.

(21) Kastner, D. W.; Nandy, A.; Mehmood, R.; Kulik, H. J. Mechanistic Insights into Substrate Positioning That Distinguish Non-Heme Fe(II)/α-Ketoglutarate-Dependent Halogenases and Hydroxylases. ACS Catal. 2023, 13 (4), 2489–2501. 10.1021/acscatal.2c06241.

(22) Smithwick, E. R.; Wilson, R. H.; Chatterjee, S.; Pu, Y.; Dalluge, J. J.; Damodaran, A. R.; Bhagi-Damodaran, A. Electrostatically Regulated Active Site Assembly Governs Reactivity in Nonheme Iron Halogenases. ACS Catal. 2023, 13 (20), 13743–13755. 10.1021/acscatal.3c02531.

(23) Hibi, M.; Kawashima, T.; Kasahara, T.; Sokolov, P. M.; Smirnov, S. V.; Kodera, T.; Sugiyama, M.; Shimizu, S.; Yokozeki, K.; Ogawa, J. A Novel Fe(II)/α-Ketoglutarate-Dependent Dioxygenase from Burkholderia ambifaria Has β-Hydroxylating Activity of N -Succinyl L-Leucine: Enzymatic Production of β-Hydroxy Amino Acids. Lett. Appl. Microbiol. 2012, 55 (6), 414–419. 10.1111/j.1472-765X.2012.03308.x.

(24) Mitchell, A. J.; Dunham, N. P.; Bergman, J. A.; Wang, B.; Zhu, Q.; Chang, W.; Liu, X.; Boal, A. K. Structure-Guided Reprogramming of a Hydroxylase to Halogenate Its Small Molecule Substrate. Biochemistry 2017, 56 (3), 441–444. 10.1021/acs.biochem.6b01173.

(25) Gomez, C. A.; Mondal, D.; Du, Q.; Chan, N.; Lewis, J. C. Directed Evolution of an Iron(II)- and α-Ketoglutarate-Dependent Dioxygenase for Site-Selective Azidation of Unactivated Aliphatic C−H Bonds. Angew. Chem. 2023, 135 (15), e202301370. 10.1002/ange.202301370.

(26) Hibi, M.; Kasahara, T.; Kawashima, T.; Yajima, H.; Kozono, S.; Smirnov, S. V.; Kodera, T.; Sugiyama, M.; Shimizu, S.; Yokozeki, K.; Ogawa, J. Multi-Enzymatic Synthesis of Optically Pure β-Hydroxy α-Amino Acids. Adv. Synth. Catal. 2015, 357 (4), 767–774. 10.1002/adsc.201400672.

(27) Zhu, X.; Liu, J.; Zhang, W. De Novo Biosynthesis of Terminal Alkyne-Labeled Natural Products. Nat. Chem. Biol. 2015, 11 (2), 115–120. 10.1038/nchembio.1718.

(28) Li, X.; Lv, J.-M.; Hu, D.; Abe, I. Biosynthesis of Alkyne-Containing Natural Products. RSC Chem. Biol. 2021, 2 (1), 166–180. 10.1039/D0CB00190B.

(29) Ignacio, B. J.; Dijkstra, J.; Mora, N.; Slot, E. F. J.; Van Weijsten, M. J.; Storkebaum, E.; Vermeulen, M.; Bonger, K. M. THRONCAT: Metabolic Labeling of Newly Synthesized Proteins Using a Bioorthogonal Threonine Analog. Nat. Commun. 2023, 14 (1), 3367. 10.1038/s41467-023-39063-7.

(30) Packer, M. S.; Liu, D. R. Methods for the Directed Evolution of Proteins. Nat. Rev. Genet. 2015, 16 (7), 379–394. 10.1038/nrg3927.

(31) Arnold, F. H. Directed Evolution: Bringing New Chemistry to Life. Angew. Chem. Int. Ed. 2018, 57 (16), 4143–4148. 10.1002/anie.201708408.

(32) Shepherd, R. A.; Fihn, C. A.; Tabag, A. J.; McKinnie, S. M. K.; Sanchez, L. M. ‘Need for Speed: High Throughput’ – Mass Spectrometry Approaches for High-Throughput Directed Evolution Screening of Natural Product Enzymes. Nat. Prod. Rep. 2025, 42 (6), 1037–1054. 10.1039/D4NP00053F.

(33) Cheung-Lee, W. L.; Kolev, J. N.; McIntosh, J. A.; Gil, A. A.; Pan, W.; Xiao, L.; Velásquez, J. E.; Gangam, R.; Winston, M. S.; Li, S.; Abe, K.; Alwedi, E.; Dance, Z. E. X.; Fan, H.; Hiraga, K.; Kim, J.; Kosjek, B.; Le, D. N.; Marzijarani, N. S.; Mattern, K.; McMullen, J. P.; Narsimhan, K.; Vikram, A.; Wang, W.; Yan, J.; Yang, R.; Zhang, V.; Zhong, W.; DiRocco, D. A.; Morris, W. J.; Murphy, G. S.; Maloney, K. M. Engineering Hydroxylase Activity, Selectivity, and Stability for a Scalable Concise Synthesis of a Key Intermediate to Belzutifan. Angew. Chem. Int. Ed. 2024, 63 (13), e202316133. 10.1002/anie.202316133.

(34) Renata, H. Combining Biocatalytic and Radical Retrosynthesis for Efficient Chemoenzymatic Synthesis of Natural Products. Chem. Soc. Rev. 2025, 54 (17), 7913–7932. 10.1039/D5CS00453E.

(35) Paton, A. E.; Boiko, D. A.; Perkins, J. C.; Cemalovic, N. I.; Reschützegger, T.; Gomes, G.; Narayan, A. R. H. Connecting Chemical and Protein Sequence Space to Predict Biocatalytic Reactions. Nature 2025, 646 (8083), 108–116. 10.1038/s41586-025-09519-5.

(36) Huls, A. J.; Soler, J.; Su, Y.; Yang, Y.; Garcia-Borràs, M.; Huang, X. Biocatalytic Olefin Difunctionalization for Synthesis of Chiral 2-Azidoamines Using Nonheme Iron Enzymes. Angew. Chem. Int. Ed. 2025, 64 (39), e202423403. 10.1002/anie.202423403.

(37) Tao, H.; Mori, T.; Chen, H.; Lyu, S.; Nonoyama, A.; Lee, S.; Abe, I. Molecular Insights into the Unusually Promiscuous and Catalytically Versatile Fe(II)/α-Ketoglutarate-Dependent Oxygenase SptF. Nat. Commun. 2022, 13 (1), 95. 10.1038/s41467-021-27636-3.

(38) Yang, D.; Chiang, C.-H.; Wititsuwannakul, T.; Brooks, C. L.; Zimmerman, P. M.; Narayan, A. R. H. Engineering the Reaction Pathway of a Non-Heme Iron Oxygenase Using Ancestral Sequence Reconstruction. J. Am. Chem. Soc. 2024, 146 (50), 34352–34363. 10.1021/jacs.4c08420.

(39) Jouaneh, T. M. M.; Rosario, M. E.; Li, Y.; Leibovitz, E.; Bertin, M. J. Incorporating LC–MS/MS Analysis and the Dereplication of Natural Product Samples into an Upper-Division Undergraduate Laboratory Course. J. Chem. Educ. 2022, 99 (7), 2636–2642. 10.1021/acs.jchemed.1c01212.

(40) Cho, Y. I.; Armstrong, C. L.; Sulpizio, A.; Acheampong, K. K.; Banks, K. N.; Bardhan, O.; Churchill, S. J.; Connolly-Sporing, A. E.; Crawford, C. E. W.; Cruz Parrilla, P. L.; Curtis, S. M.; De La Ossa, L.M.; Epstein, S. C.; Farrehi, C. J.; Hamrick, G. S.; Hillegas, W. J.; Kang, A.; Laxton, O. C.; Ling, J.; Matsumura, S. M.; Merino, V. M.; Mukhtar, S. H.; Shah, N. J.; Londergan, C. H.; Daly, C. A.; Kokona, B.; Charkoudian, L. K. Engineered Chimeras Unveil Swappable Modular Features of Fatty Acid and Polyketide Synthase Acyl Carrier Proteins. Biochemistry 2022, 61 (4), 217–227. 10.1021/acs.biochem.1c00798.

(41) Cummings, C. B.; Catania, S. S.; Ednacot, E. M. Q.; Kinsella-Johnson, A. J.; Meeds, C. E.; Reynolds, J. W.; Sanderson, A. E.; Johnson, R. A.; Watts, K. R. Improving Student Outcomes with an Adaptable Molecular Cloning Course-Based Undergraduate Research Experience. J. Vis. Exp. 2024, No. 213, 67067. 10.3791/67067.

(42) Tang, Y.; Zhong, W.; Fu, L.; Asante, E.; Kostenko, A.; Vidya, F. N. U.; Mandelare-Ruiz, P.; Adeogun, T. T.; Anderson, G. P.; Edmonds, B. E.; Fang, O.; Han, M.; Hollingsworth, A. S.; Ingham, A. R.; Kirby, C. R.; Landrum, A.; Mack, C. R.; Nobari, N. S.; Oswald, E. J.; Polevoy, C. L.; Sharifian, Y.; So, T. J.; Stokes, J. R.; Thompson, R. S.; Vuthamaraju, R.; Wang, E. C.; Yang, W. H.; Onstine, A. E.; Paul, V. J.; Wu, R.; Aron, A. T.; Agarwal, V. Phylogenomic Identification of a Highly Conserved Copper-Binding RiPP Biosynthetic Gene Cluster in Marine Microbulbifer Bacteria. ACS Chem. Biol. 2025, acschembio.5c00507. 10.1021/acschembio.5c00507.

