## Supplementary material for "CURE-ating the substrate scope and functional residues of nonheme iron(II) α-ketoglutarate dependent hydroxylase BesE": BesE-CURE-SI-PDF

**This PDF file includes:**

General Materials and Methods

Supplementary Figures S1 to S36

Chemical Synthesis

Tables S1 and S2

NMR Spectra of Characterized Compounds

References

#### Table of Contents


**General Materials & Methods** **S5**

**1. Molecular Biology/Biochemical Methods** **S6**

*besE* transformation into *E. coli* **S6**
BesE expression in BL21(DE3) *E. coli* **S6**
BesE FPLC affinity purification **S6**
Site-directed mutagenesis of *besE,* KLD and transformation into DH10β *E. coli* **S7**
Expression of *besE* mutants in BL21(DE3) *E. coli* **S7**
Nickel-NTA batch resin purification of BesE mutants **S7**
Analytical BesE assays with γ-L-glutamyl-L-propargylglycine substrate (**1**) and analogs **S8**
Marfey's derivatization of BesE *in vitro* assays **S8**
Cu-catalyzed azide-alkyne cycloaddition derivatization of BesE *in vitro* assays **S8**

**2. Supplementary Figures** **S9**

Figure S1 – Biosynthesis of β-ethynylserine in *Streptomyces cattleya* **S9**
Figure S2 – Scheme demonstrating that the radical intermediate formed by BesE does not undergo rearrangement to the allenyl resonance structure **S10**
Figure S3 – Attempted methyl esterification of β-cyanoalanine **S11**
Figure S4 – Initial *in vitro* assay with purified BesE. **S12**
Figure S5 – *in vitro* assay with purified BesE showing improved conversion with adjusted L-ascorbate and $FeSO_4$ concentrations **S13**
Figure S6 – *in vitro* assay with purified BesE comparing underivatized compounds to samples derivatized with Marfey's reagent **S14**
Figure S7 – *in vitro* assay with purified BesE post-derivatization via CuAAC with coumarin azide **S15**
Figure S8 – *in vitro* BesE assay with analog **5** **S16**
Figure S9 – *in vitro* BesE assay with analog **6** **S17**
Figure S10 – *in vitro* BesE assay with analog **7** **S18**
Figure S11 – *in vitro* BesE assay with analog **8** **S19**
Figure S12 – *in vitro* BesE assay with analog **9** **S20**
Figure S13 – *in vitro* BesE assay with analog **10** **S21**
Figure S14 – *in vitro* BesE assay with analog **11** **S22**
Figure S15 – *in vitro* BesE assay with analog **12** **S23**
Figure S16 – *in vitro* BesE assay with analog **13** **S24**
Figure S17 – *in vitro* BesE assay with analog **14** **S25**
Figure S18 – *in vitro* BesE assay with analog **15** **S26**
Figure S19 – *in vitro* BesE assay with analog **16** **S27**
Figure S20 – *in vitro* BesE assay with analog **17** **S28**

Figure S21 – *in vitro* BesE assay with analog **18** **S29**
Figure S22 – Structural depiction of BesE residues selected for site-directed mutagenesis. **S30**
Figure S23 – Conservation of BesE residues selected for site-directed mutagenesis across putative actinobacterial homologs **S31**
Figure S24 – 10% SDS-PAGE gel of purified BesE mutants **S32**
Figure S25 – Representative trace of in vitro BesE F255A assay with **1** **S33**
Figure S26 – Representative trace of in vitro BesE V146A assay with **1** **S34**
Figure S27 – Representative trace of in vitro BesE M70A assay with **1** **S35**
Figure S28 – Representative trace of in vitro BesE V240A assay with **1** **S36**
Figure S29 – Representative trace of in vitro BesE A175W assay with **1** **S37**
Figure S30 – Representative trace of in vitro BesE T150A assay with **1** **S38**
Figure S31 – Representative trace of in vitro BesE R144A assay with **1** **S39**
Figure S32 – Representative trace of in vitro BesE D155A assay with **1** **S40**
Figure S33 – Representative trace of in vitro BesE Q173A assay with **1** **S41**
Figure S34 – Representative trace of in vitro BesE Y207A assay with **1** **S42**
Figure S35 – Representative trace of in vitro BesE R249A assay with **1** **S43**
Figure S36 – in vitro BesE D155A assay with **1** is not indicative of chlorinated product formation. **S44**

**3. Chemical Synthesis** **S45**

General reaction procedures for BesE native substrate and analog syntheses **S45**
Synthesis of γ-glutamyl-L-propargylglycine (**1**) **S46**
Synthesis of γ-glutamyl-L-allylglycine (**5**) **S48**
Synthesis of γ-glutamyl-L-norvaline (**6**) **S50**
Synthesis of γ-glutamyl-L-homopropargylglycine (**7**) **S52**
Synthesis of γ-glutamyl-L-alanine (**8**) **S54**
Synthesis of γ-glutamyl-D-propargylglycine (**9**) **S56**
Synthesis of γ-glutamyl-L-propargylglycine methyl ester (**10**) **S58**
Synthesis of γ-glutamyl-(*R*)-pent-4-yn-2-amine (**11**) **S59**
Synthesis of γ-glutamyl-3-butyn-1-amine **12**) **S61**
Synthesis of γ-glutamyl-L-propargylglycine-α-amide (**13**) **S63**
Synthesis of γ-aminobutyryl-L-propargylglycine (**14**) **S65**
Synthesis of γ-glutaryl-L-propargylglycine (**15**) **S67**
Synthesis of *N*-butanoyl-L-propargylglycine (**16**) **S69**
Synthesis of β-aspartyl-L-propargylglycine (**17**) **S70**
Synthesis of γ-D-glutamyl-L-propargylglycine (**18**) **S72**

**4. Tables** **S74**

Table S1. Primers used in this study **S74**
Table S2. Specific contributions to analog synthesis and mutant generation **S75**

**5. NMR Spectra of Characterized Compounds** **S76**

γ-glutamyl-L-propargylglycine (**1**) **S76**
*O*-methyl-γ-glutamyl-L-propargylglycine (**3**) **S78**
γ-(*N*-Boc-*O*-tert-butyl-glutamyl)-*O*-methyl-L-propargylglycine (**4**) **S80**
γ-glutamyl-L-allylglycine (**5**) **S82**
γ-glutamyl-L-norvaline (**6**) **S84**
γ-glutamyl-L-homopropargylglycine (**7**) **S86**
γ-glutamyl-L-alanine (**8**) **S88**
γ-glutamyl-D-propargylglycine (**9**) **S90**
γ-glutamyl-L-propargylglycine methyl ester (**10**) **S92**
γ-glutamyl-(*R*)-pent-4-yn-2-amine (**11**) **S94**
γ-glutamyl-3-butyn-1-amine (**12**) **S96**
γ-glutamyl-L-propargylglycine-α-amide (**13**) **S98**
γ-aminobutyryl-L-propargylglycine (**14**) **S100**
γ-glutaryl-L-propargylglycine (**15**) **S102**
*N*-butanoyl-L-propargylglycine (**16**) **S104**
β-aspartyl-L-propargylglycine (**17**) **S106**
γ-D-glutamyl-L-propargylglycine (**18**) **S108**

**6. References** **S110**

#### General Materials & Methods

All chemicals, including solvents and media components, were used as received from commercial suppliers (Millipore Sigma, Thermo Fisher Scientific, Enamine). Reagents were used without further purification unless otherwise noted.

##### Synthetic and analytical methods

Anhydrous reactions were done in oven-dried glassware and carried out under an argon atmosphere. Reactions were monitored using thin-layer chromatography (TLC, Merck, 60 $F_{254}$). TLC plates were visualized with a UV lamp at 254 nm and stained with either potassium permanganate (1.5 g $KMnO_4$, 10 g $K_2CO_3$, 1.25 mL 10% NaOH in 200 mL water) or ninhydrin (1.5 g ninhydrin, 3 mL AcOH in 100 mL *n*-butanol) staining solutions. TLC $R_f$ values were rounded to the nearest 0.05. Silica ((Alfa Aesar, 60 (215-400 mesh)) was used when flash column chromatography was employed to purify compounds. Rotary evaporation was used to concentrate samples under reduced pressure. Additionally, polar final products were concentrated and dried via lyophilization (Labconco FreeZone 4.5 Liter). Nuclear magnetic resonance (NMR) spectra were obtained using an Avance III HD spectrometer (Bruker) equipped with a BBFO SmartProbe at 500 MHz ($^1$H NMR) or 125 MHz ($^{13}$C NMR) using $CDCl_3$ or $D_2O$ as solvents. Chemical shifts (δ) are reported in ppm and referenced to methanol (δ = 3.31 ppm for $^1$H, δ = 49.0 ppm for $^{13}$C NMR) as an internal standard for samples in $D_2O$ or the $CDCl_3$ solvent signal (δ = 7.26 ppm for $^1$H, δ = 77.2 ppm for $^{13}$C NMR) for samples in $CDCl_3$. NMR data are reported as follows: s = singlet, d = doublet, t = triplet, q = quartet, m = multiplet, *J* = coupling constant in Hz.

##### LCMS & HPLC instrumentation

General LCMS measurements were measured on a Bruker Elute UHPLC system coupled with a Bruker amazon SL ESI-Ion Trap mass spectrometer in positive mode. Compounds were separated via reversed-phase chromatography on a Bruker Intensity Solo C18(2), 2 µm- 2 x 100 mm column with the eluents water + 0.1% formic acid (Solvent A) and acetonitrile + 0.1% formic acid (Solvent B). The LC method uses a flow rate of 0.3 mL/min and the following gradient: 5% B (3 min), 5%-15% B (3 min), 15%-100% B (3 min), 100% B (2 min), 100%-5% B (1 min), 5% B (2 min).

#### **Molecular Biology/Biochemical Methods**

*besE* transformation into *E. coli*

A pET28a(+) plasmid containing the N-terminal hexahistidine (His6) tagged *BesE* gene from *Streptomyces cattleya* was transformed into both *E. coli* DH10β and BL21(DE3) chemically competent cell lines for plasmid production and protein expression respectively. Transformation was performed via heat shock according to the following protocol: 1 µL of plasmid was added to 100 µL chemically competent cells and the mixture was incubated on ice for 30 minutes. The cells were then heated to 42 °C for 55 sec and placed on ice again for 5 min; 700 µL of LB medium was added in the tube and the cells were incubated for 45 min at 37 °C and 200 rpm of agitation. After this step, 300 µL of the recovered cells were plated on LB agar plates supplemented with kanamycin (50 µg / mL). The plates were incubated at 37 °C overnight. A single colony was used to inoculate 5 mL of LB supplemented with kanamycin which was grown overnight at 37 °C and 200 rpm of agitation and purified following the protocol of Plasmid DNA Purification QIAprep Spin Miniprep Kit (QIAGEN). After plasmid purification, concentrations were measured by NanoDrop UV-vis spectrophotometry and stored at -20 °C.

BesE expression in BL21(DE3) *E. coli*

A single colony of BesE transformed BL21(DE3) cells was used to inoculate 20 mL of LB media supplemented with kanamycin (50 µg/mL). The culture was grown overnight at 37 °C, 200 rpm. 10 mL of the overnight culture was then added to 1 L of Terrific Broth supplemented with kanamycin (50 µg/mL). The 1 L culture was grown in a shaking incubator at 37 °C, 200 rpm until OD600 reached ~0.6. The flasks were then cooled to 18 °C for 1 h at which point 0.1 mM IPTG was added to induce protein expression. The cultures were left to grow at 18 °C and 200 rpm overnight (16 - 18 h) and pelleted in a centrifuge at 3000 x g for 30 min at 4 °C. Supernatant was discarded and the cell pellet was resuspended in 40 mL of cold buffer A (1 M NaCl, 20 mM Tris-HCl pH 8.0) and stored at -70 °C.

BesE FPLC affinity purification

BesE resuspended cell pellet was thawed and lysed on ice via sonication at 40% amplitude for 10 cycles of 15 sec on and 45 sec off (1 min total time per cycle). The lysate was pelleted via centrifugation at 4 °C and ~16,000 *x g* for 30 min. The cleared lysate was loaded onto a 5 mL HisTrap FF column (GE Healthcare Life Sciences) that was pre- equilibrated with 5 CV of buffer A. After loading, the column was washed with buffer A until the UV absorbance reached < 50 mAU. The column was then washed with 10% buffer B (1 M NaCl, 20 mM Tris-HCl pH 8.0, 250 mM imidazole) for 5 CV to remove any non-specifically bound proteins. BesE was then eluted using a linear gradient of 100% buffer A to 100% buffer B over 60 mL and 5 mL fractions were collected. The flowrate through the column was 2 mL/min. Fractions were assessed for presence of protein and purity using SDS-PAGE. Fractions containing BesE had 2 mM EDTA added and were pooled and concentrated by Amicon ultra centrifugal filters (10 kDa molecular weight cut-off) to a volume less than 2.5 mL. The concentrated protein was then further purified via a PD-10 desalting column (Cytiva) pre-equilibrated with GF buffer (50 mM HEPES-KOH pH 8.0, 300 mM KCl) following the default gravity filtration protocol. Enzyme concentration was determined via Bradford assay and pure enzyme was either used immediately for assays or aliquoted and stored at -70 °C.

Site-directed mutagenesis of *besE*, KLD, and transformation into DH10β *E. coli*

Site-directed mutagenesis PCR of the *besE* plasmid was performed using the Q5 Site-Directed Mutagenesis Kit (New England Biolabs) following the manufacturer's protocol and using the primers listed in table S1. The concentration of *besE* plasmid used as the template was 1 ng/µL. The general PCR protocol for each primer set was as follows: initial denaturation 98 °C for 30 sec, then 30 cycles of the following: denaturation at 98 °C for 10 sec, annealing at the 3-5 °C less than the $T_a$ for 10 sec (where $T_a$ is the annealing temperature determined for each primer pair), and extension at 72 °C for 2 min. Lastly, a final extension at 72 °C for 2 min followed by a hold at 4 °C. Upon completion of the PCR, a KLD reaction was performed to linearize the resulting plasmid containing the mutant gene and digest residual wildtype *besE* plasmid. The 10 µL reaction was prepared using KLD Enzyme Mix and KLD Reaction Buffer (New England Biolabs) and followed the manufacturer's KLD Enzyme Mix reaction protocol. The reaction was left to incubate at room temperature for 5 min. The entirety of the reaction was then added to a 100 µL aliquot of *E. coli* DH10β chemically competent cells and transformed as previously described. A single colony was used to generate a 5 mL inoculum from which plasmid was purified as described previously. Successfully generated mutants were confirmed via Sanger sequencing (Azenta Life Sciences) and transformed into *E. coli* Bl21(DE3) chemically competent cells as previously described for future expression.

Expression of *besE* mutants in BL21(DE3) *E. coli*

*besE* mutant expression was carried out in an identical way to the wildtype *besE* described without modification and resuspended cell pellets in A buffer were either immediately purified or stored at -80 °C.

Nickel-NTA batch resin purification of BesE mutants

Each individual resuspended BesE mutant pellet was lysed via sonication and centrifuged as previously described for the wildtype BesE. The clarified supernatant was then loaded onto an Econo-Column® Chromatography Column (2.5 x 10 cm; Bio-Rad Laboratories). containing 5 mL nickel-NTA batch resin that was pre-equilibrated with 3 - 5 CV of buffer A (20 mM Tris 1M NaCl pH 8.0). After loading, the resin was washed with an additional 5 CV buffer A. The protein was then eluted with 5 CV 20% buffer B followed by 5 CV 50%

buffer B, followed by 5 CV 100% buffer B (20 mM Tris 1M NaCl 250mM imidazole pH 8.0). All fractions were assessed for purity using a 10% SDS-PAGE gel (Fig. S3). The 100% B fraction containing BesE had 2 mM EDTA added to remove any bound iron and was concentrated and further purified via PD-10 desalting column as previously described for wildtype BesE. Enzyme concentration was determined via Bradford assay and pure mutant enzyme was aliquoted and stored at -80 °C.

Analytical BesE assays with γ-glutamyl-propargylglycine substrate **1** and analogs

Analytical BesE enzyme assays were conducted in 50 mM potassium phosphate (KPi) buffer (pH 8.0) with 10 mM L-ascorbate, 5 mM $\alpha$-ketoglutarate, and 1 mM FeSO4 with 1 mM of substrate **1** and 50 µM of purified BesE. Enzyme followed by FeSO4 were the last additions to assay and the reaction was allowed to run for 14 h. The total volume for the reaction was 100 µL. The reaction was then either quenched with 0.1 mM chloramphenicol in methanol (1 eq.) or further derivatized via Marfey's reagent or Cu-catalyzed azide-aklyne cycloaddition (CuAAC) with 7-azido-4-methylcoumarin (see below). Non-derivatized and/or derivatized reactions were centrifuged at ~ 13,000 x g for 5 min and the supernatant was then subjected to analysis by UPLC-MS. For analytical assays involving substrate analogs and BesE mutants, the same procedure was followed except 0.5 mM of substrate and 20 µM of purified enzyme were used.

Marfey's derivatization of BesE *in vitro* assays

To a 50 µL of enzyme assay reaction mixture was added 20 µL saturated sodium bicarbonate and 100 µL 1% w/v Marfey's reagent in acetone. The reaction was then incubated at 37 °C for 90 minutes followed by quench with 30 µL 1 N HCl. The sample was then centrifuged at 13,000 x *g* for 10 minutes and the supernatant was subjected to analysis by UPLC-MS.

Cu-Catalyzed Azide-Alkyne Cycloaddition for derivatization of BesE *in vitro* assays

To 25 µL of enzyme assay reaction mixture was added the following: 1.5 mM 7-azido-4-methylcoumarin, 0.2 mM $CuSO_4$, 0.6 mM sodium ascorbate. The total reaction volume was then brought to 50 µL with MilliQ water and left incubating at room temperature for 1 h. The sample was then centrifuged for 5 min at >13,000 x *g* and the supernatant was subjected to analysis by UPLC-MS.

**Supplementary Figures**

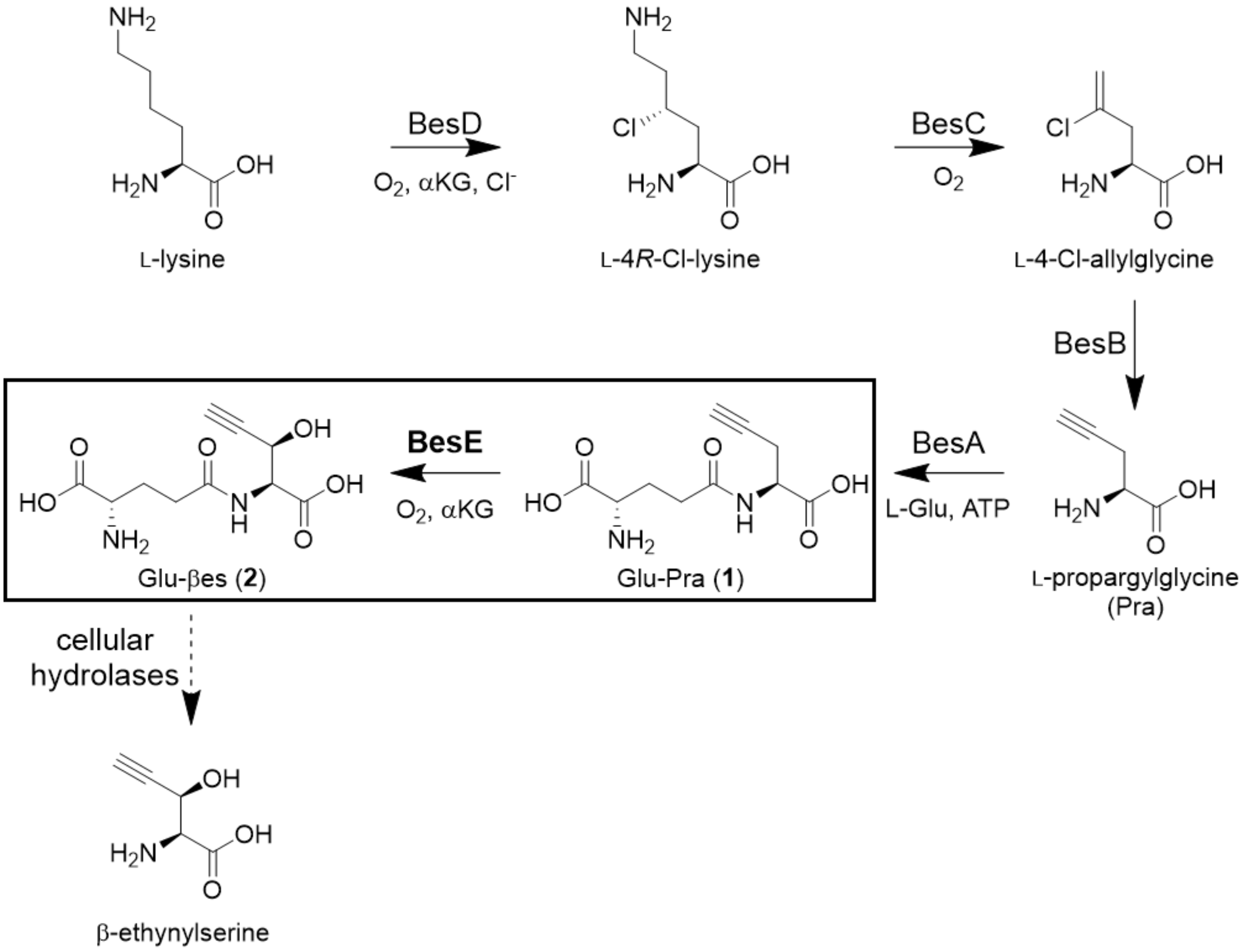


**Figure S1 – Biosynthesis of β-ethynylserine in *Streptomyces cattleya*.** BesE catalyzed hydroxylation of γ-L-Glu-L-Pra (**1**) to form γ-L-Glu-L-βes (**2**) boxed**.**

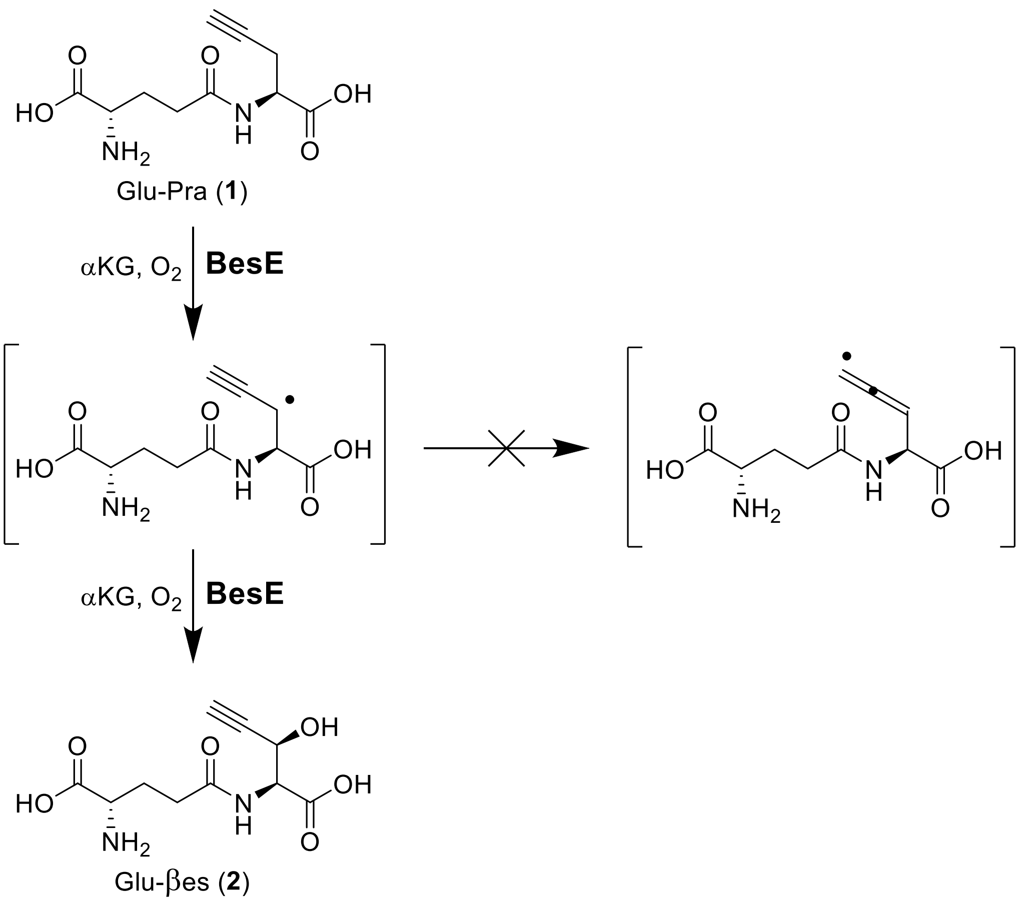


**Figure S2 – Scheme demonstrating that the radical intermediate formed by BesE does not undergo rearrangement to the allenyl resonance structure.**

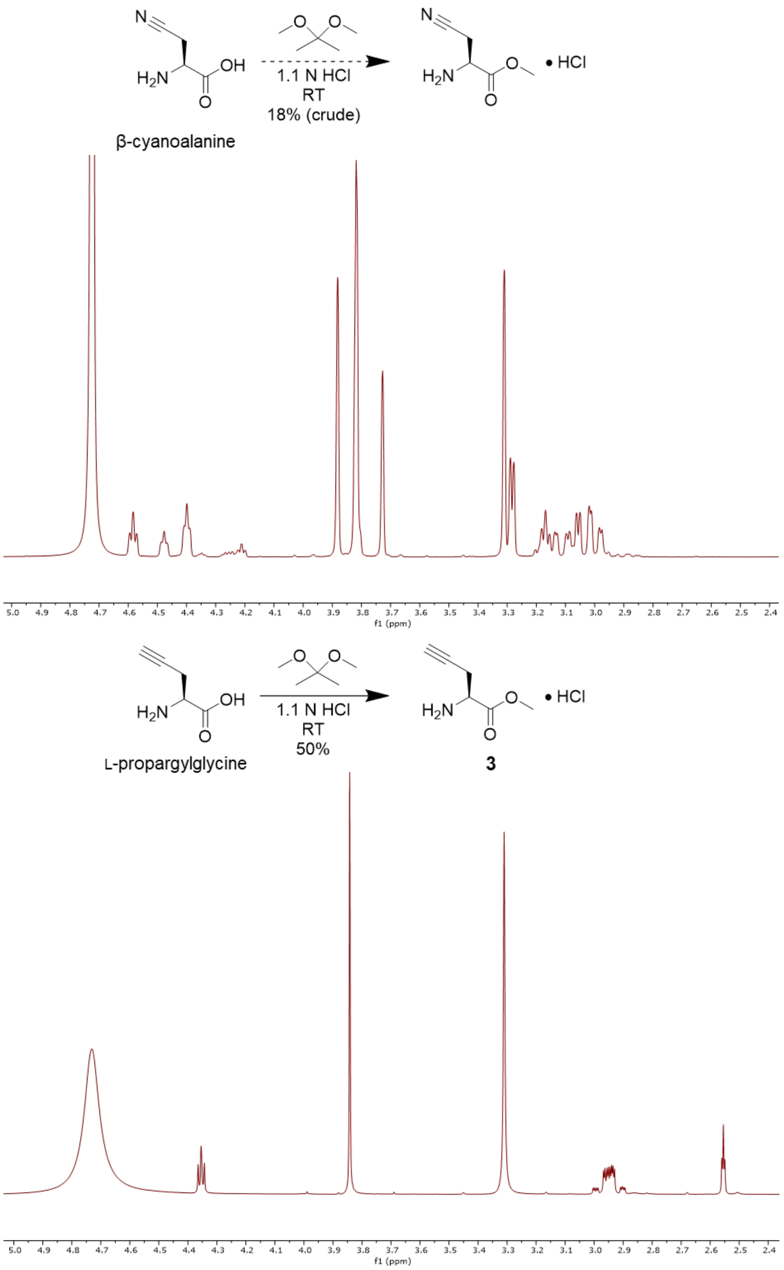


**Figure S3 – Attempted methyl esterification of β-cyanoalanine.** Synthetic scheme for methyl esterification of β-cyanoalanine with $^{1}H$ NMR (500 MHz, $D_2O$ + 0.1% MeOH) shown underneath (Top). NMR indicates the presence of additional unexpected signals. Scheme and $^{1}H$ NMR for successful methyl esterification of L-propargylglycine to form **3** shown for comparison (Bottom).

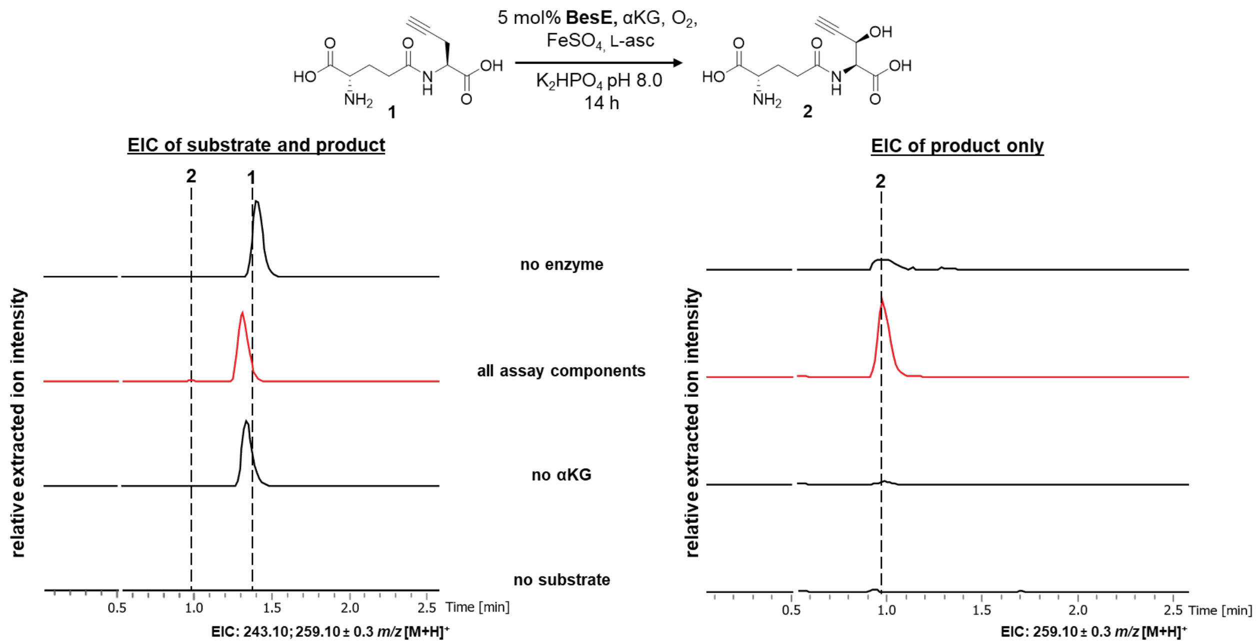


**Figure S4 – Initial *in vitro* assay with purified BesE.** Assays analyzed by UHPLC-MS in positive mode. Relative intensities were extracted for either substrate (**1**) and product (**2**) (left; EIC: 243.10; 259.10 ± 0.3 *m/z* $[M+H]^+$) or just the product mass (right; EIC: 259.10 ± 0.3 *m/z* $[M+H]^+$).

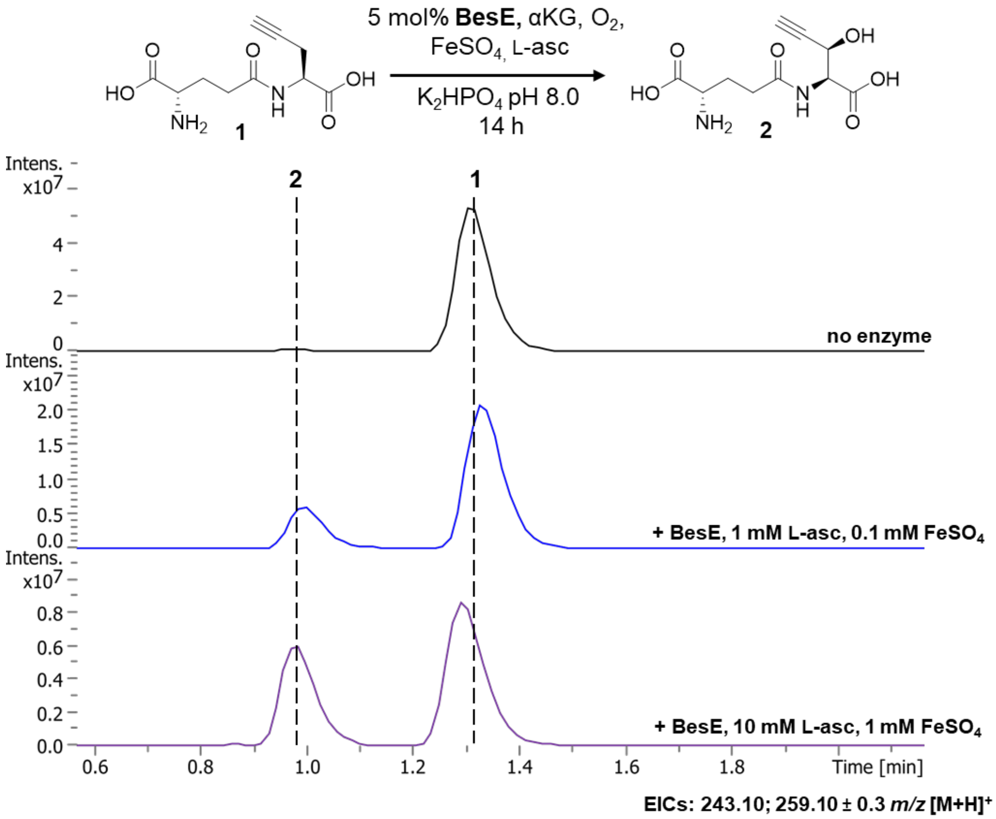


**Figure S5 – *in vitro* assay with purified BesE showing improved conversion with adjusted L-ascorbate and $FeSO_4$ concentrations.** Assays analyzed by UHPLC-MS in positive mode. Extracted ion chromatograms for substrate (**1**) and product (**2**) masses shown (EIC: 243.10; 259.10 ± 0.3 *m/z* $[M+H]^+$).

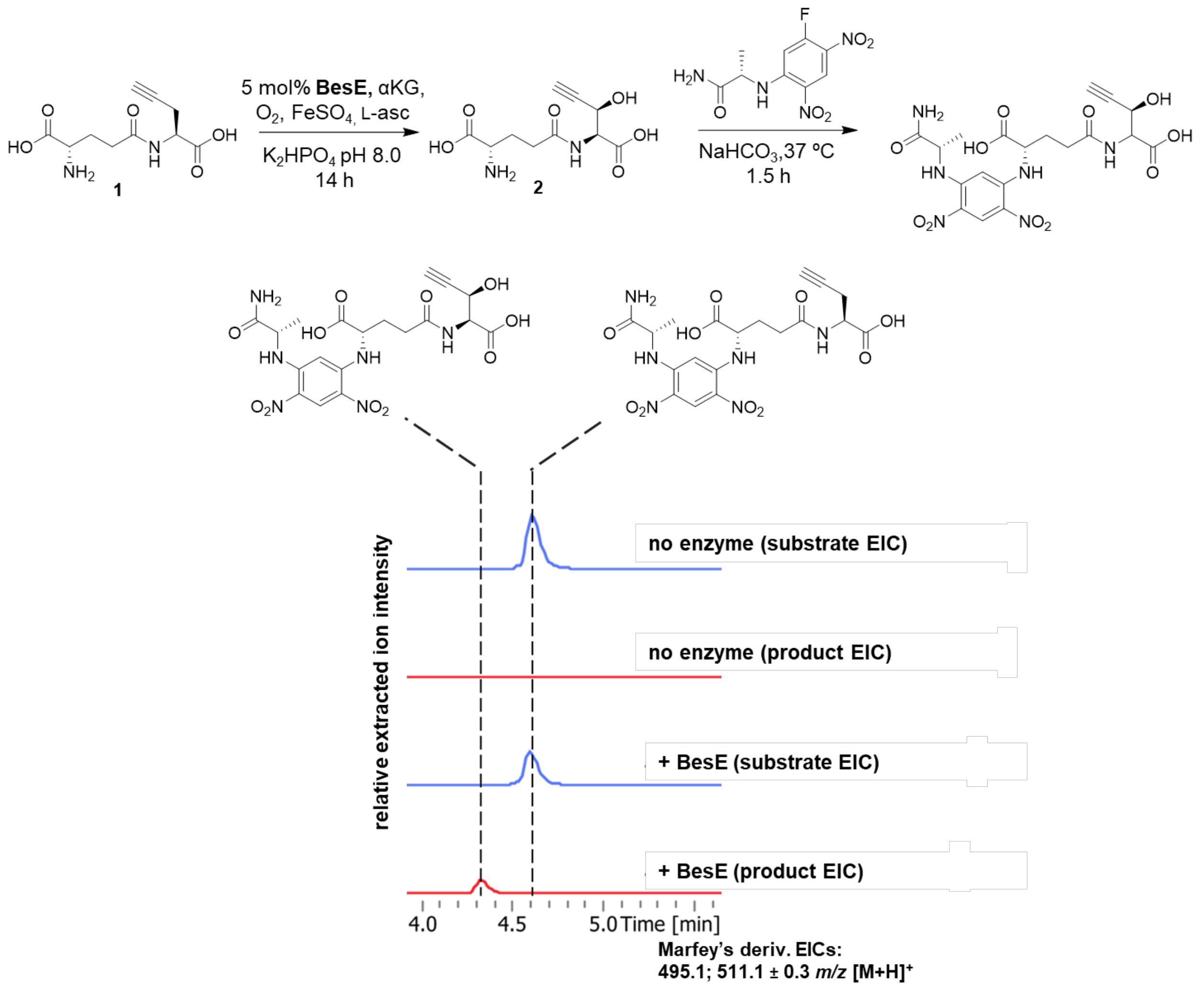


**Figure S6 – *in vitro* assay with purified BesE post-derivatization with Marfey's reagent.** Assays analyzed by UHPLC-MS in positive mode post-Marfey's derivatization Extracted ion chromatograms for Marfey's derivatized substrate (**1**) and product (**2**) masses shown (EIC: 495.1 ; 511.1 ± 0.3 *m/z* $[M+H]^+$).

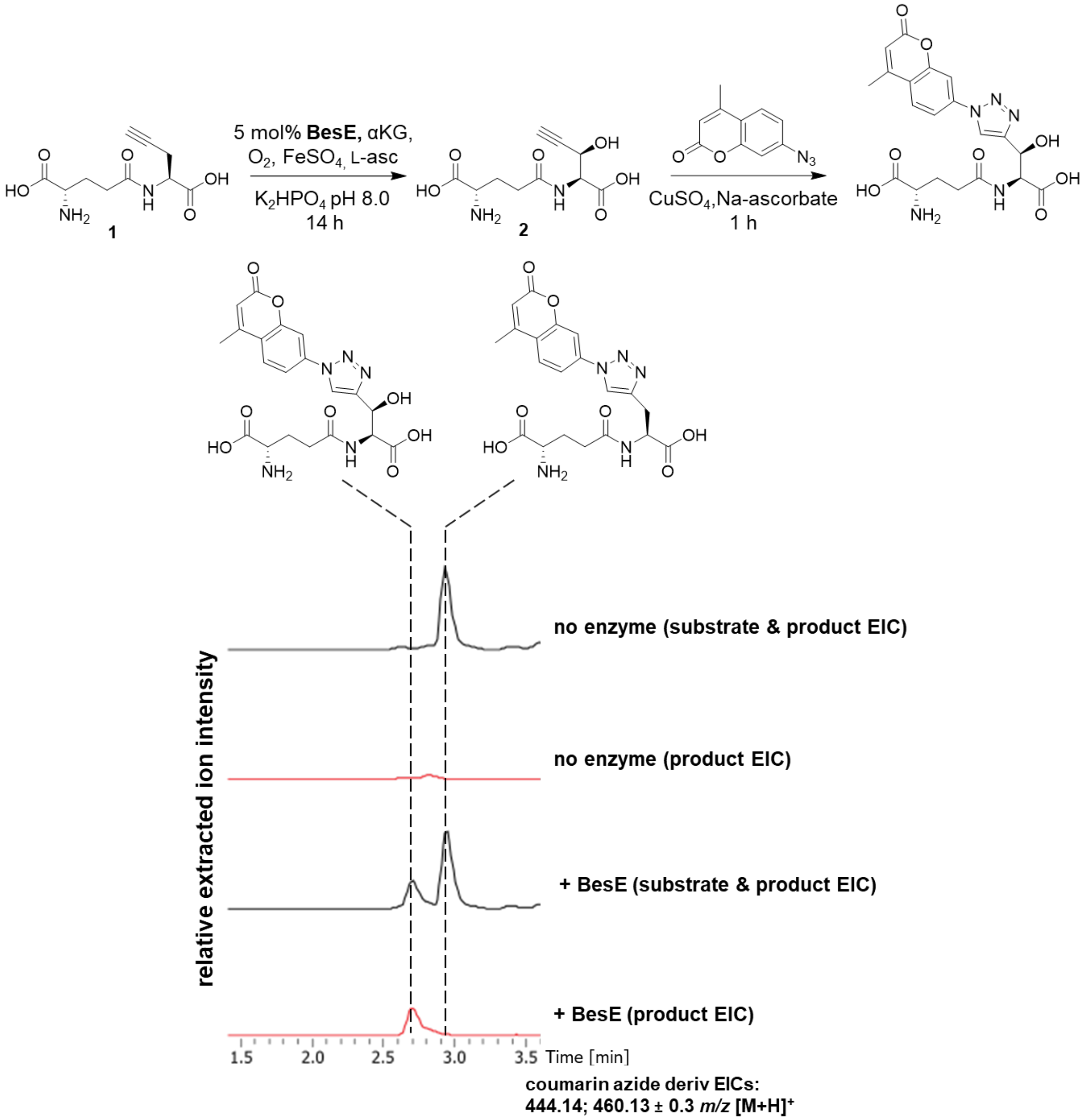


**Figure S7 – *in vitro* assay with purified BesE post-derivatization via CuAAC with 7-azido-4-methylcoumarin.** Assays analyzed by UHPLC-MS in positive mode post-derivatization. Extracted ion chromatograms for derivatized substrate (**1**) and product (**2**) masses (EIC: 444.14; 460.13 ± 0.3 *m/z* $[M+H]^+$).

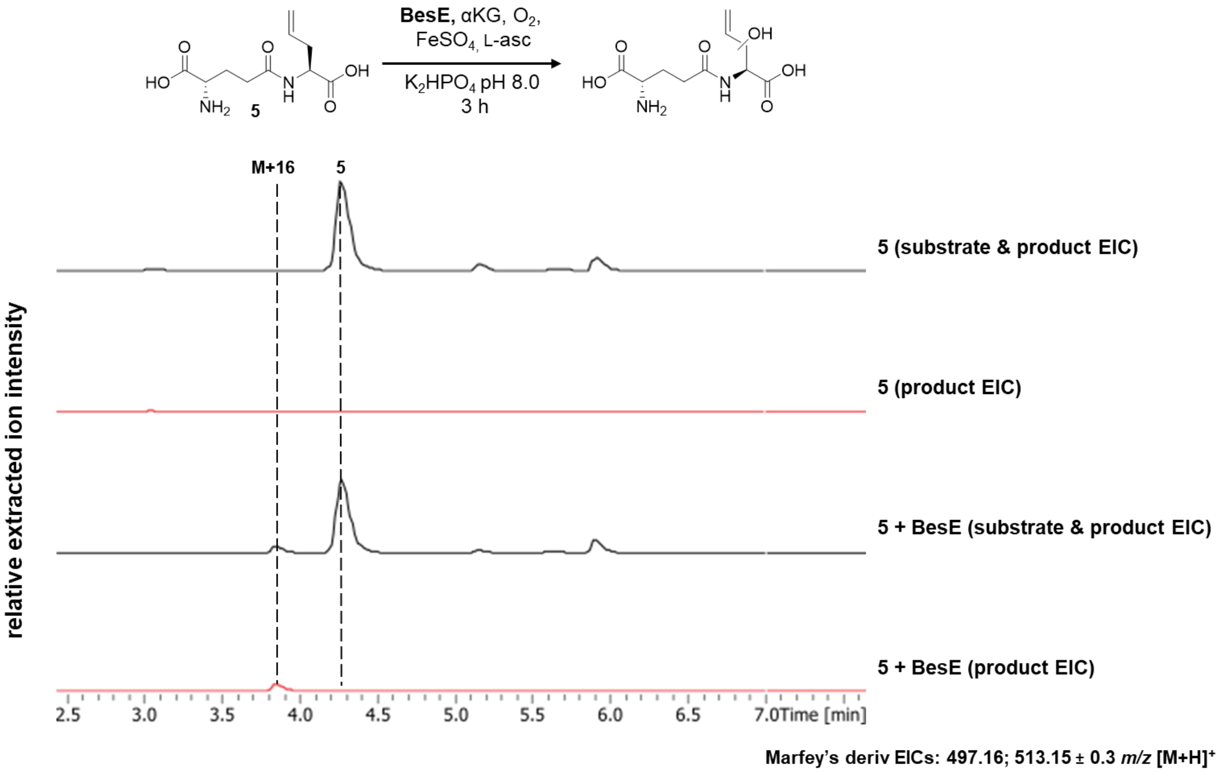


**Figure S8 – *in vitro* BesE assay with analog 5.** Assays analyzed by UHPLC-MS in positive mode post-Marfey's derivatization. Relative intensities were extracted for derivatized substrate and hydroxylated product masses (EIC: 497.16; 513.15 ± 0.3 *m/z* $[M+H]^+$).

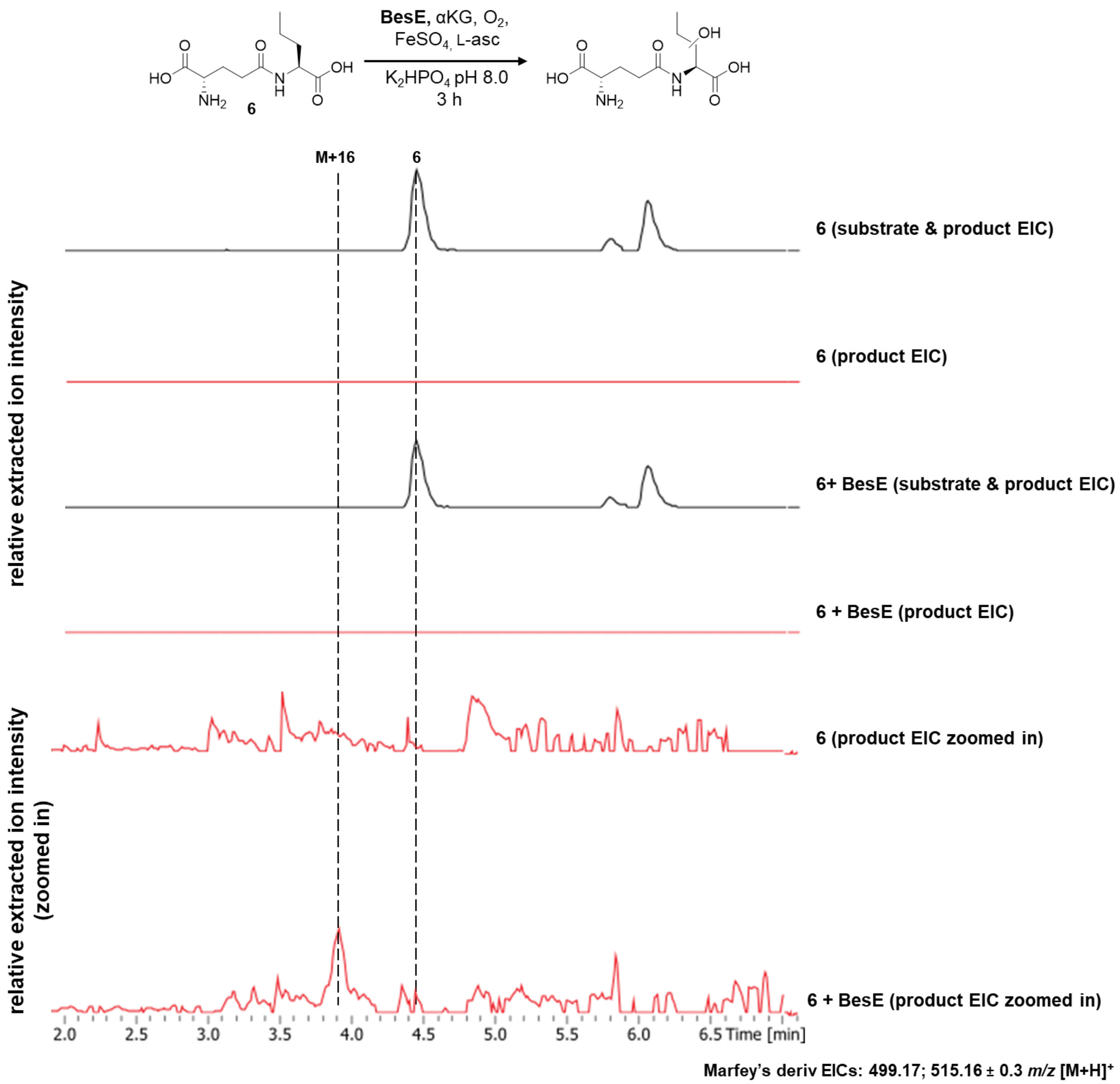


**Figure S9 – *in vitro* BesE assay with analog 6.** Additional product EICs for +/- enzyme conditions shown at a reduced scale to show the presence of a hydroxylated mass in the + BesE condition. Assays analyzed by UHPLC-MS in positive mode post-Marfey's derivatization. Relative intensities were extracted for derivatized substrate and hydroxylated product masses (EIC: 499.17; 515.16 ± 0.3 *m/z* $[M+H]^+$).

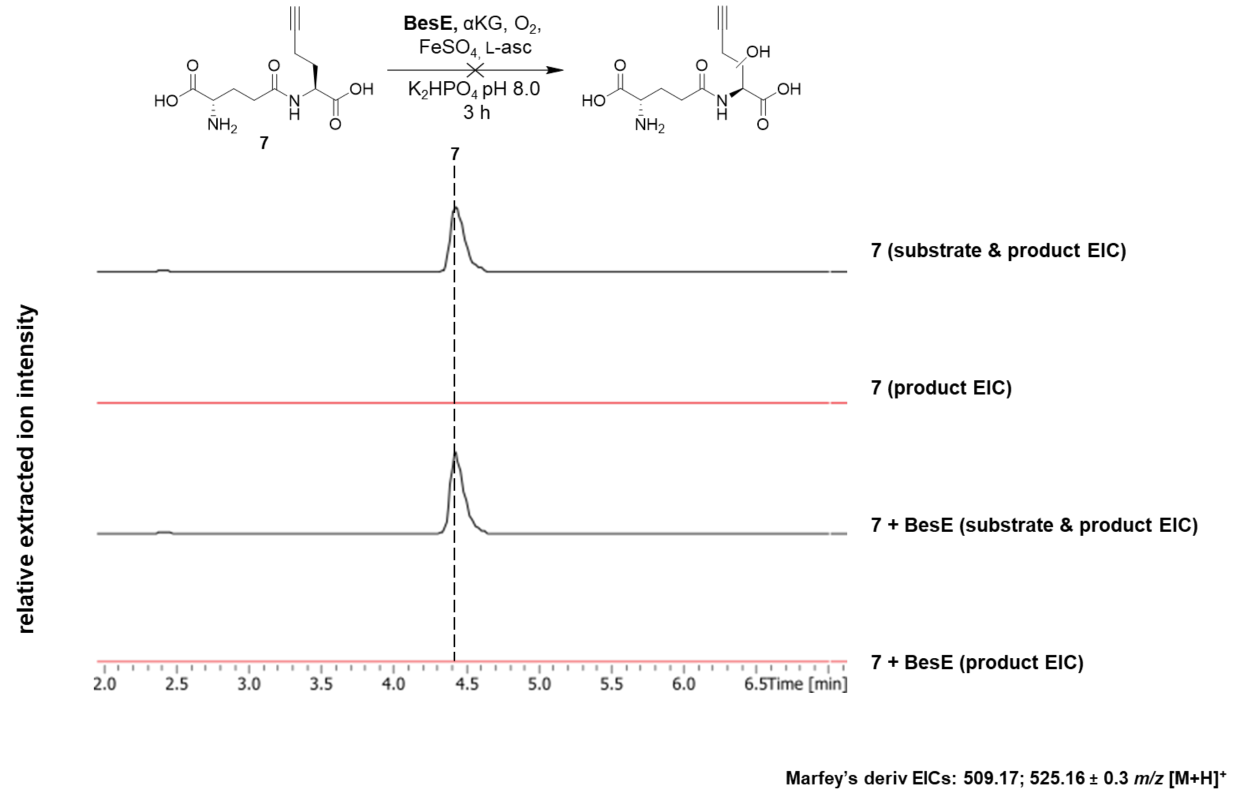


**Figure S10 – *in vitro* BesE assay with analog 7.** Assays analyzed by UHPLC-MS in positive mode post-Marfey's derivatization. Relative intensities were extracted for derivatized substrate and hydroxylated product masses (EIC: 509.17; 525.16 ± 0.3 *m/z* $[M+H]^+$).

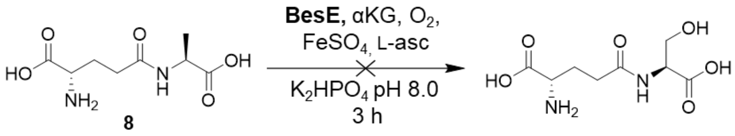


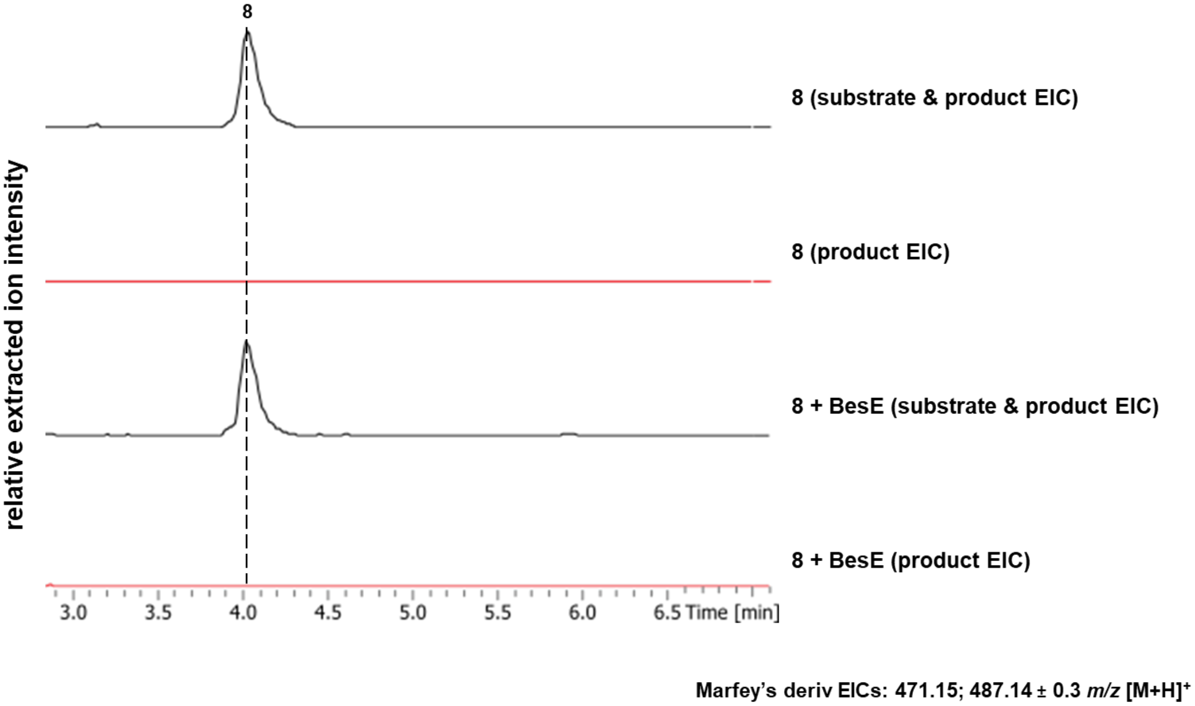


**Figure S11 – *in vitro* BesE assay with analog 8.** Assays analyzed by UHPLC-MS in positive mode post-Marfey's derivatization. Relative intensities were extracted for derivatized substrate and hydroxylated product masses (EIC: 471.15; 487.14 ± 0.3 *m/z* $[M+H]^+$).

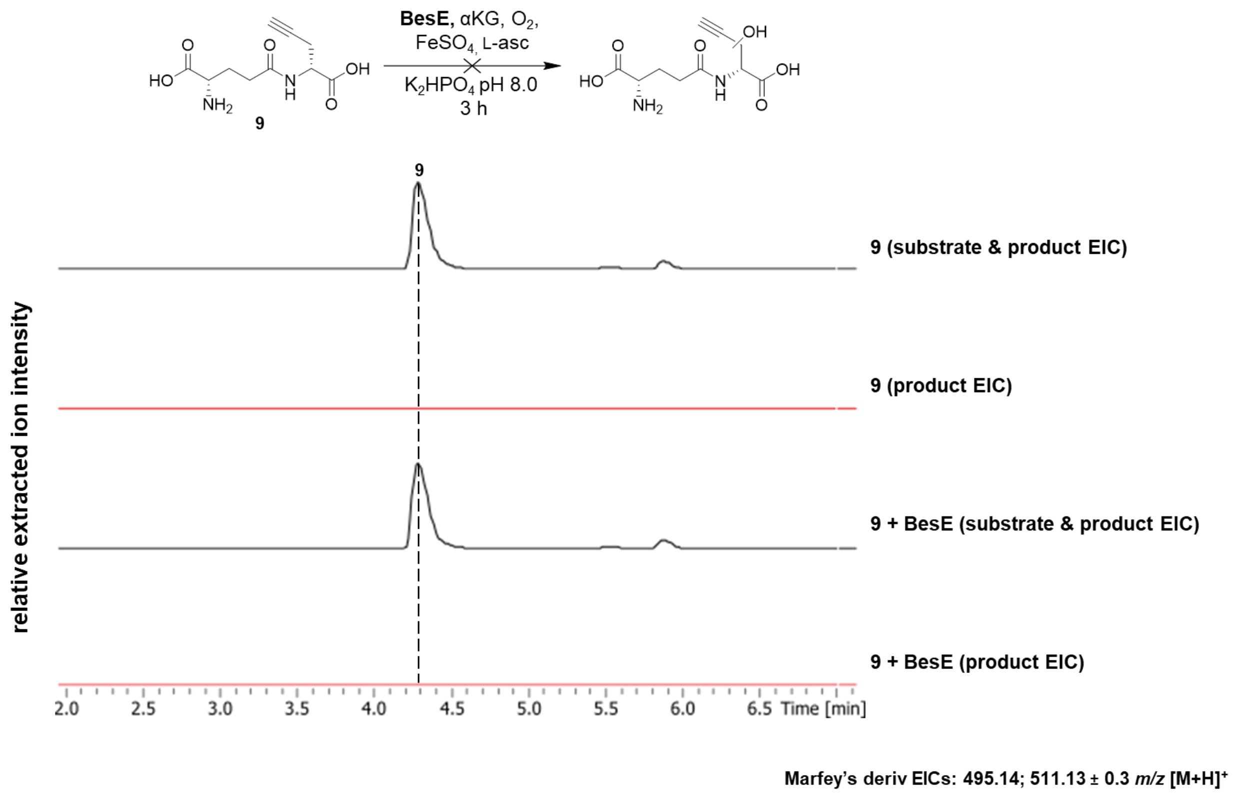


**Figure S12 – *in vitro* BesE assay with analog 9.** Assays analyzed by UHPLC-MS in positive mode post-Marfey's derivatization. Relative intensities were extracted for derivatized substrate and hydroxylated product masses (EIC: 495.14; 511.13 ± 0.3 *m/z* $[M+H]^+$).

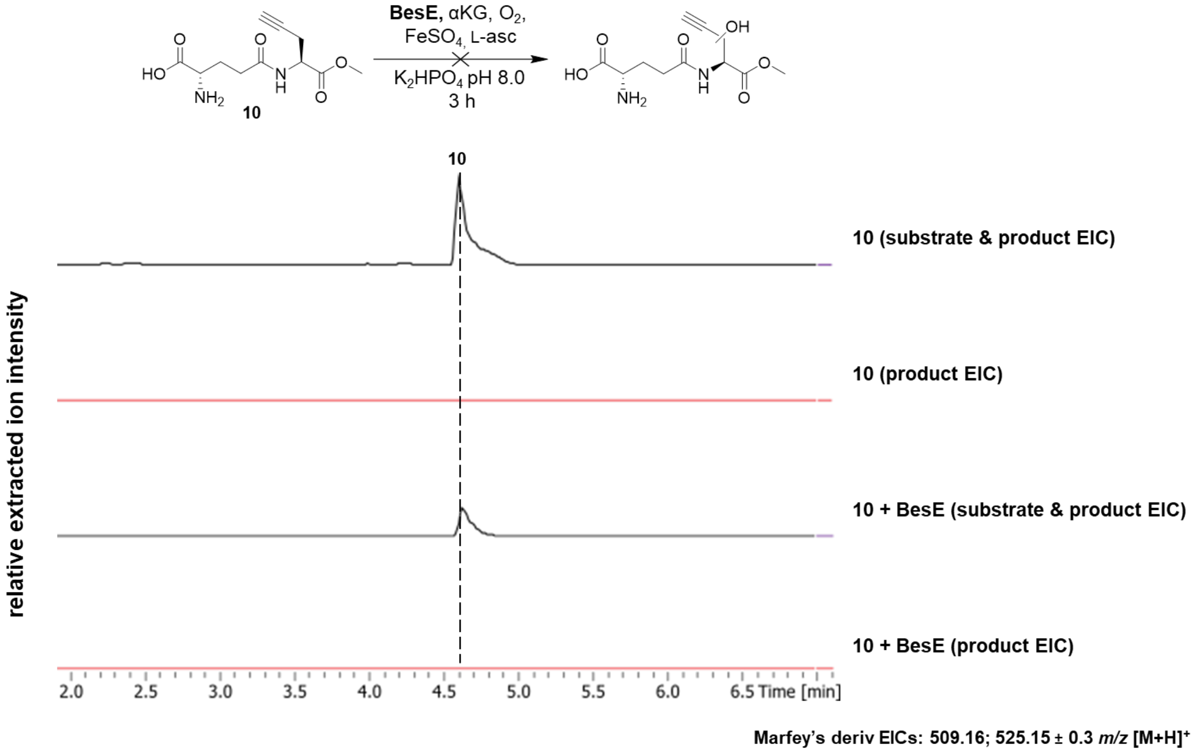


**Figure S13 – *in vitro* BesE assay with analog 10.** Assays analyzed by UHPLC-MS in positive mode post-Marfey's derivatization. Relative intensities were extracted for derivatized substrate and hydroxylated product masses (EIC: 509.16; 525.15 ± 0.3 *m/z* $[M+H]^+$).

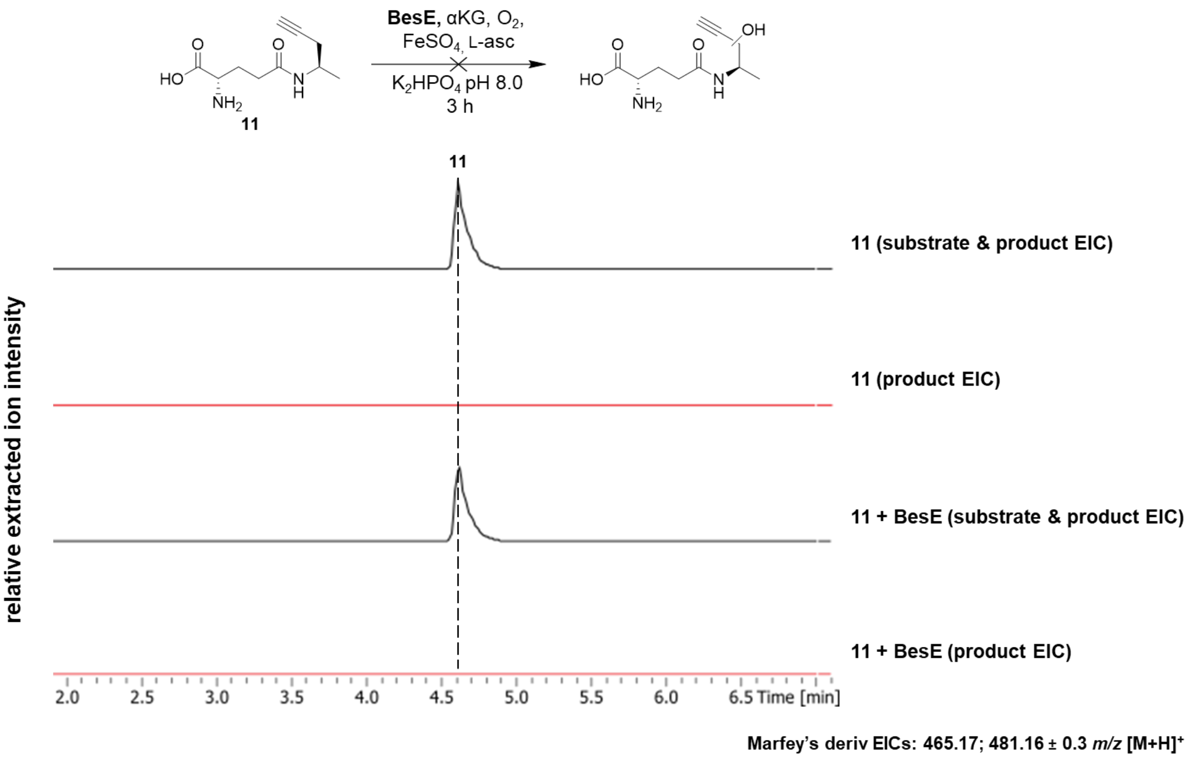


**Figure S14 – *in vitro* BesE assay with analog 11.** Assays analyzed by UHPLC-MS in positive mode post-Marfey's derivatization. Relative intensities were extracted for derivatized substrate and hydroxylated product masses (EIC: 465.17; 481.16 ± 0.3 *m/z* $[M+H]^+$).

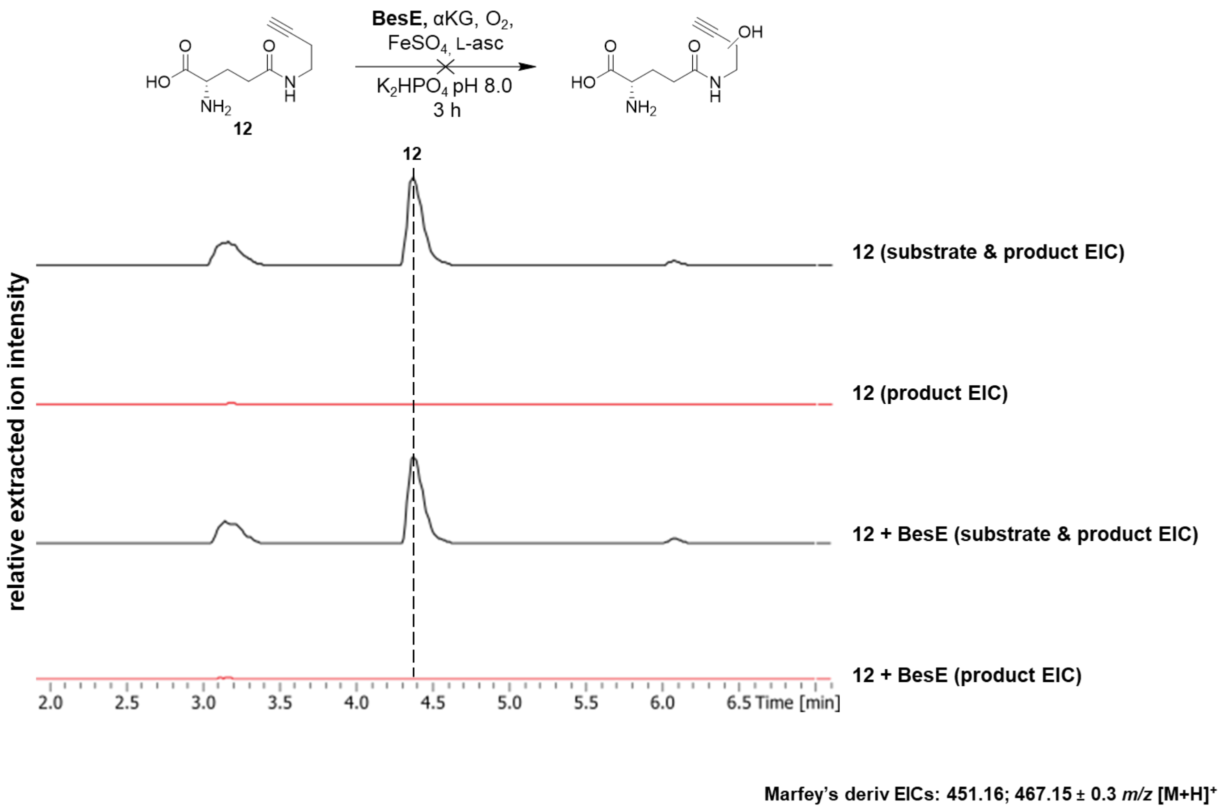


**Figure S15 – *in vitro* BesE assay with analog 12.** Assays analyzed by UHPLC-MS in positive mode post-Marfey's derivatization. Relative intensities were extracted for derivatized substrate and hydroxylated product masses (EIC: 451.16; 467.15 ± 0.3 *m/z* $[M+H]^+$).

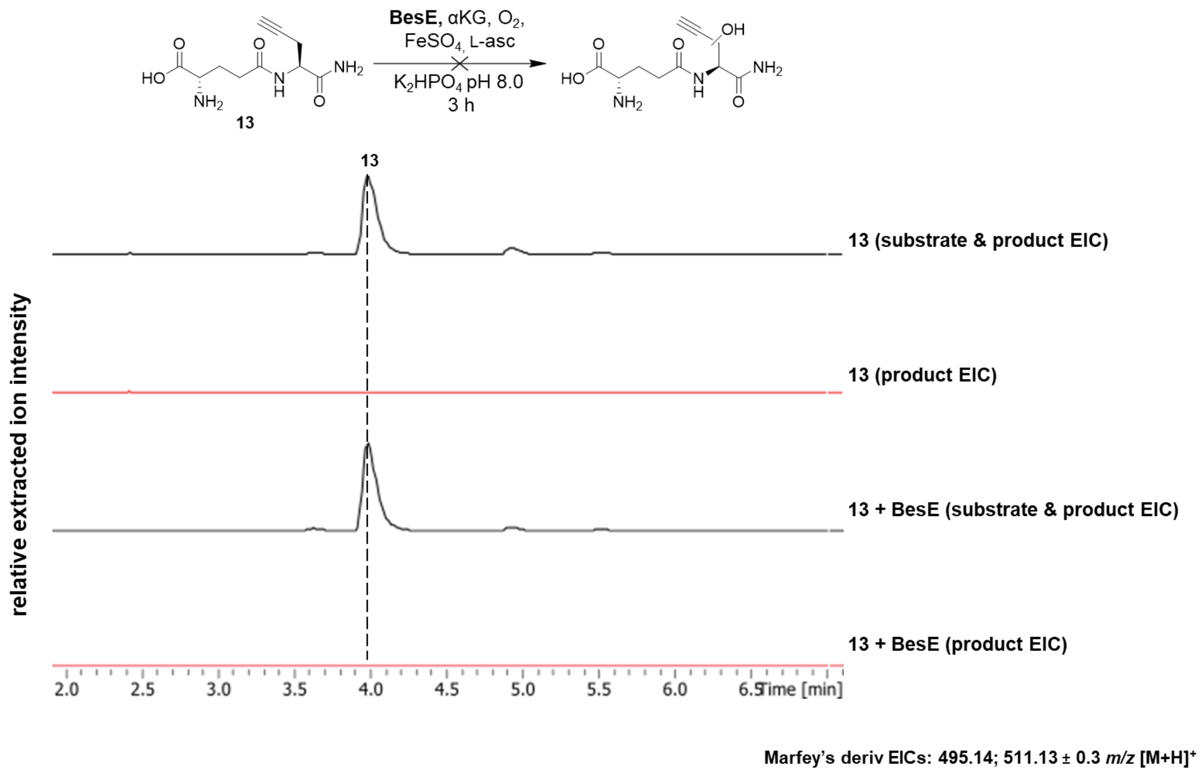


**Figure S16 – *in vitro* BesE assay with analog 13.** Assays analyzed by UHPLC-MS in positive mode post-Marfey's derivatization. Relative intensities were extracted for derivatized substrate and hydroxylated product masses (EIC: 495.14; 511.13 ± 0.3 *m/z* $[M+H]^+$).

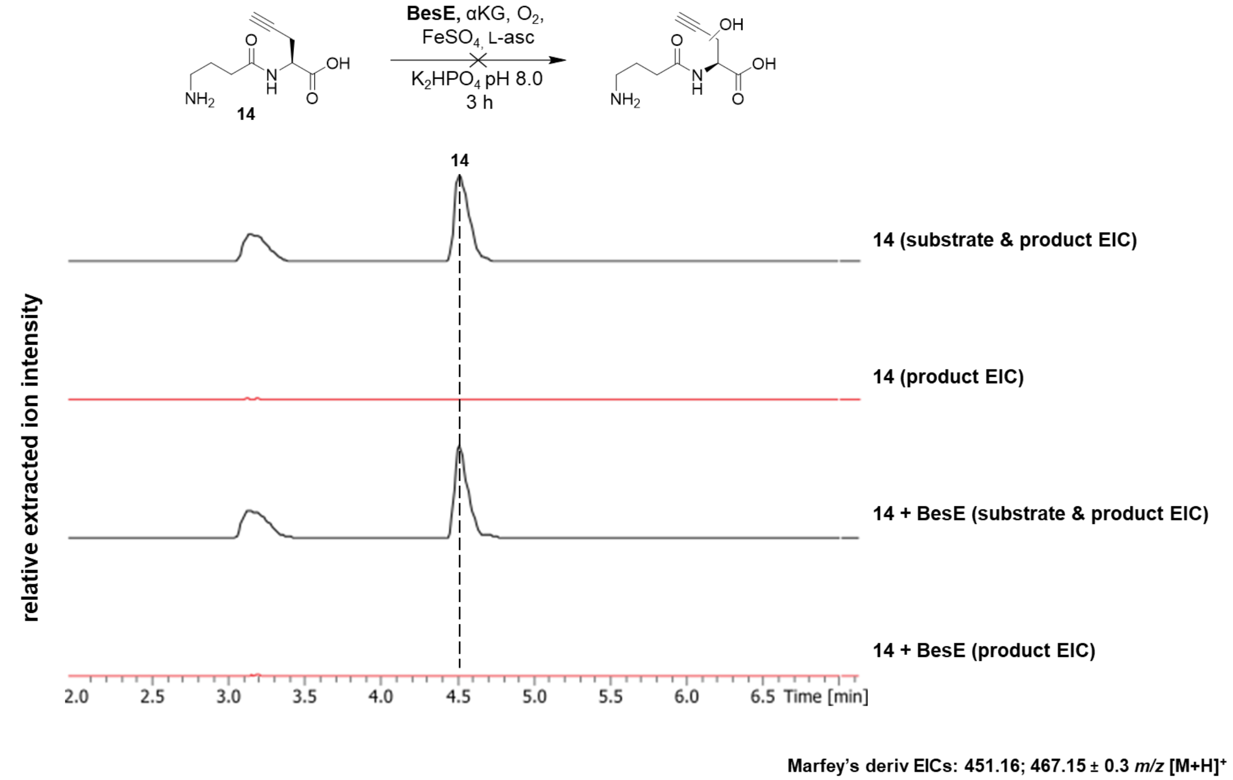


**Figure S17 – *in vitro* BesE assay with analog 14.** Assays analyzed by UHPLC-MS in positive mode post-Marfey's derivatization. Relative intensities were extracted for derivatized substrate and hydroxylated product masses (EIC: 451.16; 467.15 ± 0.3 *m/z* $[M+H]^+$).

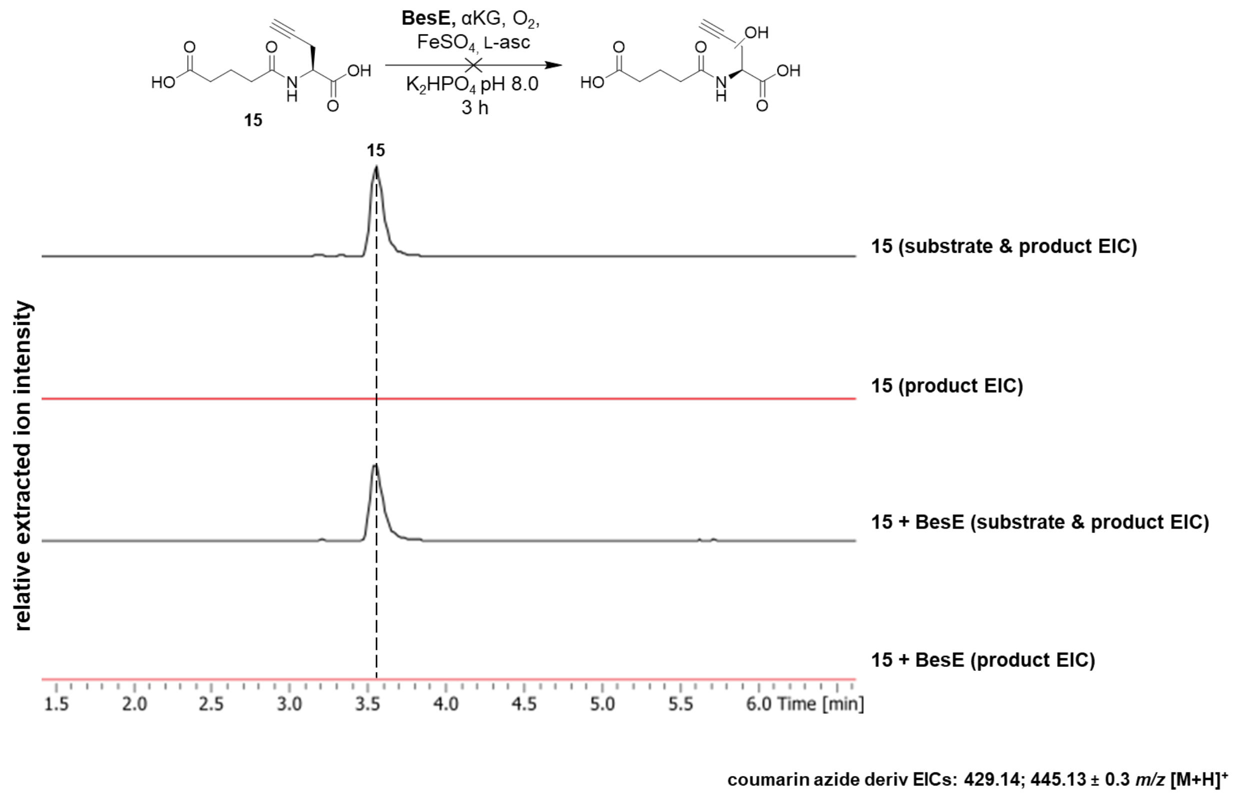


**Figure S18 – *in vitro* BesE assay with analog 15.** Assays analyzed by UHPLC-MS in positive mode post-CuAAC derivatization. Relative intensities were extracted for derivatized substrate and hydroxylated product masses (EIC: 429.14; 445.13 ± 0.3 *m/z* $[M+H]^+$).

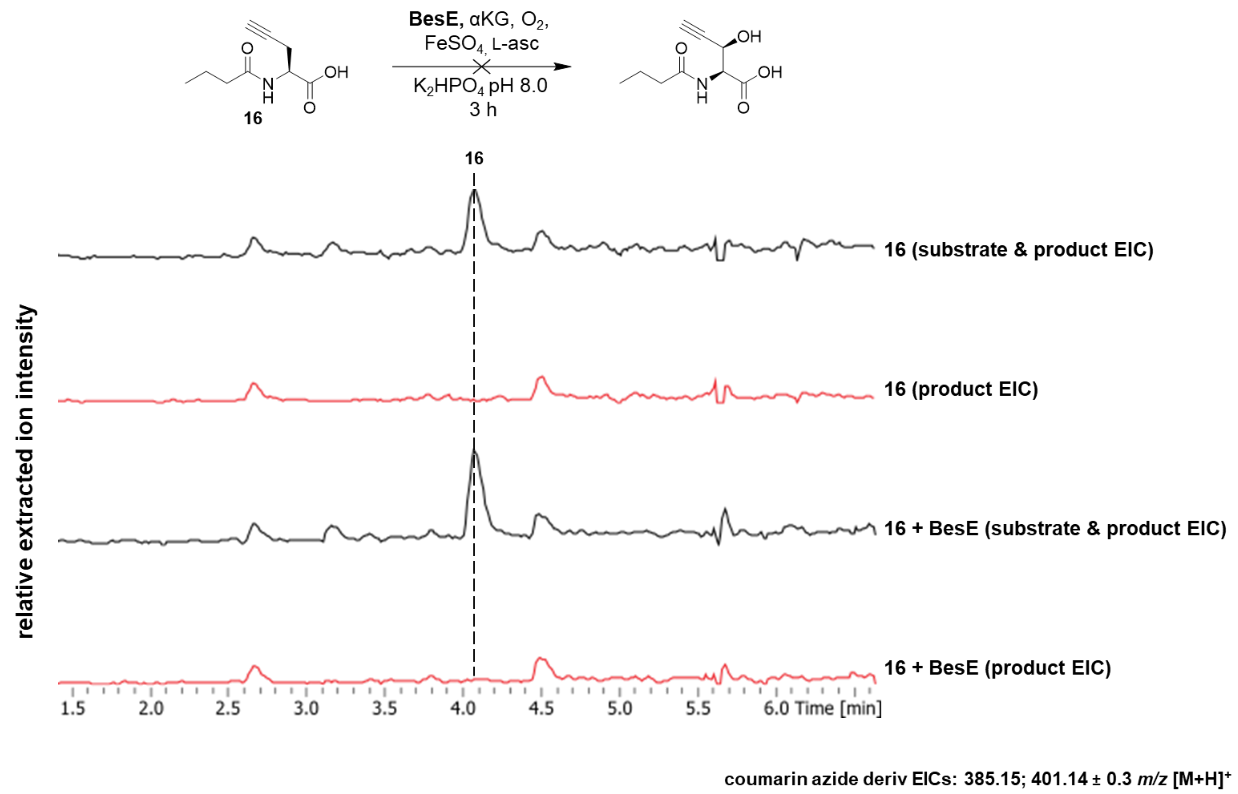


**Figure S19 – *in vitro* BesE assay with analog 16.** Assays analyzed by UHPLC-MS in positive mode post-CuAAC derivatization. Relative intensities were extracted for derivatized substrate and hydroxylated product masses (EIC: 385.15; 401.14 ± 0.3 *m/z* $[M+H]^+$).

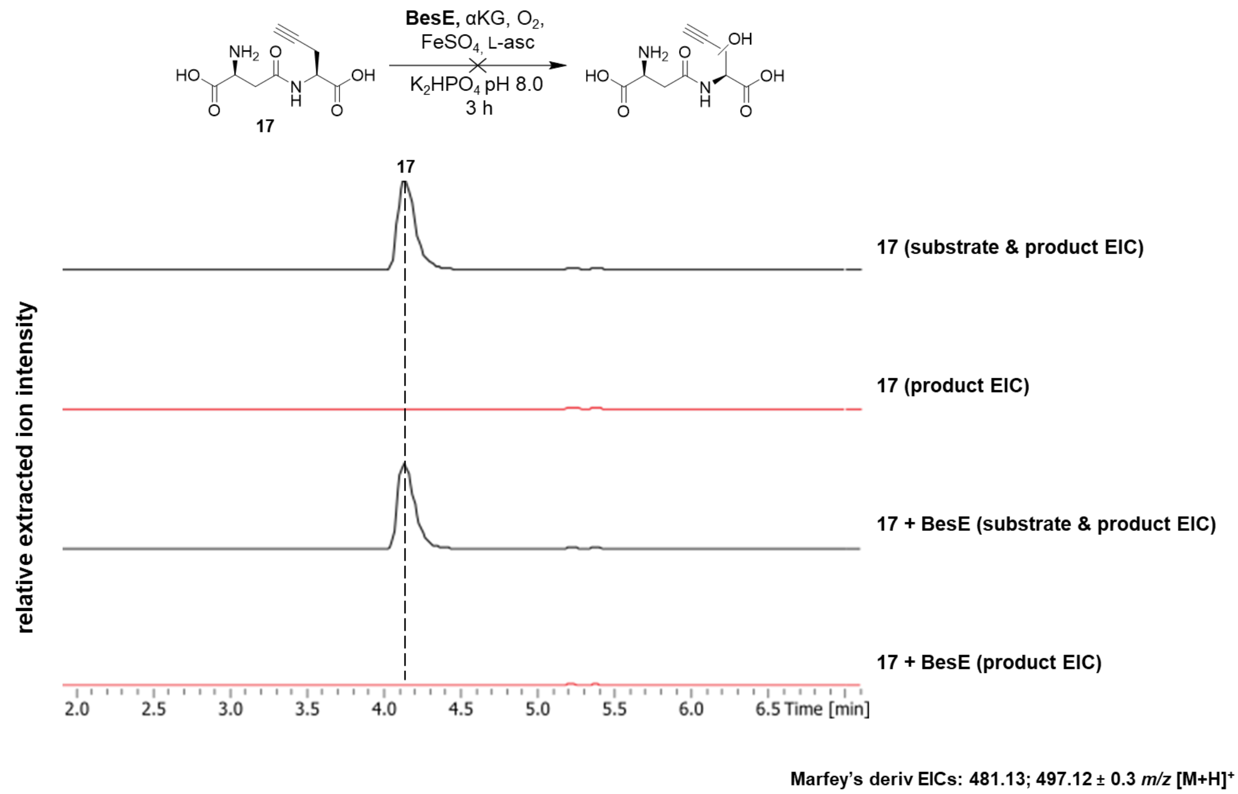


**Figure S20 – *in vitro* BesE assay with analog 17.** Assays analyzed by UHPLC-MS in positive mode post-Marfey's derivatization. Relative intensities were extracted for derivatized substrate and hydroxylated product masses (EIC: 481.13; 497.12 ± 0.3 *m/z* $[M+H]^+$).

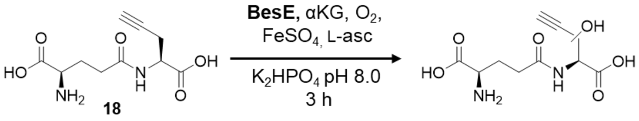


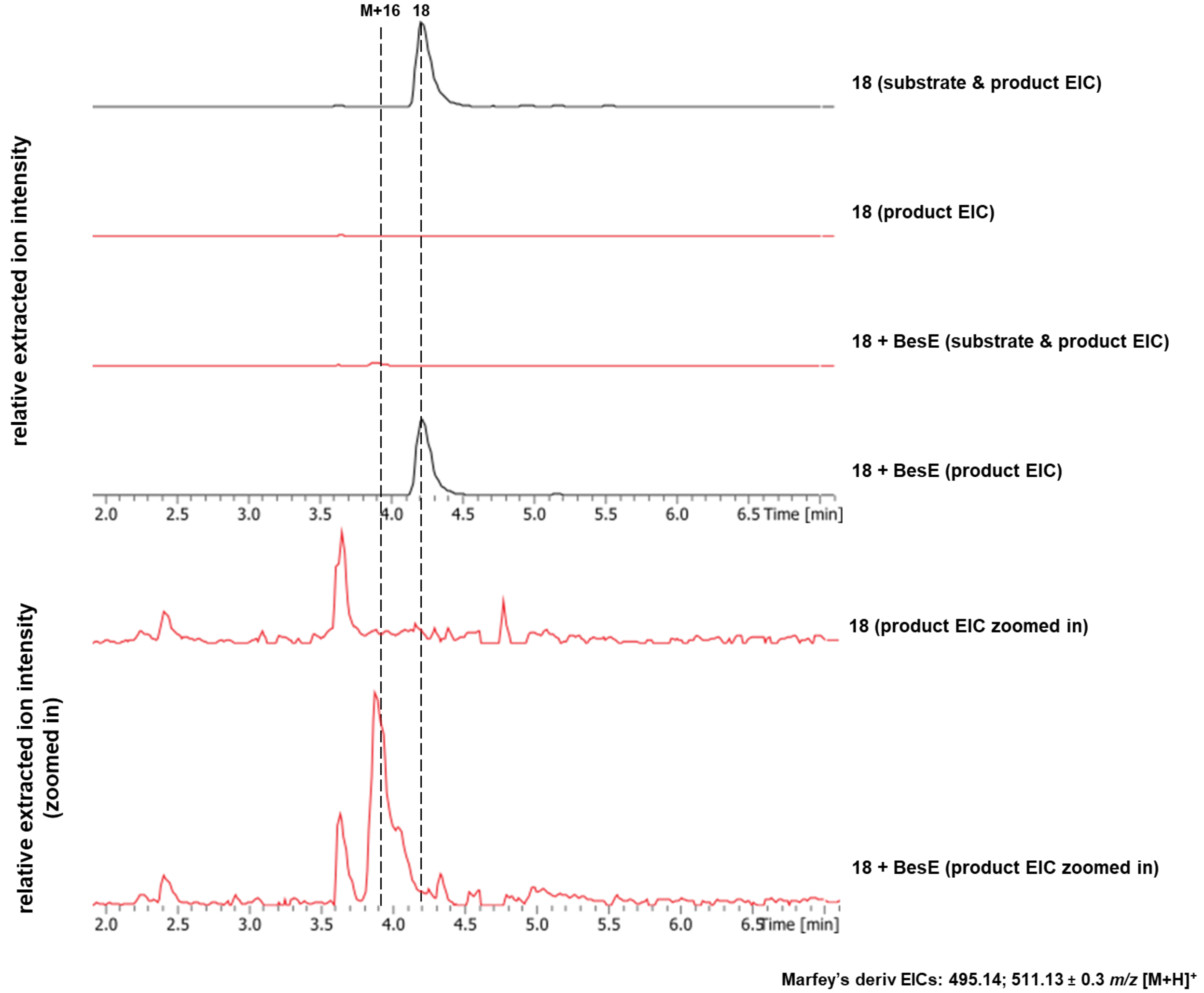


**Figure S21 – *in vitro* BesE assay with analog 18.** Additional product EICs for +/- enzyme conditions shown at a reduced scale to show the presence of a hydroxylated mass in the + BesE condition. Assays analyzed by UHPLC-MS in positive mode post-Marfey's derivatization. Relative intensities were extracted for derivatized substrate and hydroxylated product masses (EIC: 495.14; 511.13 ± 0.3 *m/z* $[M+H]^+$).

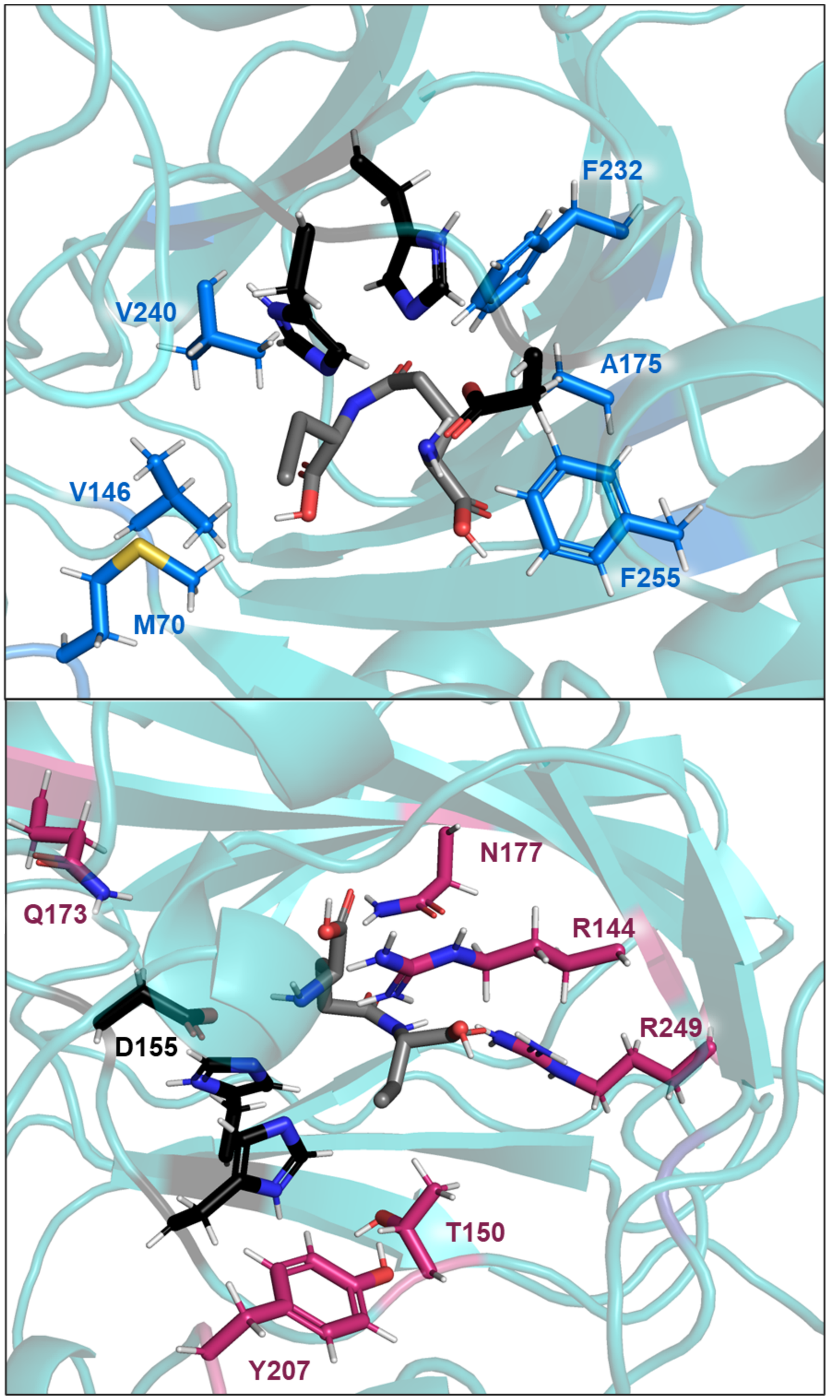


**Figure S22 – Structural depiction of BesE residues selected for site-directed mutagenesis.** Selected hydrophobic (top, blue) and polar (bottom, red) residues shown. Catalytic triad residues in black (selected residue D155 indicated in the bottom image). Docked substrate **1** in grey. All images were generated using PyMol, enzyme models were generated using AlphaFold 2.0,[1] substrate models were generated with Avogadro and substrate docking was performed with AutoDock 4.2.6.

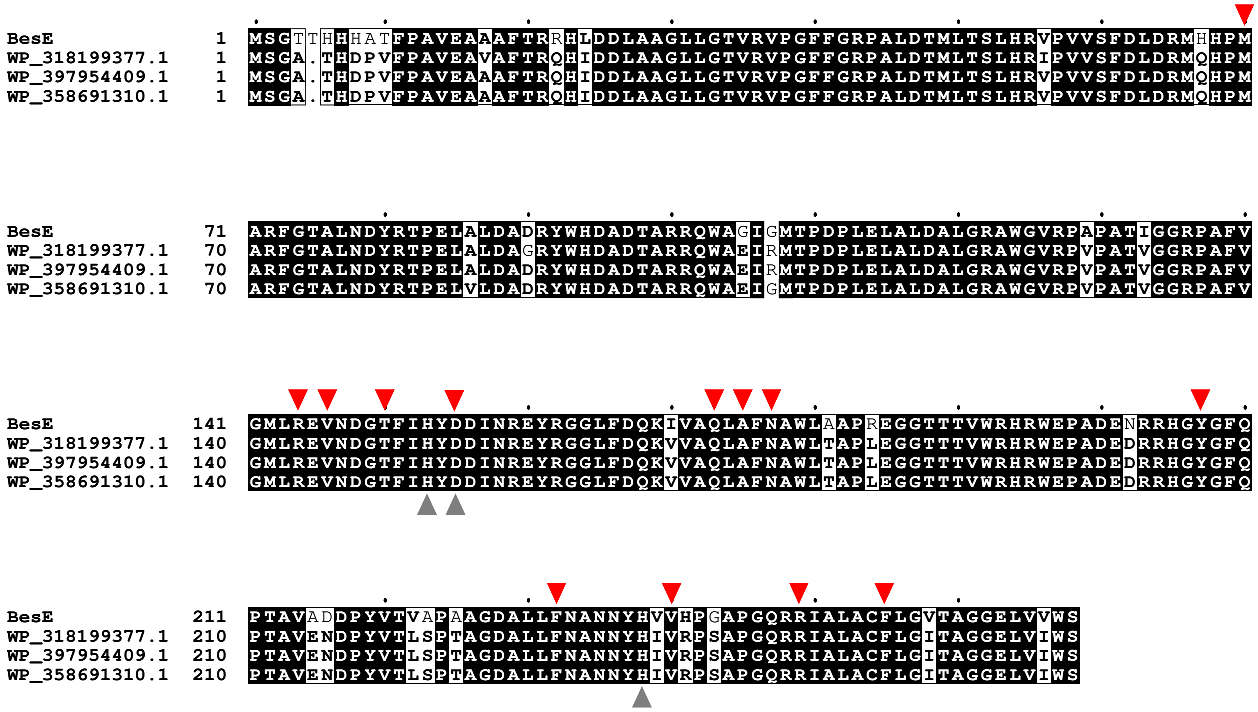


**Figure S23 – Conservation of BesE residues selected for site-directed mutagenesis across putative actinobacterial homologs.** Sequence alignment of BesE homologs from other actinobacteria. The initial sequence alignment was prepared with ClustalOmega and was further processed and refined in EsPript. BesE from *Streptomyces cattleya* (renamed to *Streptantibioticus cattleyicolor*, WP_014151498.1) was compared to three putative uncharacterized homologs annotated as 2OG-Fe(II)-dependent halogenase WelO5 family proteins from: *Streptomyces* sp. NPDC017638 (WP_397954409.1); *Streptomyces* sp. SCL15-4 (WP_318199377.1); and *Streptomyces* sp. NPDC006658 (WP_358691310.1). Active site residues in the catalytic triad are marked with grey arrows under the alignment, residues selected for site-directed mutagenesis are marked with red arrows above the alignment.

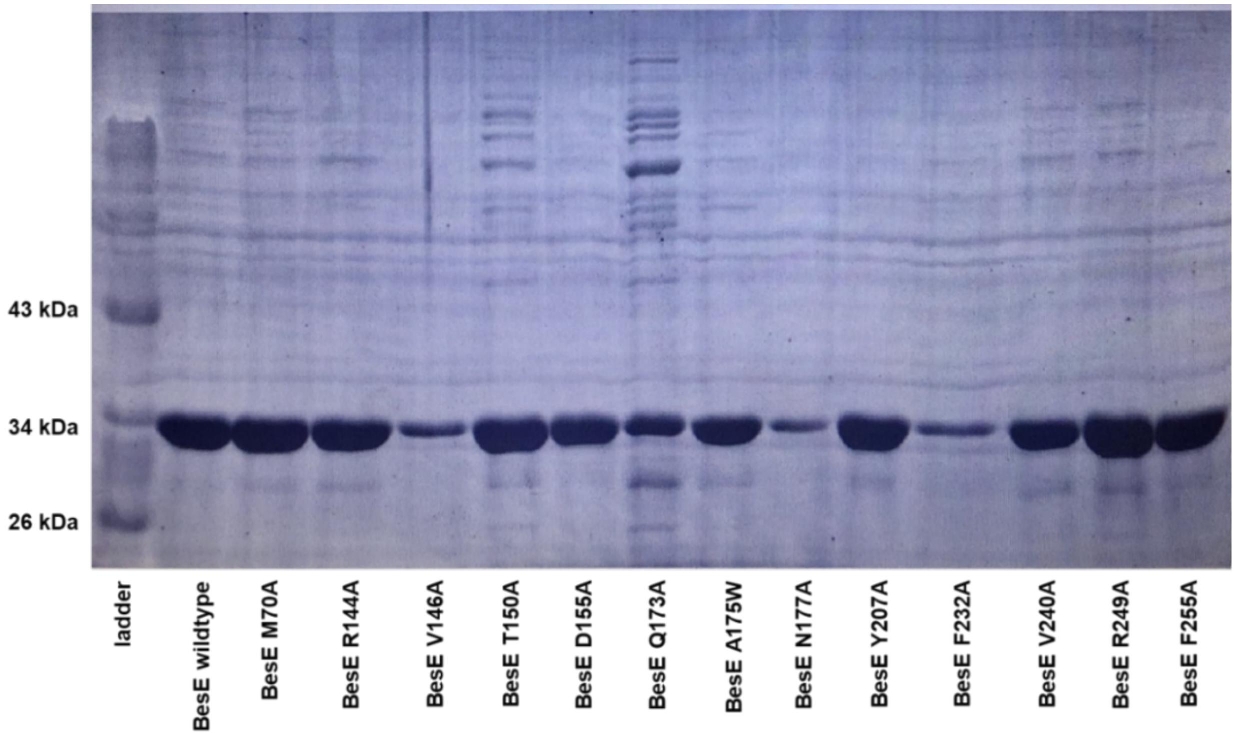


**Figure S24 – 10% SDS-PAGE gel of purified BesE mutants.** Approximately 5 μg of each purified BesE mutant was loaded per sample except for BesE mutants V146A, N177A, and F232A due to these samples being at much lower concentrations (less than 1 μg loaded for each)

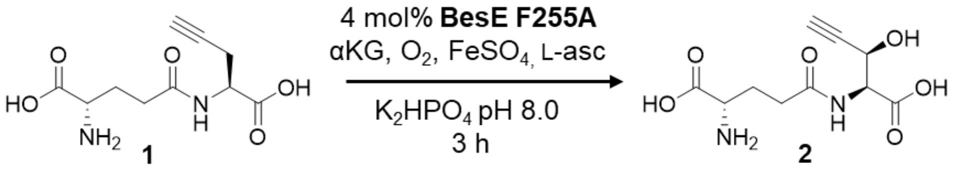


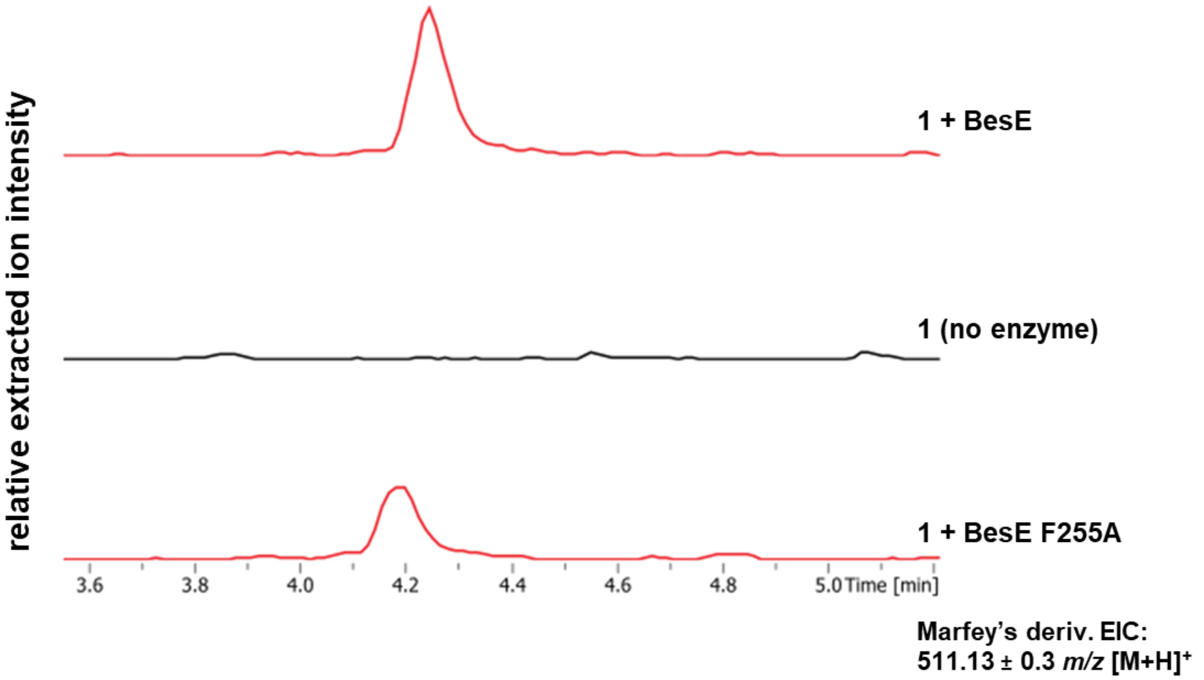


**Figure S25 – Representative trace of *in vitro* BesE F255A assay with 1.** Assays conducted in quadruplicate and analyzed by UHPLC-MS in positive mode post-Marfey's derivatization. Extracted ion chromatograms for hydroxylated product masses shown for BesE F255A assay as well as BesE wild-type and no enzyme negative control assays for comparison (EIC: 511.13 ± 0.3 *m/z* $[M+H]^+$).

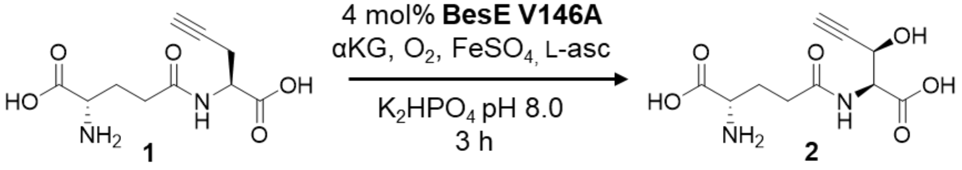


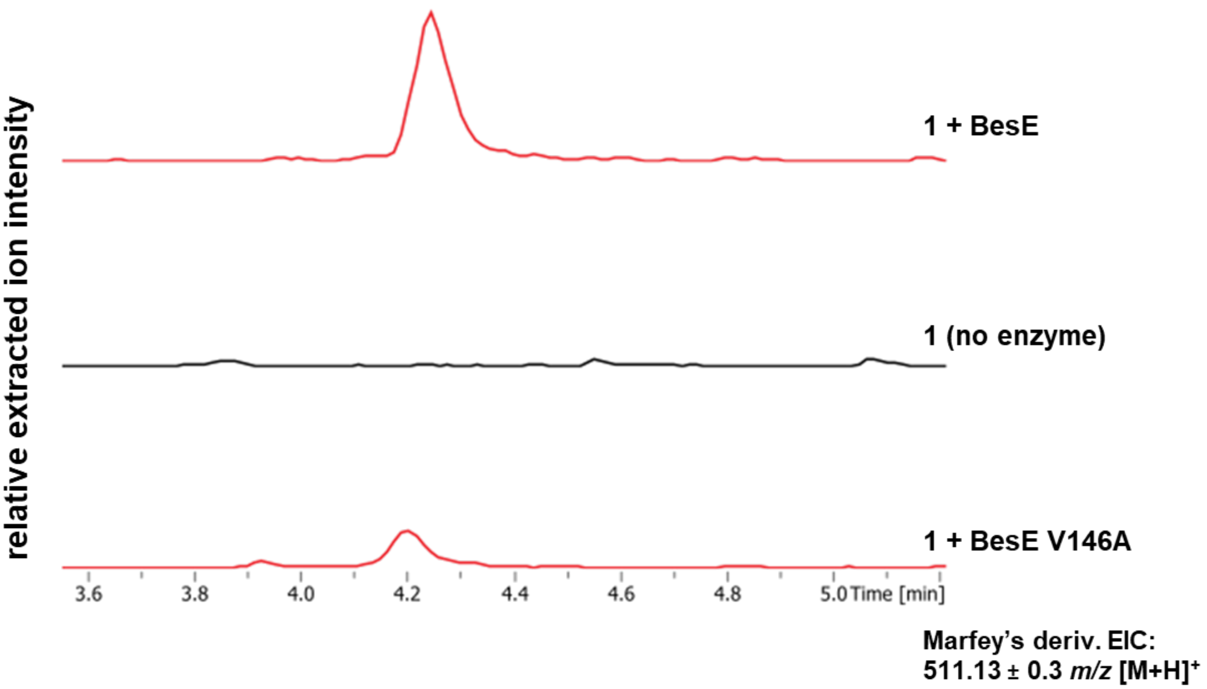


**Figure S26 – Representative trace of *in vitro* BesE V146A assay with 1.** Assays conducted in quadruplicate and analyzed by UHPLC-MS in positive mode post-Marfey's derivatization. Extracted ion chromatograms for hydroxylated product masses shown for BesE V146A assay as well as BesE wild-type and no enzyme negative control assays for comparison (EIC: 511.13 ± 0.3 *m/z* $[M+H]^+$).

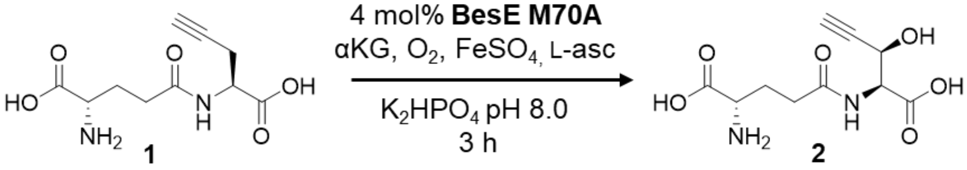


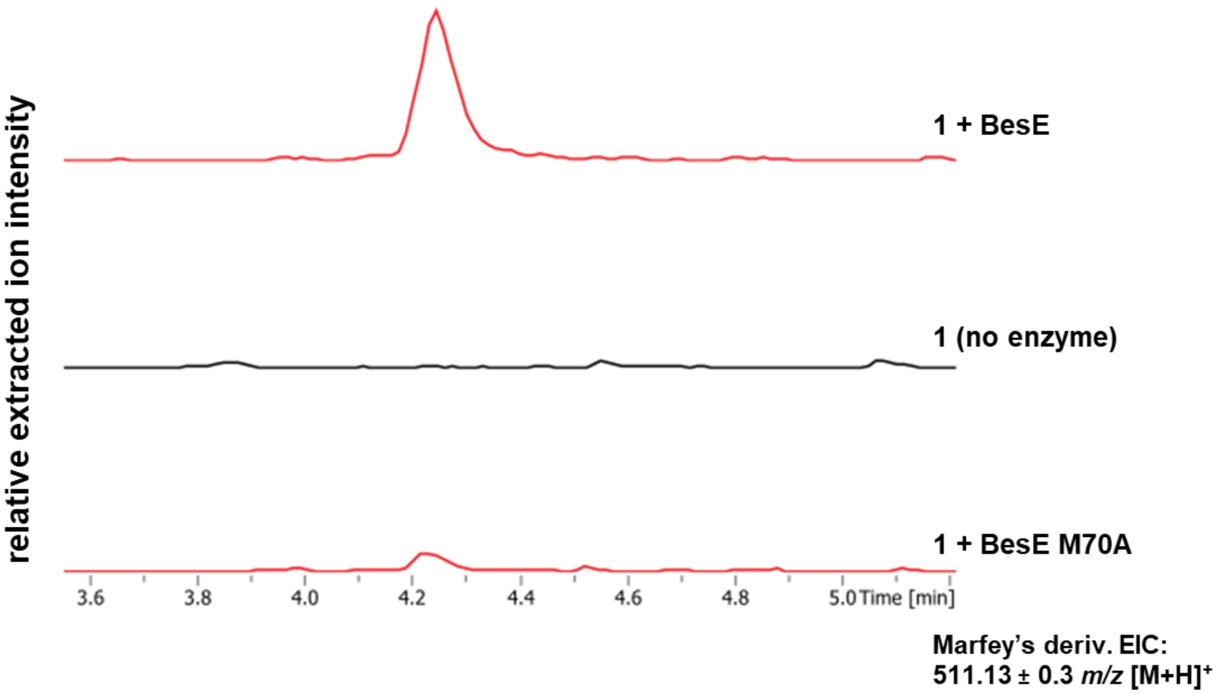


**Figure S27 – Representative trace of *in vitro* BesE M70A assay with 1.** Assays conducted in quadruplicate and analyzed by UHPLC-MS in positive mode post-Marfey's derivatization. Extracted ion chromatograms for hydroxylated product masses shown for BesE M70A assay as well as BesE wild-type and no enzyme negative control assays for comparison (EIC: 511.13 ± 0.3 *m/z* $[M+H]^+$).

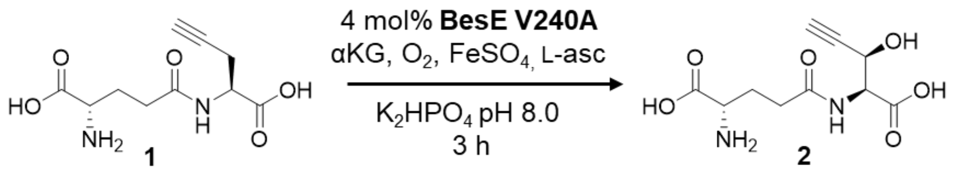


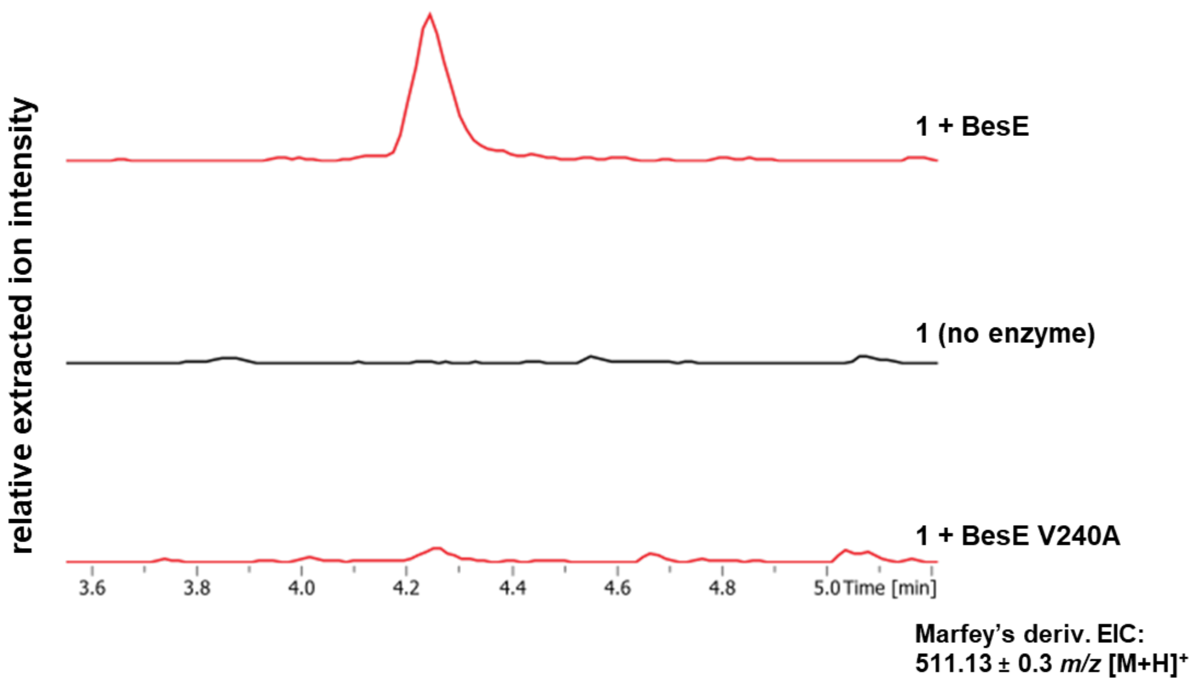


**Figure S28 – Representative trace of *in vitro* BesE V240A assay with 1.** Assays conducted in quadruplicate and analyzed by UHPLC-MS in positive mode post-Marfey's derivatization. Extracted ion chromatograms for hydroxylated product masses shown for BesE V240A assay as well as BesE wild-type and no enzyme negative control assays for comparison (EIC: 511.13 ± 0.3 *m/z* $[M+H]^+$).

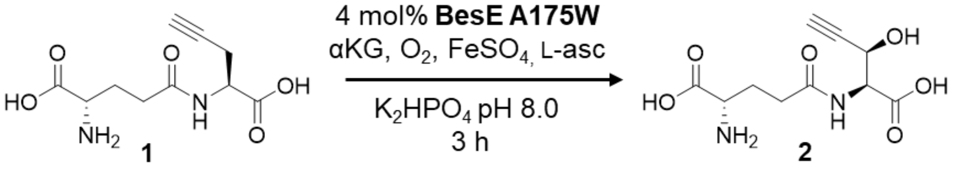


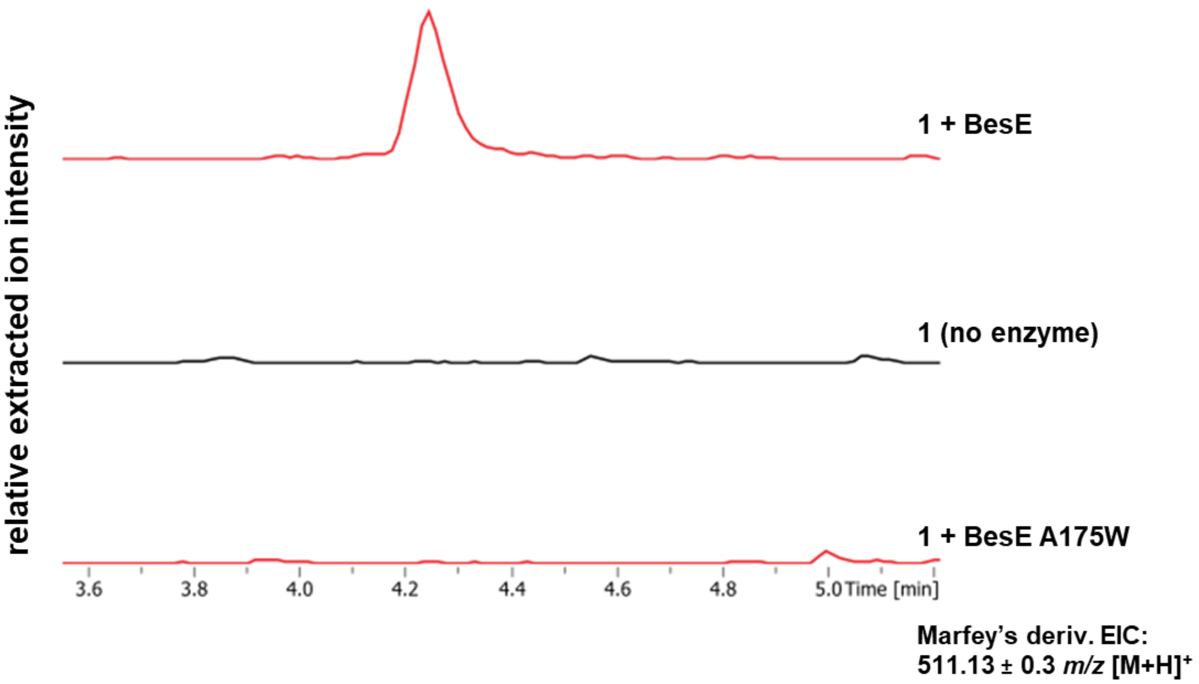


**Figure S29 – Representative trace of *in vitro* BesE A175W assay with 1.** Assays conducted in quadruplicate and analyzed by UHPLC-MS in positive mode post-Marfey's derivatization. Extracted ion chromatograms for hydroxylated product masses shown for BesE A175W assay as well as BesE wild-type and no enzyme negative control assays for comparison (EIC: 511.13 ± 0.3 *m/z* $[M+H]^+$).

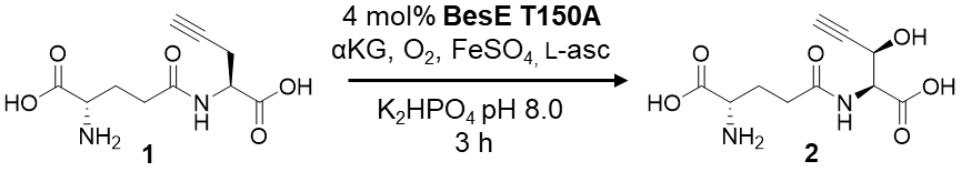


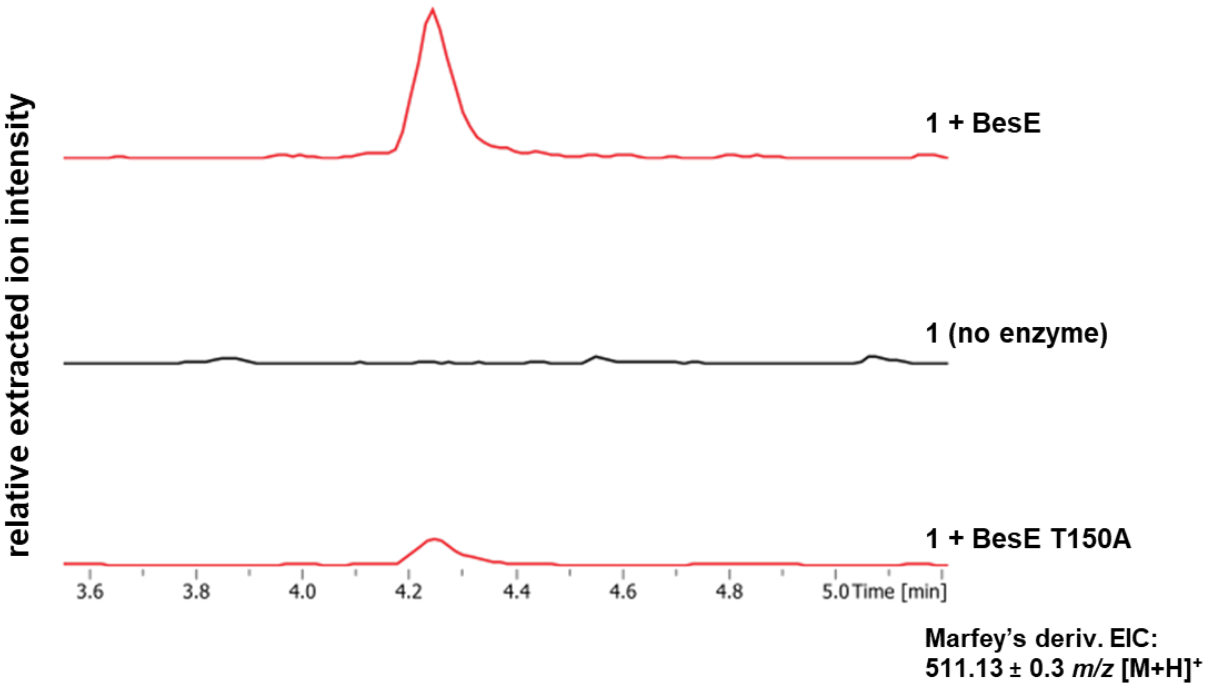


**Figure S30 – Representative trace of *in vitro* BesE T150A assay with 1.** Assays conducted in quadruplicate and analyzed by UHPLC-MS in positive mode post-Marfey's derivatization. Extracted ion chromatograms for hydroxylated product masses shown for BesE A175W assay as well as BesE wild-type and no enzyme negative control assays for comparison (EIC: 511.13 ± 0.3 *m/z* $[M+H]^+$).

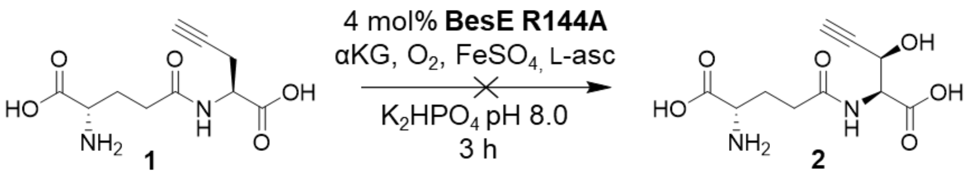


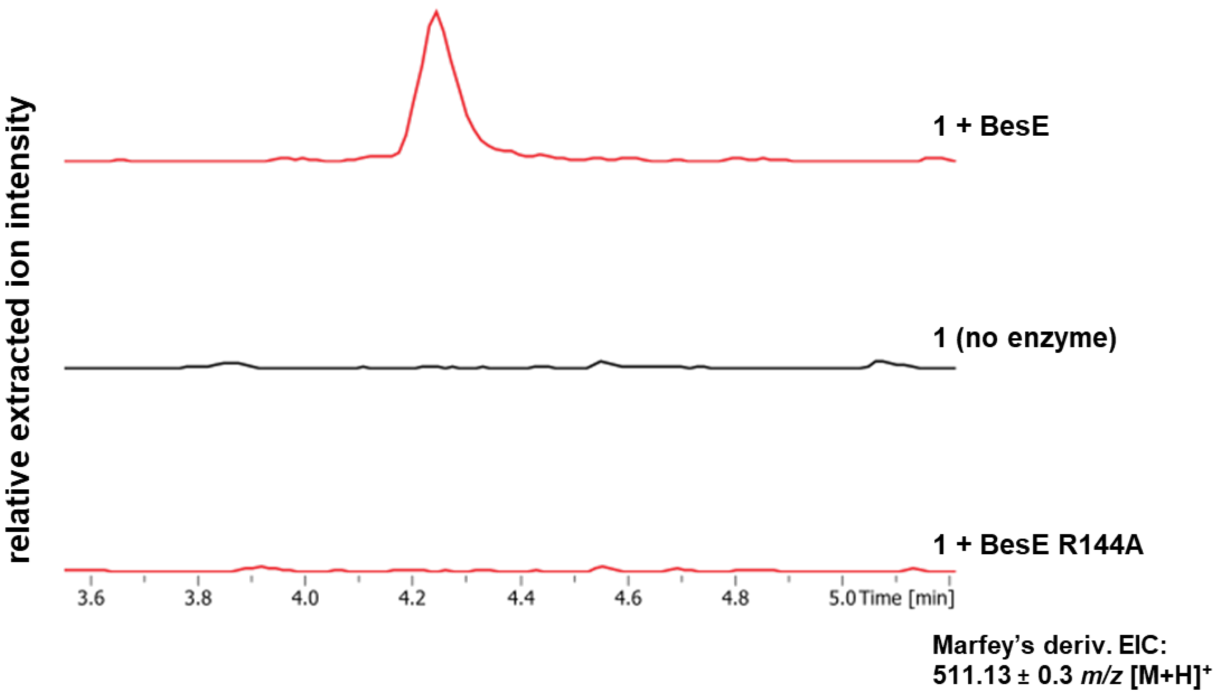


**Figure S31 – Representative trace of *in vitro* BesE R144A assay with 1.** Assays conducted in quadruplicate and analyzed by UHPLC-MS in positive mode post-Marfey's derivatization. Extracted ion chromatograms for hydroxylated product masses shown for BesE R144A assay as well as BesE wild-type and no enzyme negative control assays for comparison (EIC: 511.13 ± 0.3 *m/z* $[M+H]^+$).

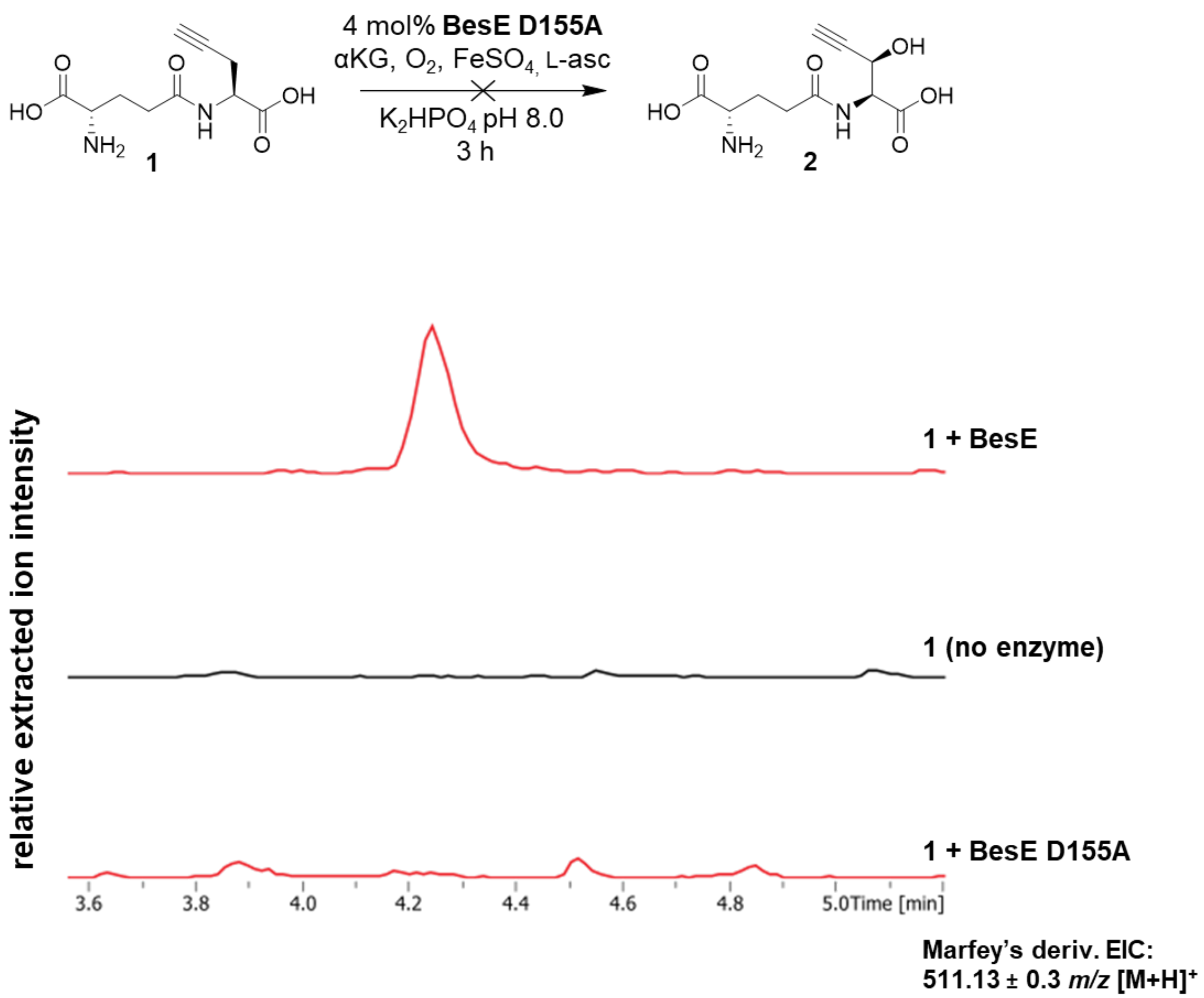


**Figure S32 – Representative trace of *in vitro* BesE D155A assay with 1.** Assays conducted in quadruplicate and analyzed by UHPLC-MS in positive mode post-Marfey's derivatization. Extracted ion chromatograms for hydroxylated product masses shown for BesE D155A assay as well as BesE wild-type and no enzyme negative control assays for comparison (EIC: 511.13 ± 0.3 *m/z* $[M+H]^+$).

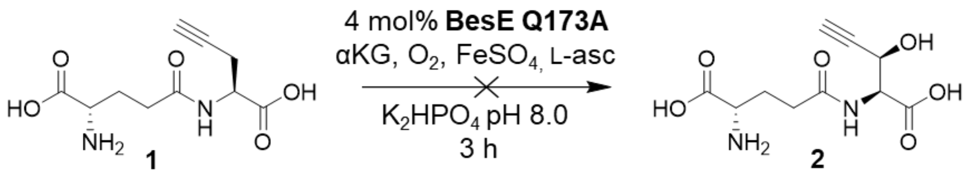


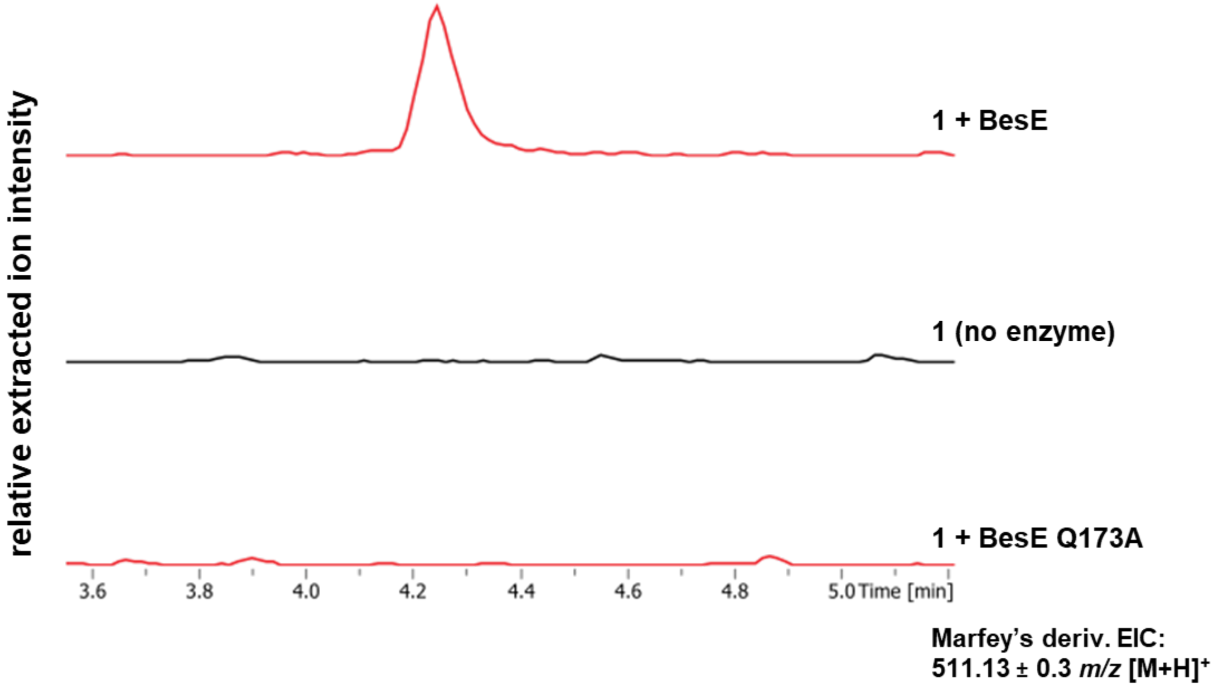


**Figure S33 – Representative trace of *in vitro* BesE Q173A assay with 1.** Assays conducted in quadruplicate and analyzed by UHPLC-MS in positive mode post-Marfey's derivatization. Extracted ion chromatograms for hydroxylated product masses shown for BesE Q173A assay as well as BesE wild-type and no enzyme negative control assays for comparison (EIC: 511.13 ± 0.3 *m/z* $[M+H]^+$).

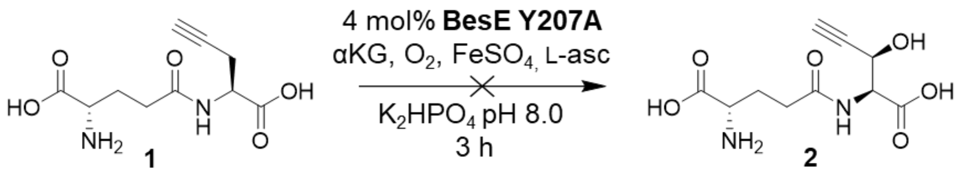


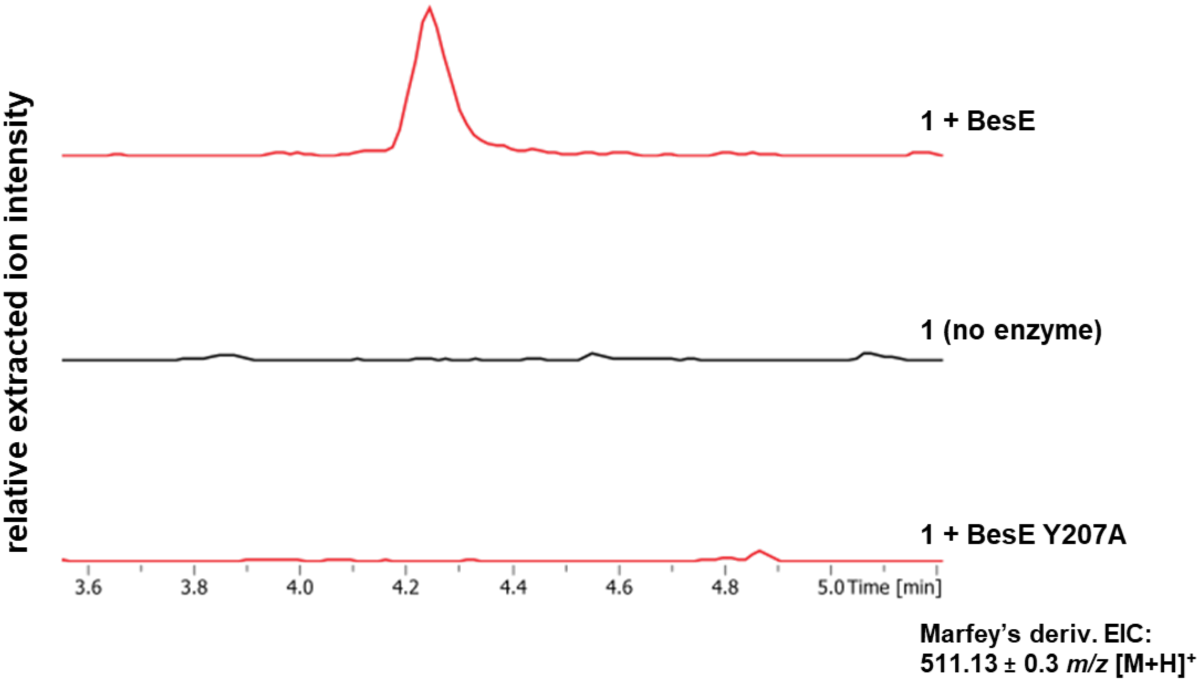


**Figure S34 – Representative trace of *in vitro* BesE Y207A assay with 1.** Assays conducted in quadruplicate and analyzed by UHPLC-MS in positive mode post-Marfey's derivatization. Extracted ion chromatograms for hydroxylated product masses shown for BesE Y207A assay as well as BesE wild-type and no enzyme negative control assays for comparison (EIC: 511.13 ± 0.3 *m/z* $[M+H]^+$).

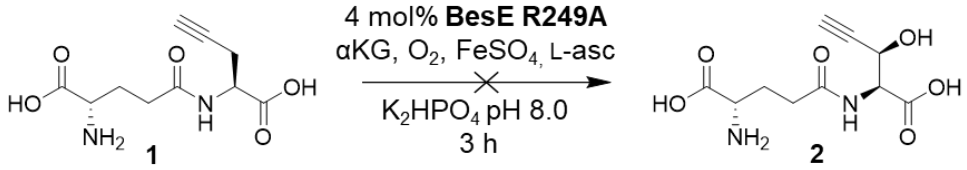


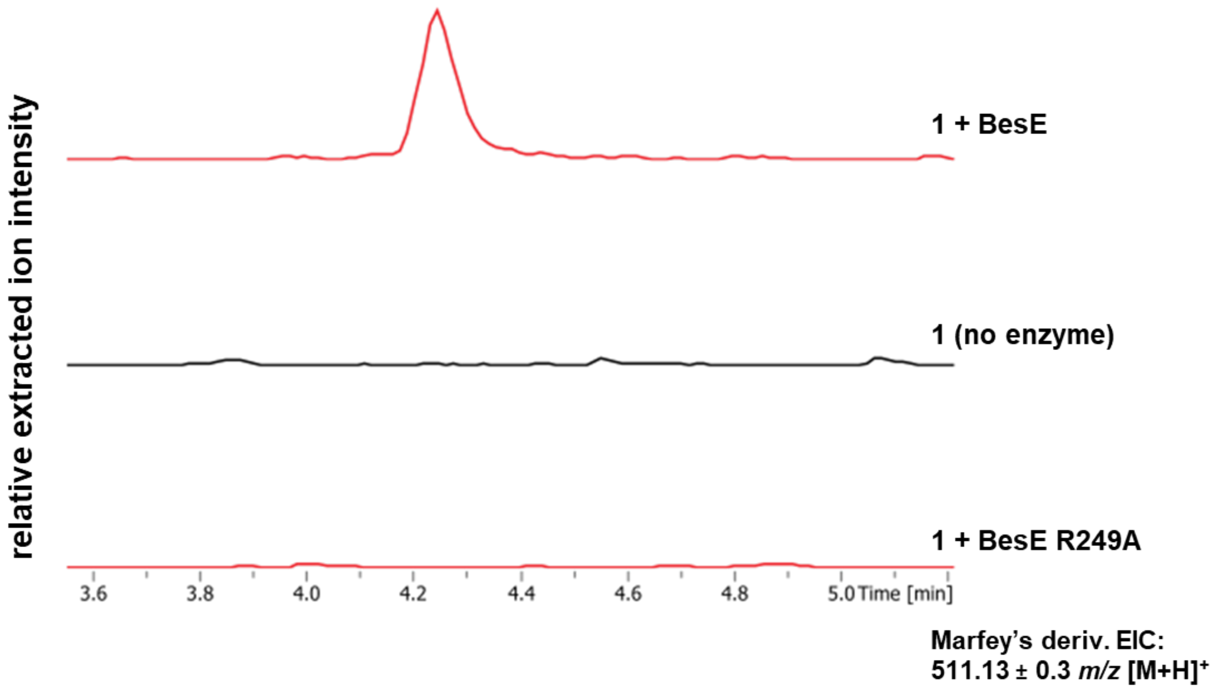


**Figure S35 – Representative trace of *in vitro* BesE R249A assay with 1.** Assays conducted in quadruplicate and analyzed by UHPLC-MS in positive mode post-Marfey's derivatization. Extracted ion chromatograms for hydroxylated product masses shown for BesE R249A assay as well as BesE wild-type and no enzyme negative control assays for comparison (EIC: 511.13 ± 0.3 *m/z* $[M+H]^+$).

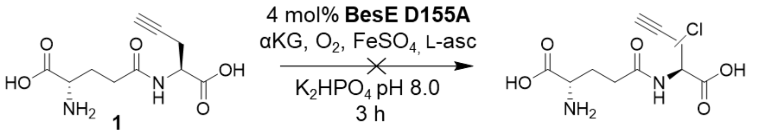


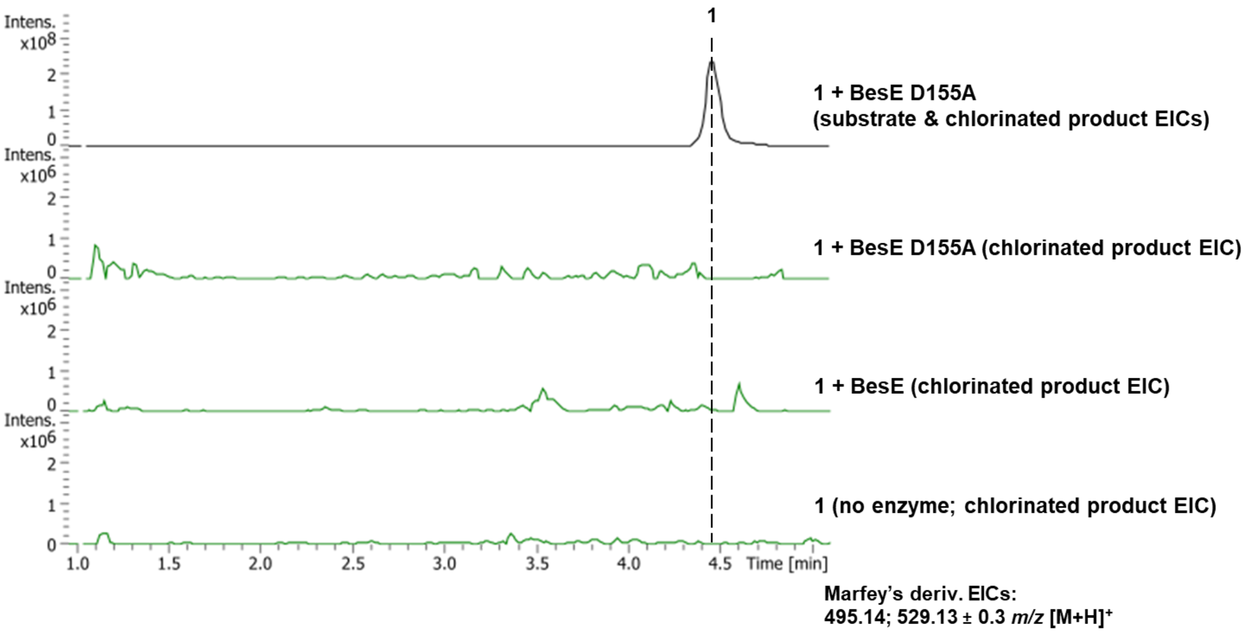


**Figure S36 – *in vitro* BesE D155A assay with 1 is not indicative of chlorinated product formation.** Assays conducted in quadruplicate and analyzed by UHPLC-MS in positive mode post-Marfey's derivatization. Chromatogram trace extracted for both substrate (EIC: 495.14 ± 0.3 *m/z* $[M+H]^+$) and chlorinated product (EIC: 529.13 ± 0.3 *m/z* $[M+H]^+$) ion masses shown for BesE D155A assay followed by traces extracted only for chlorinated product for BesE D155A, BesE wild-type and no enzyme negative control assays at a reduced relative extracted ion intensity range. No detectable signals suggestive of a chlorinated product ion were observed.

#### Chemical Synthesis

**General reaction procedures for BesE native substrate and substrate analog synthesis.**

BesE substrate general reaction A: methyl esterification of L-propargylglycine and analogs.[1,2]

Approximately 0.1 g of L-propargylglycine or analog was suspended in 10 mL of 2,2-dimethoxypropane. To the suspension was added 1 mL of concentrated (12 N) HCl. The mixture was left stirring at room temperature (RT) overnight (~16-18 h). The mixture was concentrated via rotary evaporation at 50 °C. The dark brown residue was resuspended in 10 mL water and washed with EtOAc 3 x 10 mL. The aqueous layer was frozen in a dry ice/acetone bath and lyophilized to afford the product without further purification.

BesE substrate general reaction B: coupling reaction with Boc-L-glutamic acid-α-tert-butyl-ester or alternative coupling compound.[3]

To approximately 0.05 g of *O*-methyl-L-propargylglycine or analog stirring in 10 mL $CH_2Cl_2$ was added boc-L-glutamic acid-α-tert-butyl-ester or alternative coupling compound (1 eq.) and HATU (1.05 eq.). Triethylamine (6 eq.) was added dropwise, and the reaction was left stirring at room temperature overnight (16-18 h). The reaction mixture was transferred to a separatory funnel and washed with saturated $NH_4Cl$ (2 x 10 mL), water (10 mL), and brine (10 mL). The organic layer was dried over $MgSO_4$, filtered, concentrated *in vacuo* and purified via silica flash chromatography (2:1 hexanes/EtOAc).

BesE substrate general reaction C: methyl ester deprotection.[4]

To approximately 0.035 g of Boc-L-Blu-OtBu-L-Pra-OMe or similarly protected analog in 2.5 mL tetrahydrofuran (THF) at 0 °C was added an aqueous solution of 1N LiOH at once. The reaction was brought to room temperature after 15 min and stirred for an additional 2 h. The reaction was concentrated in vacuo to remove THF. The LiOH was neutralized by addition of aqueous 1N HCl to a pH of 4. Organic compounds were extracted with EtOAc (3 x 10 mL). The organic layer was dried over $MgSO_4$, filtered, and concentrated *in vacuo* and immediately used in the next reaction (general reaction D) without further purification.

BesE substrate general reaction D: boc and/or *O*-tert-butyl deprotection.[4]

The product of reaction C was dissolved in 5 mL $CH_2Cl_2$ and 1 mL trifluoroacetic acid (TFA) was added at once. The reaction was stirred for 2 h at room temperature. The mixture was concentrated *in vacuo* re-dissolved in minimal water and lyophilized to afford the product without further purification.

#### Synthesis of γ-glutamyl-L-propargylglycine (1)

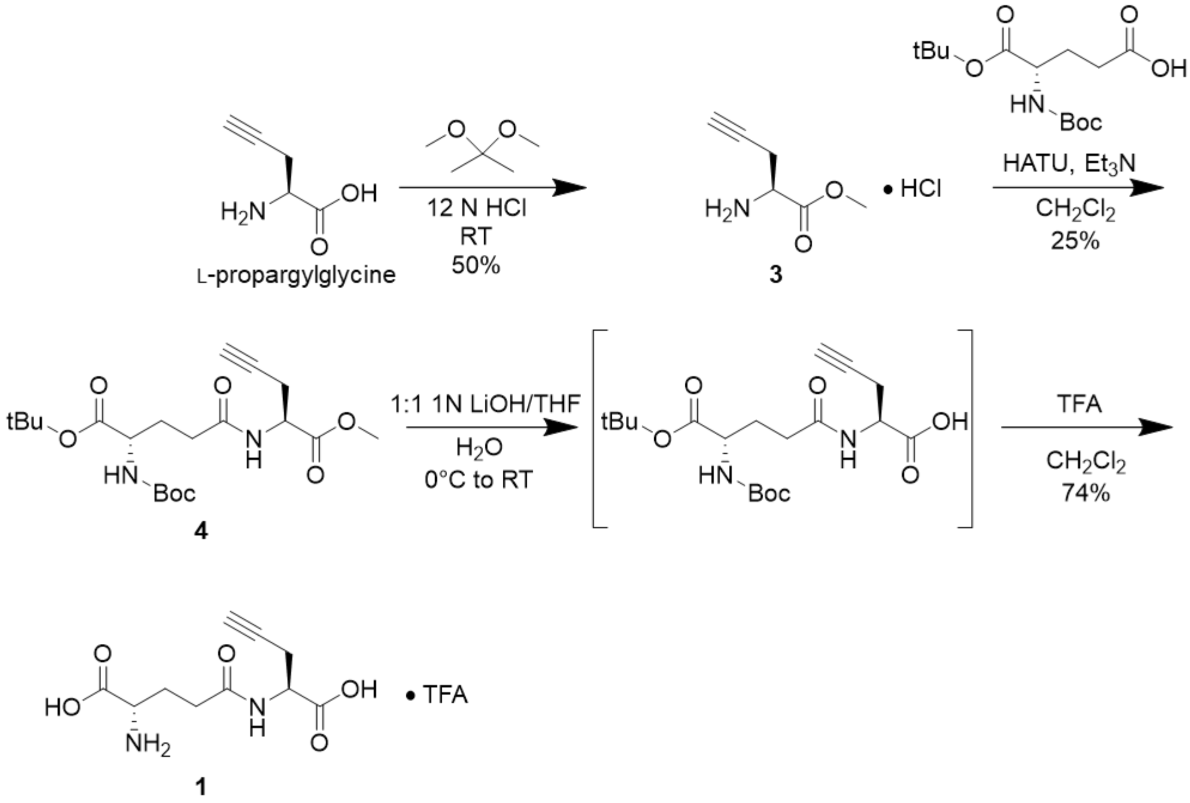


##### *O*-methyl-L-propargylglycine (3)

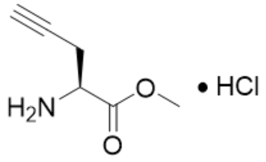

**3**

Compound was synthesized starting from from L-propargylglycine (0.120 g, 1.06 mmol) following BesE substrate general reaction A without modification to afford **3** as a sticky brown solid (0.087 g, 0.53 mmol, 50%). $^{1}$H NMR (500 MHz, $D_2O$ + 0.1% MeOH) δ 4.35 (t, *J* = 5.4 Hz, 1H), 3.84 (s, 3H), 3.02 – 2.89 (m, 2H), 2.55 (t, *J* = 2.7 Hz, 1H). $^{13}$C NMR (125 MHz, $D_2O$ + 0.1% MeOH) δ 168.9, 76.8, 75.2, 54.0, 52.6, 21.2.

**γ-(*N*-Boc-*O*-*tert*-butyl-glutamyl)-*O*-methyl-L-propargylglycine (4)**

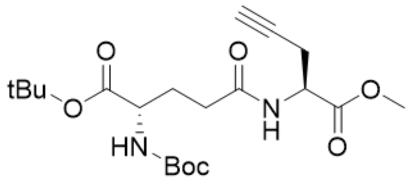


**4**

Compound was synthesized from intermediate **3** (0.05 g, 0.31 mmol) via BesE substrate general reaction B without modification to yield protected product **4** (0.031 g, 0.08 mmol, 25%). $^{1}$H NMR (500 MHz, $CDCl_3$) δ 6.70 (s, 1H), 5.20 (s, 1H), 4.74 (dt, *J* = 7.9, 5.0 Hz, 1H), 4.26 – 4.16 (m, 1H), 3.78 (s, 3H), 2.83 – 2.69 (m, 2H), 2.33 (hept, *J* = 6.9 Hz, 2H), 2.18 (tt, *J* = 13.4, 5.9 Hz, 1H), 2.03 (d, *J* = 3.2 Hz, 1H), 1.87 (dq, *J* = 15.1, 7.6 Hz, 1H), 1.45 (s, 9H), 1.44 (s, 9H). $^{13}$C NMR (125 MHz, $CDCl_3$) δ 173.44, 171.92, 171.54, 170.94, 82.38, 80.04, 78.72, 71.72, 53.53, 52.87, 50.87, 32.54, 29.30, 28.45, 28.13, 22.48.

**γ-glutamyl-L-propargylglycine (1)**

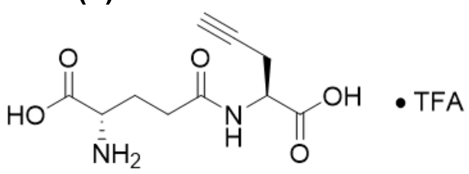


**1**

Compound was synthesized from intermediate **4** (0.035 g, 0.08 mmol) following BesE substrate general reactions C and D without modification to afford **1** as a sticky amber solid (0.021g, 0.06 mmol, 74%). $^{1}$H NMR (500 MHz, $D_2O$ + 0.1% MeOH) δ 4.55 (t, *J* = 6.0 Hz, 1H), 3.97 (t, *J* = 6.5 Hz, 1H), 2.75 (d, *J* = 5.3 Hz, 2H), 2.54 (td, *J* = 7.7, 2.7 Hz, 2H), 2.41 (s, 1H), 2.19 (dtt, *J* = 22.0, 14.6, 7.3 Hz, 2H). $^{13}$C NMR (125 MHz, $D_2O$ + 0.1% MeOH) δ 174.5, 174.0, 172.4, 79.6, 72.1, 53.0, 51.7, 31.1, 25.9, 21.0. MS (ESI) calculated for $C_{19}H_{23}N_6O_{10}^+$ (Marfey's derivatized) 495.15, found 495.19 $(M+H)^+$.

#### Synthesis of γ-glutamyl-L-allylglycine (5)

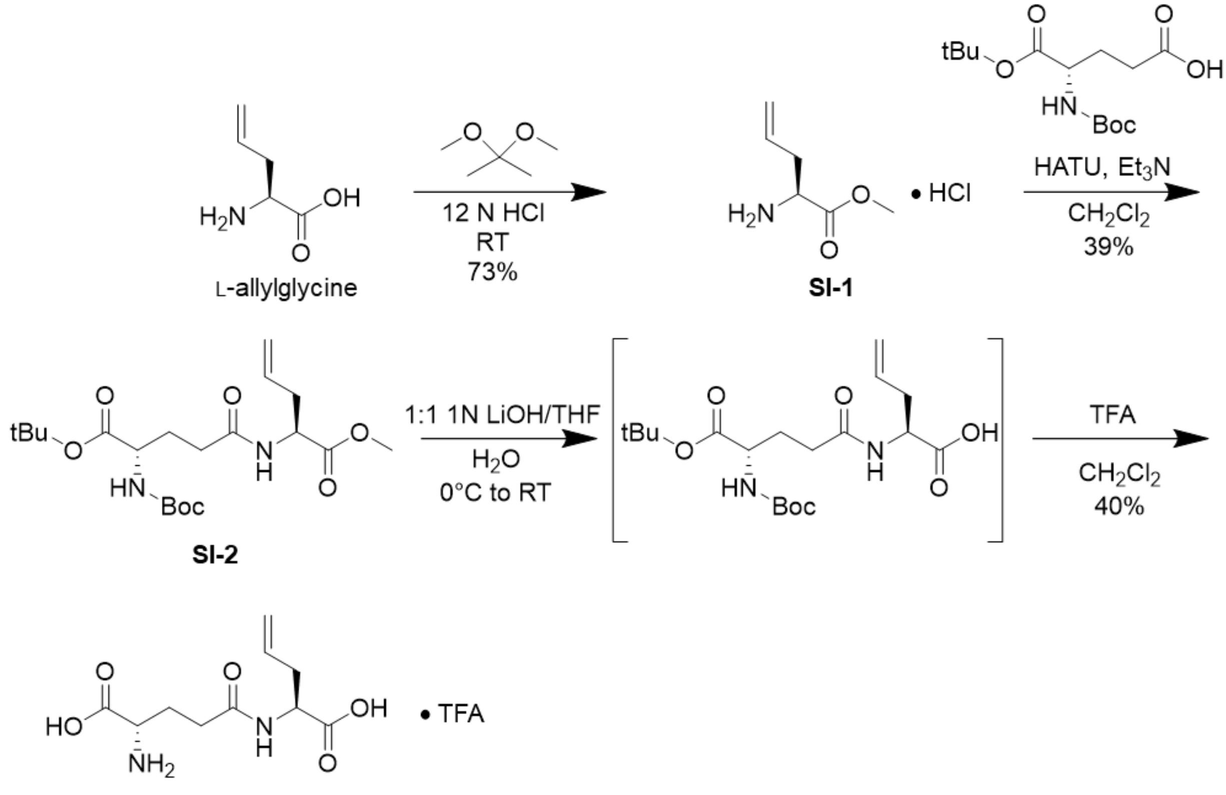


**5**

##### *O*-methyl-L-allylglycine (SI-1)

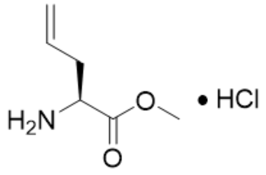


**SI-1**

Compound was synthesized via BesE substrate general reaction A starting from L-allylglycine (0.126 g) to afford **SI-1** as a brown sticky solid (0.1302g, 73%). $^{1}$H NMR (500 MHz, D2O + 0.1% MeOH) δ 5.73 (s, 1H), 5.27 (d, *J* = 11.9 Hz, 2H), 4.73 (d, *J* = 3.4 Hz, 13H), 4.23 (s, 1H), 3.82 (d, *J* = 3.2 Hz, 3H), 3.31 (s, 1H), 2.70 (d, *J* = 28.2 Hz, 2H). $^{13}$C NMR (125 MHz, D2O + 0.1% MeOH) δ 170.17, 130.07, 121.50, 53.57, 52.23, 48.88, 34.29, 34.00.

**γ-(*N*-Boc-*O*-*tert*-butyl-glutamyl)-*O*-methyl-L-allylglycine (SI-2)**

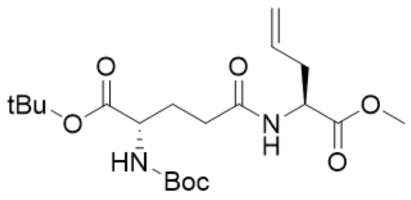

**SI-2**

Compound was synthesized via BesE substrate general reaction B without modification starting from methyl ester intermediate **SI-1** (0.1310 g) to afford **SI-2** (0.129 g, 39%).[1]H NMR (500 MHz, $CDCl_3$) δ 6.49 (s, 1H), 5.78 – 5.66 (m, 1H), 5.12 (s, 4H), 4.68 (s, 1H), 4.13 (s, 3H), 3.75 (s, 4H), 2.57 (d, *J* = 34.4 Hz, 2H), 2.30 (s, 3H), 2.17 (s, 4H), 2.04 (s, 2H), 1.85 (s, 2H), 1.56 (s, 8H), 1.45 (s, 24H), 1.26 (s, 3H). $^{13}C$ NMR (125 MHz, $CDCl_3$) δ 77.27, 77.01, 76.76, 52.34, 36.47, 28.32, 28.01.

**γ-glutamyl-L-allylglycine (5)**

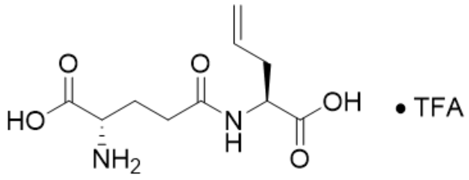

**5**

Compound was synthesized following BesE substrate general procedures C and D starting from protected intermediate **SI-2** (0.116 g) to afford **5** (0.029 g, 40%). $^1H$ NMR (500 MHz, D2O + 0.1% MeOH) δ 5.78 (s, 1H), 5.18 (s, 1H), 5.14 (s, 2H), 4.73 (s, 362H), 4.42 (s, 1H), 4.00 (s, 1H), 3.31 (s, 17H), 2.60 (s, 1H), 2.51 (s, 3H), 2.18 (s, 2H), 1.47 (s, 2H). $^{13}C$ NMR (125 MHz, D2O + 0.1% MeOH) δ 174.4, 132.8, 118.9, 52.7, 52.6, 49.0, 34.9, 31.0, 27.2, 25.78. MS (ESI) calculated for $C_{19}H_{25}N_6O_{10}^+$ (Marfey's derivatized) 497.16, found 497.19 $(M+H)^+$.

**Synthesis of γ-glutamyl-L-norvaline (6)**

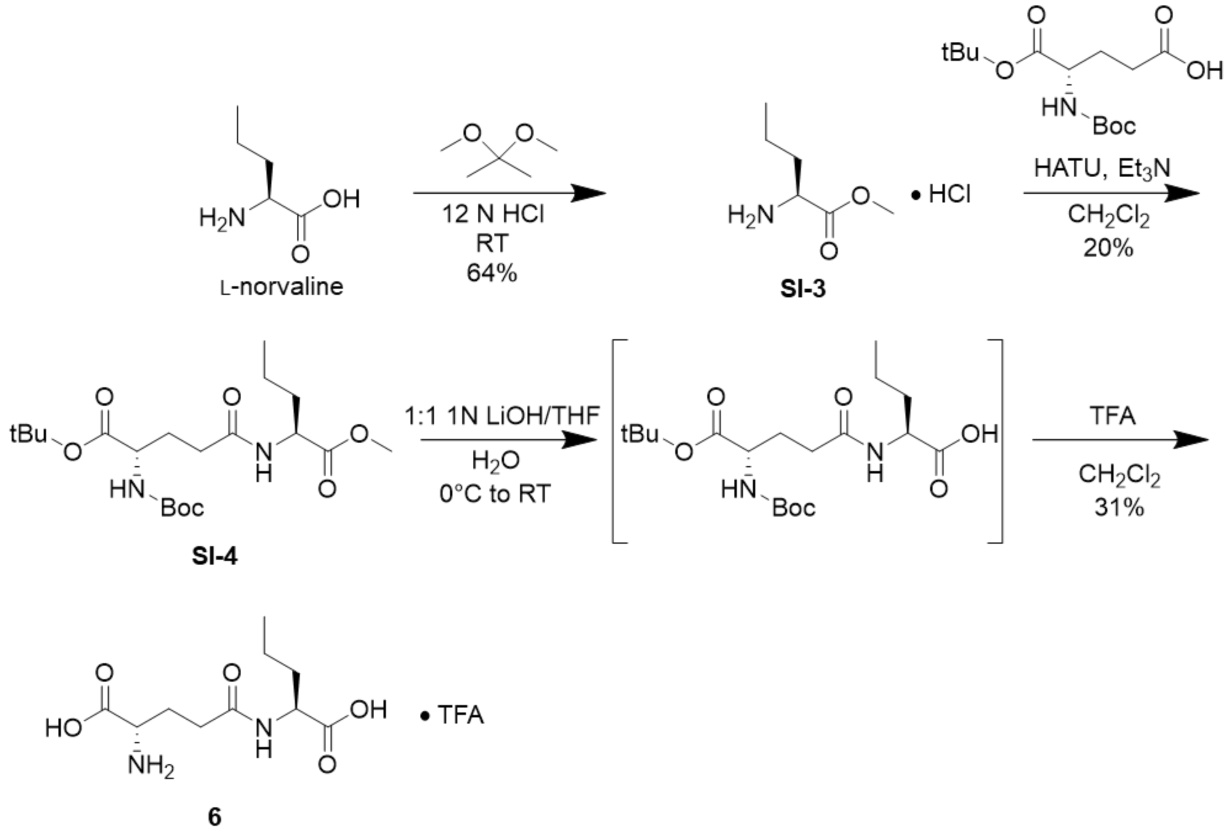


***O*-methyl-L-norvaline (SI-3)**

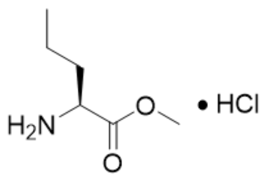


**SI-3**

Compound was synthesized via BesE substrate general reaction A starting from L-norvaline (0.109 g) to afford **SI-3** as a brown sticky solid (0.069 g, 64%). $^1$H NMR (500 MHz, $D_2O$ + 0.1 MeOH) δ 4.11 (t, *J* = 6.6 Hz, 1H), 3.81 (s, 3H), 1.98 – 1.75 (m, 3H), 1.39 (th, *J* = 13.0, 6.7 Hz, 3H), 0.91 (t, *J* = 7.4 Hz, 4H). $^{13}$C NMR (125 MHz, $D_2O$ + 0.1 MeOH) δ 170.9, 53.6, 52.8, 31.8, 17.8, 12.8.

**γ-(*N*-Boc-*O*-*tert*-butyl-glutamyl)-*O*-methyl-L-norvaline (SI-4)**

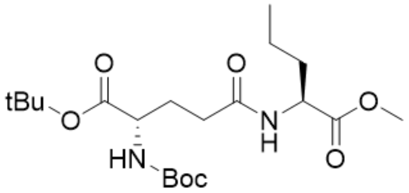


**SI-4**

Compound was synthesized via BesE substrate general reaction B without modification starting from methyl ester intermediate **SI-3** (0.069 g) to afford **SI-4** (0.100 g, 20%). $^{1}H$ NMR (500 MHz, $CDCl_3$) δ 6.48 (s, 1H), 5.20 (s, 1H), 4.60 (t, *J* = 10.4 Hz, 1H), 4.21 – 4.14 (m, 1H), 3.74 (s, 3H), 2.36 – 2.24 (m, 2H), 2.23 – 2.12 (m, 1H), 1.93 – 1.77 (m, 2H), 1.71 – 1.63 (m, 1H), 1.48 – 1.43 (m, 18H), 1.39 – 1.32 (m, 2H), 0.93 (t, *J* = 8.9 Hz, 3H).

**γ-glutamyl-L-norvaline (6)**

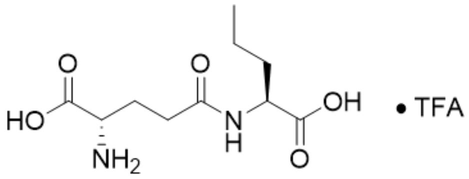


**6**

Compound was synthesized following BesE substrate general procedures C and D starting from protected intermediate **SI-4** (0.051 g) to afford **6** (0.036 g, 31%). $^{1}H$ NMR (500 MHz, $D_2O$ + 0.1% MeOH) δ 4.29 (dd, *J* = 9.0, 5.1 Hz, 1H), 3.97 (dt, *J* = 23.2, 6.5 Hz, 1H), 2.51 (qd, *J* = 7.4, 4.2 Hz, 2H), 2.23 – 2.11 (m, 2H), 1.83 – 1.65 (m, 2H), 1.44 – 1.29 (m, 3H), 0.88 (t, *J* = 7.4 Hz, 3H). $^{13}C$ NMR (125 MHz, $D_2O$ + 0.1% MeOH) δ 176.48, 174.28, 168.61, 53.00, 52.82, 32.58, 31.08, 25.94, 18.56, 12.84. MS (ESI) calculated for $C_{19}H_{27}N_6O_{10}^{+}$ (Marfey's derivatized) 499.18, found 499.22 $(M+H)^{+}$.

#### Synthesis of γ-glutamyl-L-homopropargylglycine (7)

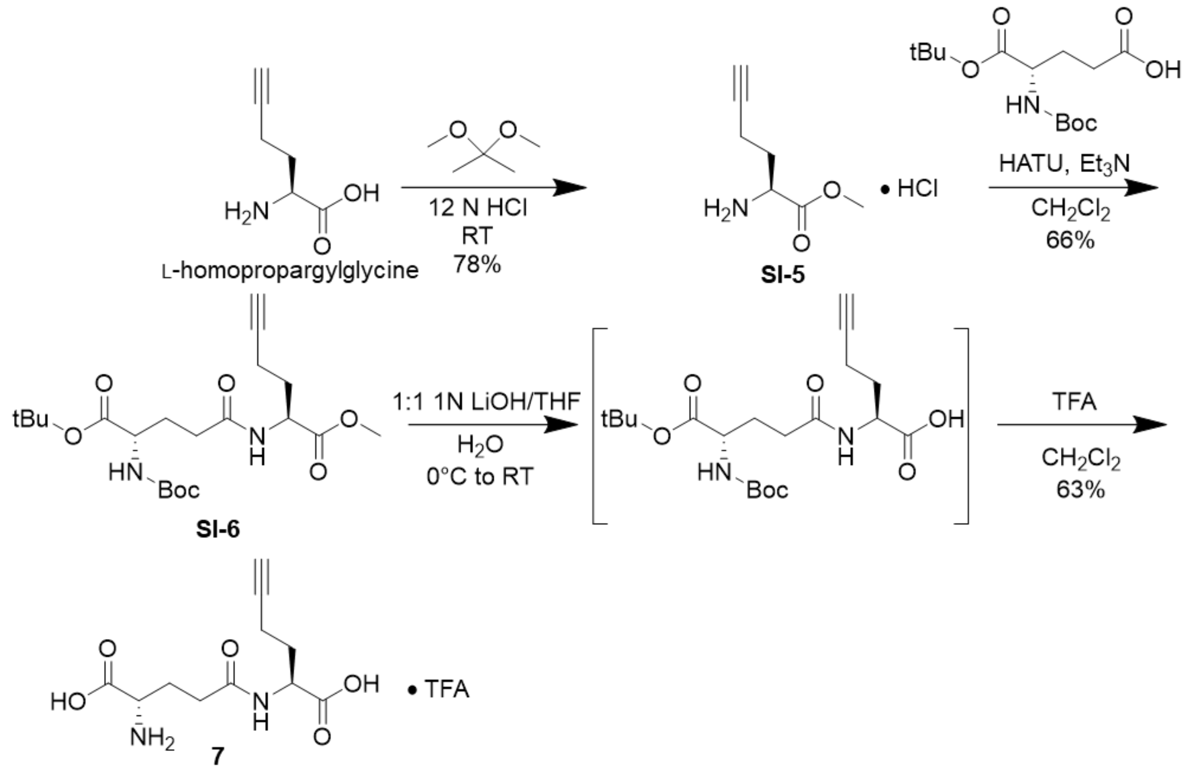


##### *O*-methyl-L-homopropargylglycine (SI-5)

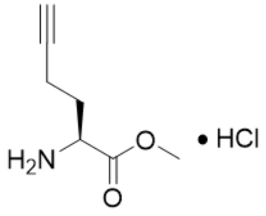


**SI-5**

Compound was synthesized via BesE substrate general reaction A starting from L-homopropargylglycine (0.05 g, 0.31 mmol) to afford **SI-5** (0.043 g, 0.24 mmol, 78%). $^{1}H$ NMR (500 MHz, $D_2O$ + 0.1 MeOH) δ 4.27 (t, *J* = 6.4 Hz, 1H), 3.82 (s, 3H), 2.45 – 2.40 (m, 3H), 2.15 (ddt, *J* = 41.6, 14.7, 7.7 Hz, 3H). $^{13}C$ NMR (125 MHz, $D_2O$ + 0.1 MeOH) δ 170.50, 82.38, 71.32, 53.72, 51.96, 28.52, 14.15.

**γ-(N-Boc-*O*-*tert*-butyl-glutamyl)-*O*-methyl-L-homopropargylglycine (SI-6)**

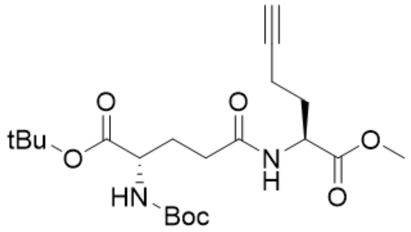


**SI-6**

Compound was synthesized via BesE substrate general reaction B without modification starting from methyl ester intermediate **SI-5** (0.043 g, 0.24 mmol) to afford **SI-6** (0.068 g, 0.16 mmol, 66%). $^1$H NMR (500 MHz, $CDCl_3$) δ 6.65 (s, 1H), 5.31 – 5.15 (m, 1H), 4.72 – 4.64 (m, 1H), 4.18 – 4.07 (m, 1H), 3.76 (s, 3H), 2.38 – 2.23 (m, 4H), 2.23 – 2.08 (m, 2H), 1.99 (s, 1H), 1.98 – 1.92 (m, 1H), 1.90 – 1.79 (m, 1H), 1.47 (s, 9H), 1.45 (s, 9H).

**γ-glutamyl-L-homopropargylglycine (7)**

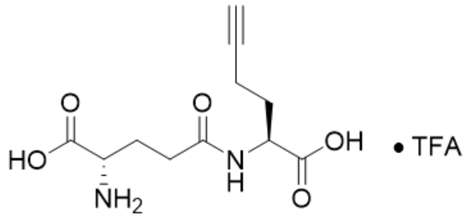


**7**

Compound was synthesized from protected intermediate **SI-6** (0.068 g, 0.16 mmol) following BesE substrate general reactions C and D with slight modification to reaction D only: The TFA deprotection was allowed to run overnight. Subsequent concentration *in vacuo* yielded **7** (0.026 g, 0.10 mmol, 63%). $^1$H NMR (500 MHz, $D_2O$ + 0.1% MeOH) δ 4.51 (dq, *J* = 10.6, 6.2, 5.1 Hz, 1H), 4.07 (dq, *J* = 9.8, 6.1, 4.0 Hz, 1H), 2.54 (qd, *J* = 7.1, 4.7, 3.6 Hz, 2H), 2.36 (s, 1H), 2.31 (dt, *J* = 9.2, 4.3 Hz, 2H), 2.21 (dd, *J* = 14.1, 6.9 Hz, 2H), 2.14 – 2.06 (m, 1H), 1.92 (s, 1H). $^{13}$C NMR (125 MHz, $D_2O$ + 0.1% MeOH) δ 175.45, 174.56, 171.67, 83.68, 70.45, 52.39, 52.01, 31.01, 29.17, 25.62, 14.69. MS (ESI) calculated for $C_{20}H_{25}N_6O_{10}^+$ (Marfey's derivatized) 509.16, found 509.21 $(M+H)^+$.

#### Synthesis of γ-glutamyl-L-alanine (8)

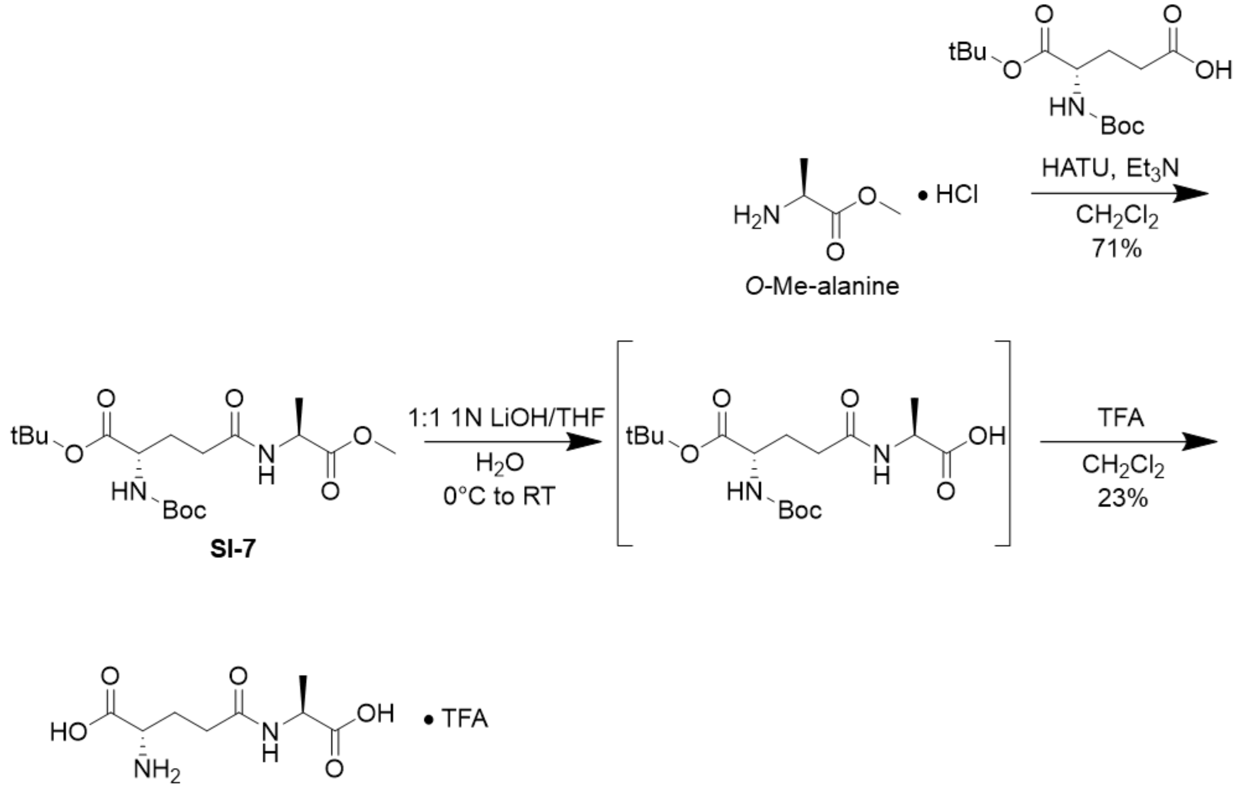


**8**

##### γ-(*N*-Boc-*O*-*tert*-butyl-glutamyl)-*O*-methyl-L-alanine (SI-7)

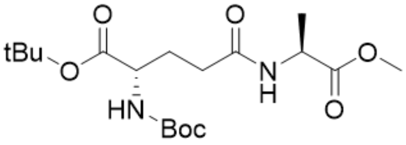


**SI-7**

Compound was synthesized via BesE substrate general reaction B without modification starting from *O*-Me-alanine (0.074 g) to afford **SI-7** (0.147g, 71%). 1H NMR (500 MHz, $CDCl_3$) δ 6.50 (s, 1H), 5.20 (d, J = 7.9 Hz, 1H), 4.58 (p, J = 7.2 Hz, 1H), 4.16 (s, 1H), 3.75 (s, 3H), 2.30 (t, J = 7.7 Hz, 2H), 2.17 (q, J = 6.9, 5.4 Hz, 1H), 1.86 (q, J = 7.4 Hz, 1H), 1.62 (s, 2H), 1.46 (s, 9H), 1.44 (s, 9H), 1.42 (d, J = 7.2 Hz, 3H). $^{13}C$ NMR (125 MHz, $CDCl_3$) δ 173.64, 171.70, 171.56, 155.95, 82.44, 80.09, 53.54, 52.57, 48.27, 32.62, 29.41, 28.46, 28.14, 18.46.

**γ-glutamyl-L-alanine (8)**

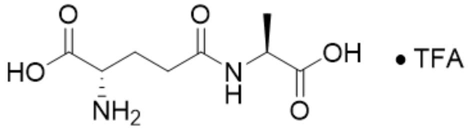

**8**

Compound was synthesized following BesE substrate general procedures C and D starting from protected intermediate **SI-7** (0.036 g, 0.09 mmol) to afford **8** (0.125g, 23%). $^{1}$H NMR (500 MHz, $D_2O$ + 0.1% MeOH) δ 4.28 (q, *J* = 7.3 Hz, 1H), 3.83 (t, *J* = 6.4 Hz, 1H), 2.46 (t, *J* = 7.3 Hz, 2H), 2.14 (dh, *J* = 15.0, 7.2, 6.8 Hz, 2H), 1.37 (d, *J* = 7.4 Hz, 3H). $^{13}$C NMR (125 MHz, $D_2O$ + 0.1% MeOH) δ 177.27, 174.42, 173.37, 53.74, 49.14, 31.23, 26.11, 16.25. MS (ESI) calculated for $C_{17}H_{23}N_6O_{10}^+$ (Marfey's derivatized) 471.15, found 471.20 $(M+H)^+$.

#### Synthesis of γ-glutamyl-D-propargylglycine (9)

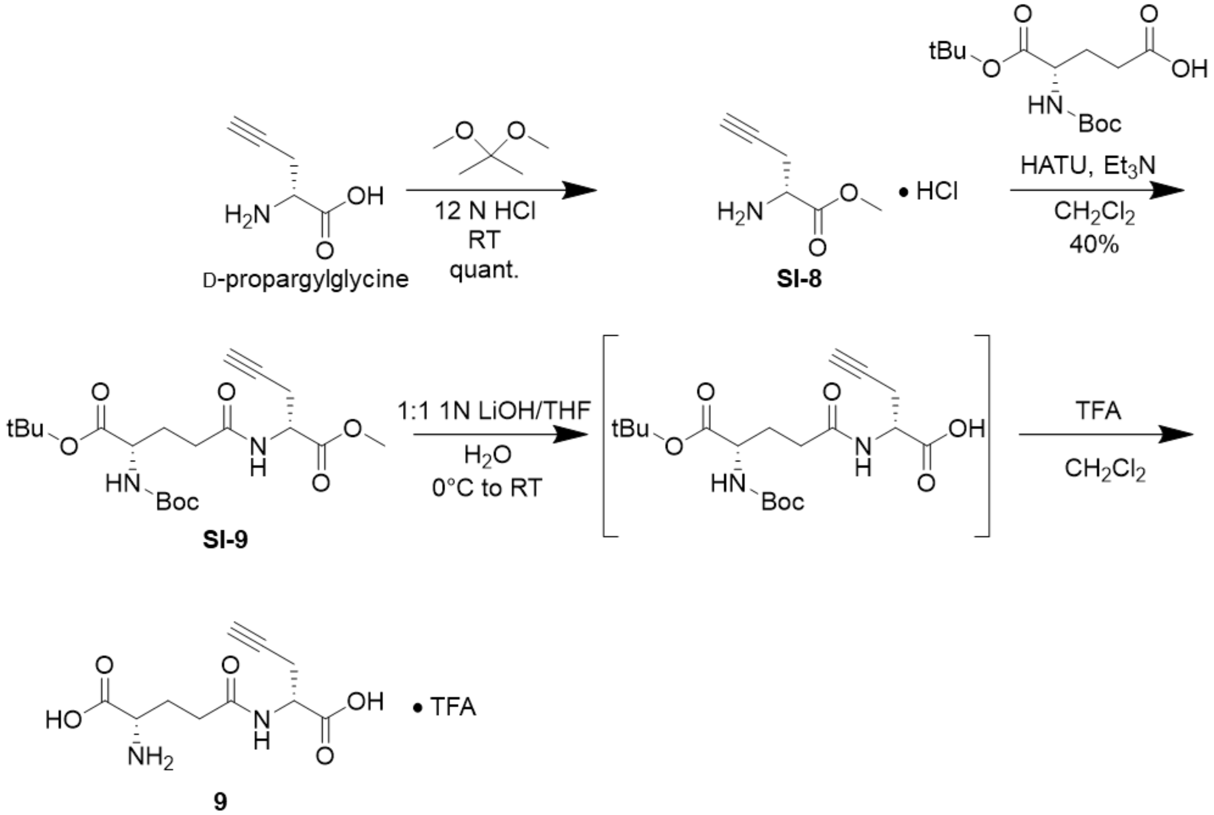


#### *O*-methyl-γ-glutamyl-D-propargylglycine (SI-8)

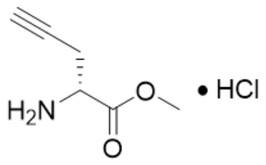


**SI-8**

Compound was synthesized via BesE substrate general reaction A starting from D-propargylglycine (0.200 g, 1.8 mmol) to afford **SI-8** as a yellow/orange sticky solid (0.295 g, 1.8 mmol, quant.) $^1$H NMR (500 MHz, $D_2O$ + 0.1 MeOH) δ 4.37 (t, *J* = 5.6 Hz, 1H), 3.84 (s, 3H), 3.02 – 2.89 (m, 2H), 2.56 (s, 1H). $^{13}$C NMR (125 MHz, $D_2O$ + 0.1 MeOH) δ 169.14, 76.40, 74.45, 54.01, 51.46, 19.99.

**γ-(*N*-Boc-*O*-*tert*-butyl-glutamyl)-*O*-methyl-D-propargylglycine SI-9)**

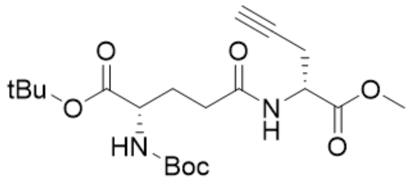


**SI-9**

Compound was synthesized via BesE substrate general reaction B without modification starting from methyl ester intermediate **SI-8** (0.285 g, 1.74 mmol) to afford **SI-9** (0.284 g, 0.69 mmol, 40%). $^1$H NMR (500 MHz, $CDCl_3$) δ 6.80 (s, 1H), 5.22 (s, 1H), 4.74 (p, *J* = 5.3, 4.3 Hz, 1H), 4.23 (s, 1H), 3.78 (s, 3H), 2.83 – 2.71 (m, 2H), 2.41 – 2.27 (m, 2H), 2.25 – 2.09 (m, 1H), 2.03 (s, 1H), 1.95 – 1.83 (m, 1H), 1.46 (s, 9H), 1.44 (s, 9H).$^{13}$C NMR (125 MHz, $CDCl_3$) δ 172.10, 171.57, 170.98, 82.39, 80.06, 78.64, 71.72, 60.53, 53.58, 52.89, 50.88, 32.62, 28.46, 28.14, 22.46.

**γ-glutamyl-D-propargylglycine (9)**

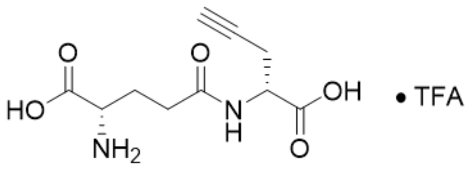


**9**

Compound was synthesized from protected intermediate **SI-9** (0.274g, 0.66 mmol) following BesE substrate general reactions C and D with slight modification to reaction D only: The TFA deprotection was allowed to run overnight. Subsequent concentration *in vacuo* yielded **9** (0.006 g, 0.025 mol, 4%). $^1$H NMR (500 MHz, $D_2O$ + 0.1% MeOH) δ 4.58 (q, *J* = 5.5 Hz, 1H), 4.09 (dt, *J* = 7.9, 5.0 Hz, 1H), 2.76 (p, *J* = 3.8, 3.4 Hz, 2H), 2.57 (tt, *J* = 7.4, 3.3 Hz, 2H), 2.47 – 2.39 (m, 1H), 2.32 – 2.14 (m, 2H). $^{13}$C NMR (125 MHz, $D_2O$ + 0.1% MeOH) δ 174.30, 173.75, 171.54, 79.55, 72.16, 52.22, 51.52, 30.81, 25.58, 20.93. MS (ESI) calculated for $C_{19}H_{23}N_6O_{10}^+$ (Marfey's derivatized) 495.15, found 495.21 $(M+H)^+$.

#### Synthesis of γ-glutamyl-L-propargylglycine methyl ester (10)

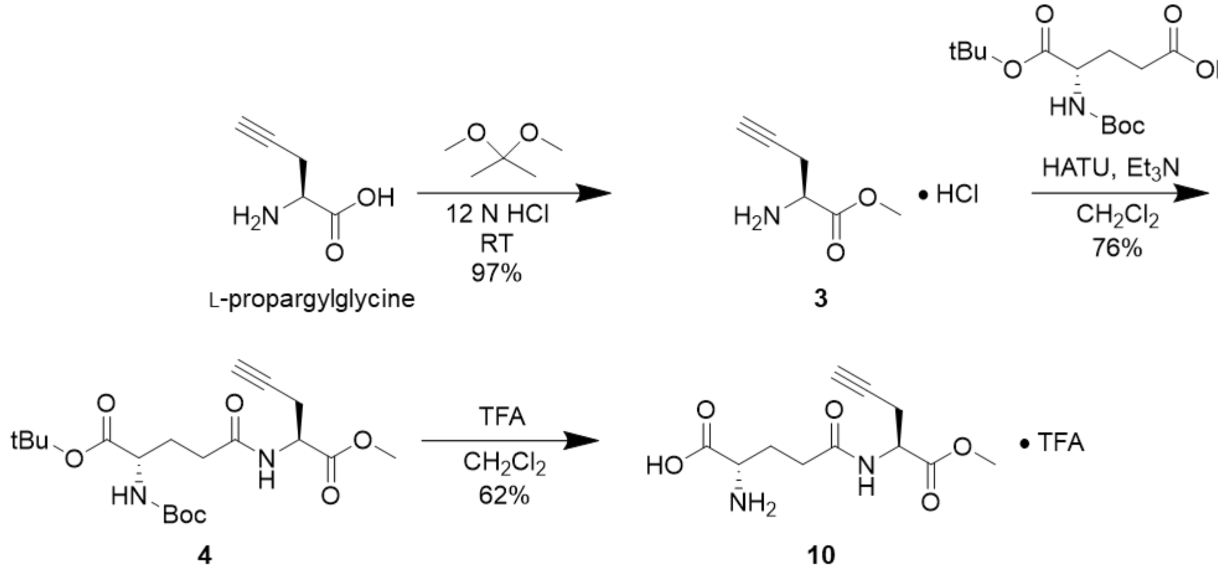


##### γ-glutamyl-L-propargylglycine methyl ester (10)

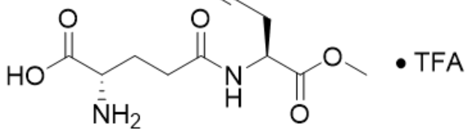


**10**

Compound was synthesized from protected intermediate **4** (0.050 g, 0.12 mmol) via BesE substrate general reaction D with slight modification: the TFA deprotection was allowed to run overnight. Concentration *in vacuo* followed by resuspension in water and lyophilization yielded **10** (0.028 g, 0.08 mmol, 62%). $^1$H NMR (500 MHz, $D_2O$ + 0.1% MeOH) δ 4.61 (t, $J$ = 6.2 Hz, 1H), 4.02 (t, $J$ = 6.6 Hz, 1H), 3.75 (s, 3H), 2.82 – 2.72 (m, 2H), 2.55 (td, $J$ = 7.7, 3.2 Hz, 2H), 2.42 (d, $J$ = 3.2 Hz, 1H), 2.20 (th, $J$ = 14.7, 7.2 Hz, 2H). $^{13}$C NMR (125 MHz, $D_2O$ + 0.1% MeOH) δ 174.47, 172.58, 171.95, 79.42, 72.23, 53.32, 52.58, 51.64, 30.95, 25.74, 20.86. MS (ESI) calculated for $C_{20}H_{25}N_6O_{10}^+$ (Marfey's derivatized) 509.16, found 509.22 $(M+H)^+$.

#### Synthesis of γ-glutamyl-(*R*)-pent-4-yn-2-amine (11)

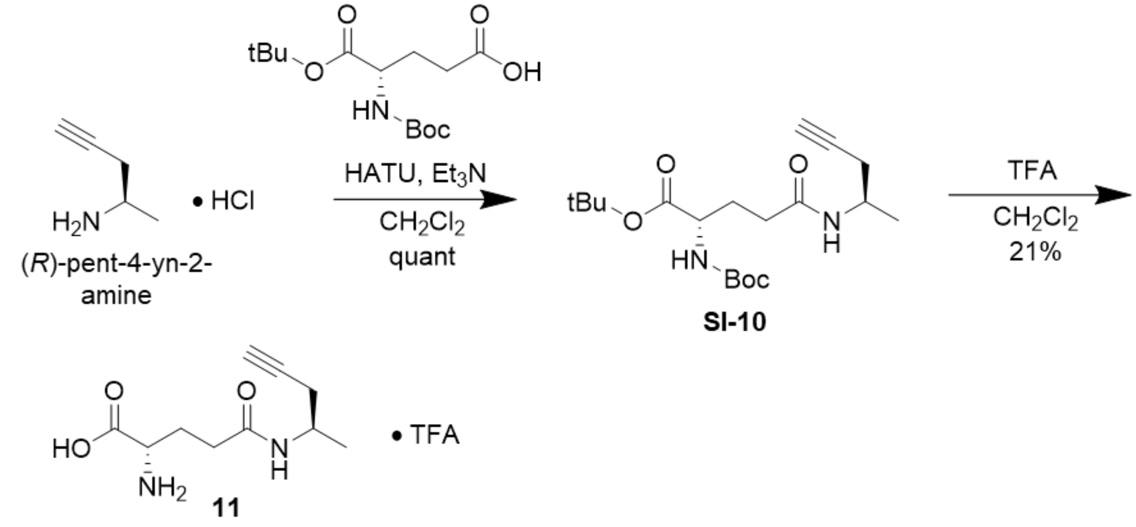


##### γ-(*N*-Boc-*O*-*tert*-butyl-glutamyl)-(*R*)-pent-4-yn-2-amine (SI-10)

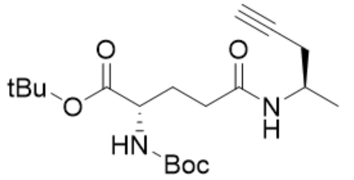


**SI-10**

Compound was synthesized via BesE substrate general reaction B without modification starting from (*R*)-pent-4-yn-2-amine (0.035 g, 0.29 mmol) to afford **SI-10** (0.120 g, 0.33 mmol, quant.). $^{1}$H NMR (500 MHz, $CDCl_3$) δ 6.33 – 6.18 (m, 1H), 5.22 (d, J = 8.2 Hz, 1H), 4.27 – 4.10 (m, 2H), 2.48 (ddd, J = 16.7, 5.8, 2.6 Hz, 1H), 2.36 (ddd, J = 16.7, 4.6, 2.7 Hz, 1H), 2.30 – 2.18 (m, 2H), 2.18 – 2.09 (m, 1H), 2.00 (t, J = 2.6 Hz, 1H), 1.86 (d, J = 9.9 Hz, 1H), 1.44 (d, J = 8.6 Hz, 18H), 1.24 (d, J = 6.7 Hz, 3H). $^{13}$C NMR (125 MHz, $CDCl_3$) δ 171.61, 171.52, 156.04, 82.37, 80.64, 80.07, 77.36, 70.89, 53.61, 43.59, 33.01, 29.63, 28.45, 28.13, 25.79, 19.58.

**γ-glutamyl-(*R*)-pent-4-yn-2-amine (11)**

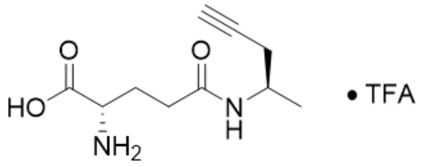

**11**

Compound was synthesized from protected intermediate **SI-10** (0.05 g, 0.14 mmol) via BesE substrate general reaction D with slight modification: the TFA deprotection was allowed to run overnight. Concentration *in vacuo* followed by resuspension in water and lyophilization yielded **11** (0.009 g, 0.03 mmol, 21%). $^{1}$H NMR (500 MHz, $D_2O$ + 0.1% MeOH) δ 4.00 (dt, *J* = 8.6, 6.2 Hz, 2H), 2.51 – 2.37 (m, 3H), 2.35 (s, 1H), 2.34 – 2.29 (m, 1H), 2.25 – 2.10 (m, 2H), 1.18 (d, *J* = 6.6 Hz, 3H). $^{13}$C NMR (125 MHz, $D_2O$ + 0.1% MeOH) δ 173.60, 171.98, 81.82, 71.14, 52.66, 44.46, 31.41, 26.03, 25.03, 18.61. MS (ESI) calculated for $C_{19}H_{25}N_6O_8^+$ (Marfey's derivatized) 465.17, found 465.23 $(M+H)^+$.

#### Synthesis of γ-glutamyl-but-3-yn-1-amine (12)

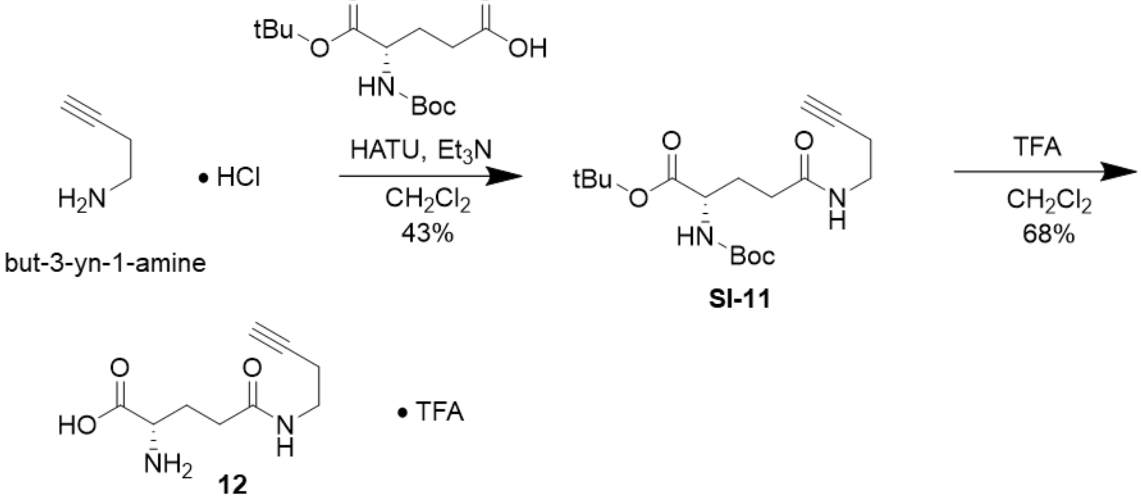


##### γ-(*N*-Boc-*O*-*tert*-butyl-glutamyl)-but-3-yn-1-amine (SI-11)

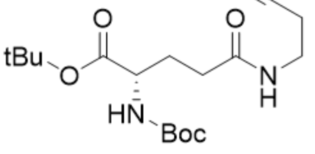


**SI-11**

Compound was synthesized via BesE substrate general reaction B without modification starting from but-3-yn-1-amine (0.098 g, 0.93 mmol) to afford protected intermediate **SI-11** (0.143 g, 0.40 mmol, 43%). $^{1}$H NMR (500 MHz, $CDCl_3$) δ 6.44 (s, 1H), 5.22 (d, *J* = 7.6 Hz, 1H), 4.18 (s, 1H), 3.42 (d, *J* = 5.1 Hz, 2H), 2.42 (s, 2H), 2.27 (s, 2H), 2.15 (s, 1H), 1.99 (s, 1H), 1.87 (s, 1H), 1.46 (s, 9H), 1.45 (s, 9H).$^{13}$C NMR (125 MHz, $CDCl_3$) δ 172.34, 171.57, 156.10, 82.47, 81.76, 80.13, 70.06, 53.60, 38.27, 32.91, 29.75, 28.46, 28.14, 19.56.

**γ-glutamyl-but-3-yn-1-amine (12)**

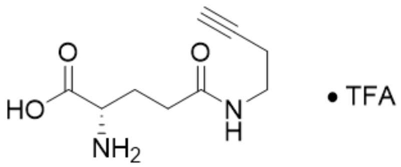


**12**

Compound was synthesized from protected intermediate **SI-11** (0.143 g, 0.40 mmol) via BesE substrate general reaction D with slight modification: the TFA deprotection was allowed to run for 24 h. Concentration *in vacuo* followed by resuspension in water and lyophilization yielded **12** (0.054 g, 0.27 mmol, 68%). $^{1}H$ NMR (500 MHz, $D_2O$ + 0.1% MeOH) δ 4.06 – 4.01 (m, 1H), 3.35 – 3.31 (m, 2H), 2.53 – 2.42 (m, 2H), 2.42 – 2.36 (m, 2H), 2.35 – 2.30 (m, 1H), 2.27 – 2.11 (m, 2H). $^{13}C$ NMR (125 MHz, $D_2O$) δ 174.42, 171.81, 82.64, 70.48, 52.49, 38.12, 31.32, 25.93, 18.47. MS (ESI) calculated for $C_{18}H_{23}N_6O_8^+$ (Marfey's derivatized) 451.16, found 451.22 $(M+H)^+$.

#### Synthesis of γ-glutamyl-L-propargylglycine-α-amide (13)

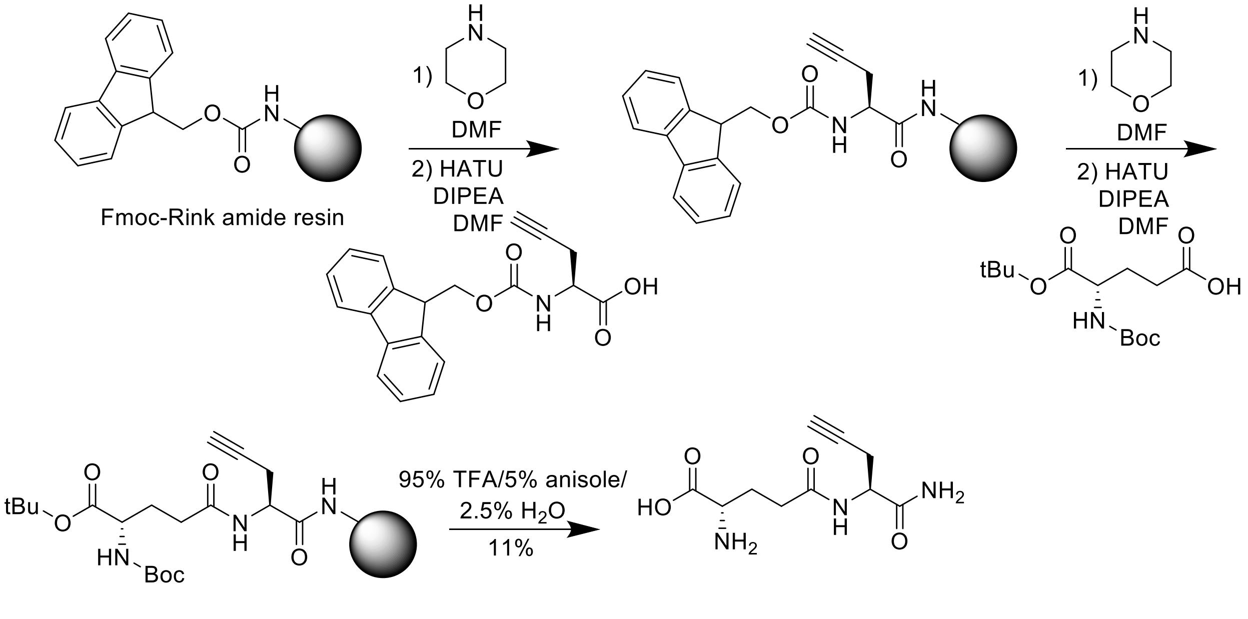


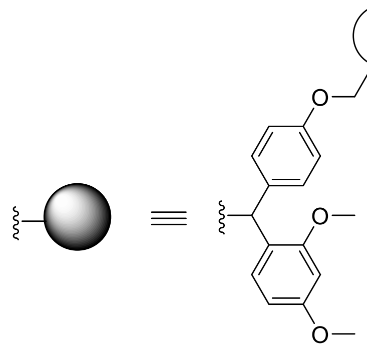

**γ-glutamyl-L-propargylglycine-α-amide (13)**

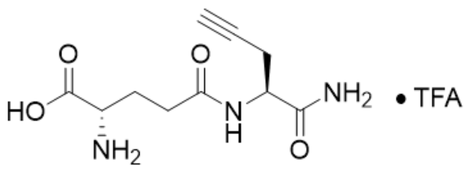


**13**

Compound was generated via solid phase peptide synthesis using Rink amide resin. 0.256 g of the resin was used in the reaction and it was then swelled with DMF washes (3 x 5 mL) while bubbling with argon gas. The Fmoc-group on the resin was then deprotected with a 20% morpholine in DMF solution. The first deprotection was washed with the 20% morpholine in DMF solution 8 times at 5 mL for 5-15 min each to achieve deprotection. After the deprotection, the resin was then washed with DMF (3 x 5 mL). The resin was then coupled with Fmoc-Pra-OH (5 eq.) with HATU (5 eq.), and DPEA (10 eq.) for 1 h. Another deprotection was performed with 20% morpholine in DMF solution (3 x 5 mL) for 15 min each. After deprotection, wash resin with DMF (3 x 5 mL). Boc-Glu-OtBu (5 eq.) was then coupled with HATU (5 eq.), and DiPEA (10 eq.) for 1 h. After the coupling, the resin was washed with DMF (3 x 5 mL). The resin was then prepared for overnight storage. It was dried and stored with argon gas and then put in a -20 °C freezer. The cleavage and final deprotection was done with a 10 mL solution of 95% TFA/2.5 % anisole/ 2.5% water. The resin and solution mixture was then rocked for 1.25 h. The resin was pipet filtered with glass wool into a round bottomed flask and concentrated *in vacuo* until 10% of the initial volume. Cold diethyl ether was then added to the mixture to triturate out peptide product. The resulting precipitate was then collected, resuspended into water and lyophilized to yield the product **13** (0.027 g, 0.11 mmol, 11%). $^{1}H$ NMR (500 MHz, $D_2O$ + 0.1% MeOH) δ 4.46 (q, $J$ = 5.6 Hz, 1H), 4.02 – 3.91 (m, 1H), 2.69 (dd, $J$ = 6.2, 3.4 Hz, 2H), 2.55 (p, $J$ = 8.5 Hz, 2H), 2.43 (s, 1H), 2.18 (d, $J$ = 8.9 Hz, 2H). $^{13}C$ NMR (125 MHz, $D_2O$+ 0.1% MeOH) δ 175.00, 174.67, 174.65, 79.40, 72.31, 52.93, 52.30, 31.00, 25.72, 21.17. MS (ESI) calculated for $C_{19}H_{24}N_7O_9^+$ (Marfey's derivatized) 494.16, found 494.21 $(M+H)^+$.

#### Synthesis of γ-aminobutyryl-L-propargylglycine (14)

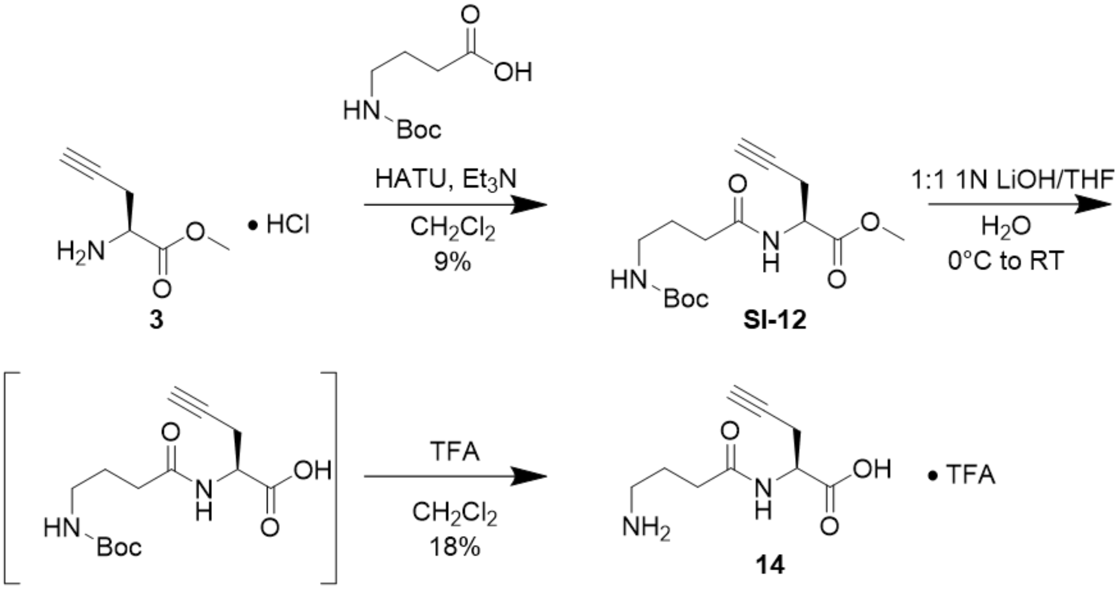


**γ-(*N*-Boc-glutamyl)-aminobutyryl-*O*-methyl-L-propargylglycine (SI-12)**

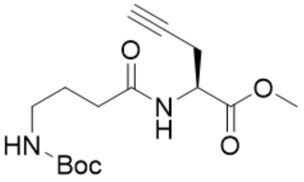


**SI-12**

Compound was synthesized via BesE substrate general reaction B without modification starting from **3** (0.101g, 0.62 mmol) to afford protected intermediate **SI-12** (0.017 g, 0.053 mmol, 9%). $^{1}H$ NMR (500 MHz, $CDCl_3$) δ 6.63 (s, 1H), 4.76 – 4.70 (m, 1H), 3.79 (s, 3H), 3.19 (s, 2H), 2.85 – 2.73 (m, 2H), 2.38 – 2.27 (m, 2H), 2.03 (s, 1H), 1.86 – 1.78 (m, 2H), 1.44 (s, 9H). $^{13}C$ NMR (125 MHz, $CDCl_3$) δ 172.51, 171.04, 156.42, 78.64, 71.77, 52.93, 50.80, 39.88, 33.60, 28.55, 26.20, 22.49.

**γ-aminobutyryl-L-propargylglycine (14)**

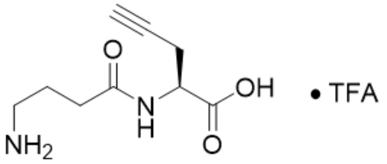


**14**

Compound was synthesized from protected intermediate **SI-12** (0.035 g, 0.08 mmol) following BesE substrate general reactions C and D with slight modification to reaction D only: The TFA deprotection was allowed to run overnight, however stirring was stopped after 4 h due to a power outage. Subsequent concentration *in vacuo* yielded **14** (0.003 g, 0.015 mmol, 18%). $^{1}$H NMR (500 MHz, $D_2O$ + 0.1% MeOH) δ 4.52 (q, *J* = 5.2 Hz, 1H), 3.01 (d, *J* = 8.7 Hz, 2H), 2.74 (s, 2H), 2.42 (dt, *J* = 12.5, 5.4 Hz, 3H), 1.95 (q, *J* = 8.9, 8.2 Hz, 2H). $^{13}$C NMR (126 MHz, $D_2O$) δ 174.86, 173.76, 79.70, 71.87, 51.89, 38.77, 32.10, 22.83, 20.95. MS (ESI) calculated for $C_{18}H_{23}N_6O_8^+$ (Marfey's derivatized) 451.16, found 451.21 $(M+H)^+$.

**Synthesis of γ-glutaryl-L-propargylglycine (15)**

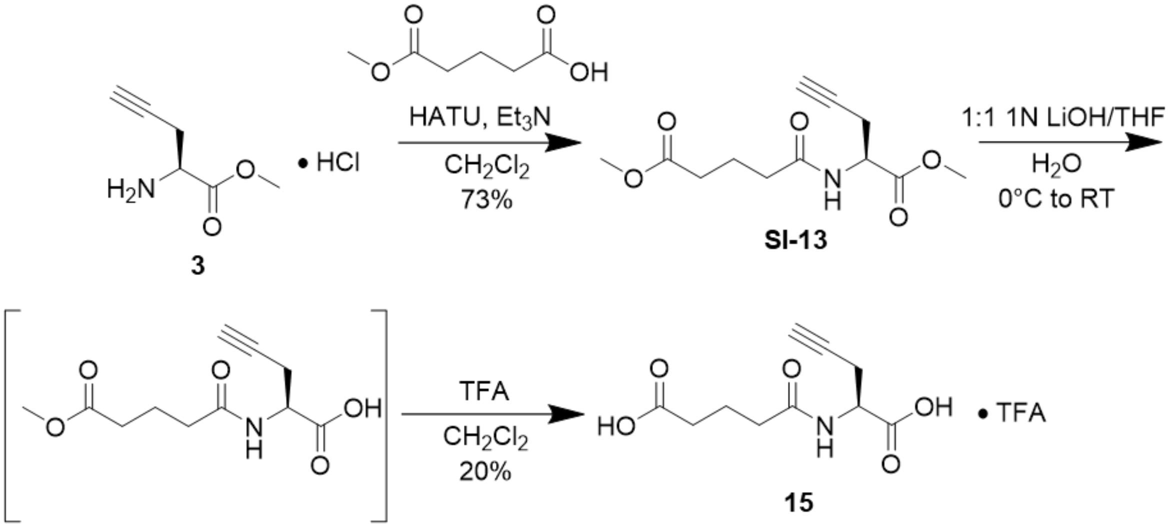

**γ-(*O*-methyl-glutaryl)-*O*-methyl-L-propargylglycine (SI-13)**

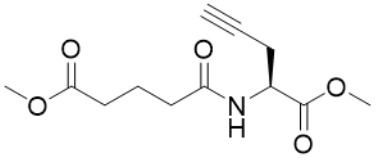

**SI-13**

Compound was synthesized starting from methyl ester **3** following modified BesE substrate general reaction B with the following modification: methyl hydrogen glutarate was used in place of Boc-L-glutamic acid-α-tert-butyl-ester. **SI-13** was isolated as a white solid (0.075 g, 0.29 mmol, 73%). $^{1}$H NMR (500 MHz, $CDCl_3$) δ 6.33 (d, J = 7.7 Hz, 1H), 4.75 (dt, J = 7.8, 4.7 Hz, 1H), 3.79 (s, 3H), 3.68 (s, 3H), 2.77 (dt, J = 4.7, 2.5 Hz, 2H), 2.41 (t, J = 7.2 Hz, 2H), 2.34 (t, J = 7.4 Hz, 2H), 2.03 (t, J = 2.6 Hz, 1H), 1.99 (q, J = 7.3 Hz, 2H). $^{13}$C NMR (125 MHz, $CDCl_3$) δ 173.69, 171.99, 170.99, 78.54, 71.80, 52.95, 51.75, 50.66, 35.32, 33.07, 22.56, 20.82.

**γ-glutaryl-L-propargylglycine (15)**

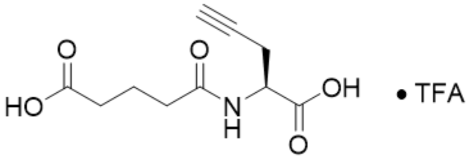


**15**

Compound was synthesized starting from methyl ester protected intermediate **SI-13** (0.020 g, 0.08 mmol) via BesE substrate general reaction C without modification to afford product **15** (0.087 g, 0.38 mmol, 20%). $^{1}$H NMR (500 MHz, $D_2O$ + 0.1% MeOH) δ 4.49 (t, *J* = 6.3 Hz, 1H), 2.77 – 2.67 (m, 2H), 2.42 (d, *J* = 7.6 Hz, 2H), 2.40 (s, 1H), 2.36 (t, *J* = 7.4 Hz, 2H), 1.88 (p, *J* = 7.6 Hz, 2H). $^{13}$C NMR (126 MHz, $D_2O$) δ 178.04, 175.97, 174.67, 79.95, 71.92, 52.05, 34.52, 32.90, 21.19, 20.65. MS (ESI) calculated for $C_{20}H_{21}N_4O_7^+$ (coumarin azide derivatized) 429.14, found 429.22 $(M+H)^+$.

#### Synthesis of *N*-butanoyl-L-propargylglycine (16)

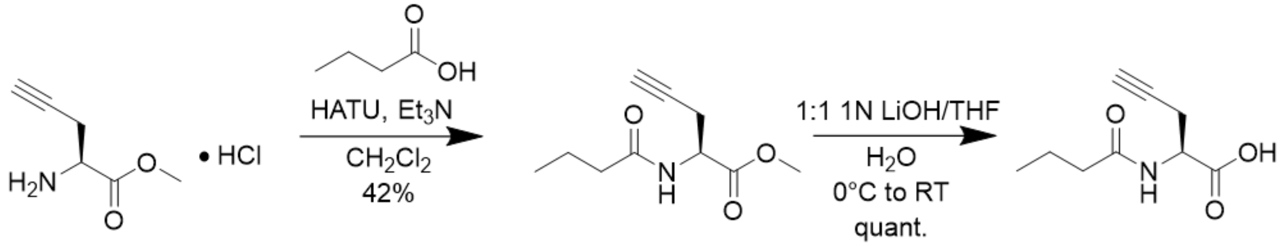


##### *N*-butanoyl-*O*-methyl-L-propargylglycine (SI-14)

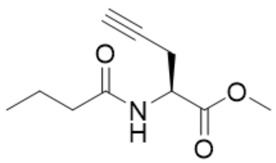

**SI-14**

Compound was synthesized via BesE substrate general reaction B without modification starting from **3** (0.105 g, 0.64 mmol) to afford protected intermediate **SI-14** (0.053 g, 0.27 mmol, 42%). $^1$H NMR (500 MHz, $CDCl_3$) δ 6.26 (s, 1H), 4.79 – 4.72 (m, 1H), 3.79 (s, 3H), 2.83 – 2.73 (m, 2H), 2.24 (t, *J* = 7.3 Hz, 2H), 2.03 – 2.01 (m, 1H), 1.69 (h, *J* = 7.2 Hz, 2H), 0.97 (t, *J* = 7.3 Hz, 3H). $^{13}$C NMR (125 MHz, $CDCl_3$) δ 172.84, 171.14, 78.63, 71.68, 52.93, 50.57, 38.56, 22.63, 19.15, 13.80.

##### *N*-butanoyl-L-propargylglycine (16)

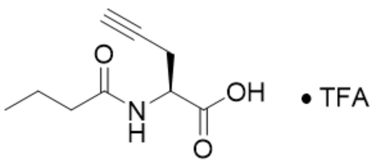


**16**

Compound was synthesized starting from methyl ester protected intermediate (**SI-14**) (0.053 g, 0.27 mmol) via BesE substrate general reaction C without modification to afford product **16** (0.135 g, 0.74 mmol, quant.). Given the purity of the resulting NMR analysis, the unexpected gain in mass is likely due to the presence of salts or incomplete lyophilizate of the final product. $^1$H NMR (500 MHz, $D_2O$ + 0.1% MeOH) δ 4.53 (dt, *J* = 9.0, 5.0 Hz, 1H), 2.73 (dq, *J* = 5.4, 2.7 Hz, 2H), 2.39 (s, 1H), 2.25 (dq, *J* = 7.5, 4.3, 3.6 Hz, 2H), 1.64 – 1.49 (m, 2H), 0.94 – 0.78 (m, 3H). $^{13}$C NMR (125 MHz, $D_2O$ + 0.1% MeOH) δ 177.40, 173.87, 79.62, 72.10, 51.43, 37.34, 20.91, 18.99, 12.81. MS (ESI) calculated for $C_{19}H_{21}N_4O_5^+$ (coumarin azide derivatized) 385.15, found 385.21 $(M+H)^+$.

#### Synthesis of β-aspartyl-L-propargylglycine (17)

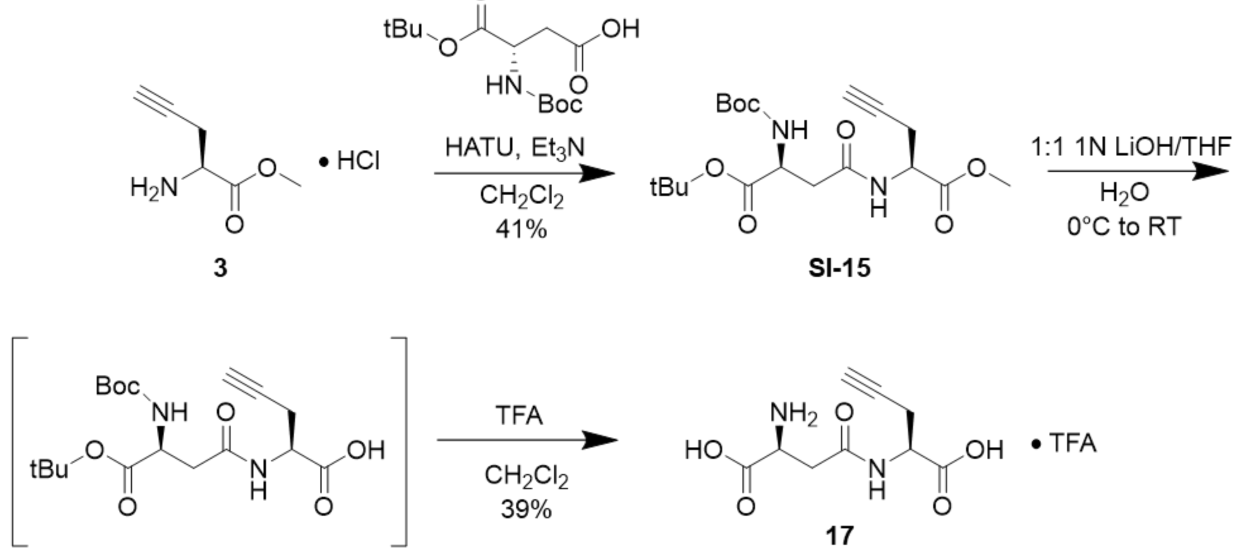


##### β-(*N*-Boc-*O*-*tert*-butyl- aspartyl)-*O*-methyl-L-propargylglycine (SI-15)

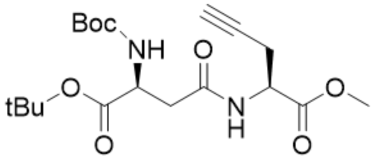


**SI-15**

Compound was synthesized via BesE substrate general reaction B without modification starting from **3** (0.024 g, 0.15 mmol) to afford protected intermediate **SI-15** (0.023 g, 0.06 mmol, 41%). $^1$H NMR (500 MHz, $CDCl_3$) δ 6.39 (s, 1H), 5.62 (s, 1H), 4.75 – 4.67 (m, 1H), 4.43 – 4.35 (m, 1H), 3.79 (s, 3H), 2.94 – 2.85 (m, 1H), 2.78 – 2.71 (m, 3H), 2.03 (s, 1H), 1.46 (s, 9H), 1.44 (s, 9H).$^{13}$C NMR (125 MHz, $CDCl_3$) δ 175.26, 170.71, 170.31, 82.81, 82.31, 78.43, 71.93, 53.00, 50.97, 50.82, 44.59, 38.22, 28.48, 28.05, 22.55.

**β-aspartyl-L-propargylglycine (17)**

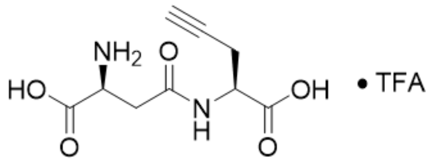

**17**

Compound was synthesized starting from methyl ester protected intermediate **SI-15** (0.035 g, 0.08 mmol) via BesE substrate general reactions C and D with slight modification to reaction D only: The TFA deprotection was allowed to run overnight. Concentration *in vacuo* and lyophilization yielded product **17** (0.007 g, 0.03 mmol, 39%). $^{1}$H NMR (500 MHz, $D_2O$ + 0.1% MeOH) δ 4.62 – 4.55 (m, 1H), 4.30 (q, *J* = 4.7, 4.0 Hz, 1H), 3.06 (d, *J* = 5.2 Hz, 2H), 2.80 – 2.71 (m, 2H), 2.42 (h, *J* = 3.1, 2.3 Hz, 1H). $^{13}$C NMR (126 MHz, $D_2O$ + 0.1% MeOH) δ 173.77, 171.39, 170.94, 79.47, 72.18, 51.57, 49.83, 34.23, 21.07. MS (ESI) calculated for $C_{18}H_{21}N_6O_{10}^+$ (Marfey's derivatized) 481.13, found 481.18 $(M+H)^+$.

#### Synthesis of γ-D-glutamyl-L-propargylglycine (18)

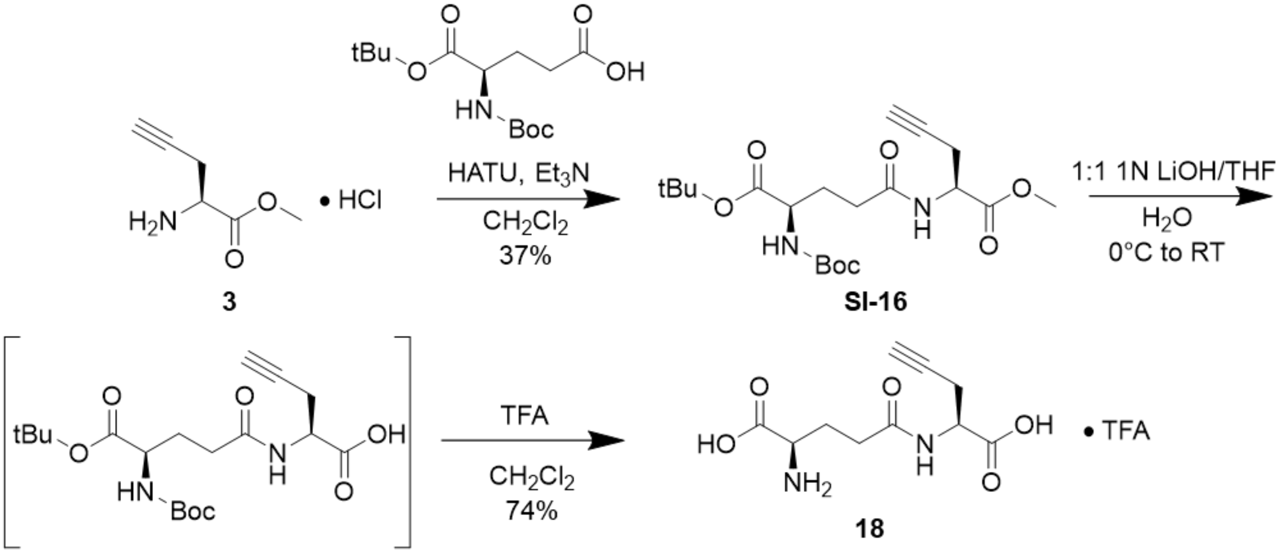


**γ-(*N*-Boc-*O*-*tert*-butyl-D-glutamyl)-L-propargylglycine (SI-16)**

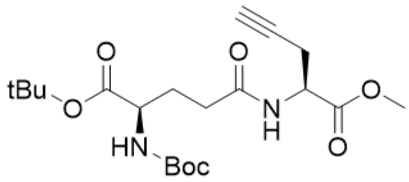

**SI-16**

Compound was synthesized via BesE substrate general reaction B without modification starting from **3** (0.109 g, 0.67 mmol) to afford protected intermediate **SI-16** (0.102 g, 0.25 mmol, 37%). $^1$H NMR (500 MHz, $CDCl_3$) δ 6.84 (s, 1H), 5.24 (s, 1H), 4.78 – 4.70 (m, 1H), 4.26 – 4.16 (m, 1H), 3.78 (s, 3H), 2.81 – 2.70 (m, 2H), 2.39 – 2.27 (m, 2H), 2.21 – 2.11 (m, 1H), 2.02 (s, 1H), 1.96 – 1.82 (m, 1H), 1.46 (s, 9H), 1.43 (s, 9H).$^{13}$C NMR (125 MHz, $CDCl_3$) δ 176.30, 172.20, 171.58, 170.98, 82.40, 80.07, 78.62, 71.73, 53.57, 52.89, 50.89, 32.60, 29.35, 28.44, 28.13, 22.44.

**γ-D-glutamyl-L-propargylglycine (18)**

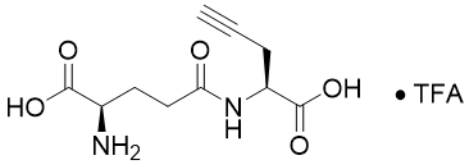


**18**

Compound was synthesized starting from methyl ester protected intermediate **SI-16** (0.102 g, 0.08 mmol) via BesE substrate general reactions C and D with slight modification to reaction D only: The TFA deprotection was allowed to run overnight. Concentration *in vacuo* and lyophilization yielded product **18** (0.015 g, 0.06 mmol, 74%). H NMR (500 MHz, $D_2O$ + 0.1% MeOH) δ 4.58 (q, *J* = 5.4 Hz, 1H), 4.12 – 4.06 (m, 1H), 2.82 – 2.72 (m, 2H), 2.65 – 2.52 (m, 3H), 2.42 (s, 1H), 2.29 – 2.11 (m, 3H). $^{13}$C NMR (125 MHz, $D_2O$ + 0.1% MeOH) δ 174.32, 173.78, 171.62, 79.56, 72.15, 52.29, 51.54, 30.82, 25.61, 20.94. MS (ESI) calculated for $C_{19}H_{23}N_6O_{10}^+$ (Marfey's derivatized) 495.15, found 495.19 $(M+H)^+$.

#### Tables

| | |
|---|---|
| BA-T150A-BesE-F | GAATGATGGCgcgTTCATCCATT |
| BA-T150A-BesE-R | ACTTCGCGCAACATACCAACG |
| GB-R144A-BesE-F | TGGTATGTTGgcgGAAGTGAATGATGGC |
| GB-R144A-BesE-R | ACGAATGCCGGTCTCCCTCC |
| AN- F232A-BesE-F | TGCCTTGCTCgcgAATGCAAATAATTATCATGTTGTGCACCCGG |
| AN- F232A-BesE-R | TCGCCGGCGGCTGGC |
| CG-F255A-BesE-F | CCTTGCCTGTgcgCTGGGGGTCA |
| CG-F255A-BesE-R | GCAATACGGCGTTGACCAGGTGCACC |
| IG-N177A-BesE-F | ACTGGCATTCgcgGCGTGGTTAG |
| IG-N177A-BesE-R | TGAGCTACAATCTTCTGATCAAACAGGCCGC |
| SB-R249A-BesE-F | TGGTCAACGCgcgATTGCCCTTGCCTGTTTCC |
| SB-R249A-BesE-R | GGTGCACCCGGGTGCACAACATG |
| MD-A175W-BesE-F | AGCTCAACTGtggTTCAACGCGTG |
| MD-A175W-BesE-R | ACAATCTTCTGATCAAACAGGCCGC |
| EP-Q173A-BesE-F | GATTGTAGCTgcgCTGGCATTCAAC |
| EP-Q173A-BesE-R | TTCTGATCAAACAGGCCGCC |
| EA-V146A-BesE-F | GTTGCGCGAAgcgAATGATGGCA |
| EA-V146A-BesE-R | ATACCAACGAATGCCGGTCTCCCTC |
| LD-Y207A-BesE-F | TCGCCATGGAgcgGGATTTCAAC |
| LD-Y207A-BesE-R | CGATTTTCATCTGCCGGTTCCCAG |
| KH-D155A-BesE-F | CATCCATTATgcgGACATTAACCGC |
| KH-D155A-BesE-R | AACGTGCCATCATTCACTTCGC |
| CC-V240A-BesE-F2 | TTA TCA TGT TgcgCA CCC GGG TGC ACC |
| CC-V240A-BesE-R2 | TTATTTGCATTGAAGAGCAAGGCATCGCCGG |
| BA-M70A-F | GCATCATCCGgcgGCCCGCTTTG |
| BA-M70A-R | ATGCGATCCAGATCAAAGCTGACAACTGGC |

**Table S1.** Primers used in this study. Red indicates the altered codon for mutagenesis.

| Student | Analogs synthesized | BesE mutants |
|---|---|---|
| Beau Andre | Efforts towards cyanoalanine analog (Figure S3) | M70, T150A |
| Emily Avalos | Analog 6 | V146A |
| Surina Beal | Analog 5 | R249A |
| George Beck | Analog 12 | R144A |
| Charles Choi | Analog 18 | V240A |
| Elizabeth Dolzhansky | Analog 17 | Y207A |
| Michelle Du | Efforts towards analog 15 | A175W |
| Isabella Gallardo | Analog 13 | N177A |
| Cyrus Ghorbani | Analog 7 | F255A |
| Kyle Hinaga | Analog 9 | D155A |
| Analynn Nguyen | Analog 16 | F232A |
| Edward Pham | Analog 14 | Q173A |

**Table S2.** Individual CURE lab student contributions.

#### NMR Spectra of Characterized Compounds

##### γ-glutamyl-L-propargylglycine (1)

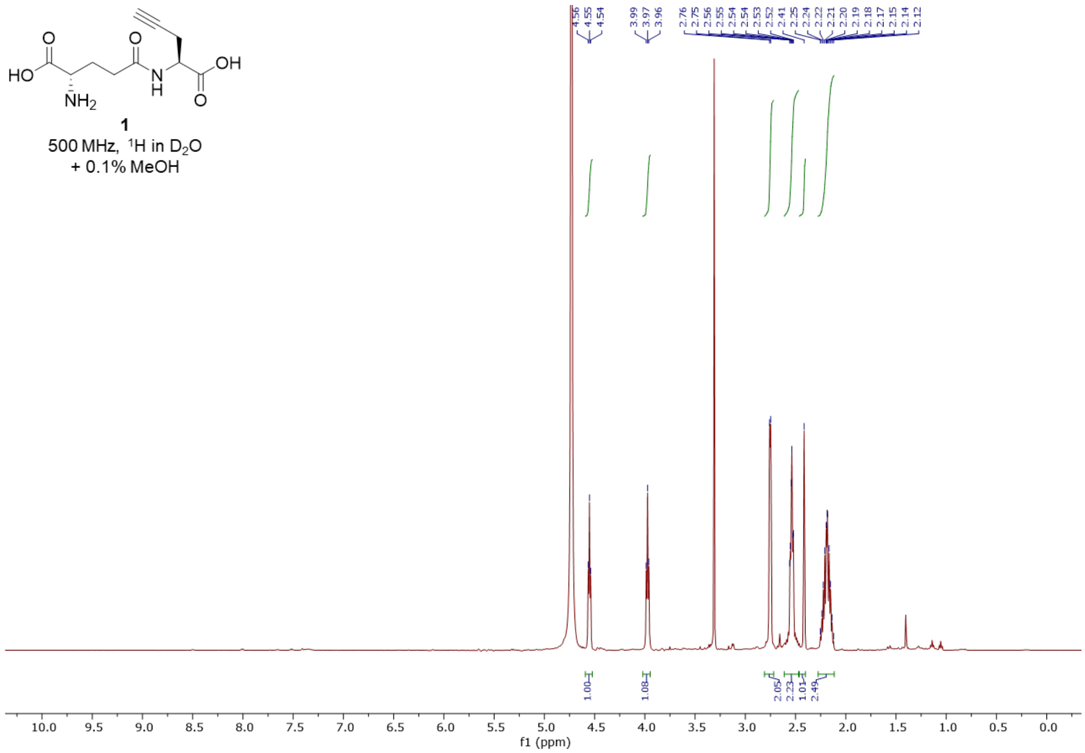


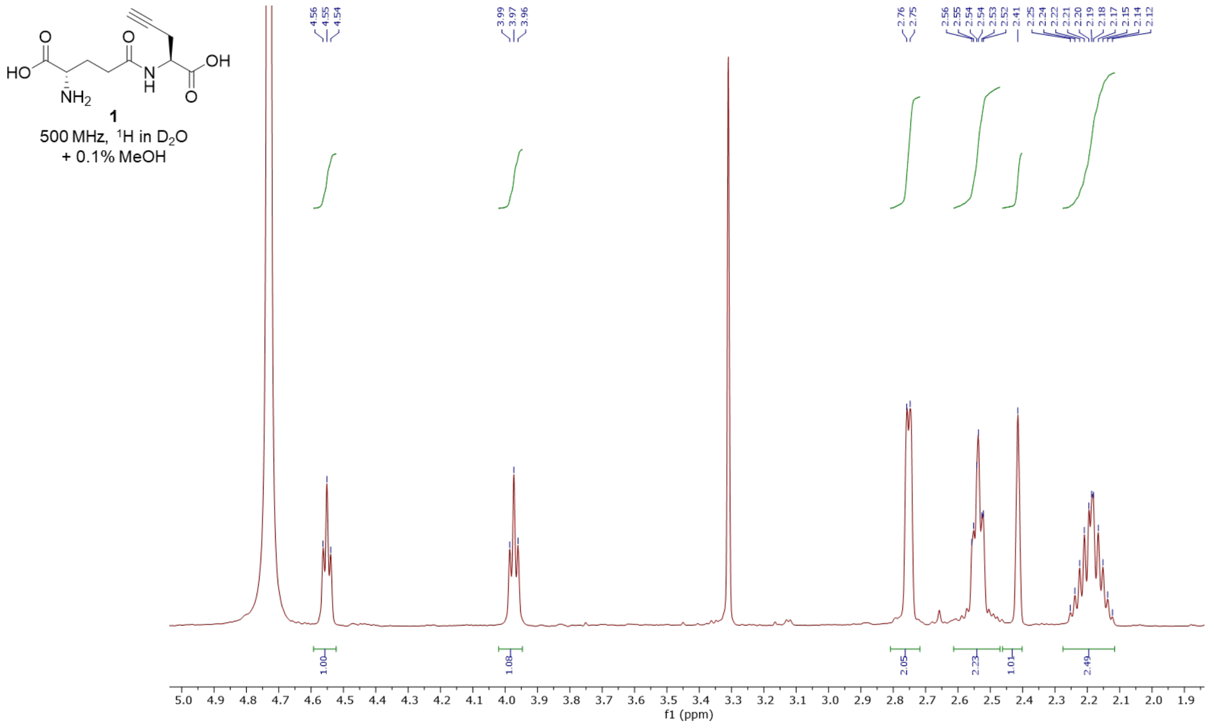

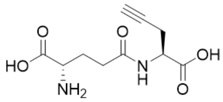

**1**

125 MHz, $^{13}C$ in $D_2O$
+ 0.1% MeOH

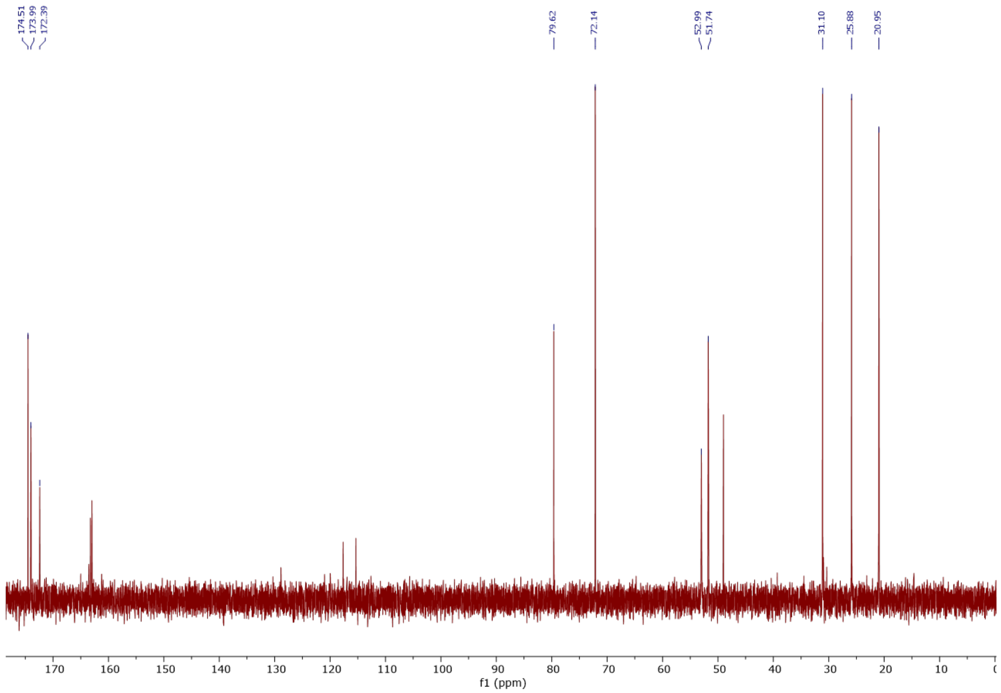

#### *O*-methyl-L-propargylglycine (3)

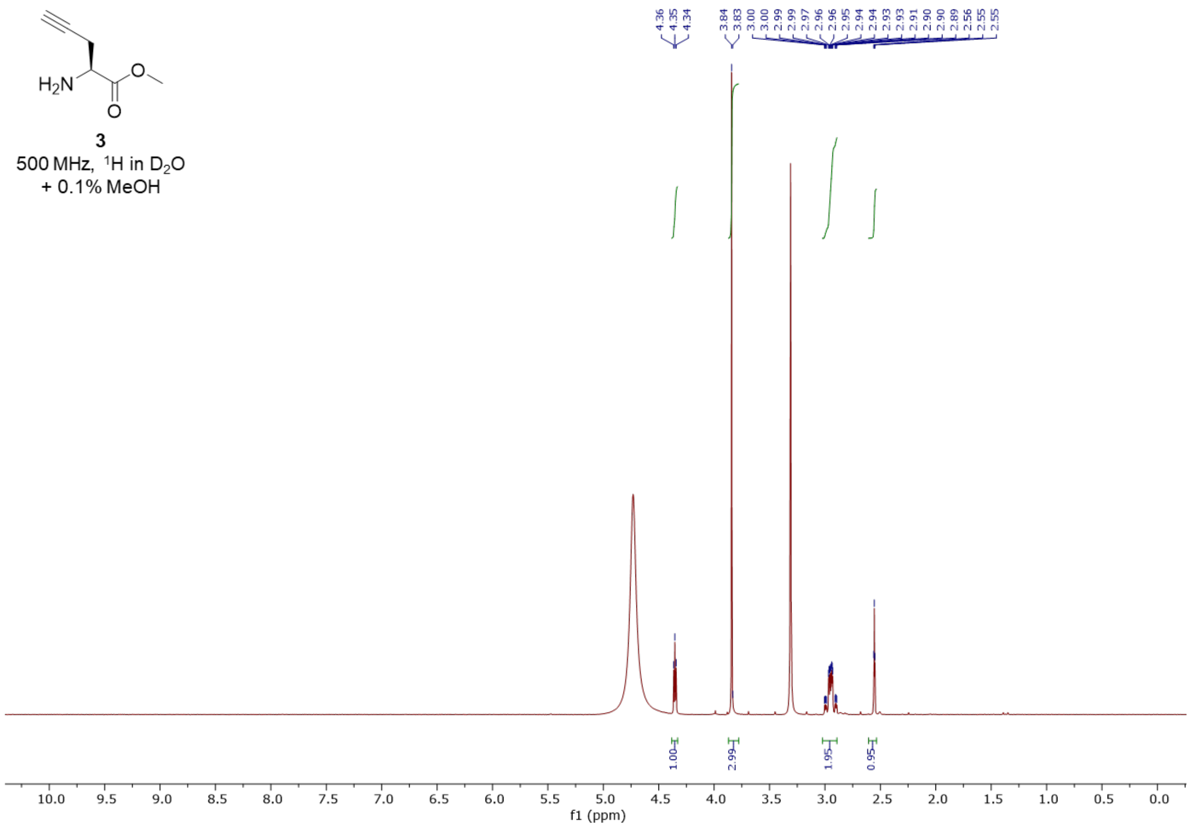


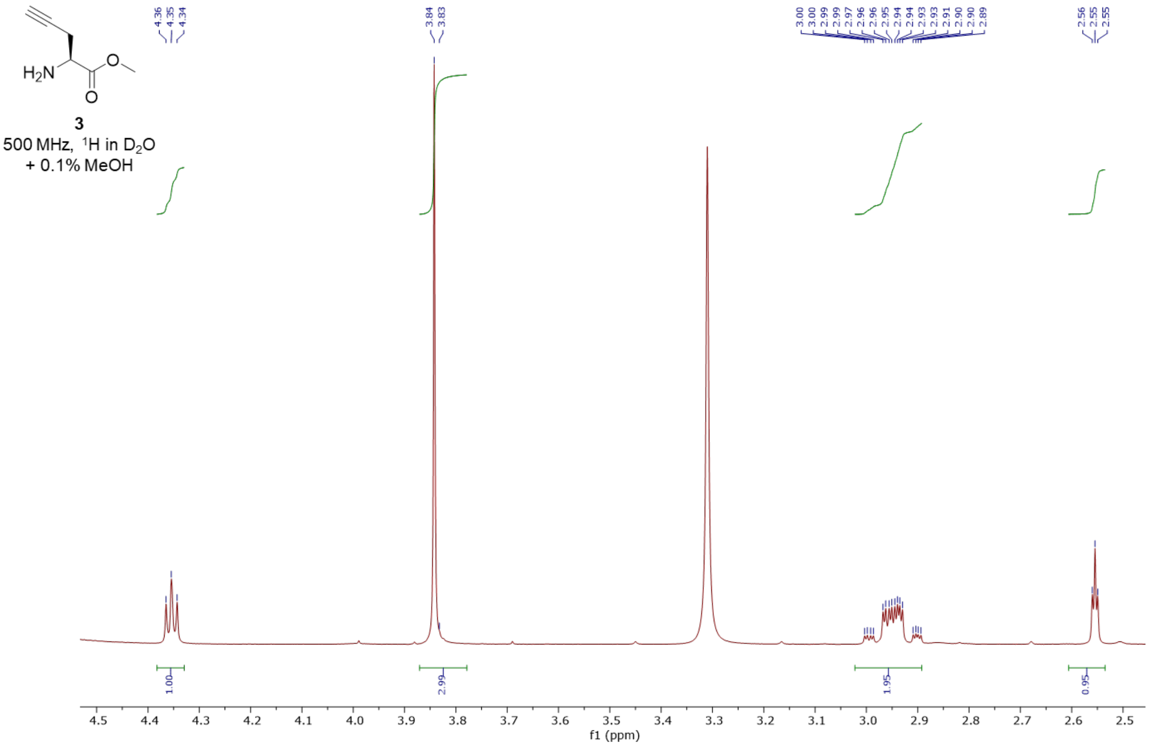

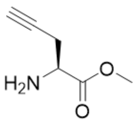


**3**
125 MHz, $^{13}$C in $D_2O$
+ 0.1% MeOH

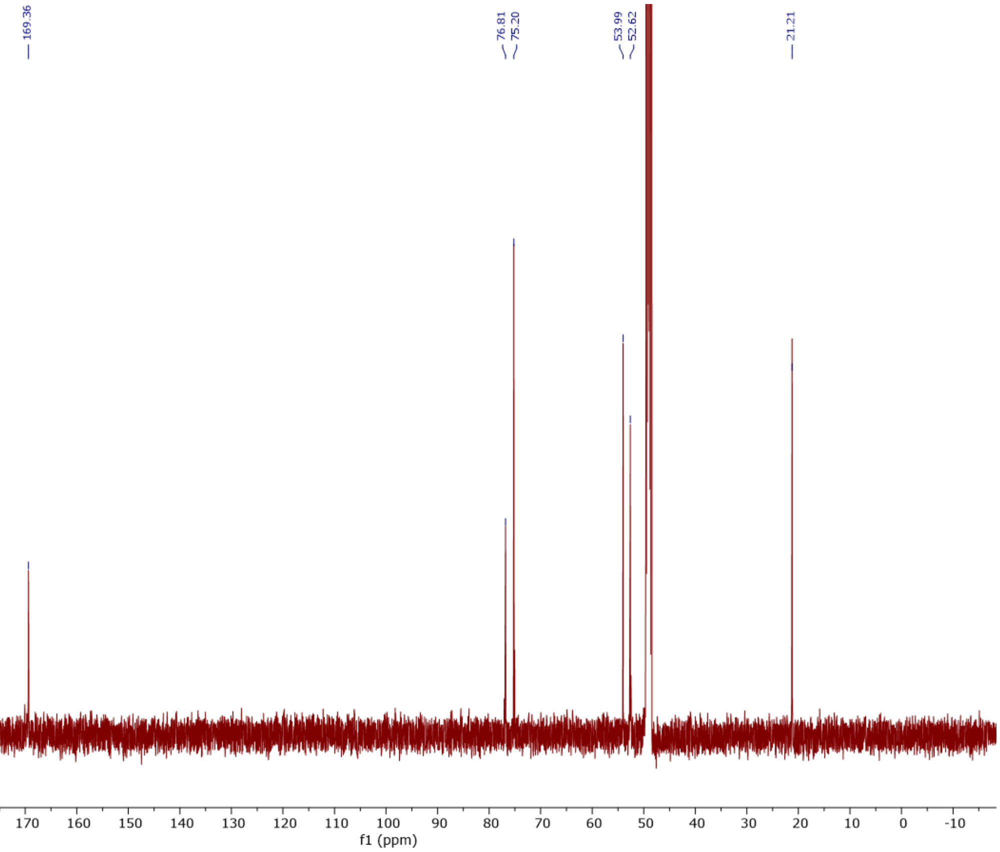

#### γ-(*N*-Boc-*O*-tert-butyl-glutamyl)-*O*-methyl-L-propargylglycine (4)

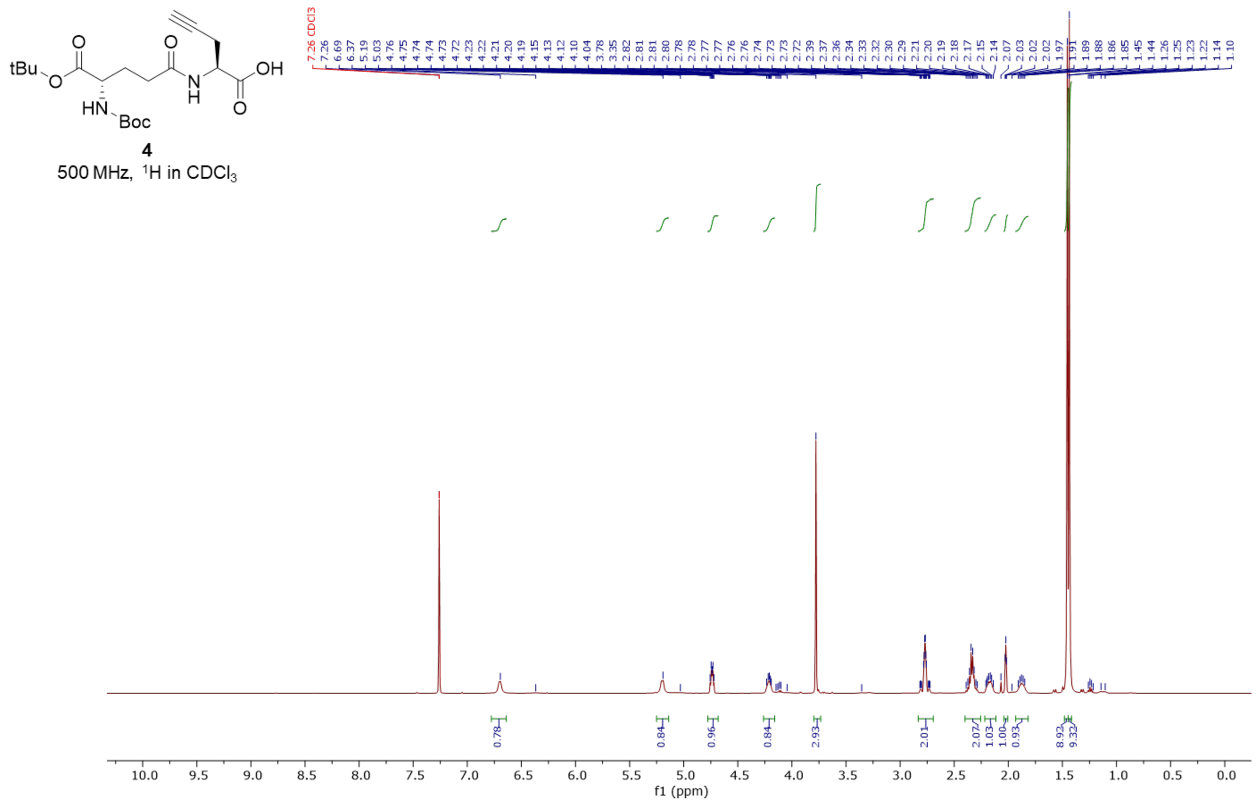


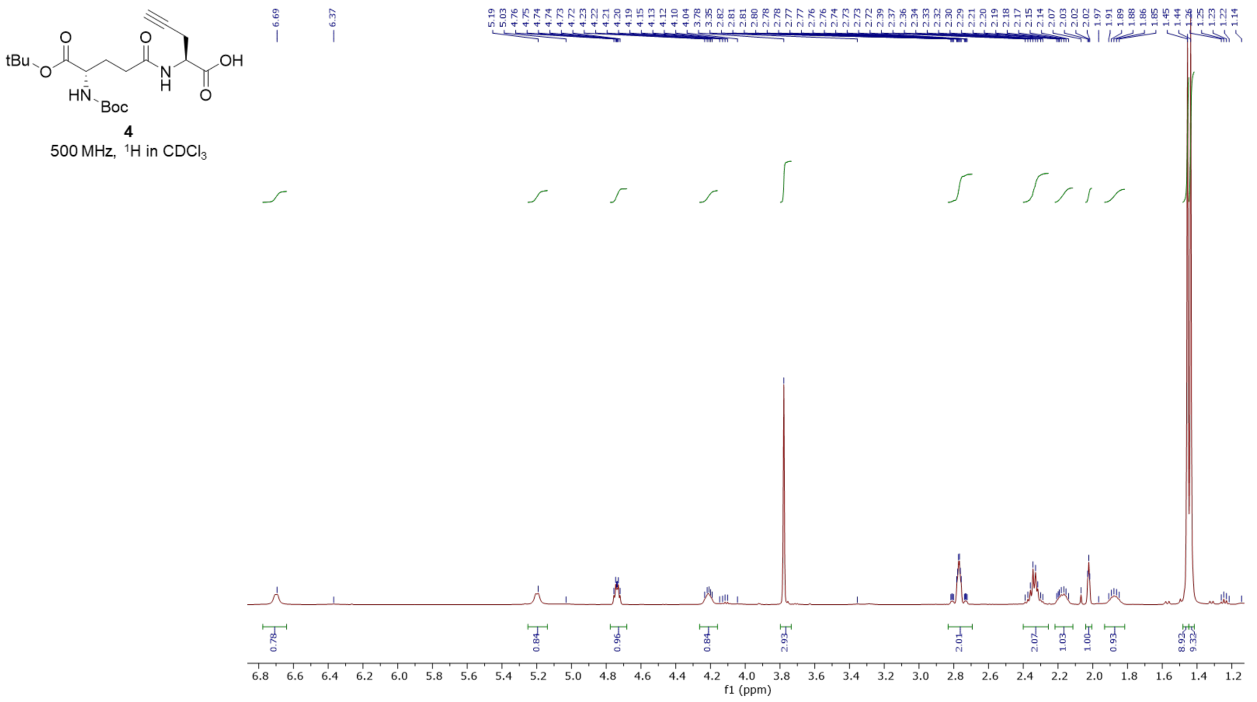

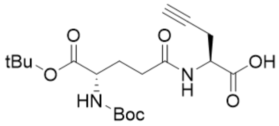


**4**

125 MHz, $^{13}C$ in $CDCl_3$

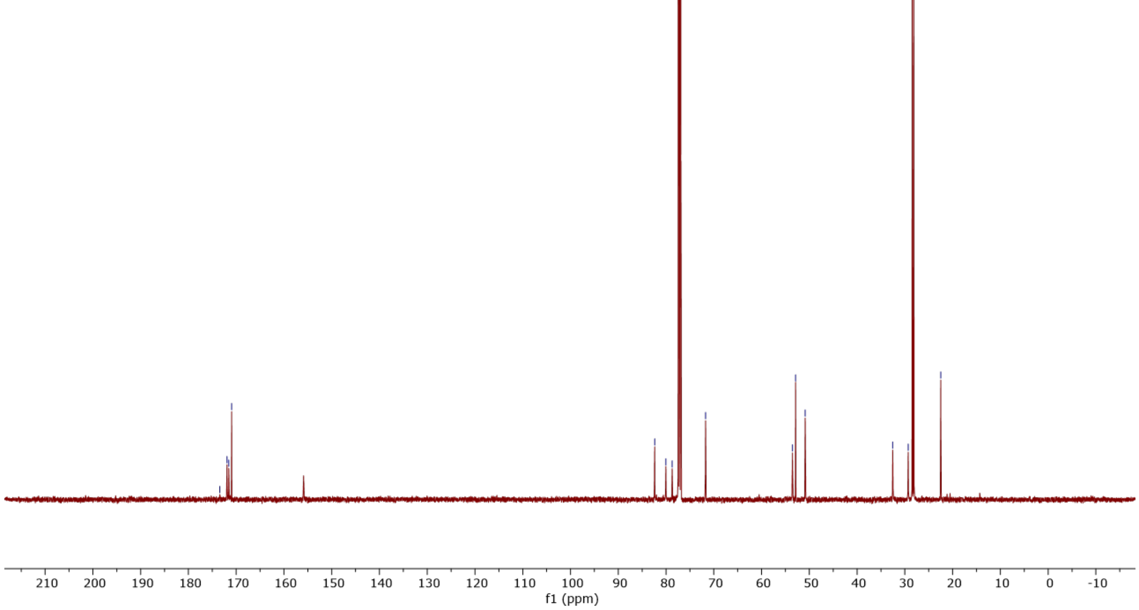

### γ-glutamyl-L-allylglycine (5)

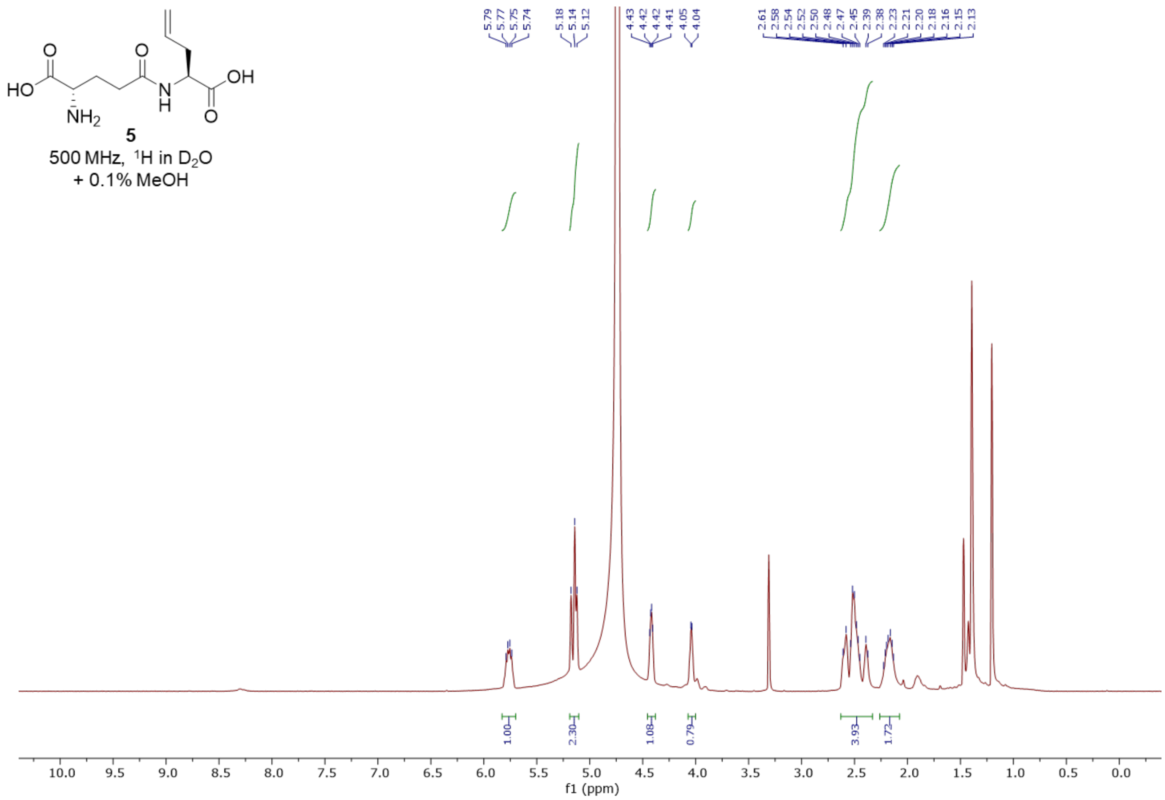


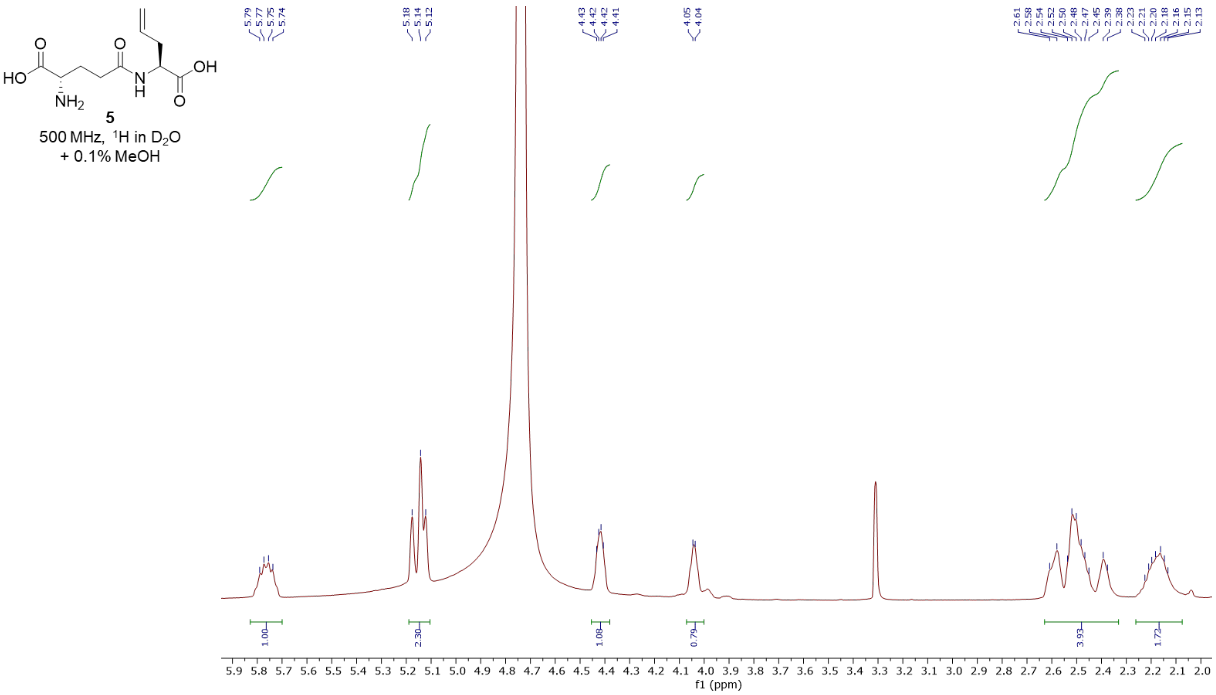

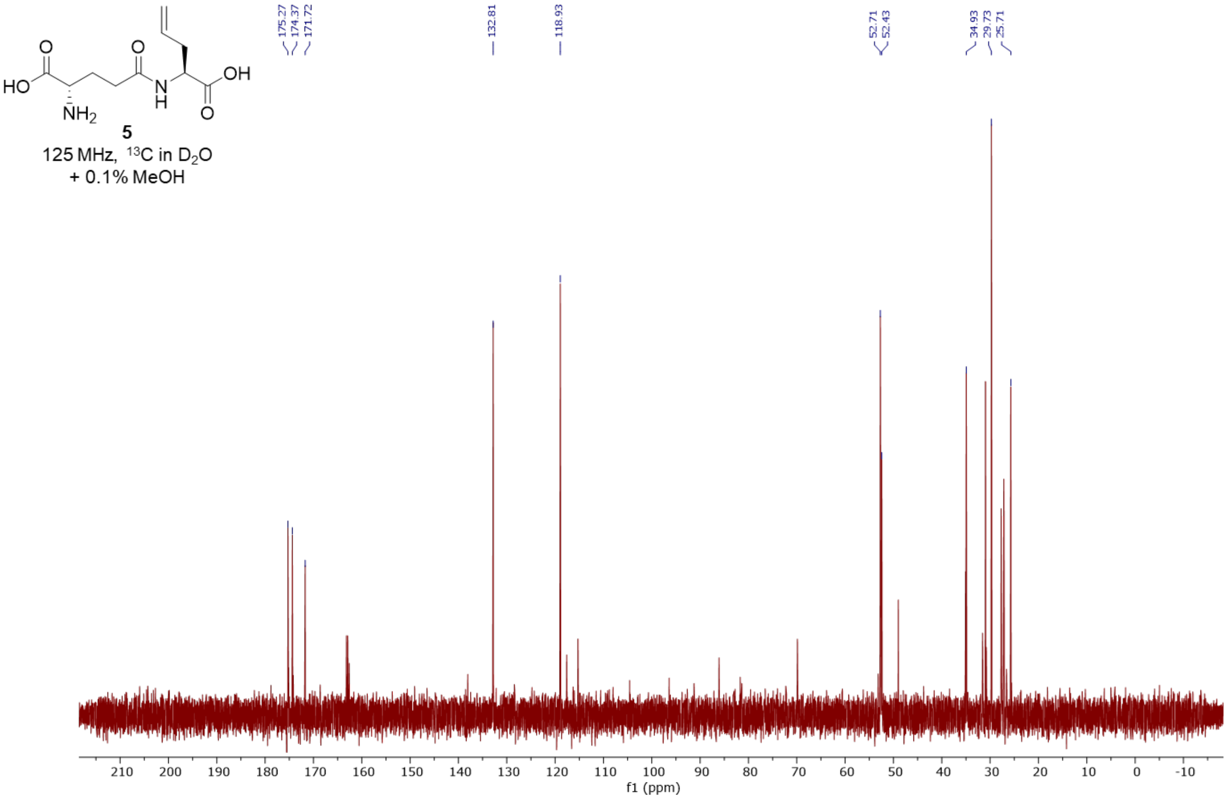


**5**

125 MHz, $^{13}C$ in $D_2O$
+ 0.1% MeOH

#### γ-glutamyl-L-norvaline (6)

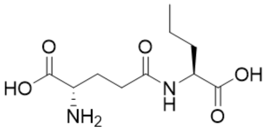

**6**
500 MHz, $^{1}$H in $D_2O$
+ 0.1% MeOH

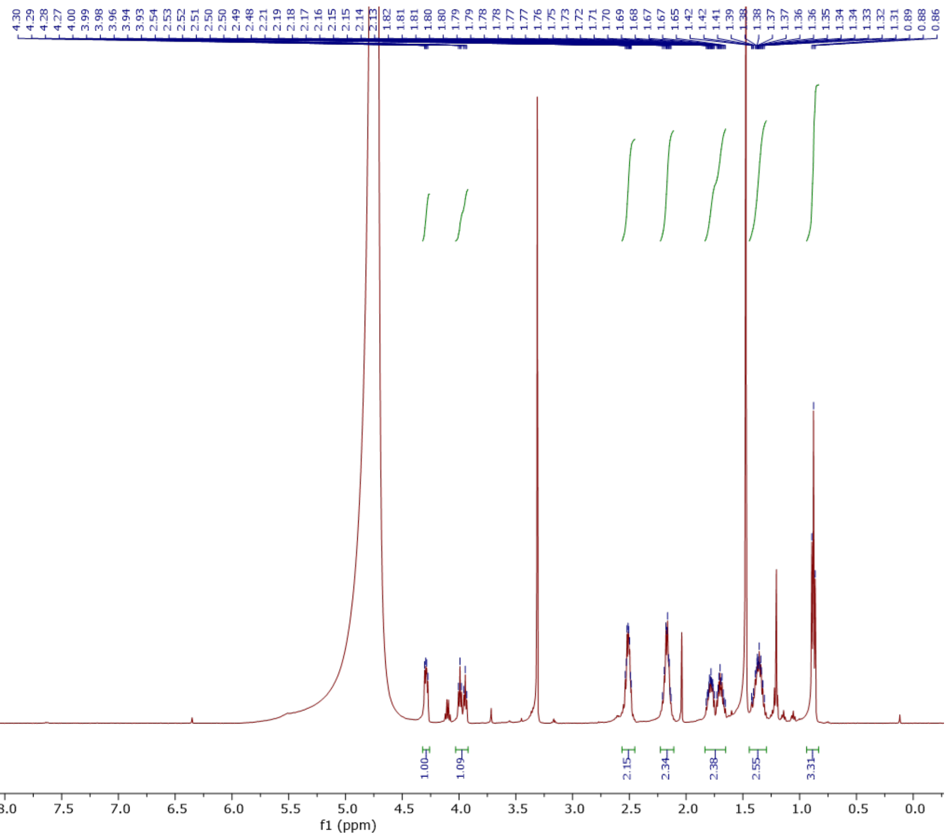


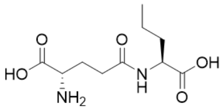

**6**
500 MHz, $^{1}$H in $D_2O$
+ 0.1% MeOH

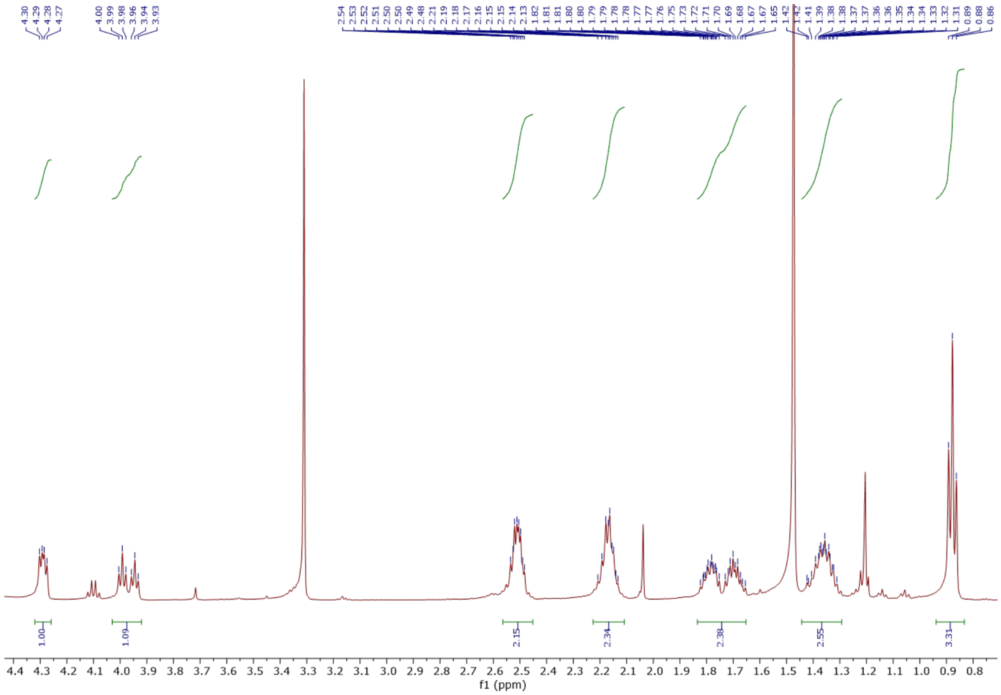

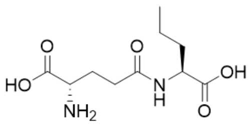


**6**

125 MHz, $^{13}C$ in $D_2O$
+ 0.1% MeOH

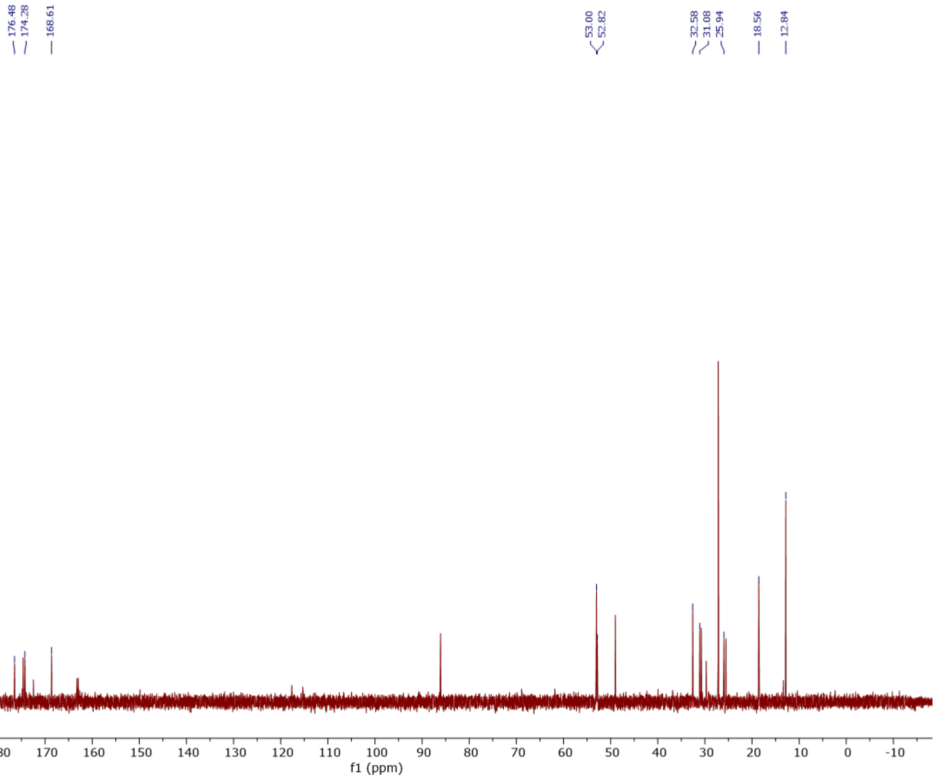

### γ-glutamyl-L-homopropargylglycine (7)

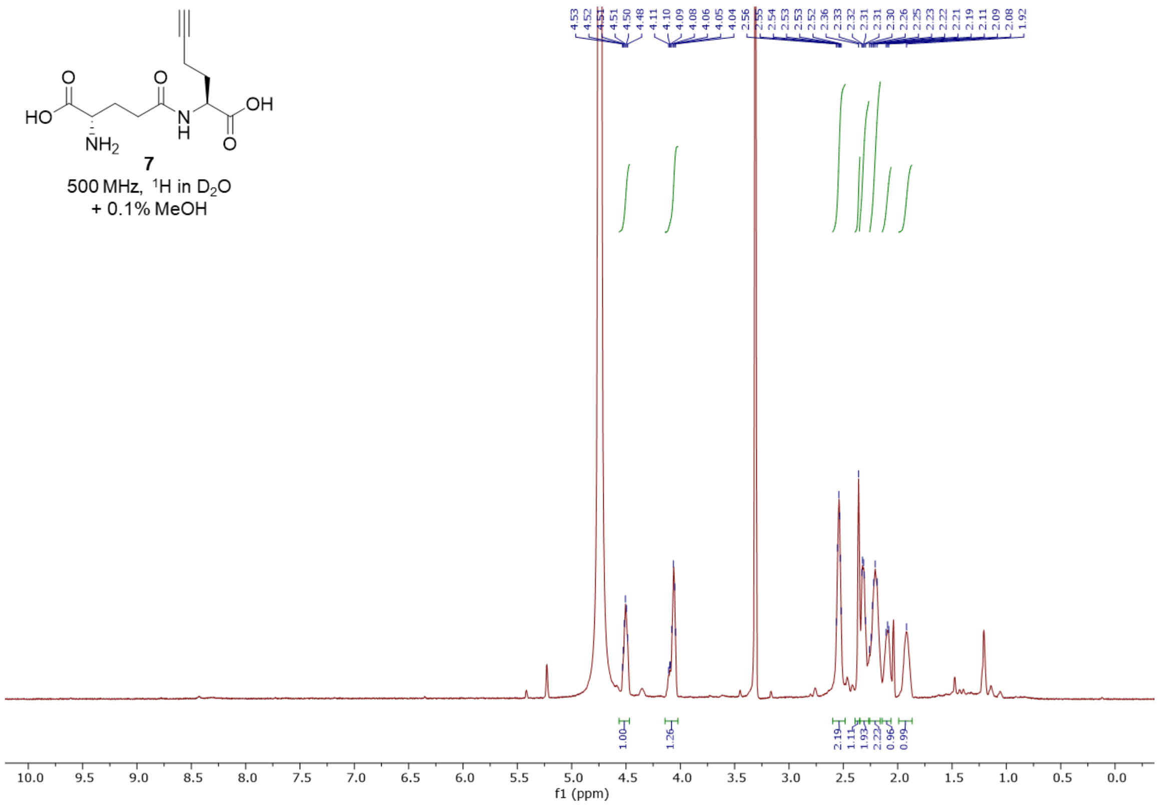


**7**
500 MHz, $^1H$ in $D_2O$
+ 0.1% MeOH

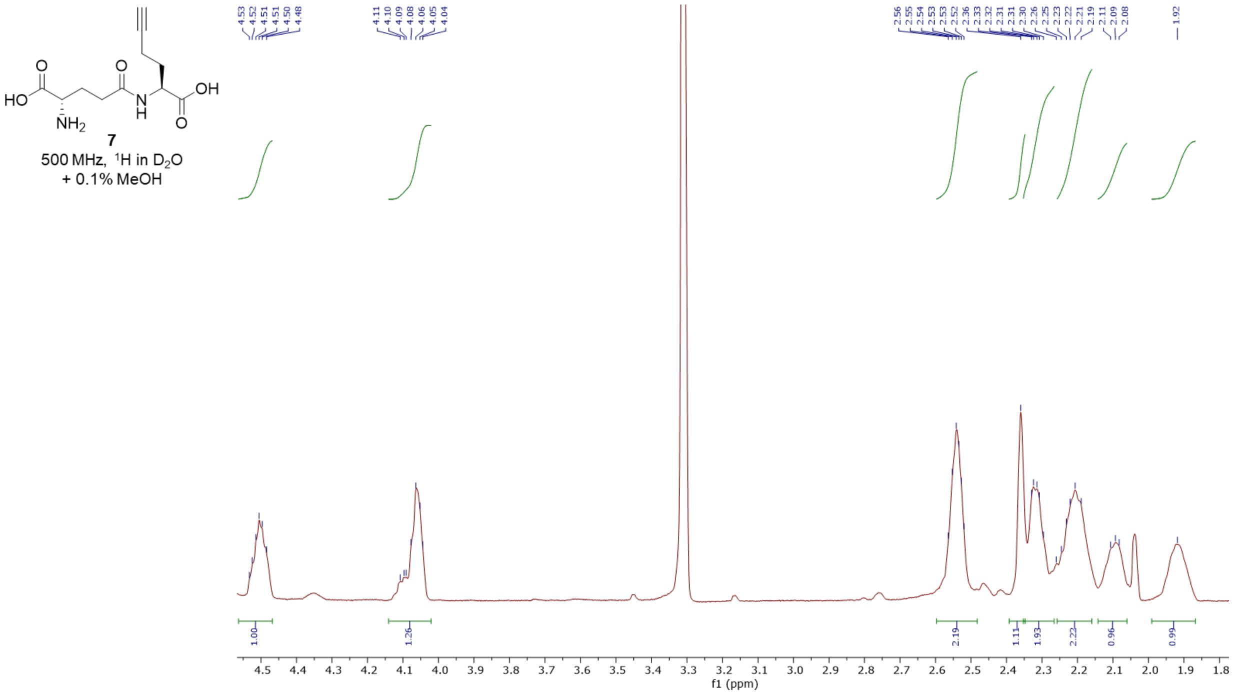


**7**
500 MHz, $^1H$ in $D_2O$
+ 0.1% MeOH

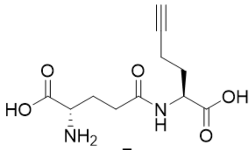


**7**

125 MHz, ${}^{13}$C in $D_2O$
+ 0.1% MeOH

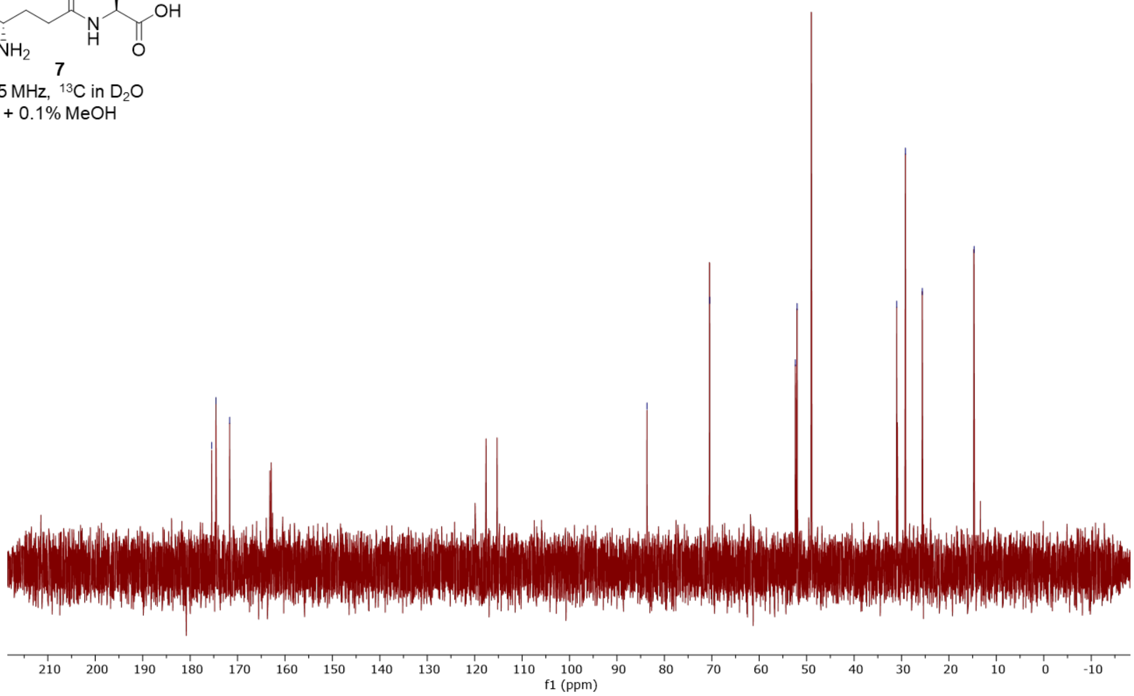

**γ-glutamyl-L-alanine (8)**

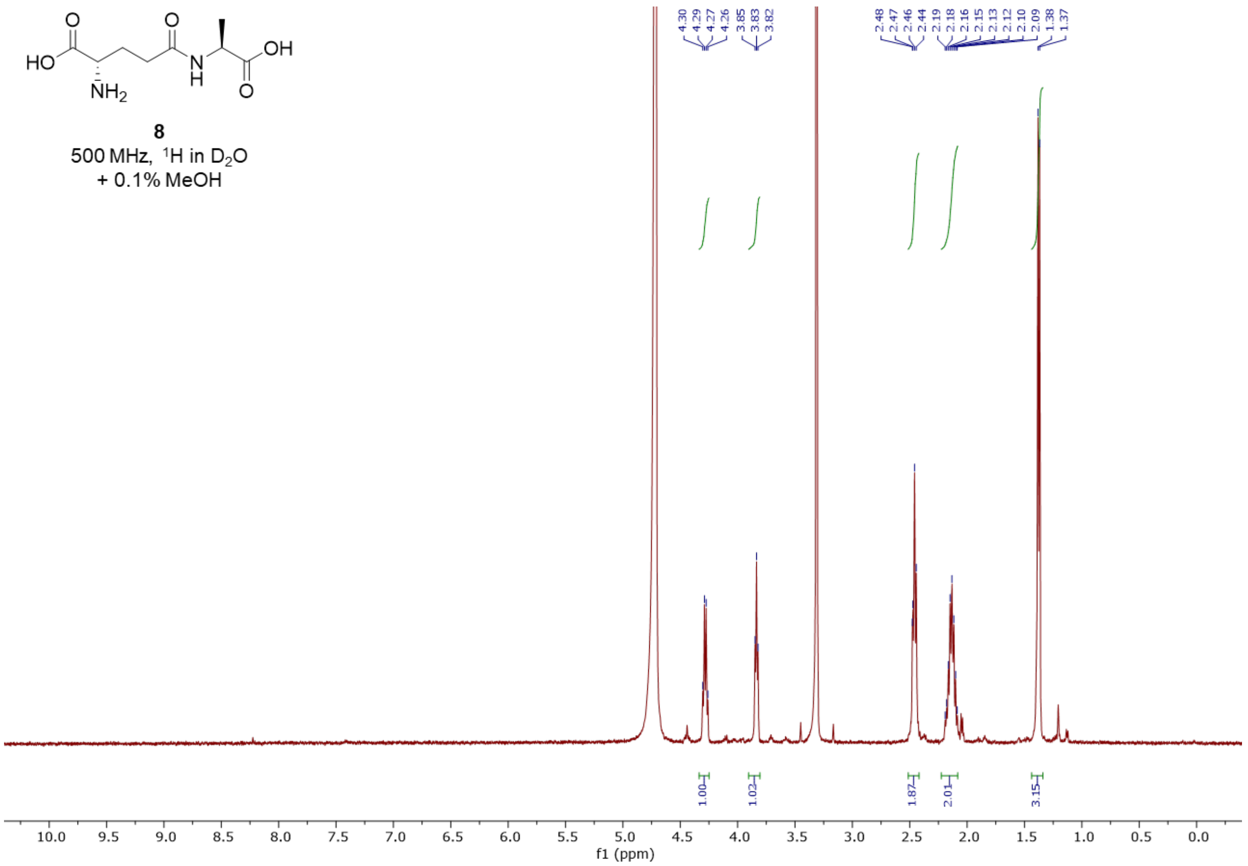


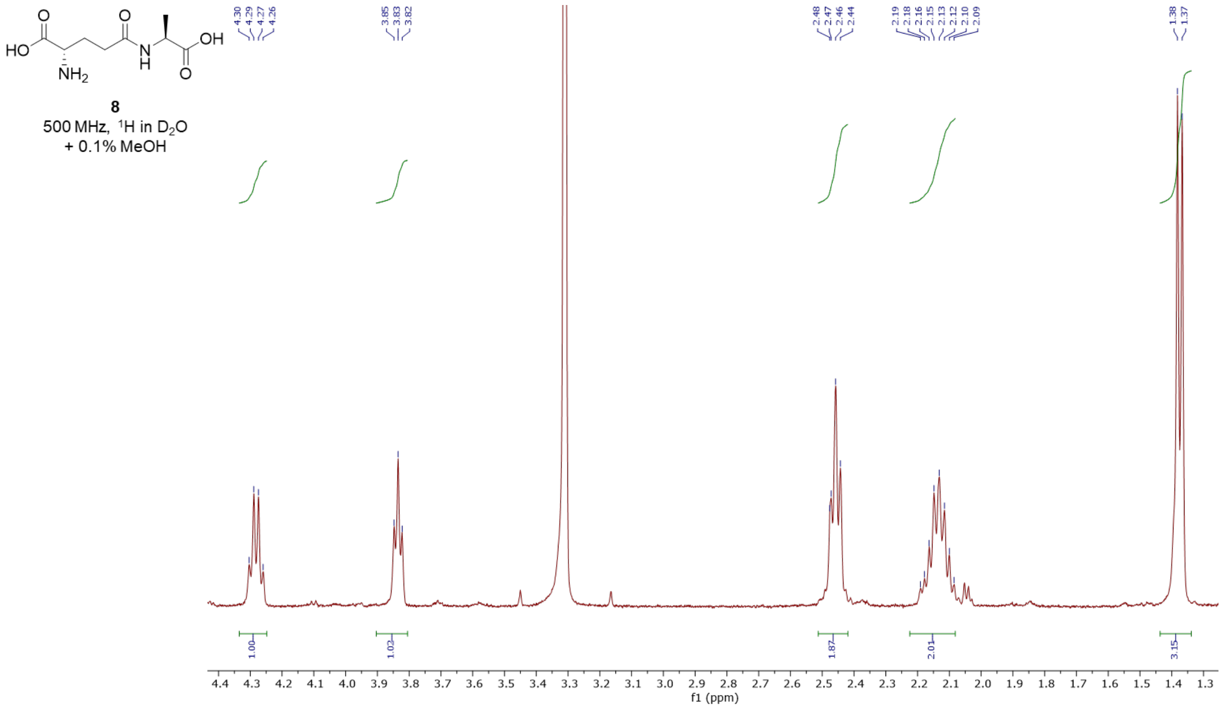

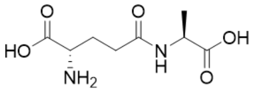


**8**

125 MHz, $^{13}C$ in $D_2O$
+ 0.1% MeOH

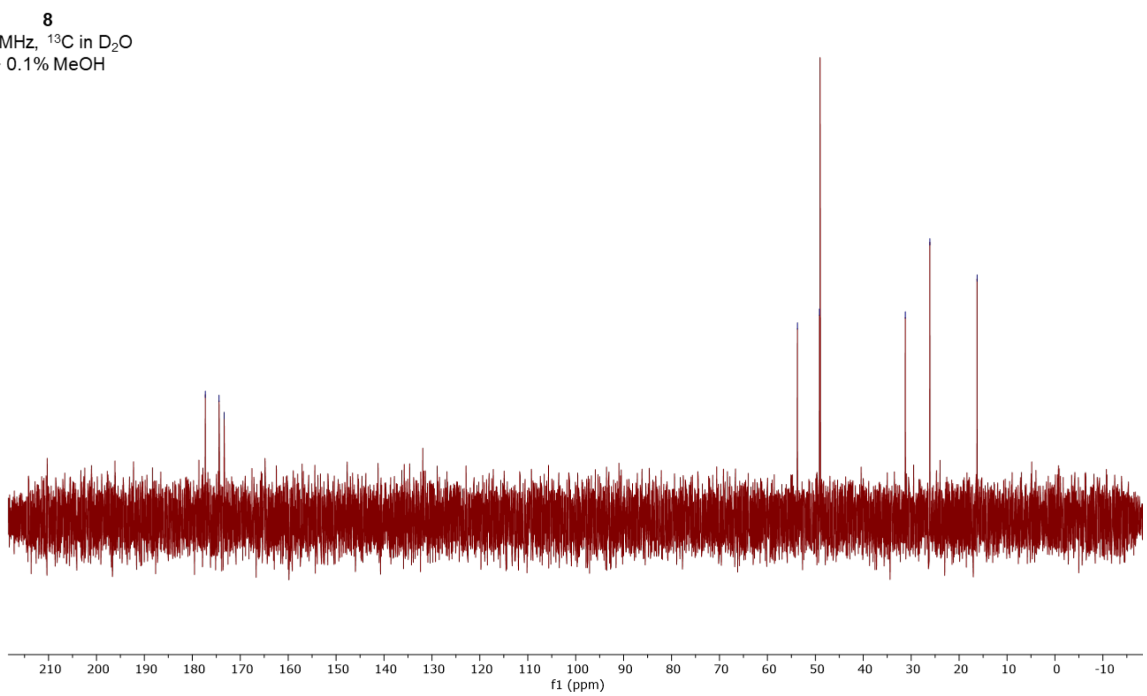

#### γ-glutamyl-D-propargylglycine (9)

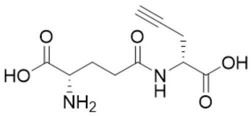

**9**

500 MHz, $^{1}H$ in $D_2O$
+ 0.1% MeOH

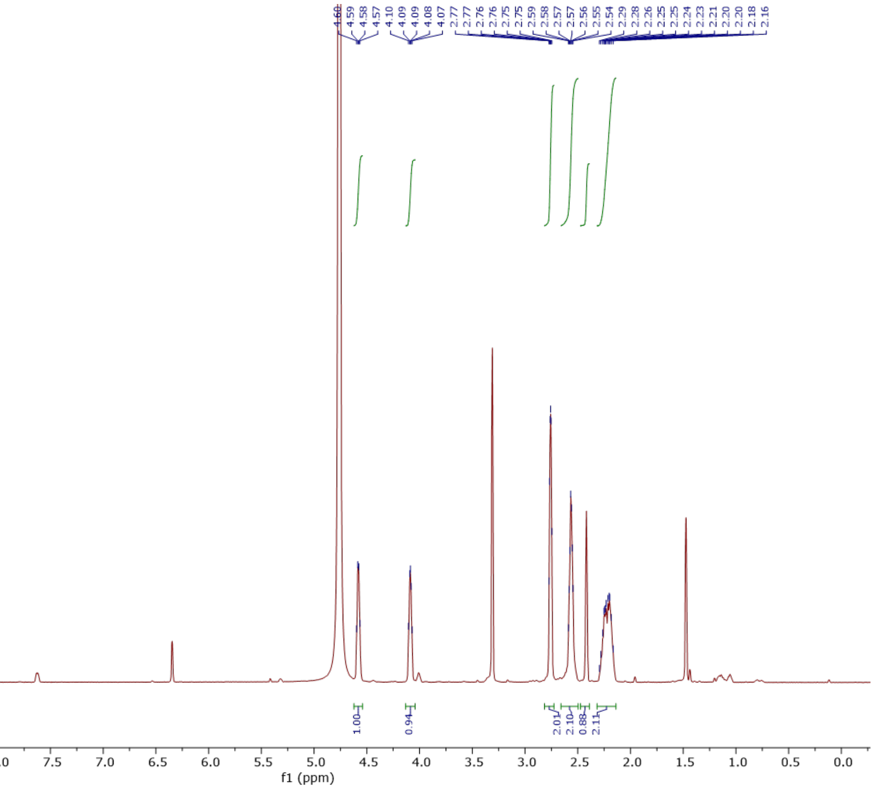


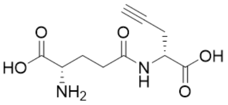

**9**

500 MHz, $^{1}H$ in $D_2O$
+ 0.1% MeOH

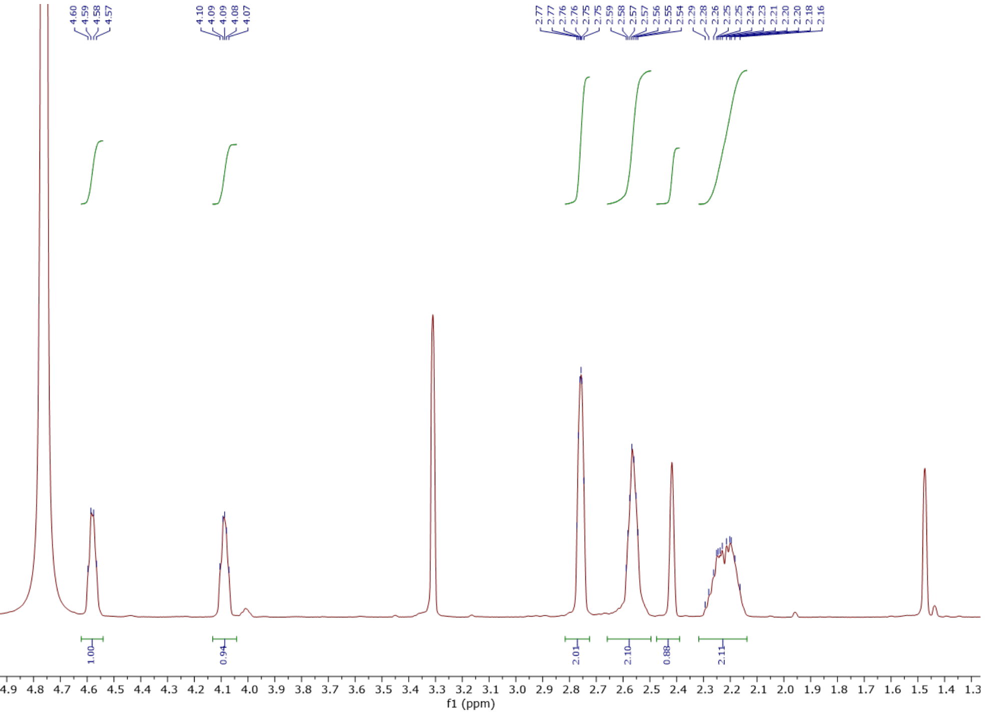

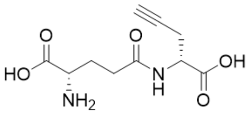


**9**

125 MHz, $^{13}C$ in $D_2O$
+ 0.1% MeOH

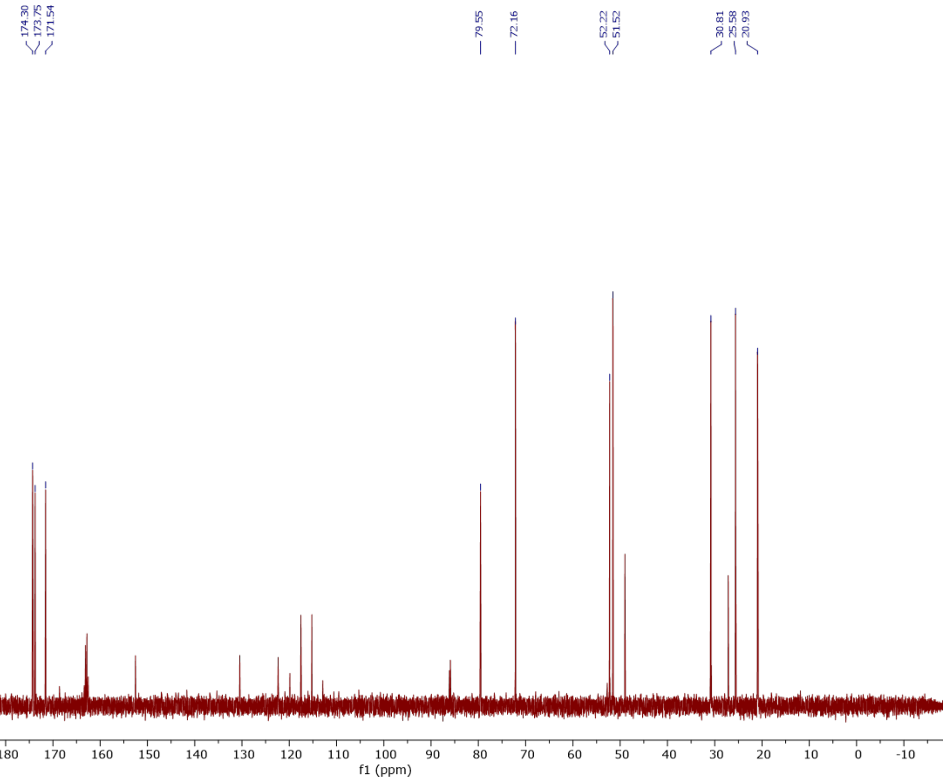

### γ-glutamyl-L-propargylglycine methyl ester (10)

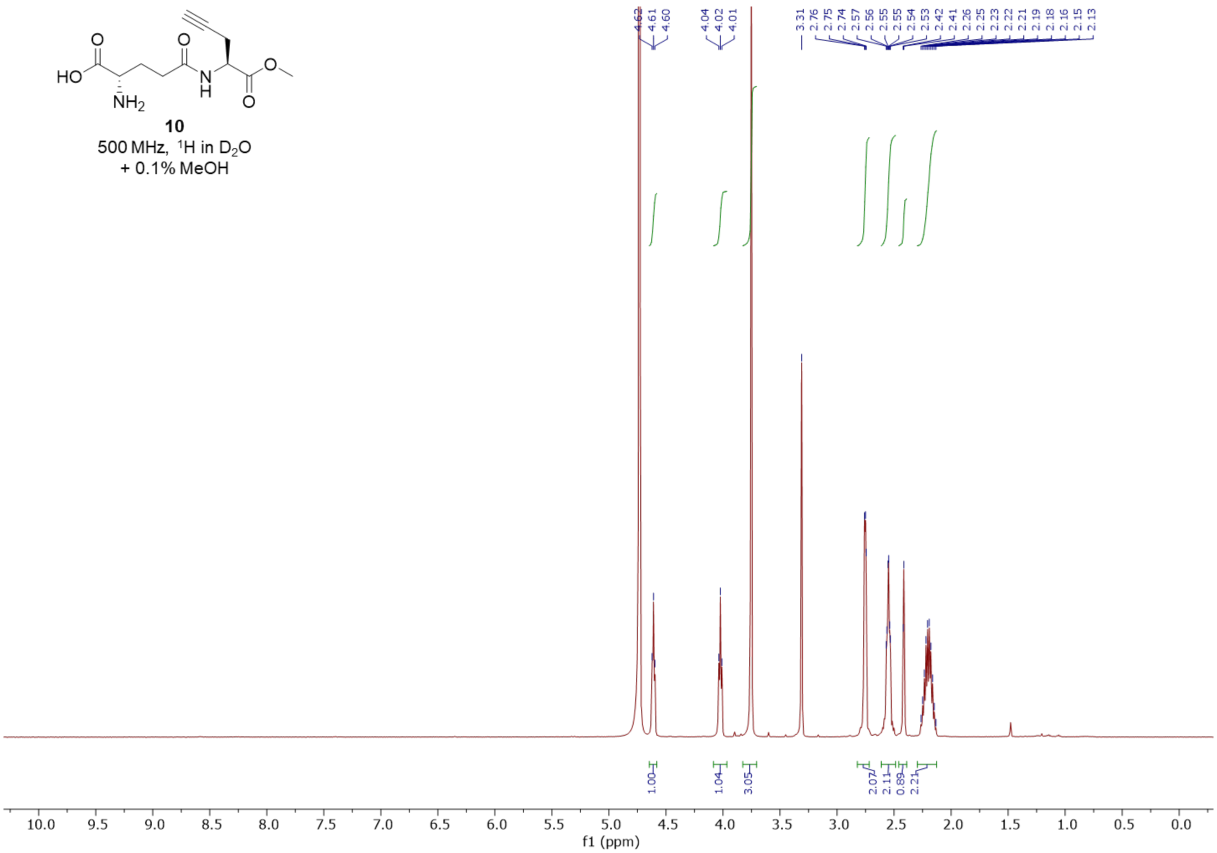


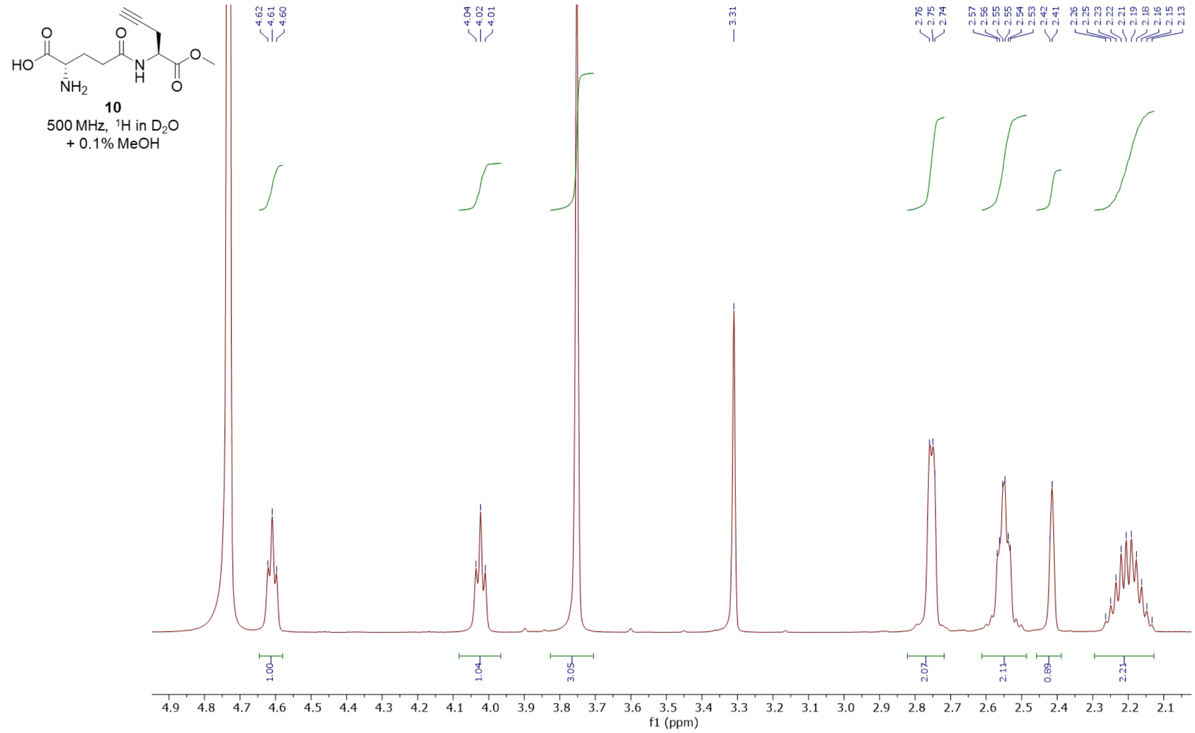

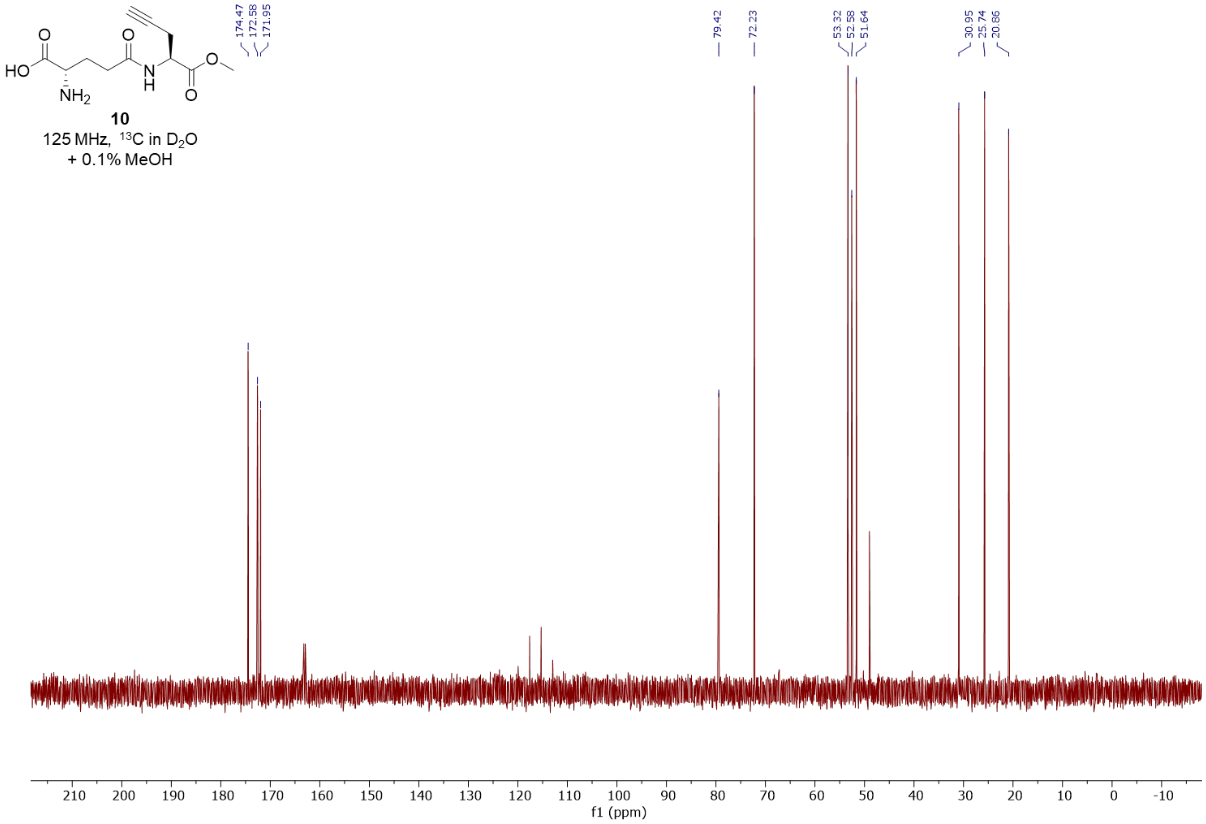
HO
NH2
O
O
N
H
O
O
10
125 MHz, 13C in D2O
+ 0.1% MeOH
174.47
172.58
171.95
79.42
72.23
53.32
52.58
51.64
30.95
25.74
20.86
210
200
190
180
170
160
150
140
130
120
110
100
90
80
70
60
50
40
30
20
10
0
-10
f1 (ppm)

**γ-glutamyl-(*R*)-pent-4-yn-2-amine (11)**

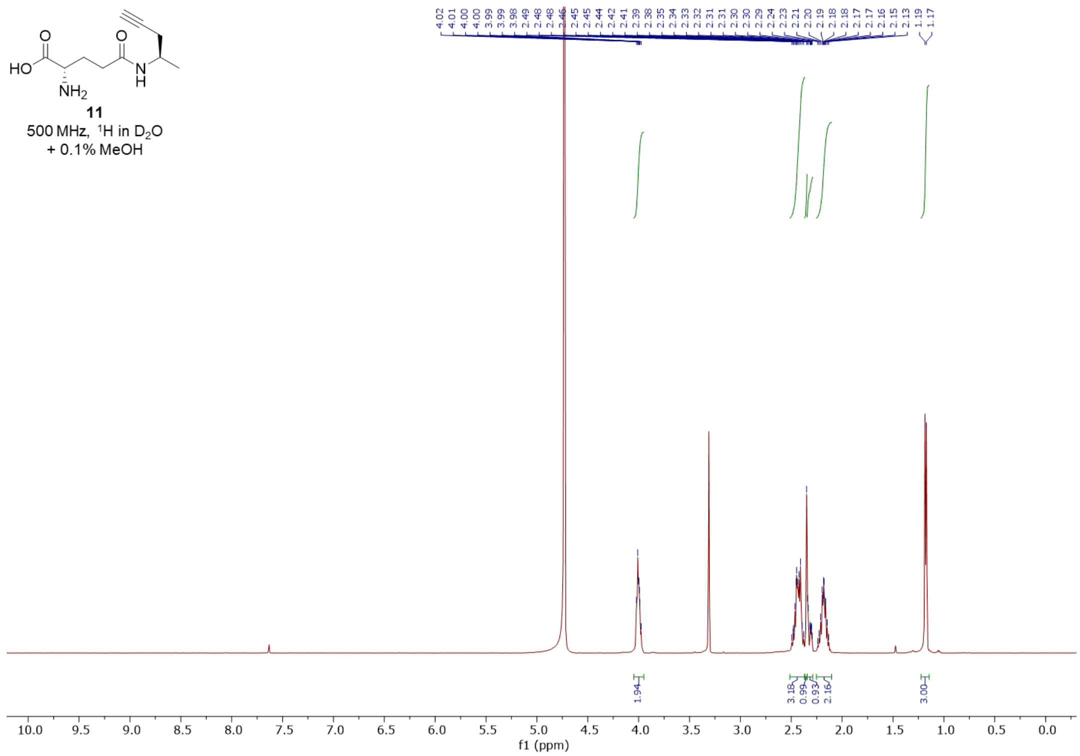


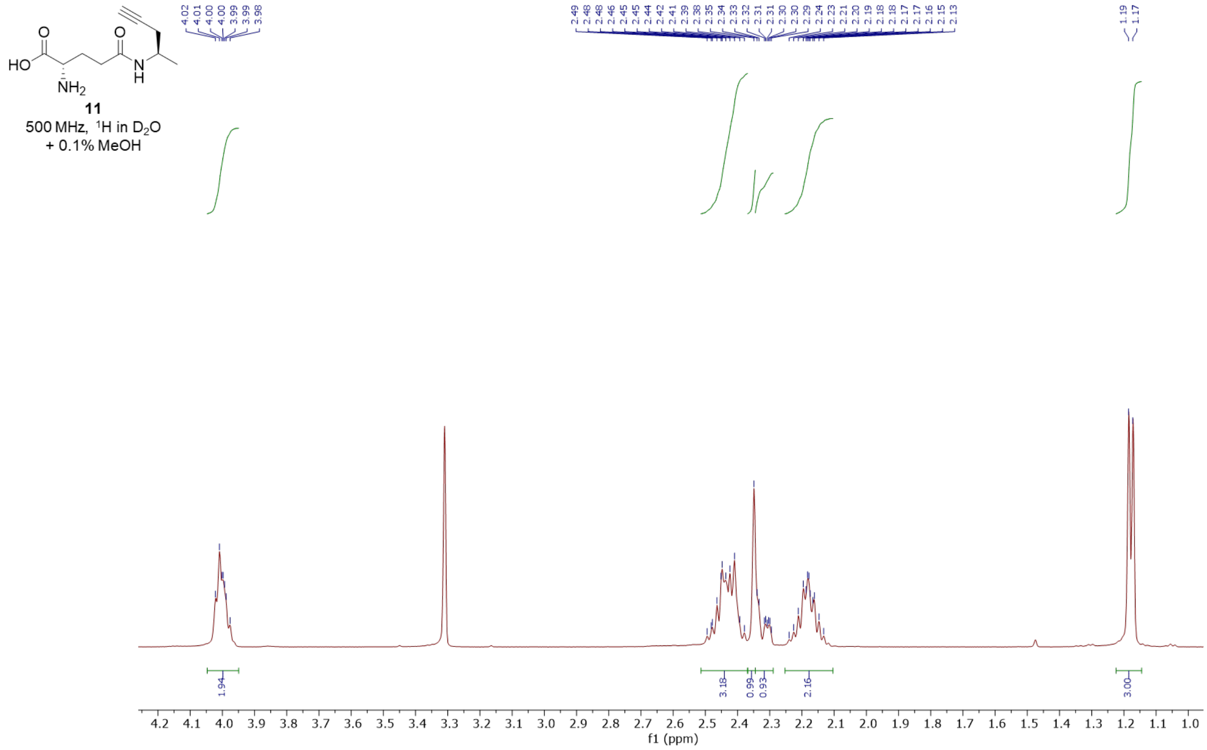

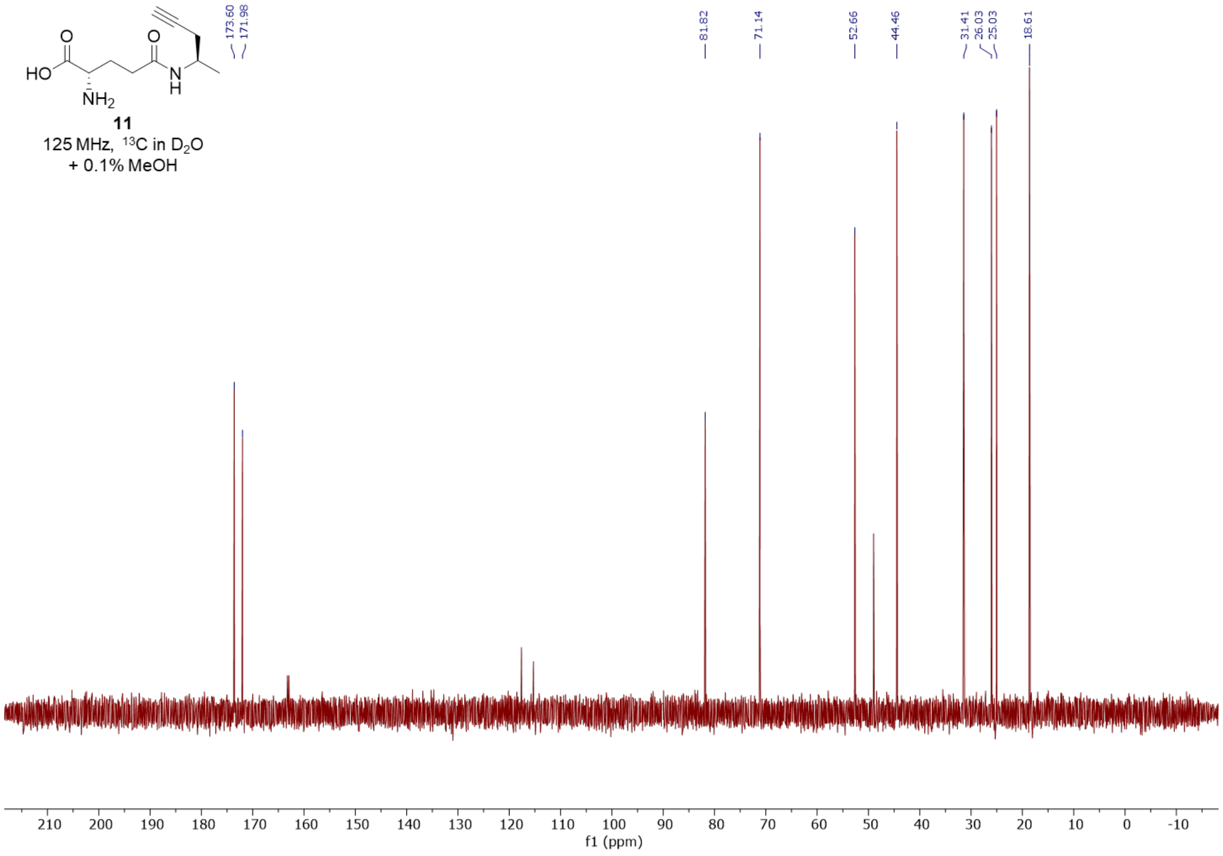
HO
O
O
NH2
N
H
11
125 MHz, 13C in D2O
+ 0.1% MeOH
173.60
171.98
81.82
71.14
52.66
44.46
31.41
26.03
25.03
18.61
210
200
190
180
170
160
150
140
130
120
110
100
90
80
70
60
50
40
30
20
10
0
-10
f1 (ppm)

**γ-glutamyl-3-butyn-1-amine (12)**

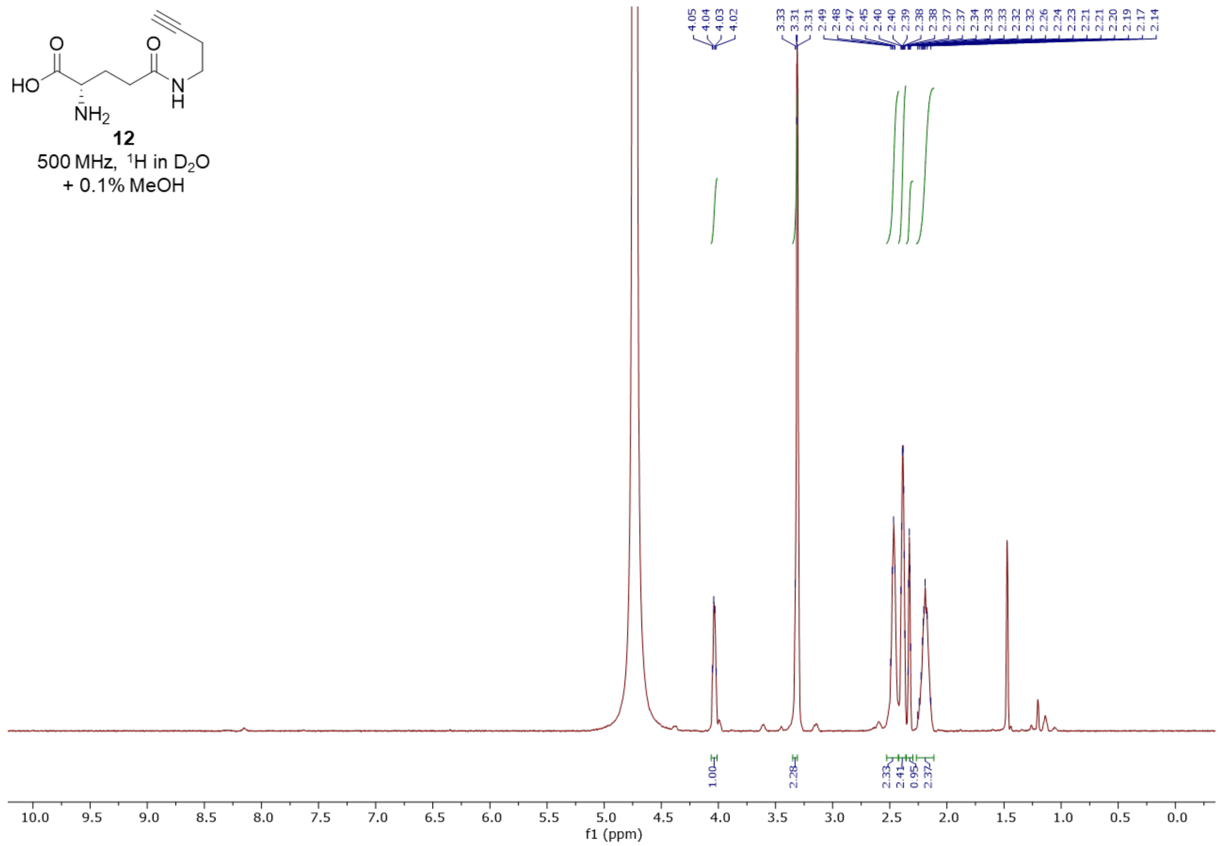


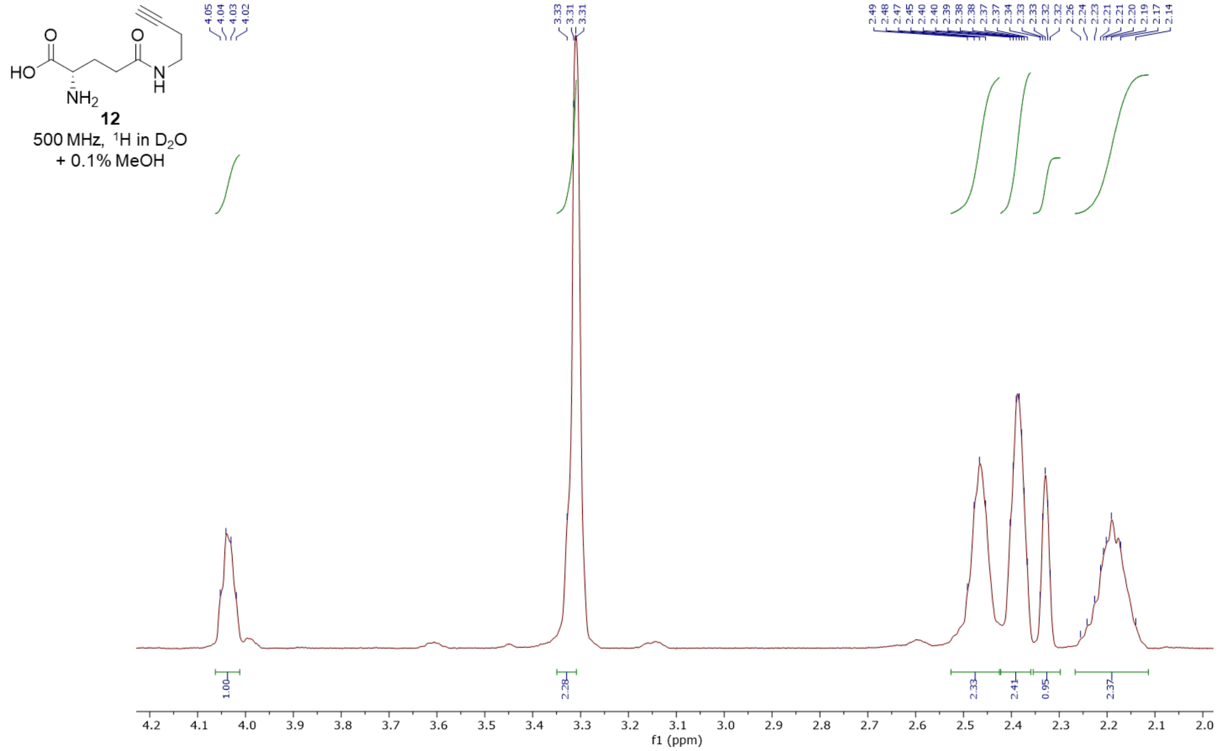

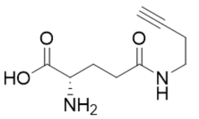

**12**

125 MHz, $^{13}C$ in $D_2O$

+ 0.1% MeOH

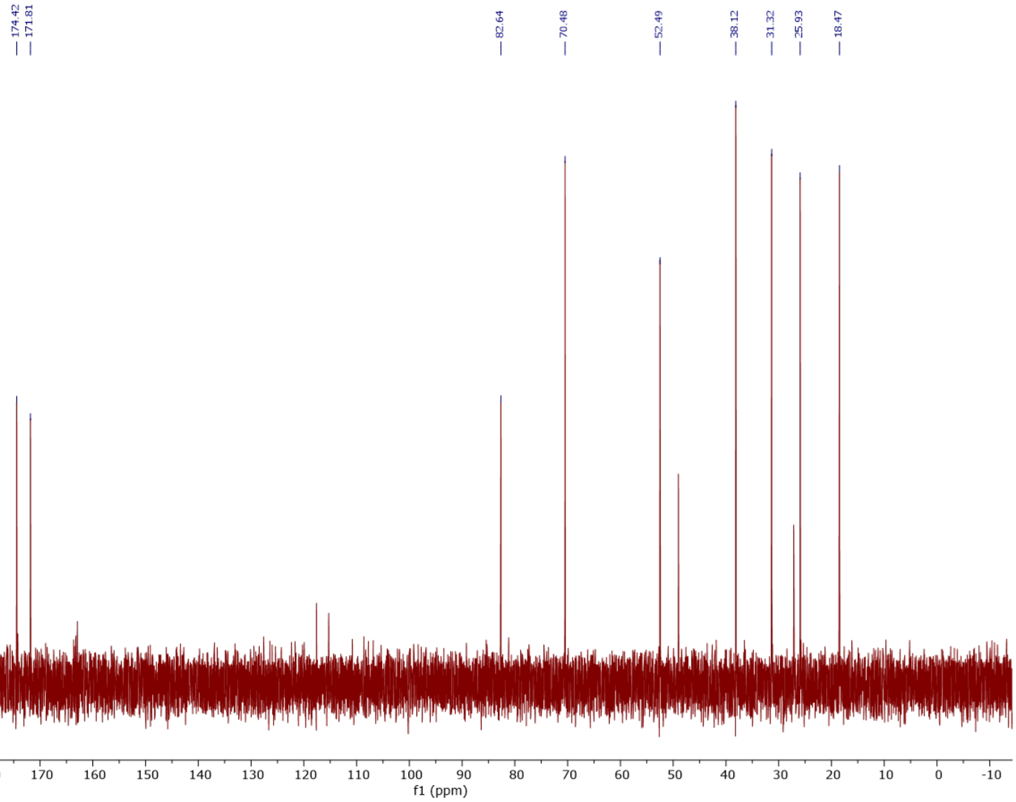


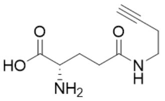

**12**

500 MHz, HSQC in $D_2O$

+ 0.1% MeOH

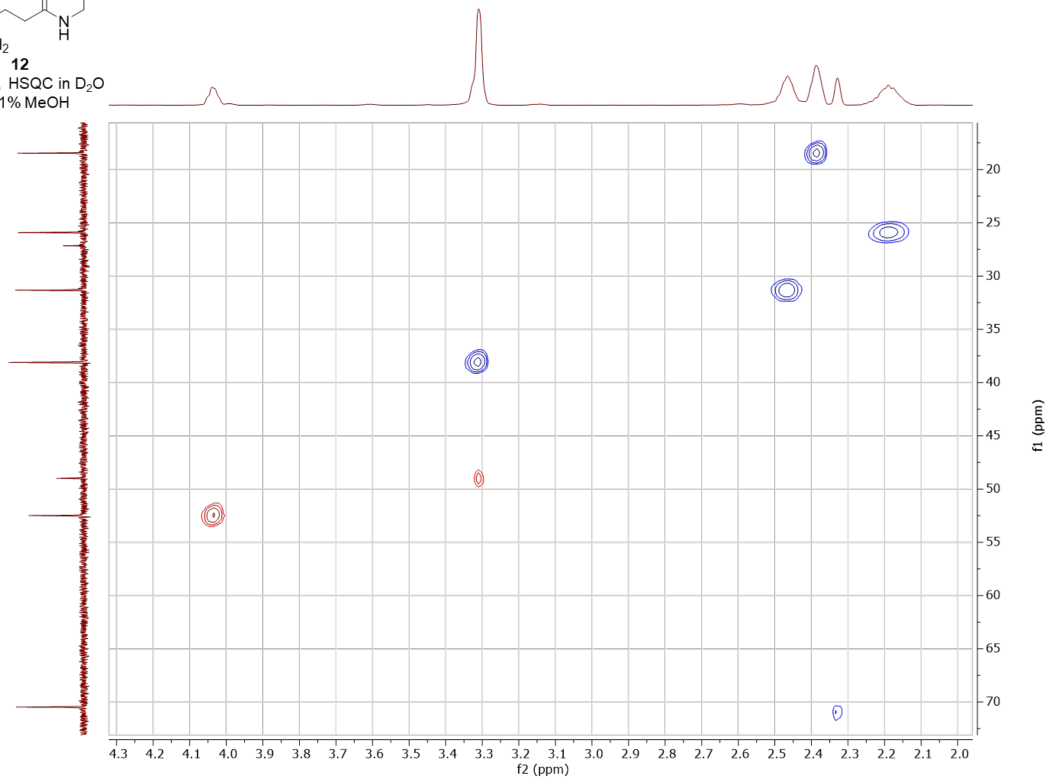

#### γ-glutamyl-L-propargylglycine-α-amide (13)

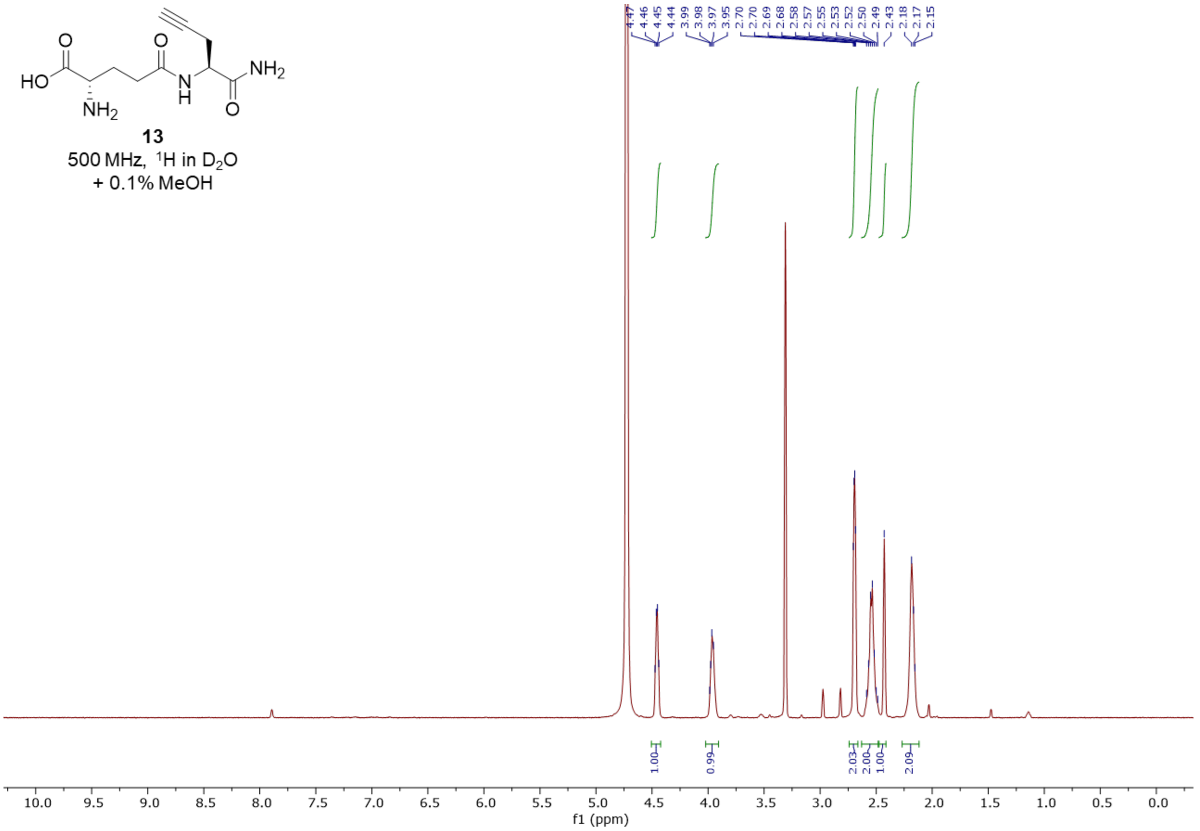


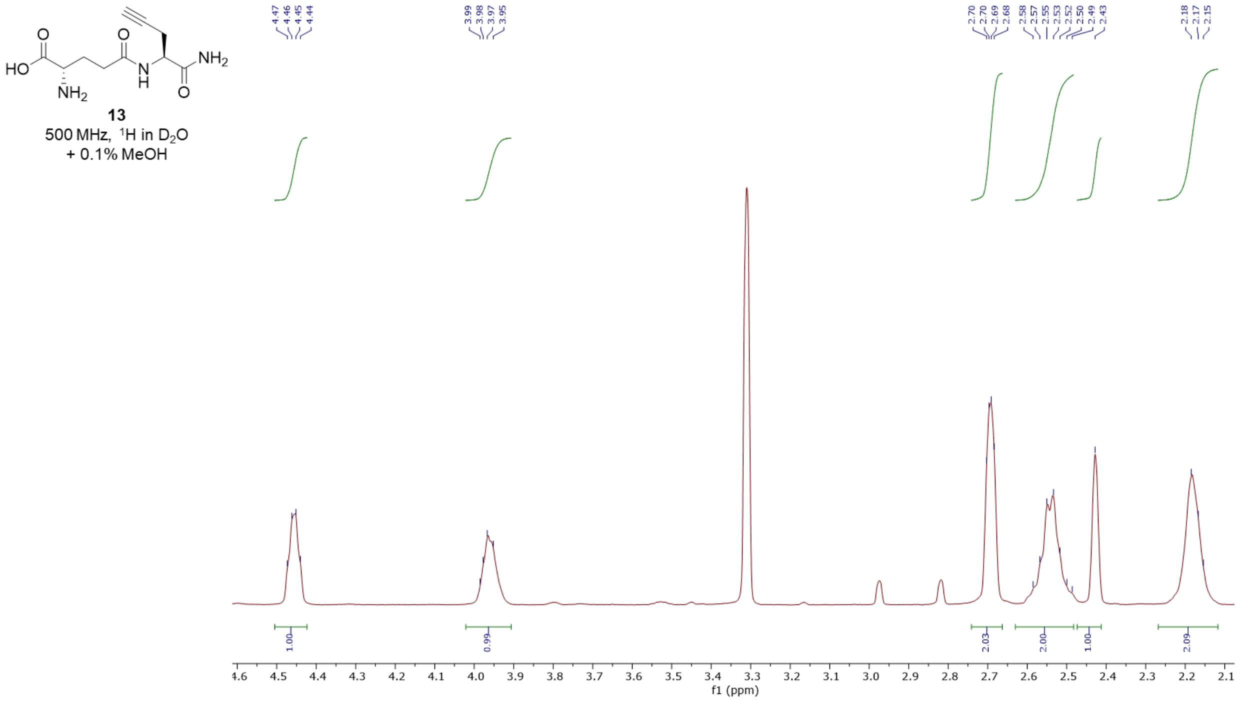

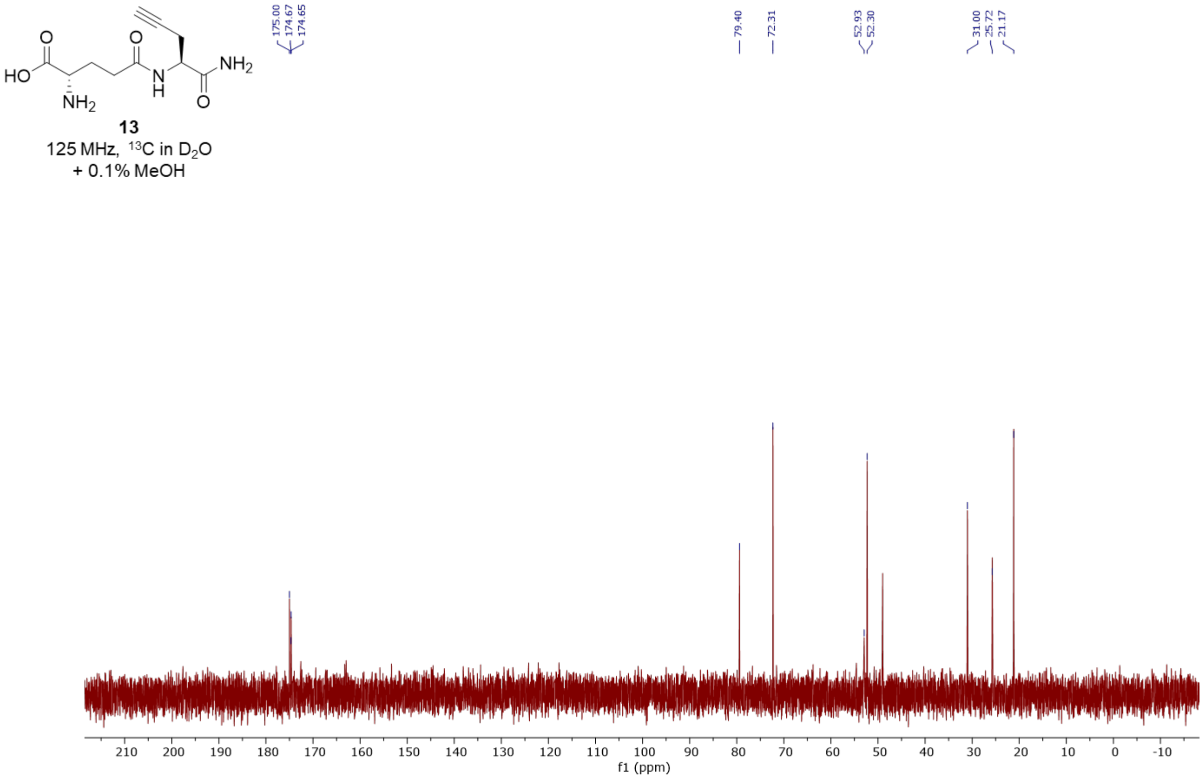

HO
NH2
N
H
NH2
O
13
125 MHz, 13C in D2O
+ 0.1% MeOH
175.00
174.67
174.65
79.40
72.31
52.93
52.30
31.00
25.72
21.17
210 200 190 180 170 160 150 140 130 120 110 100 90 80 70 60 50 40 30 20 10 0 -10
f1 (ppm)

#### γ-aminobutyryl-L-propargylglycine (14)

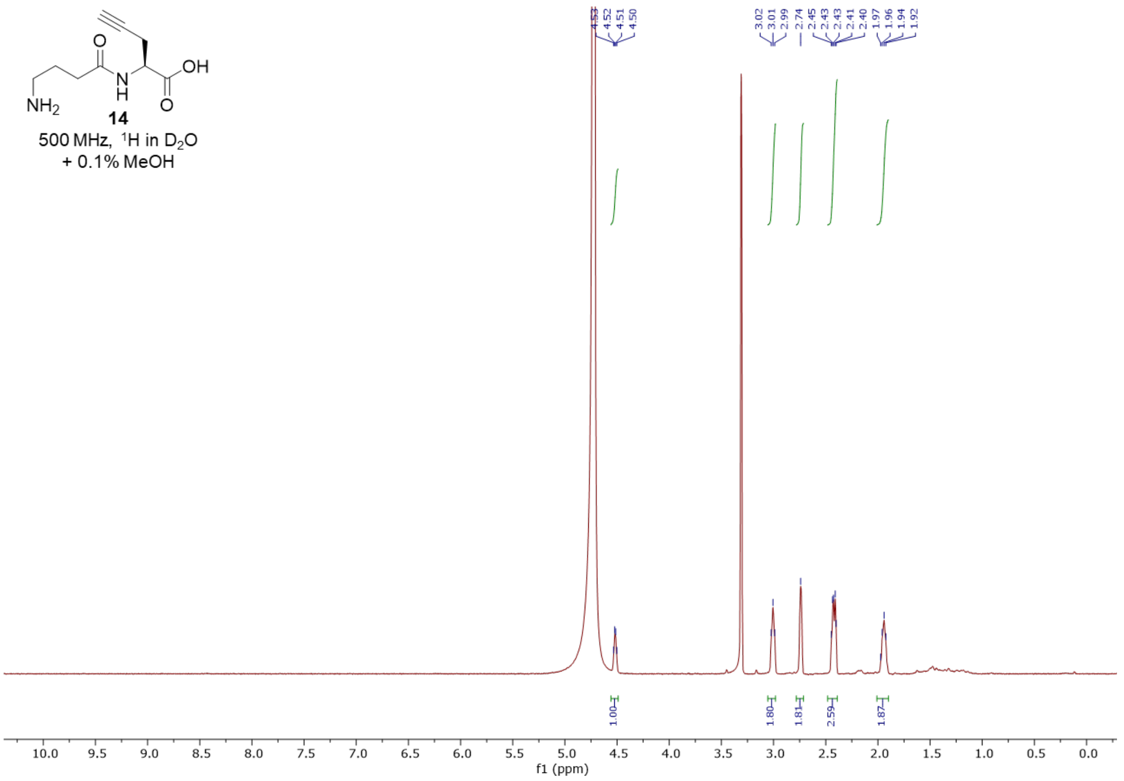


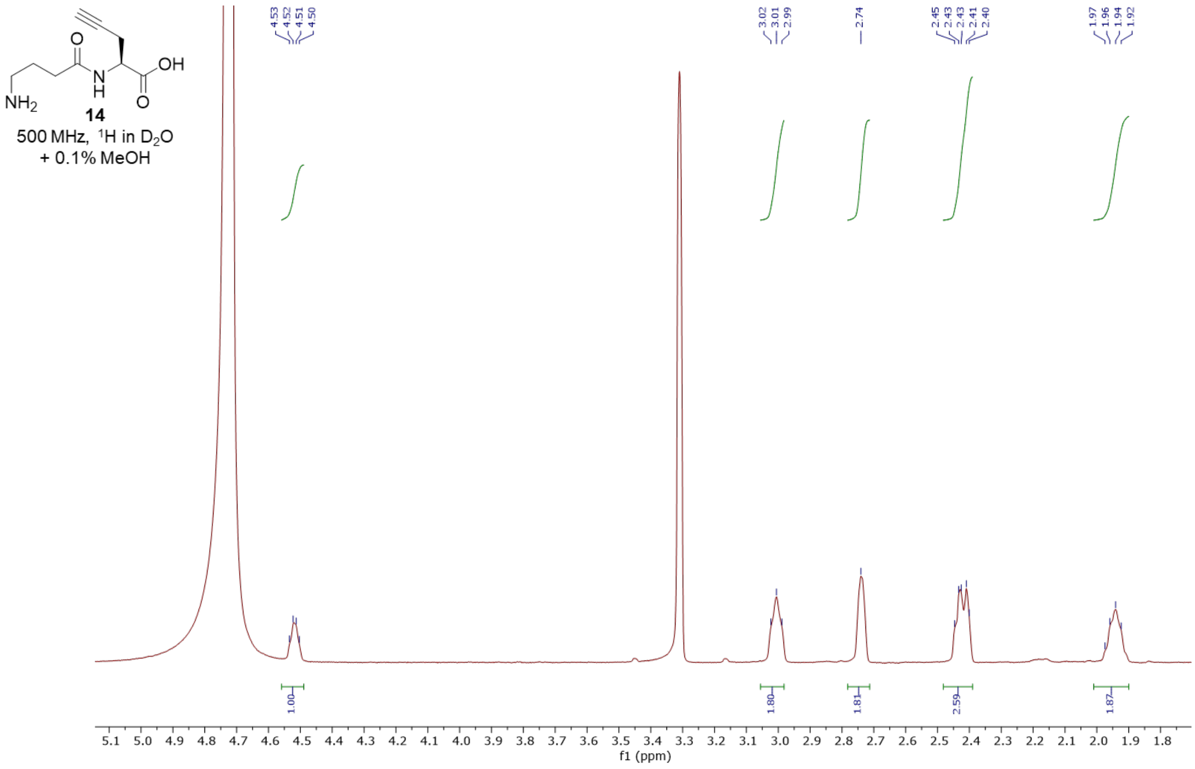

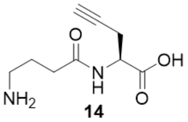


**14**

125 MHz, $^{13}C$ in $D_2O$
+ 0.1% MeOH

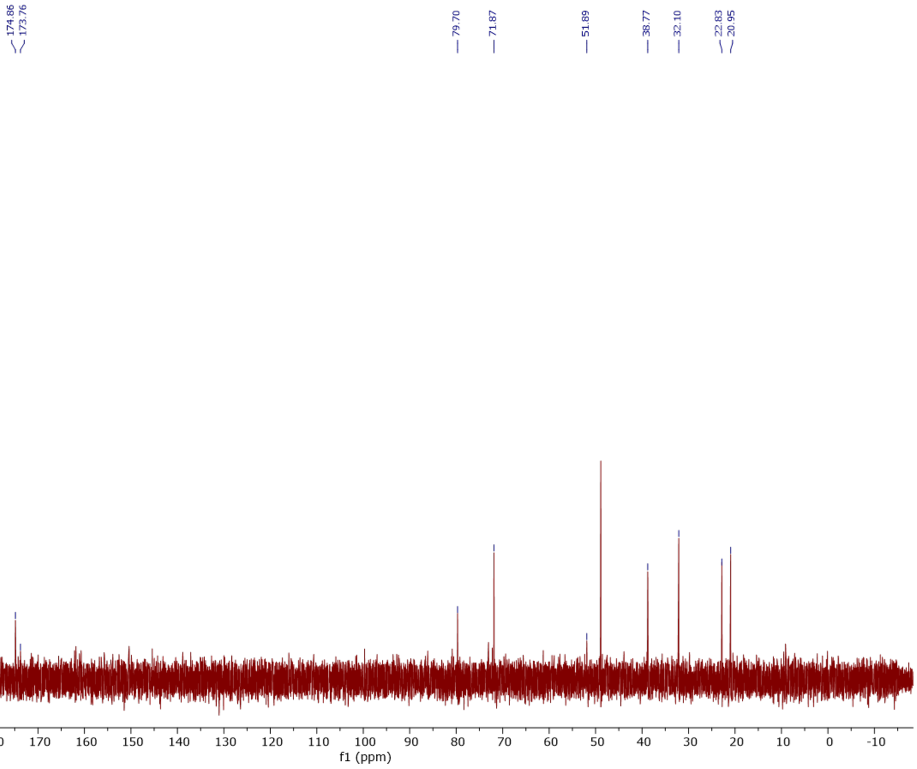

#### γ-glutaryl-L-propargylglycine (15)

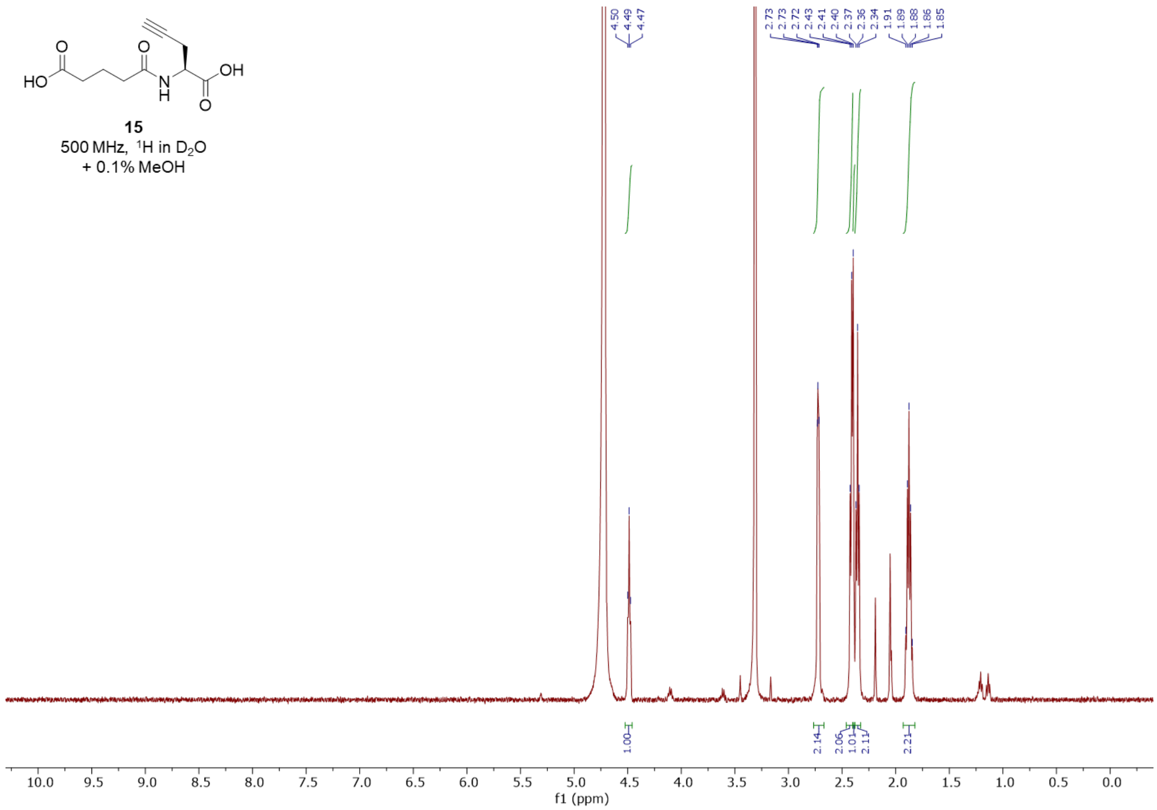


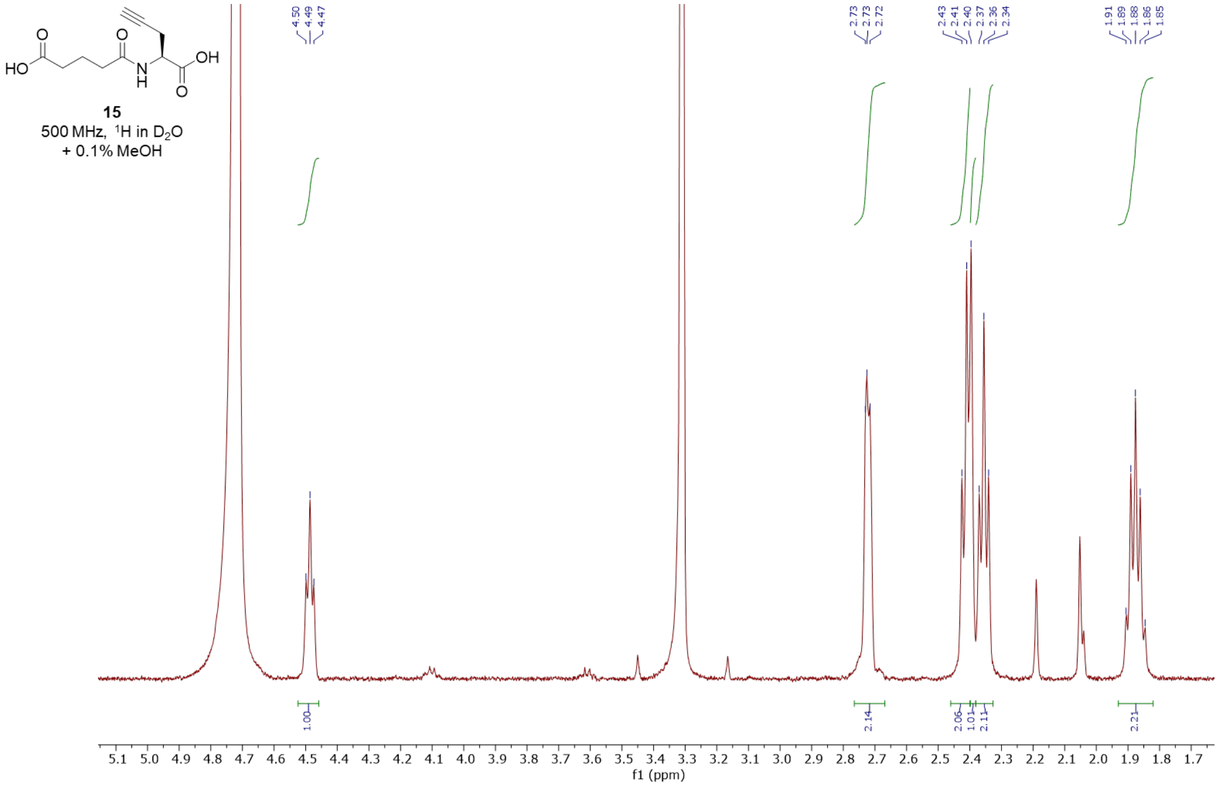

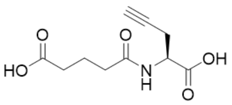

**15**

125 MHz, $^{13}C$ in $D_2O$

+ 0.1% MeOH

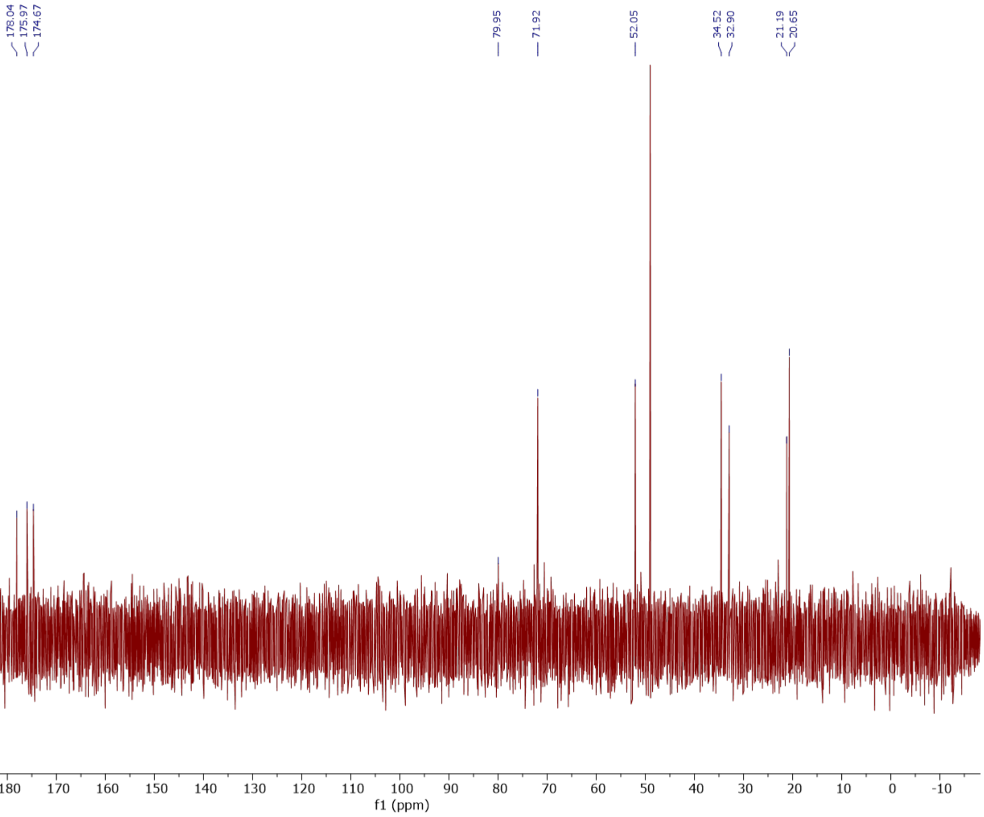

#### ***N*-butanoyl-L-propargylglycine (16)**

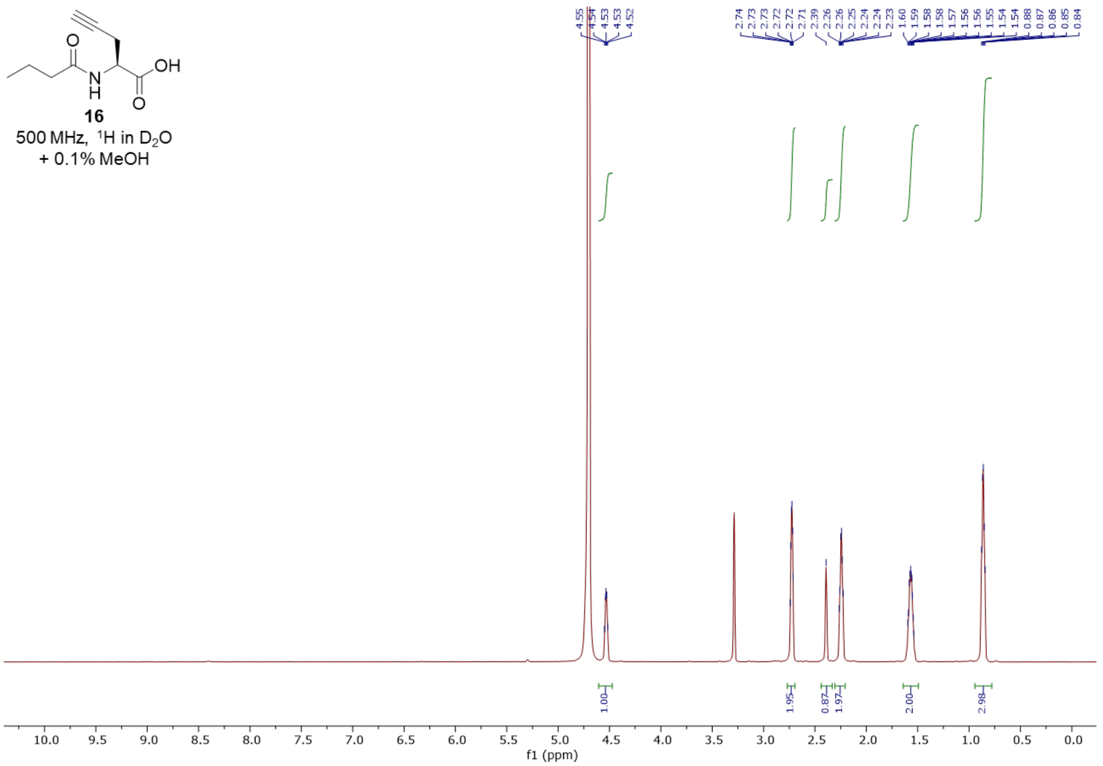


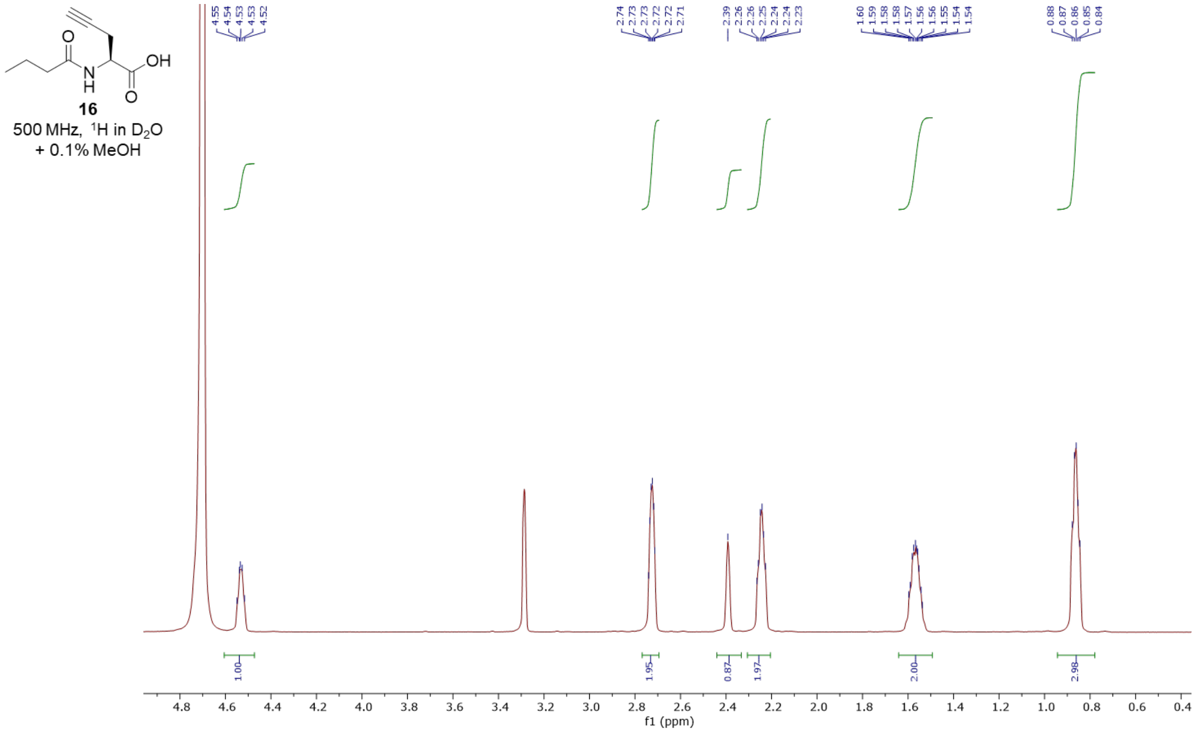

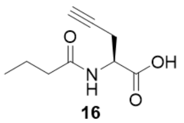

**16**

125 MHz, $^{13}$C in $D_2O$
+ 0.1% MeOH

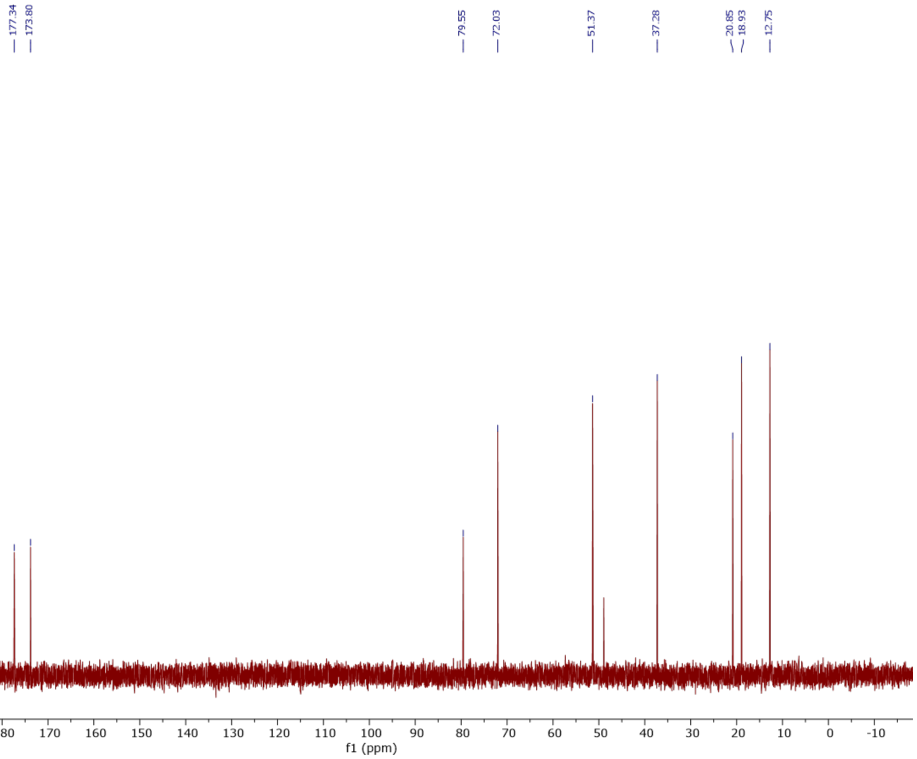

#### β-aspartyl-L-propargylglycine (17)

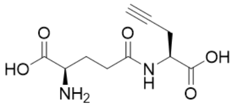

**17**
500 MHz, ¹H in $D_2O$
+ 0.1% MeOH

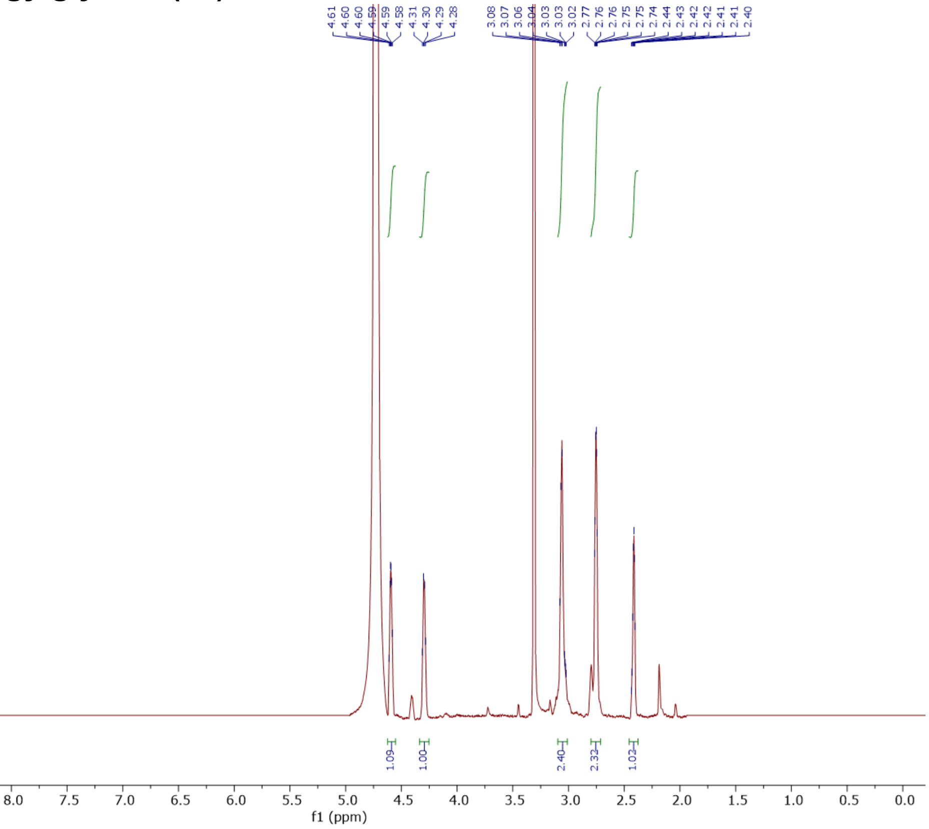


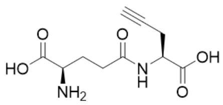

**17**
500 MHz, ¹H in $D_2O$
+ 0.1% MeOH

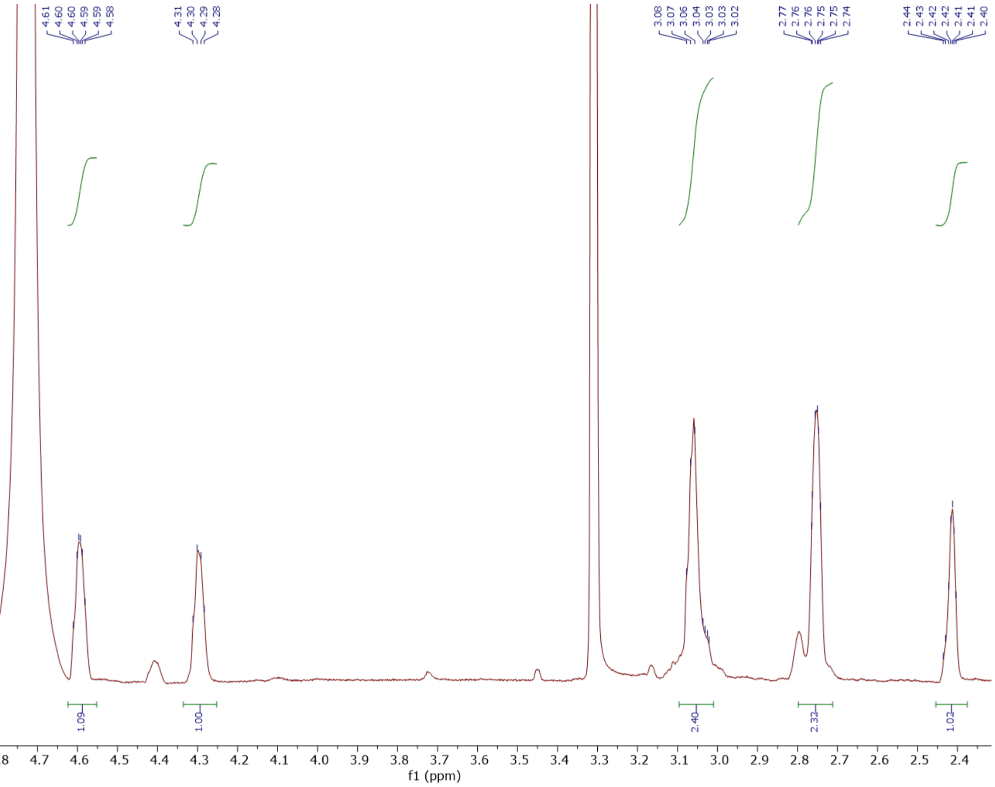

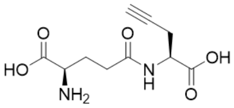

**17**
125 MHz, $^{13}C$ in $D_2O$
+ 0.1% MeOH

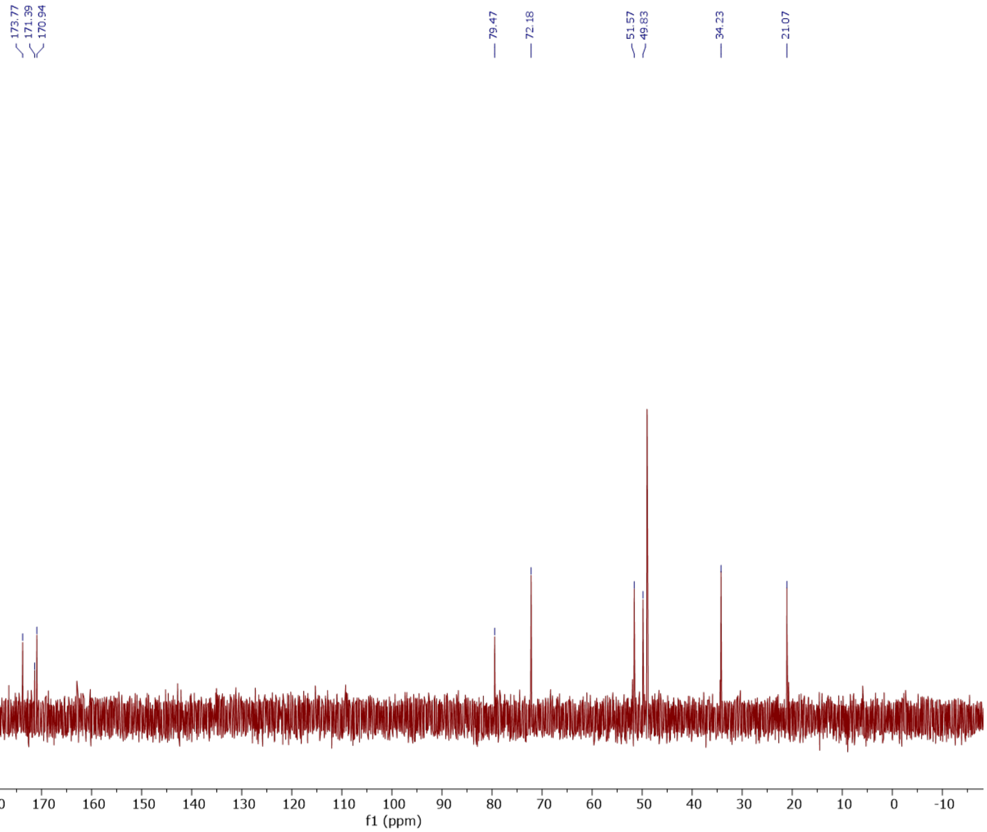

#### γ-D-glutamyl-L-propargylglycine (18)

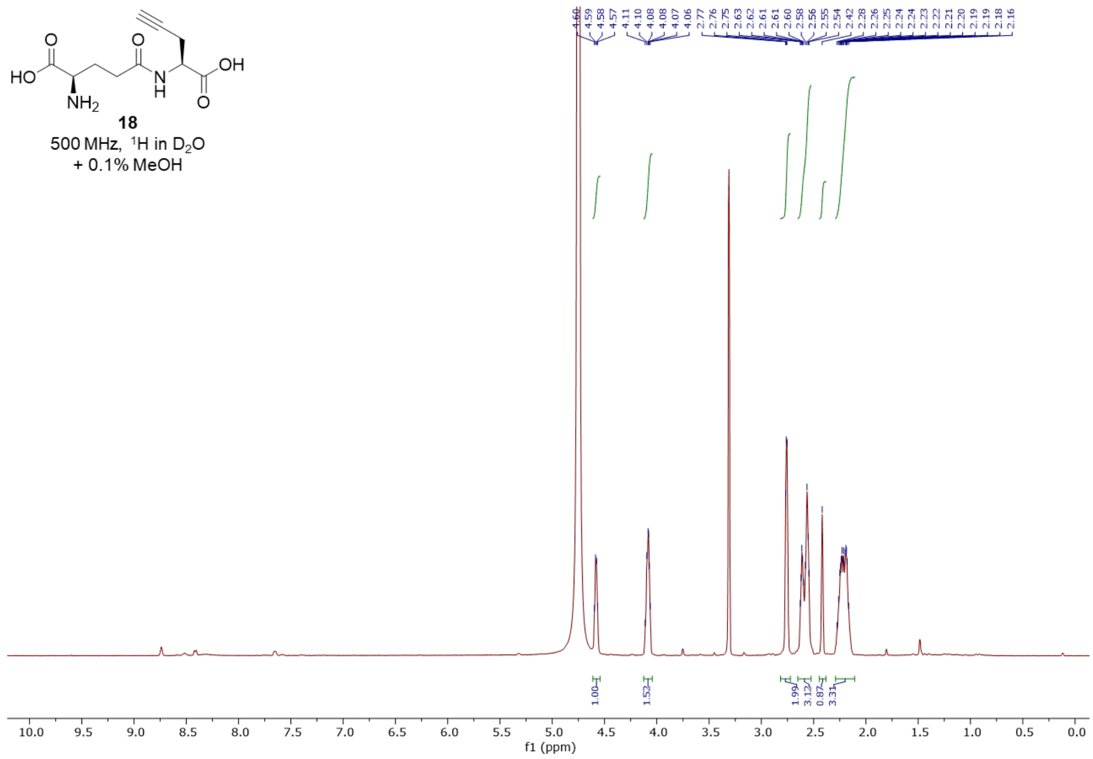


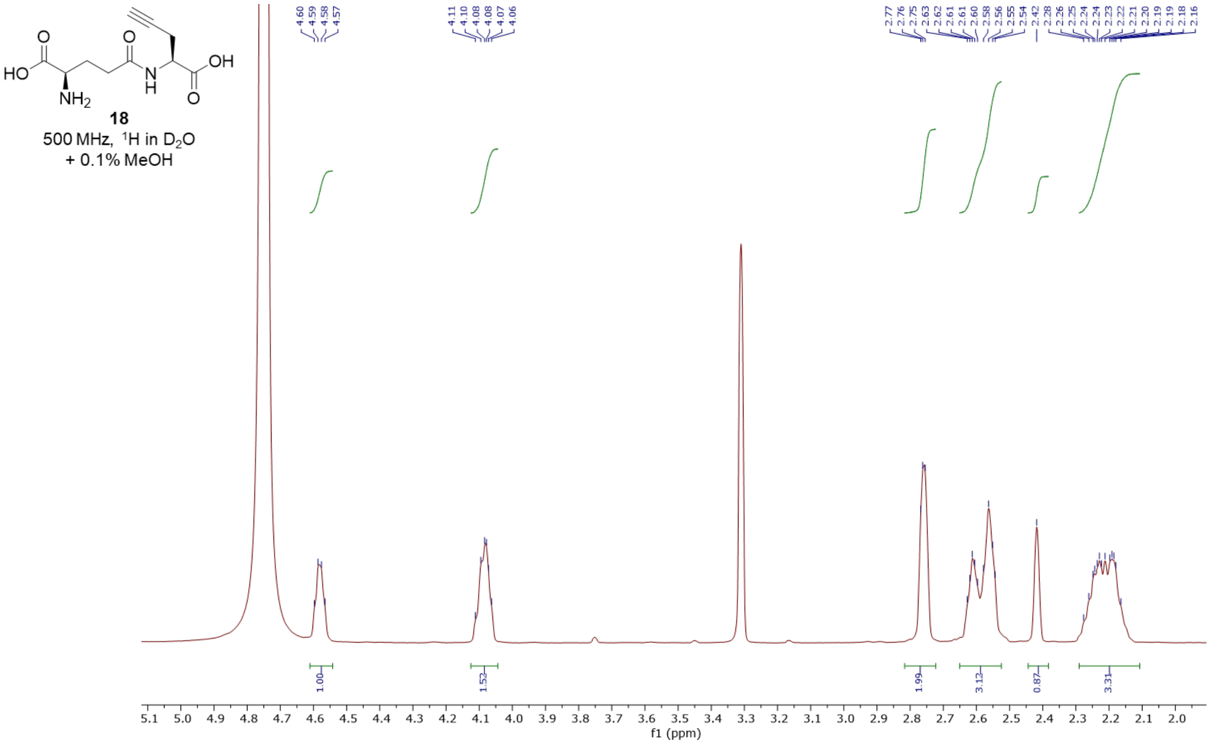

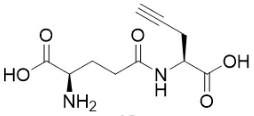

**18**
125 MHz, $^{13}C$ in $D_2O$
+ 0.1% MeOH

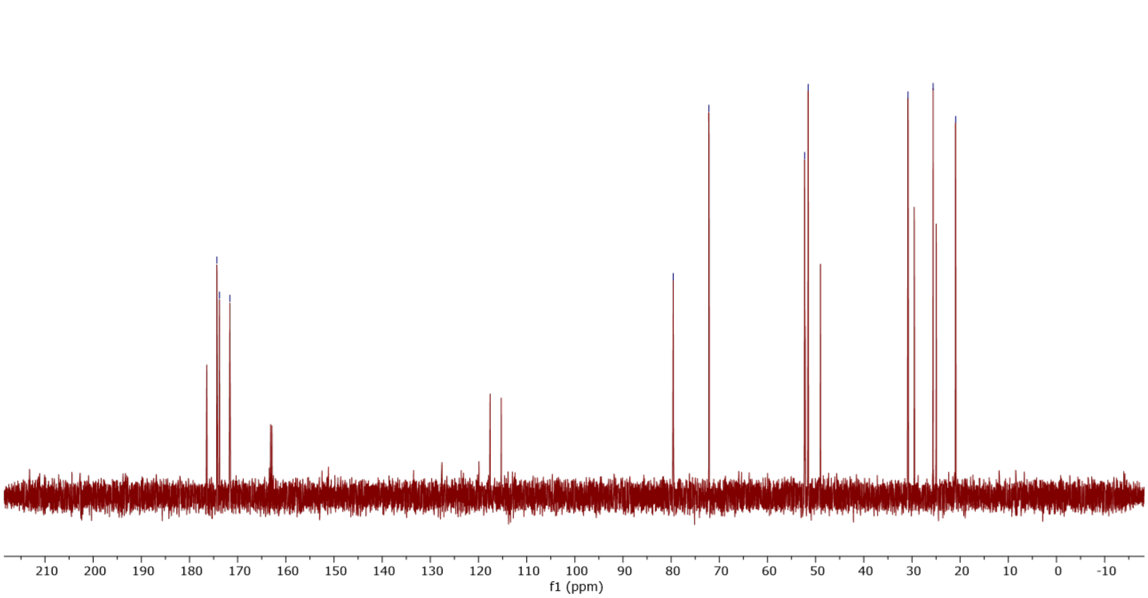

#### References


[1] J. Jumper, R. Evans, A. Pritzel, T. Green, M. Figurnov, O. Ronneberger, K. Tunyasuvunakool, R. Bates, A. Žídek, A. Potapenko, A. Bridgland, C. Meyer, S. A. A. Kohl, A. J. Ballard, A. Cowie, B. Romera-Paredes, S. Nikolov, R. Jain, J. Adler, T. Back, S. Petersen, D. Reiman, E. Clancy, M. Zielinski, M. Steinegger, M. Pacholska, T. Berghammer, S. Bodenstein, D. Silver, O. Vinyals, A. W. Senior, K. Kavukcuoglu, P. Kohli, D. Hassabis. Highly accurate protein structure prediction with AlphaFold. *Nature* **2021**, *596*, 583–589.

[2] J. R. Rachele. The Methyl Esterification of Amino Acids with 2,2-Dimethoxypropane and Aqueous Hydrogen Chloride. *J. Org. Chem.* **1963**, *28*, 2898–2898.

[3] T. Rehm, V. Stepanenko, X. Zhang, F. Würthner, F. Gröhn, K. Klein, C. Schmuck. A New Type of Soft Vesicle-Forming Molecule: An Amino Acid Derived Guanidiniocarbonyl Pyrrole Carboxylate Zwitterion. *Org. Lett.* **2008**, *10*, 1469–1472.

[4] W.-W. Zhang, T.-T. Gao, L.-J. Xu, B.-J. Li. Macrolactonization of Alkynyl Alcohol through Rh(I)/Yb(III) Catalysis. *Org. Lett.* **2018**, *20*, 6534–6538.

[5] J. C. Maza, J. R. McKenna, B. K. Raliski, M. T. Freedman, D. D. Young. Synthesis and Incorporation of Unnatural Amino Acids To Probe and Optimize Protein Bioconjugations. *Bioconjugate Chem.* **2015**, *26*, 1884–1889.
